# Complex population structure and haplotype patterns in Western Europe honey bee from sequencing a large panel of haploid drones

**DOI:** 10.1101/2021.09.20.460798

**Authors:** David Wragg, Sonia E. Eynard, Benjamin Basso, Kamila Canale-Tabet, Emmanuelle Labarthe, Olivier Bouchez, Kaspar Bienefeld, Małgorzata Bieńkowska, Cecilia Costa, Aleš Gregorc, Per Kryger, Melanie Parejo, M. Alice Pinto, Jean-Pierre Bidanel, Bertrand Servin, Yves Le Conte, Alain Vignal

## Abstract

Honey bee subspecies originate from specific geographic areas in Africa, Europe and the Middle East. The interest of beekeepers in specific phenotypes has led them to import subspecies to regions outside of their original range. The resulting admixture complicates population genetics analyses and population stratification can be a major problem for association studies. As a typical example, the case of the French population is studied here. We sequenced 870 haploid drones for SNP detection and identified nine genetic backgrounds in 629 samples. Five correspond to subspecies, two to isolated populations and two to human-mediated population management. We also highlight several large haplotype blocks, some of which coincide with the position of centromeres. The largest is 3.6 Mb long on chromosome 11, representing 1.6 % of the genome and has two major haplotypes, corresponding to the two dominant genetic backgrounds identified.

## Introduction

The honey bee *Apis mellifera* comprises more than 30 subspecies, each of which defined according to morphological, behavioural, physiological and ecological characteristics suited to their local habitat [1–4]. European subspecies broadly group into two evolutionary lineages representing on one side western and northern Europe (M lineage), and on the other eastern and southern Europe (C lineage) [1]. The two European M lineage subspecies are the Dark European or ’black’ honey bee *A. m. mellifera* and the Iberian honey bee *A. m. iberiensis*, while the C lineage subspecies include amongst others, the Italian honey bee *A. m. ligustica* and the Carniolan honey bee *A. m. carnica* [4]. Prior to the involvement of apiarists, the Alps are thought to have presented a natural barrier between *A. m. mellifera* to the north, *A. m. carnica* to the southeast, and *A. m. ligustica* to the southwest [5]. Before the turn of the 19^th^ century, French honey bee populations were solely represented by the native *A. m. mellifera*, for which regional ecotypes have previously been described [6, 7]. However, during the 20^th^ century much interest arose amongst apiarists in developing hybrids between the endemic *A. m. mellifera* and other subspecies including *A. m. ligustica*, *A. m. carnica* and the Caucasian *A. m. caucasia* from Georgia [1,8,9]. Apiarists found the hybrids to perform better with regards to the production of honey and royal jelly than the native *A. m. mellifera*, spurring further interest in these subspecies which were also reported to be more docile and easier to manage [1]. *A. m. ligustica* is a very popular subspecies worldwide amongst apiarists because of its adaptability to a wide range of climatic conditions, its ability to store large quantities of honey without swarming, and its docile nature if disturbed [10]. *A. m. ligustica* queens are also frequently exported worldwide, and most of the honey bees imported during the last centuries into the New World were also of Italian origin [10, 11]. Apiculture involving *A. m. carnica*, is also very popular among apiarists [12]. *A. m. carnica* became increasingly popular for further selection throughout central and western Europe [13, 14] on account of their calm temperament and higher honey yield compared to *A. m. mellifera* [1], to the point where *A. carnica* almost replaced entirely *A.m. mellifera* in Germany [15]. *A. m. caucasia* is a subspecies that was also imported to France, to generate *A. m. ligustica* x *A. m. caucasia* hybrids, that were themselves crossed naturally to the *A. m. mellifera* present in the local environment. Another popular hybrid used in apiculture is the so-called Buckfast, created and bred by Brother Adam of Buckfast Abbey in England [16]. Following the extensive imports of queens from “exotic” subspecies, the genetic makeup of honey bee populations in France became complex, and the genetic pollution of local populations followed with clear phenotypic consequences such as changes in the colour of the cuticle [17]. The increasing admixture of divergent honey bee subspecies has fostered conservationists to protect the native genetic diversity of regional ecotypes, such as *A. m. iberiensis* in Spain and Portugal, *A. m. ligustica* and *A. m. siciliana* in Italy (Fontana et al., 2018), and *A. m. mellifera* in France, Scotland and Switzerland amongst other places [18–22]. As a result of the different breeding practices, the necessity for a study targeted towards *A. m. mellifera* conservatories and French bee breeders specialized in rearing and selling queens arose and in this context the genomic diversity project “SeqApiPop” emerged. Within this project, samples from French conservatories, from individual French breeders and breeder organisations were analysed, including Buckfast samples. Traditionally, such wide diversity studies have been performed using a small number of molecular markers such as microsatellites [23] or limited sets of single-nucleotide polymorphisms (SNPs) [24–27], enabling population stratification, introgression and admixture levels to be characterized. However, to understand complex population admixture events, as has happened for the managed honey bee populations in France and elsewhere, or to identify signatures of natural [25,28–30] or artificial [31, 32] selection in the genome, a much higher density of markers is required. As no high-density SNP chip was available for honey bee at the onset of the project, and as the honey bee genome is very small compared to most animal genomes, being only 226.5 Mb long [33], we employed a whole-genome sequencing approach [28, 34]. Although the sequencing of honey bee workers has proved successful for detecting selection signatures or admixture events [28,34–36], analysing haploid drones allows to sequence at a lower depth and with greater accuracy in variant detection [20,29,31]. An additional advantage of sequencing haploids is that the alleles are phased, which is invaluable for studies investigating genome dynamics such as recombination hotspots and haplotype structure. Although some insights into recombination patterns in the honey bee have been made through the analysis of drones from individual colonies [37, 38] and linkage disequilibrium (LD)-based approaches [39, 40], a deep understanding of the recombination landscape, essential for fine-scale genetic analyses, requires hundreds of phased genomes. Such ‘HapMap’ projects have been conducted in humans and cattle, initially using SNP arrays [41, 42] and more recently by whole-genome sequencing as in the “1000 genome” projects [43, 44].

Therefore, as a first step towards a deep understanding of French and Western European managed honey bee populations and of their genome dynamics, we undertook the sequencing of a large dataset of haploid drones. This data comprised samples from French conservatories and commercial breeders in addition to samples from several European countries each representing potentially pure *A. m. ligustica*, *A. m. carnica, A. m. mellifera* and *A. m. caucasia* populations typically imported by French breeders. Finally, *A. m. iberiensis*, the Iberian subspecies only separated from the native French *A. m. mellifera* by the natural barrier of the Pyrenees was also studied. In total, 870 samples were sequenced for SNP detection and 629 were used for a detailed genetic analysis of present-day honey bee populations in France.

## Methods

### Sampling and sequencing

For the population genomics analyses, one individual drone per colony was sampled before emergence, from colonies throughout France, Spain, Germany, Switzerland, Italy, the UK, Slovenia, Poland, Denmark, China and from a French beekeeper having imported queens from Georgia, amounting to a total of 642 samples (Supplementary figure 1). To improve the robustness of the primary SNP detection and filtering steps, a further 30 “duplicate” samples were collected from colonies already samples for this study, in addition to 198 samples of similar genetic backgrounds from two other ongoing projects. Thus, although 642 colonies were included for population genomics analyses, in total 870 samples were used for SNP detection (supplementary table 1).

DNA was extracted from the thorax of adult bees or from pupae as described in Wragg *et al*. (2016). Briefly, drones were sampled at either the pupae/nymph or larval stage and stored in absolute ethanol at -20°C. DNA was extracted from the thorax or from diced whole larvae. Tissue fragments were first incubated 3 hours at 56° in 1 mL of a solution containing 4 M urea, 10 mM Tris-HCl pH 8, 300 mM NaCl, 1% SDS, 10 mM EDTA and 0.25 mg proteinase K, after which 0.25 mg proteinase K was added for an incubation over-night at 37°C. Four hundred μL of a saturated NaCl solution was added to the incubation, which was then gently mixed and centrifuged for 30 minutes at 15000 g. The supernatant was treated for 5 minutes at room temperature with RNAse (Qiagen) and then centrifuged again, after which the DNA in the supernatant was precipitated with absolute ethanol and re-suspended in 100 μL TE 10/0.1. Pair-end sequencing was performed on Illumina^TM^ HiSeq 2000, 2500 and 3000 sequencing machines with 20 samples per lane, or on a NovaSeq machine with 96 samples per lane, following the manufacturer’s protocols for library reparations.

### Mapping and genotype calling

Sequencing reads were mapped to the reference genome Amel_HAv3.1 [33] using BWA-MEM (v0.7.15) [45], and duplicates marked with Picard (v2.18.2;) (http://broadinstitute.github.io/picard/). Libraries that were sequenced in multiple runs were merged with Samtools (v1.8) merge [46] prior to marking duplicates. Local realignment and base quality score recalibration (BQSR) were performed using GATK (v4.1.2.) [47], using single-nucleotide polymorphisms (SNPs) called with GATK HaplotypeCaller as covariates for BQSR. Each drone was processed with the pipeline independently, and genotyped independently with HaplotypeCaller. Although the drones sequenced are haploid, variant calling was performed using a diploid model to allow the detection and removal of SNPs that called heterozygous genotypes in > 1% of samples, which might have arisen for example as a result of short-tandem repeats repeats (STRs) and could highlight copy number variants (CNVs) on the genome. Individual gVCF files were combined with CombineGVCFs, and then jointly genotyped with GenotypeGVCFs, resulting in a single VCF file for the 870 samples containing 14.990.574 raw variants. After removing Indels with GATK SelectVariants, 10.601.454 SNPs remained. Sequencing depth was estimated using Mosdepth [48]. Further details are given in supplementary file SeqApiPop_1_MappingCalling.pdf.

### Quality filters on SNPs

The first run of filters concerns technical issues related to the sequencing and alignment steps and were therefore used for the total dataset of 870 samples, to benefit from its larger size for SNP detection and validation (supplementary figure 2). These filters included (i) strand biases and mapping quality metrics (SOR ≥ 3; FS ≤ 60 and MQ ≥ 40), (ii) genotyping quality metrics (QUAL > 200 and QD < 20) and (iii) individual SNP genotyping metrics (heterozygote calls < 1%; missing genotypes < 5%, allele number < 4 and genotypes having individual GQ < 10 < 20%). Distribution and ECDF plots of values for all the filters used on the dataset were used to select thresholds and are shown in supplementary file SeqApiPop_2_VcfCleanup.pdf.

### Haplotype block detection, LD pruning, PCA, Admixture, Treemix, RFMix

Haplotype blocks were detected with Plink (v1.9) [49] using the blocks function, “--blocks no-pheno-req no-small-max-span”, with the parameter “--blocks-max-kb 5000”. LD pruning was performed with Plink using the indep-pairwise function. Principal component analyses were performed with Plink and the contribution of individual SNPs to the principal components were estimated using smartpca from the eigensoft package v7.2.1. Further details are given in supplementary file SeqApiPop_3_LDfilterAndPCAs.pdf. Admixture analysis was performed with the program Admixture v 1.3.0 [50], with values of K ranging from 2 to 16. Fifty runs were performed each time using a unique random seed. The Pong software [51] was used for aligning runs with different K values and for grouping results from runs into clustering modes, setting the similarity threshold to -s = 0.98.

Further details are given in supplementary file SeqApiPop_4_Admixture.pdf. Population migration analysis was performed with TreeMix [52], with the option for grouping SNPs set to -k=500, testing between 0 and 9 migrations and performing 100 runs per migration with a unique random seed. The optimum number of migrations was estimated with the R package OptM (Fitak, R. R.: https://github.com/cran/OptM) using the Evanno method provided [53]. Tree summaries for the 100 runs per migration tested were performed with DendroPy [54] and drawn with FigTree v1.4.4 (http://tree.bio.ed.ac.uk/software/figtree/). Further details are given in supplementary file SeqApiPop_5_TreeMix.pdf. Local ancestry inference and positioning of haplotype switches were performed with RFMix v2.03-r0 [55]. Three main genetic backgrounds were considered for this analysis, corresponding to the three major groups highlighted in the PCA analysis.

Reference samples were selected as having > 95 % pure background. Although most diploid data was removed and data is already phased, shapeit.v2.904 [56] was run to format the vcf files for RFMix.

RFMix was run using genetic maps generated from the data of Liu et al. [37]. Briefly, reads from the project SRP043350 were retrieved from Short Read Archive (SRA) (https://www.ncbi.nlm.nih.gov/sra), aligned to the reference genome for SNP detection and recombinants were detected with the custom script find_crossing_overs.py to produce a genetic map. Further details on the RFMix analysis are given in supplementary file SeqApiPop_6_RFMix.pdf.

## Results

### Sequencing and genotyping

Sequencing of the honey bee drones for the SeqApiPop diversity project began in 2014 on Illumina HiSeq instruments and some of the first samples had such low coverage that a second run (or even three in the case of OUE8) sequencing was performed. For these samples, the resulting BAM files were merged prior to variant calling. Only four samples of the diversity project were sequenced on Novaseq instruments, for which higher sequencing depths were achieved. Therefore, to improve the robustness of the SNP detection pipeline, we included drone genome sequences from other ongoing subsequent projects using the same genetic types, that were produced with Novaseq instruments. Samples sequenced with the HiSeq and NovaSeq instruments had mean sequencing depths of 12.5 ± 6.1 and 33.5 ± 10.2 respectively (Supplementary Table ST1, Supplementary figures 3 and 4). Genotyping the whole dataset of 870 drones with the GATK pipeline allowed the detection of 10,601,454 raw SNPs (supplementary figure 2). Results of the subsequent filtering steps are shown in the Venn diagrams in supplementary figures 5, 6 and 7. A total of 7,023,976 high-quality SNPs remained after filtering. The 198 samples from the other projects and 30 within-colony duplicate samples from the present diversity project were removed from the dataset for downstream analyses. Although a filter on genotyping rate ≥ 95% was applied in the primary filtering steps, the final filter on heterozygote calls was set to keep SNPs with up to 1% of heterozygote samples, and these remaining heterozygous genotypes were set to missing (supplementary figure 2). After this, a final filter on missing data in samples was applied and 15 samples were removed due to the fraction of missing genotypes exceeding 10 %. The final diversity dataset comprised 629 drones (supplementary table 1) and 7,012,891 SNPs, and was used for all subsequent analyses unless stated otherwise.

### Contribution of SNPs to the variance in PCAs: detection of large haplotype blocks

Principal component analysis was performed on the 629 samples and 7 million SNPs, results in a clear differentiation of three groups of samples. The first principal component, representing 10.8 % of the total variance, broadly differentiates M lineage bees, *A. m. mellifera* and *A. m. iberiensis*, from the *A. m. ligustica*, *A. m. carnica* and *A. m. caucasia* bees. The second principal component, representing 3.1 % of the variance, separates the O lineage *A. m. caucasia* bees from the C lineage *A. m. ligustica* and *A. m. carnica* bees (supplementary figure 8). PC3 represents 1.2 % of the variance and the remaining principal components each represent 0.7 % or less. When looking at the individual contributions of SNPs to the variance, we can see that only a very small proportion of the ∼7 million markers contribute significantly to PC1 (red lines on supplementary figure 9) and that this proportion is even much smaller for PCs 2 and 3. Two reasons for such a limited contribution to the variance of the majority of markers is the low informativity of markers of low minor allele frequency (MAF) and the redundancy of markers that are in strong linkage disequilibrium (LD). Therefore, to thin the dataset, we tested the effect of several MAF filters and chose the most pertinent one for subsequent testing of various LD pruning values. The effects of these filters were estimated by inspecting the contributions of the SNPs to the principal components. The MAF filters tested showed clearly that datasets containing only SNPs with MAF > 0.01 or MAF > 0.05 are sufficient to allow a higher proportion of markers contributing to the PCs, with a notable increase of SNPs contributing to PC2 and PC3 (supplementary figure 9). To avoid losing too many potential population-specific markers present at low frequency in the data, we chose to use the lowest MAF threshold tested, leaving a dataset of 3,285,296 SNPs having MAF > 0.01 for subsequent analyses. On inspecting the contributions of individual SNPs to principal components along the genome, a striking feature we observe is that for several large chromosomal regions, five of which being larger than 1 Mb, most SNPs make a significant contribution, with the observed values being amongst the strongest observed genome-wide (Supplementary Figures 10 and 11). Such observations suggest the existence of large haplotype blocks driving differentiation along principal components, in particular principal component 1. To explore this further we compared these genomic regions to the haplotype blocks detected with Plink (supplementary table 2) revealing significant overlap by visual inspection (Supplementary Figure 12). The largest of these blocks spans 3.6 Mb on chromosome 11, which is close to 1.6 % of the honey bee genome size, and four others on chromosomes 4, 7 and 9 are larger than 1 Mb (figure 1, supplementary figures 10, 11, 12, supplementary table 2).

**Figure 1:**
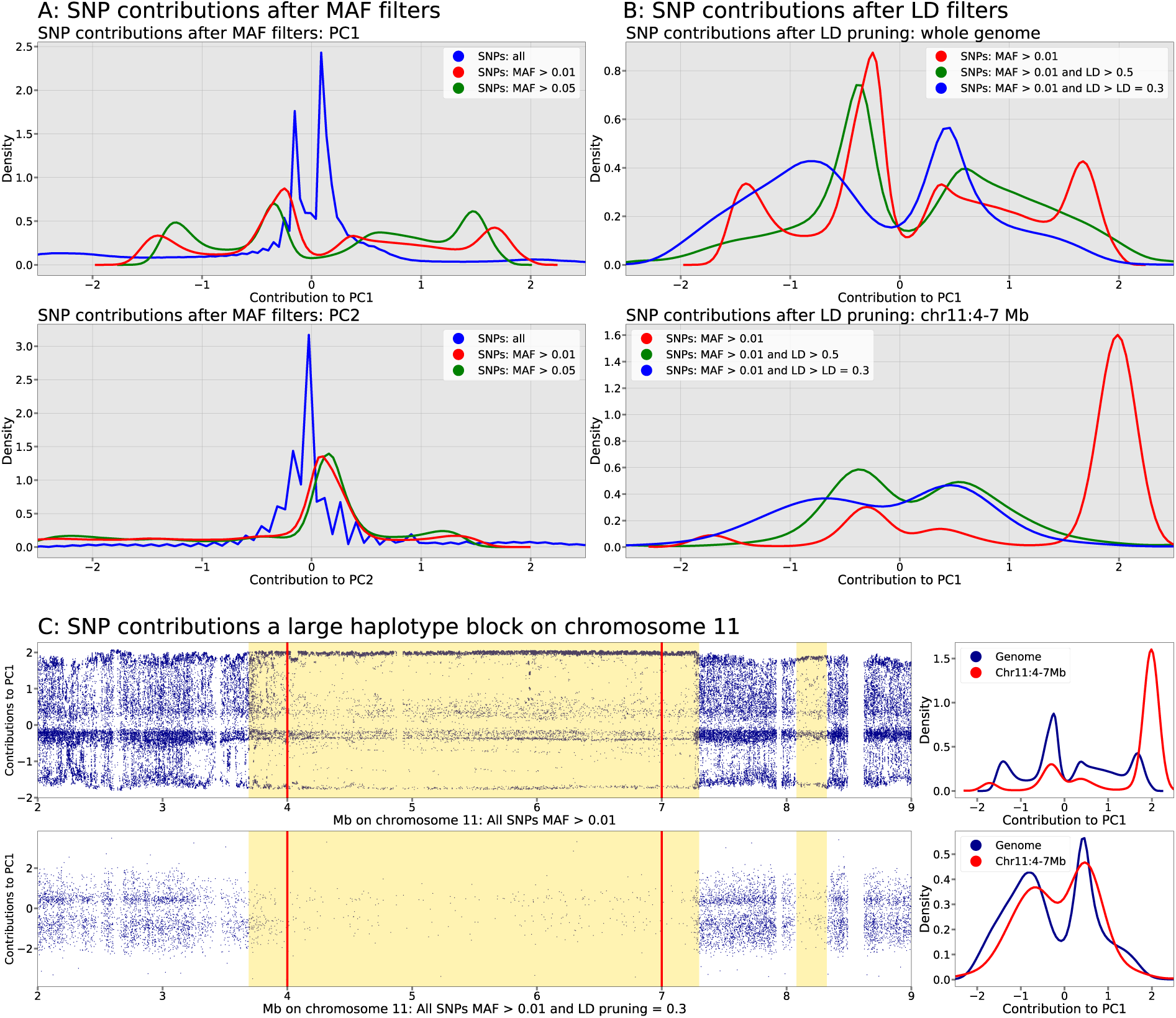
Contribution of SNPs to the principal components, MAF and LD filters and detection of large haplotype blocks. A: contribution to PC1 (top) and PC2 (bottom). When all 7 million SNPs are analysed simultaneously, the majority share a small contribution to PC1 and have no contribution to PC2 (blue). When retaining only markers with MAF > 0.01 or 0.05, (3,285,296 and 2,525418 SNPs; green and red lines, respectively), markers retained have a stronger contribution to PC1 and a higher proportion of markers contribute to PC2. B: LD pruning on the 3,285,296 SNPs with MAF > 0.01 (red line). Top: out of the 1,011,918 and 601,945 SNPs retained after pruning at LD = 0.5 (green line) or 0.3 (blue line), the distribution of the contribution of the markers is more even. Bottom: contribution of SNPs to PC1 in a 3 Mb region of chromosome 11. Almost all markers in this region show contributions are among the highest genome-wide (red line). The distribution of these contributions is improved by LD pruning (green and blue lines). C: blue points show the contribution of individual SNPs along a 6 Mb region of chromosome 11 containing two haplotype blocks of > 3 Mb and ∼200 kb (yellow backgrounds) before (top) and after (bottom) LD pruning. The LD pruning eliminates successfully the markers in the haplotype blocks and the distribution of marker approaches that of the rest of the genome, as shown in the corresponding density plots on the right.

### LD filtering

Population structure and admixture analyses rely largely on the assumption that markers along the genome are independent. Indeed, markers in strong LD such as those in haplotype blocks, can influence genetic structure. Therefore, we sought to investigate the impact of LD pruning on population structure inference. The number of SNPs used in a window for LD pruning was determined such that most windows would correspond to a physical size of 100 kb. To achieve this, we used the mode of the distribution of the number of SNPs in 100 kb bins, which is 1749 for the dataset of 3,285,296 SNPs with MAF > 0.01 (supplementary figures 13 and 14). LD pruning was thus performed with a window size of 1749 SNPs and 175 bp (10 %) overlap and various values were tested, spanning between 0.1 ≤ LD r^2^ ≤ 0.9. PCA following these various thresholds show that with LD r^2^ < 0.3 the global structure of the dataset is lost, with only one population (*A. m. iberiensis*) contributing strongly to the variance (supplementary figure 15), whereas with LD r^2^ > 0.3, the contributions to the variance in PC1 is not so widely distributed (supplementary figure 16). The effect of LD pruning on the haplotype blocks is drastic, with the few SNPs retained having a distribution of their contributions to the variance in PC1 and PC2 similar to that of the rest of the genome (supplementary figure 17). After pruning for LD r^2^ < 0.3, 601,945 SNPs were left in the dataset which were subsequently used in the analysis of population structure.

### Analysis of population structure

The PCA revealed distinct population structure within the data. For instance, some populations from French breeding organisations, such as the Royal Jelly breeder organisation (GPGR: Groupement des Producteurs de Gelée Royale), and the Corsican breeder’s organization (AOP Corse), appear quite homogenous (figure 2), with GPGR samples clustering close to the *A. m. ligustica* and *A. m. carnica* reference populations and, while AOP Corse samples appear as a distinct group between the C lineage *A. m. ligustica* and *A. m. carnica* on one side and the M lineage *A. m. mellifera* and *A. m. iberiensis* on the other. Other populations from French breeders appear much less homogenous, with individuals scattered across the whole graph, see for examples Tarn 2 on figure 2, suggesting various degrees of admixture between the three principal genetic groups (supplementary figure 18).

**Figure 2:**
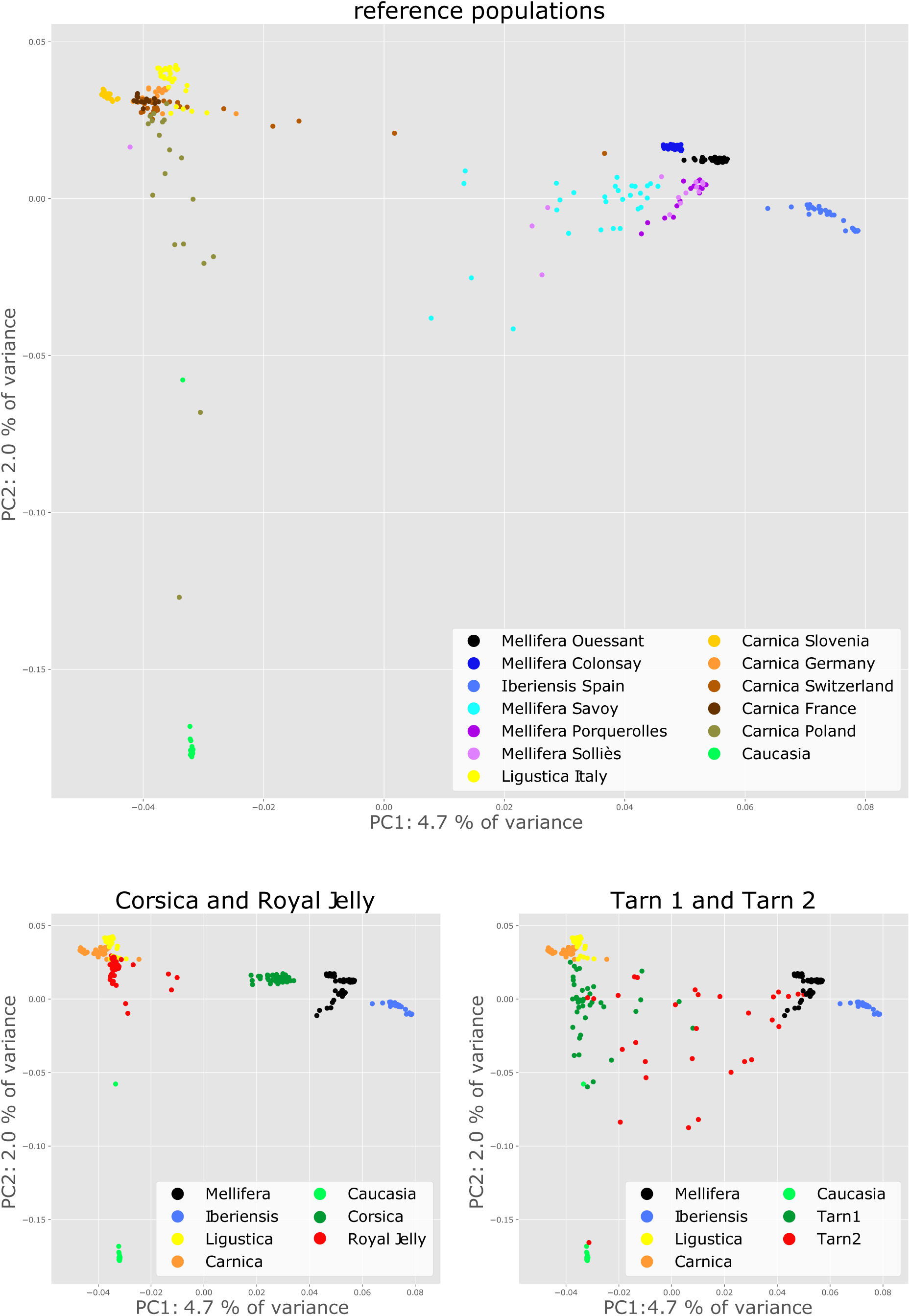
PCA on the reference populations and on a sample of representative breeder populations. The 601,945 SNPs obtained after MAF filtering and LD pruning were used. Left: reference populations only, with a colouring scheme according to their origin. Middle and left: only the reference populations with a high proportion of pure background individuals, as observed after Admixture analysis, were kept and coloured according to the five subspecies. Some breeder populations appear homogeneous, such as the honey bees selected for Royal Jelly or those from Corsica. Others are heterogeneous, such as populations Tarn1 and Tarn2, from breeders.

To further investigate the genetic structure and the effects of human-mediated breeding, we performed admixture analyses. Our dataset consists of reference samples from thirteen origins, including two islands, in addition to samples from several commercial breeders and conservatories. The genetic makeup is therefore expected to be complex and the first task was to estimate the optimal number of genetic backgrounds (K). We performed 50 independent runs with the Admixture software for each value of 2 ≤ K ≤ 16 on the LD-pruned dataset, totaling 750 independent analyses. Cross-validation (CV) error estimates of the results computed by the software are shown in figure 3A. Results suggest that the most likely number of genetic backgrounds is 8 or 9, with K = 8 having runs with the lowest CV values overall, and K = 9 having the lowest median CV value over its 50 runs. The resulting Q matrices were jointly analyzed using Pong [51], where for each value of K runs are grouped together by similarity into modes and the mode containing the largest number of similar runs is defined as the major mode. As Pong failed to find disjoint modes with the default similarity threshold of 0.97, we increased the stringency of this value to 0.98 for our analyses. Naturally, for low values of K, such as 2 or 3, most of the Q matrices are very similar and the major modes contain most runs, if not all.

**Figure 3:**
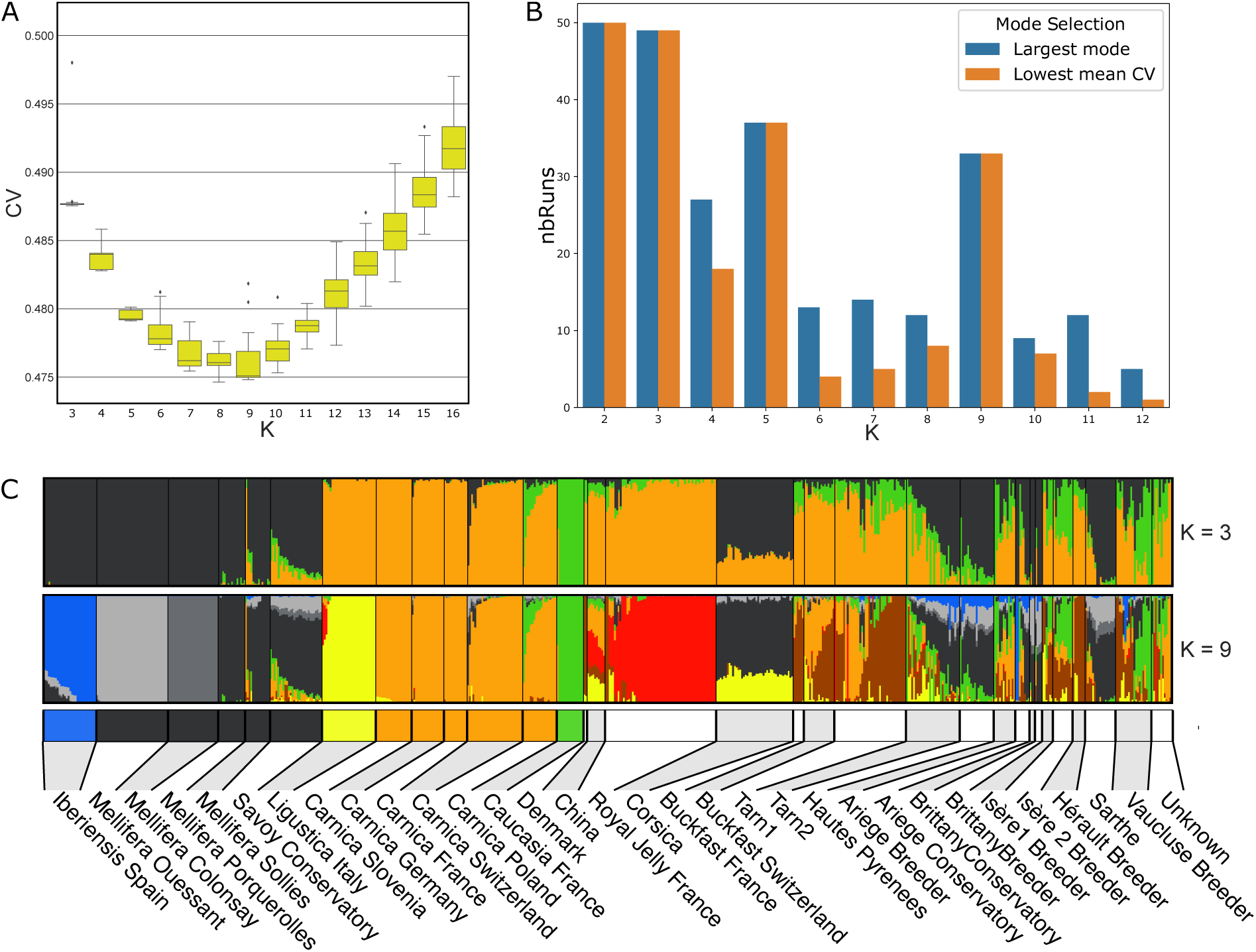
Admixture analysis. A: estimation of Cross validation error for 50 runs of Admixture for 3 ≤ K ≤ 16. B: Major modes and modes with the lowest mean cross-validation (CV) error for Admixture runs. For each value of K ranging between 2 and 12, Q matrices from Admixture runs were grouped by similarity in modes by using the Pong software (Behr et al. 2016). Blue: number of runs in the major mode; orange: number of runs in the major or minor mode having the lowest mean CV value. Amongst the values of K having the lowest CV values from Admixture runs (see figure 12), K = 9 stands out as having a major mode containing 33 runs out of 50, which is also the mode having the lowest mean CV value from the Admixture runs. For other values of K, such as 4, 6, 7, 8, the major modes do not have the lowest mean CV values. C: Admixture plots for all 629 samples for K = 3 (major mode containing 49 out of 50 runs) and K = 9 (major mode containing 33 out of 50 runs). Reference populations on the left have a colour code under the admixture plot that recapitulates their colour on the PCA plots of figure 2; other populations are indicated with alternating grey and white colours.

Typically, for K = 2, all 50 runs are in a single mode and for K = 3, the major mode contains 49 out of all 50 runs and reflects the three main groups from the PCA analysis. Amongst the values of K having the lowest CV values (figure 3A), K = 9 stands out as having a major mode containing 33 runs out of 50. While K = 8 had the lowest overall mean CV value, its major mode contained only 12 runs, indicating K = 9 to be the better model (figure 3B and supplementary table 3). Interestingly, the pattern observed when considering only K = 3 genetic backgrounds, recapitulates the general pattern observed in the PCAs, in which the reference populations separate into three groups. These groups reflect the main evolutionary lineages present in the dataset, being the M lineage (*A. m. mellifera* and *A. m. iberiensis*), C lineage (*A. m. ligustica*, *A. m. carnica*), and O lineage (*A. m. caucasia*). For K = 2, these *A. m. caucasia* bees are considered as having the same genetic background as the *A. m. ligustica* and *A. m. carnica* samples, also reflecting the results from the PCA (figure 2, supplementary figure 19). Our results support the assumption that *A. m. caucasia* bees are assigned to the O lineage by morphometry 22/10/2021 17:15:00 and to the C lineage by mtDNA [57]. Some admixture can be observed for a small proportion of the reference samples. For instance, the reference samples from the Savoy conservatory appear to have a small proportion of genetic background from *A. m. ligustica* and/or *A. m. carnica,* which is consistent with the PCA results (figure 2). Likewise, the *A. m. carnica* samples from Poland have a small proportion of genetic background from *A. m. caucasia*. Finally, the *A. m. carnica* from Switzerland show some proportion of *A. m. mellifera* genetic background. When examining the admixture pattern representing the 33 runs at K = 9 genetic backgrounds, the three main groups are now separated. The M lineage group from the K = 3 backgrounds is now composed of four genetic backgrounds: *A. m. iberiensis* is now separated from *A. m. mellifera* and the *A. m. mellifera* bees are separated in three groups from mainland France, and the two islands of Ouessant and Colonsay. The other three subspecies *A. m. ligustica*, *A. m. carnica* and *A. m. caucasia* each have their own genetic background. An eighth background corresponds to the samples from the bees selected for the production of royal jelly and a ninth appears in the two populations that were noted as Buckfast bees. Although it is a major background in these two populations, a majority of samples have also a large proportion of *A. m. carnica* and, to a lesser extent, of *A. m. ligustica* backgrounds. This ninth background can also be found in other breeder’s populations principally in Hérault and Tarn1 (figure 3C). Apart from the royal jelly population, all honey bees from breeders show high levels of admixture. Moreover, there is a great variability in the genetic origins and proportions of backgrounds, even for samples coming from a same location (figure 4). The exception is the population from Corsica, for which all samples show proportions close to 75% - 25% of *A. m. mellifera* and *A. m. ligustica* backgrounds respectively.

**Figure 4:**
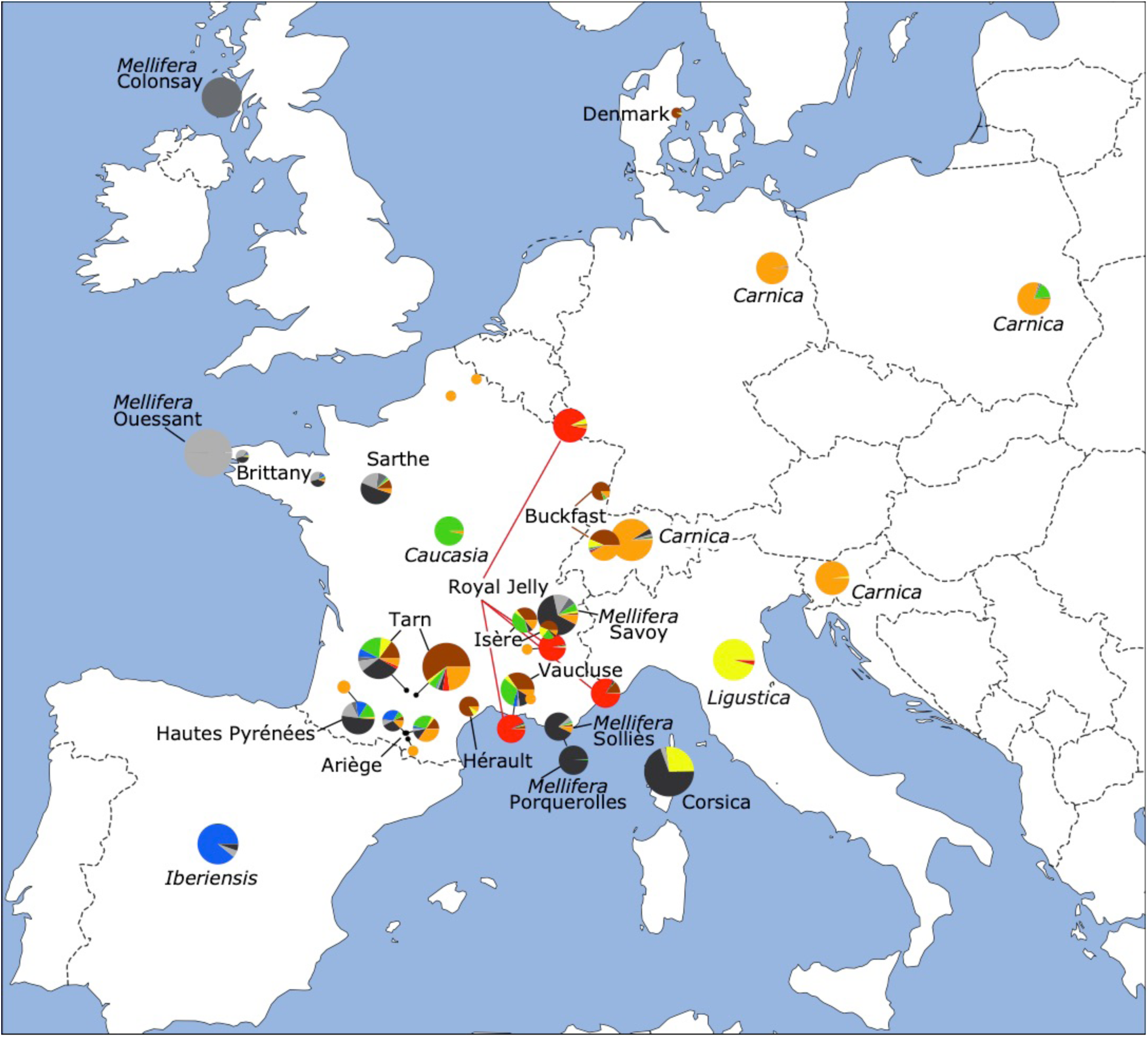
Admixture proportions and location of sample populations used in the diversity study. The size of the pie charts indicates the number of samples from a given location, with the number ranging from 2 samples (e.g. Denmark) to 43 samples (Corsica). Positions in France indicate the coordinates of the breeder or honey bee conservatory sampled. In other countries, reference samples are all grouped together, unless two genetic types were sampled (e.g. Switzerland). Colours in the pie charts correspond to the backgrounds found in the admixture analysis for K = 9, as presented in figure 3. Reference populations for the five subspecies are indicated in italics. Two Buckfast populations in France and Switzerland are indicated, so as the four breeders from the Royal Jelly breeder organisation (GPGR: Groupement des Producteurs de Gelée Royale) having provided samples.

### Migrations between populations

Due to the commercial interest expressed by beekeepers for the Buckfast bees and the peculiar genetic composition observed in the Corsican population, we performed a population migration analysis with TreeMix [52]. All samples having more than 80% ancestry from one of the 9 backgrounds detected in the Admixture analysis were selected from one of the K = 9 major mode Q-matrices (supplementary table 4), and the list supplemented with the 43 Corsican samples, making our data set composed of ten representative groups for the European populations.

Estimations on the number of migrations (m) between the populations in the dataset, based on the Evanno method [53], return a mode of m = 1, strongly suggesting a single migration, and a relatively high Δm value for m = 2 supports the existence of a second migration. The Δm values for 3 or more migrations are close to zero, suggesting that more than 2 migrations between populations in the dataset is unlikely (figure 5A). For m = 1 the 100 TreeMix runs indicated a migration from *A. m. ligustica* to the Corsican population. For m = 2 the 100 TreeMix runs show the two migrations as being from *A. m. ligustica* to the Corsica population, and from *A. m. caucasia* to the Buckfast bees (figure 5B).

**Figure 5:**
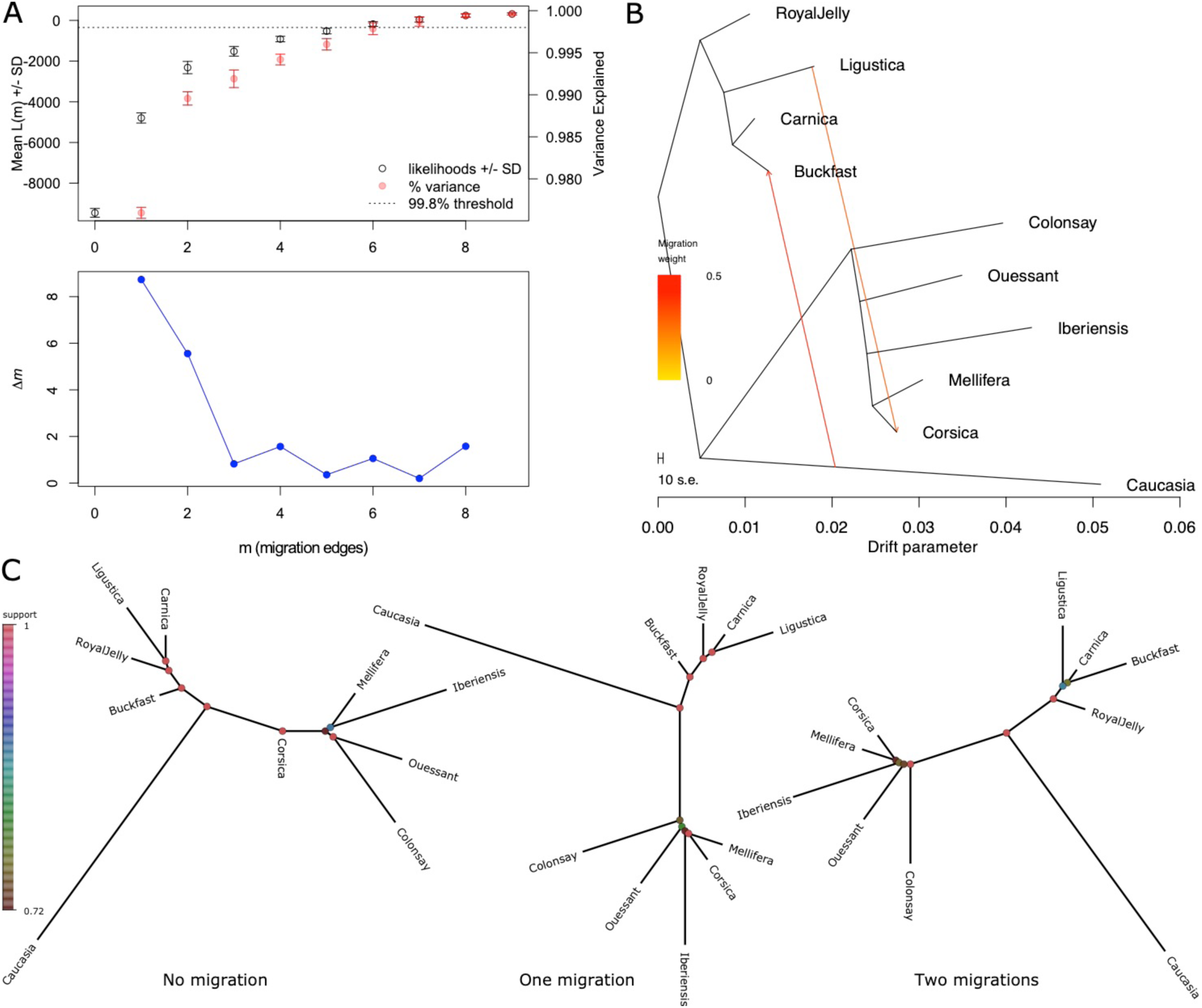
Analysis of migrations with Treemix. A: the OptM package was used to determine the optimal number of migrations between populations and backgrounds. The Δm values suggest one or two migrations. B: TreeMix graph selected amongst the 100 runs showing the two migrations identified. C: summaries of trees from TreeMix, estimated from 100 runs per migration with DendroPy.

Summaries of the resulting trees with DendroPy [54] are shown in figure 5C, indicating that when the two migrations are taken into account, the Corsican samples are grouped with the *A. m. mellifera* M lineage bees, and the Buckfast bees group with the *A. m. ligustica* and *A. m. carnica* C lineage bees.

### Haplotype conservation in the admixed populations

To investigate further the haplotype blocks detected, we performed a local ancestry inference on our dataset with RFMix. Reference samples were selected as bees having > 95 % ancestry for a given background following the Admixture analysis at K = 3 (figure 3C), resulting in 131 samples for group 1, 148 for group 2 and 17 for group 3, while the remaining 333 samples formed the query dataset. To perform the local ancestry inference, we constructed a genetic map from cross overs identified in the sequence data of 43 males from 3 colonies [37] aligned to the HAv3.1 reference genome. Results indicate that few historical recombination events have occurred in the large haplotype blocks since the admixture between the subspecies. The most notable example is that of the 3.6 Mb haplotype block between positions 3.7 and 7.3 Mb on chromosome 11, in which almost all 333 samples from the query dataset show one continuous stretch for one of the three backgrounds. Only one of the 43 samples from Corsica presents two different ancestral haplotypes within this interval, with a switch from a group 1 to a group 2 haplotype at position ∼4.5 Mb on chromosome 11, within the 3.6 Mb haplotype block, whereas numerous switches can be observed on the rest of the chromosome (figure 6A). When counting the haplotype switches detected in all 333 query samples, only 28 are situated within the 3.6 Mb haplotype block on chromosome 11, whereas other regions of the chromosome can have more than 50 switches per 100 kb (figure 6B and see supplementary figure 20 for the other chromosomes). Interestingly, *LOC724287,* which is the largest gene described in the Gnomon annotation set for the genome assembly HAv3.1, is found in this block at position 11:5,292,072-6,161,805. This gene is 869,734 bp long and encodes protein rhomboid transcript variant X2, its large size being largely due to intron 4, which is 596,047 bp long. However, on investigating a possible relationship between haplotype block and gene sizes in the honey bee genome no obvious association could be found (data not shown).

**Figure 6:**
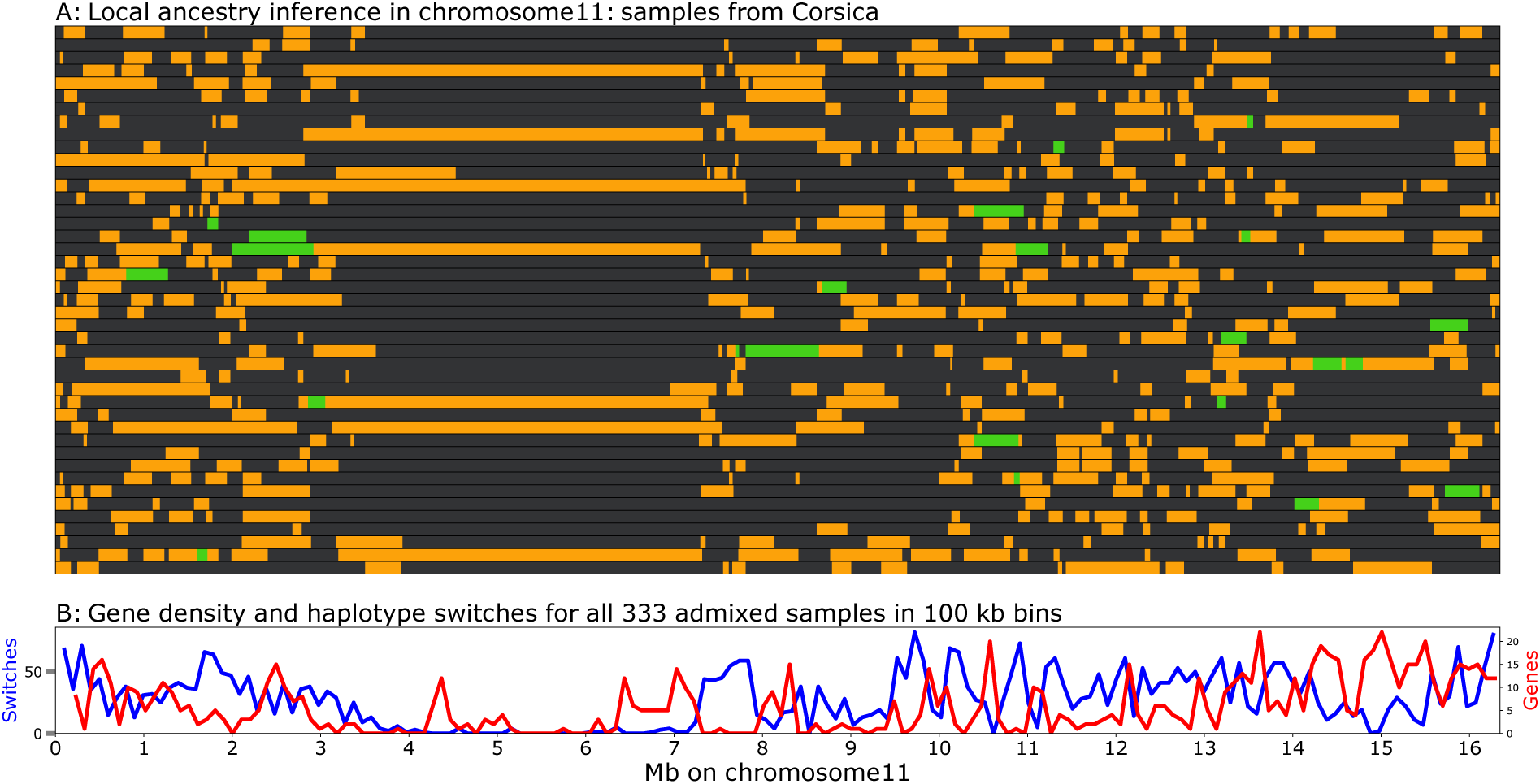
Local ancestry inference on chromosome 11 in the admixed samples from Corsica. A: each horizontal line represents the ancestry inference on one of the 43 individual samples from Corsica. Grey: *A. m. mellifera* and *A. m. iberiensis* backgrounds; yellow: *A. m. ligustica* and *A. m. carnica* backgrounds; green: *A. m. caucasia* background. B: haplotype switches in all 412 admixed samples analysed. The 3 Mb haplotype block at positions 4-7 Mb on chromosome 11 shows very little historical recombination.

## Discussion

### SNP detection in a haploid dataset

Our complete dataset of haploid drones is composed of 870 samples sequenced using Illumina’s HiSeq and NovaSeq technologies. Results clearly show that although a few of the early sequences produced on the HiSeq are of lower depth, only 15 samples were eliminated due to the fraction of missing genotypes exceeding 10%. By contrast, the fraction of missing genotypes over the ∼7 million SNPs detected was considerably lower in samples sequenced on the NovaSeq sequencing platform. Having sequenced haploids, the removal of heterozygote SNPs in individual samples is recommended to reduce the likelihood of “pseudo SNPs”, as we have shown previously that heterozygote SNPs tend to cluster together [31] and co-locate with repetitive elements (data not shown). This set of ∼7 million markers can now be used as a basis for the realization of high-density SNP chips, allowing selections of markers according to optimized spacing and to defined MAF values in the main subspecies of interest. Indeed, an important technical issue in SNP chip design, is that very high SNP densities, such as found in the honey bee, can potentially cause allele dropout when genotyping, due to interference in the probe designs. Deep knowledge of SNP and indel positions will help select candidates flanked by monomorphic sequences. Conversely, for lower density chips, the spacing of markers can be optimized by taking the haplotype structure into account, thus avoiding redundancy while maintaining the highest possible level of genetic information. Another advantage of sequencing haploid samples, is that the whole dataset represents phased chromosomes. Notably, the present dataset will be invaluable for genotype imputation in future studies using lower density genotyping, such as DNA chips or low-pass sequencing [58–60].

### Population structure in managed honey bees

The deep understanding of European honey bee populations and of their recent admixture via imports of genetic stocks by breeders is not a simple task. The analyses of admixture events in complex population structures can be sensitive to a number of parameters and sometimes yields misleading results, especially if one or several populations went through a recent bottleneck [61]. PCA on all ∼7 million markers indicate that our dataset is structured into three main genetic types (supplementary figure 8). The first principal component, representing 10.8 % of the variance, separates two major groups corresponding respectively to subspecies from north-western (M lineage) and south-eastern Europe (C lineage). These two groups are represented by several populations, including the Savoy and Porquerolles conservatories from South-East France on one side, and bees that are not so far geographically from Italy or Slovenia on the other. This large genetic distance despite relatively close geographic proximity of the populations supports the hypothesis of the colonization of Europe by honey bees via distinct western and eastern routes [1,24,62,63], and the separation between subspecies due to the Alps forming a natural barrier preventing genetic exchange [5]. Along the second principal component, representing 3.1 % of the variance, the population originating from *A. m. caucasia* separates from the south-eastern European populations (supplementary figure 8). Prior to investigating admixture we pruned SNPs in LD taking care to maximize the removal of redundancy while maintaining the general structure of the data (figure 2 and supplementary figures 15, 16, 17).

We explored a range of K number of genetic backgrounds, running multiple iterations of each, to determine the most likely admixture pattern (figure 3). Our results indicate that this approach is necessary to ensure the results from each K model are stable prior to interpretation. We observe from our Admixture analyses that CV outliers within a K model are common. For instance, at K = 8, the mode with the lowest CV is only represented by 8 out of 50 Admixture runs, whereas the major mode has 12 runs. On examining the admixture patterns from these two modes, the major mode suggests the *A. m. mellifera* bees from conservatories on mainland France to be hybrids between bees from Ouessant and Spain, with roughly 50% of each genetic background moreover on the same mode, the *A. m. iberiensis* background represents also 50% of the M lineage background in the bees from Corsica (supplementary figure 19). This is extremely unlikely given the geography of Western Europe and our knowledge of the history of the bees of Ouessant. Indeed, Ouessant is a very small island (15.6 km^2^) off the coast of western Brittany, isolated from the rest of the French honey bee population since its installation in 1987 and the prohibition of imports since 1991 mostly for sanitary reasons. In contrast, the mode with the 8 runs and lowest CV presents a better separation of *A. m. mellifera* and *A*. *m. iberiensis,* which is also found in the major mode at K = 9 backgrounds. A smaller level of admixture can still be found between *A. m. mellifera* and *A. m. iberiensis,* that is quite likely due to the shared ancestry between these two subspecies.

The major mode at K = 9 is represented by 33 out of 50 runs and returned the lowest mean CV value. This mode identifies mainland France *A. m. mellifera* samples as having a distinct genetic background and suggests that honey bees from Ouessant may have been re-introduced in the mainland conservatories. This mode also identifies a distinct genetic background in French and Swiss Buckfast bees. Buckfast bees were developed by Brother Adam, and are described on page 14 of “Beekeeping at the Buckfast Abbey” as a cross performed around 1915 between “the leather-coloured Italian bee and the old native English variety” [16]. Brother Adam also notes that the Italian bees that were imported in later years were distinct from the ones used in the development of the Buckfast strain. Our analysis of migrations between populations with TreeMix suggests that the Buckfast in our dataset were subject to introgression with genetic material from *A. m. caucasia* (figure 5B), although the timing of this potential admixture event could not be determined. When the two migrations of *A. m. ligustica* into Corsica and *A. m. caucasia* into the Buckfast are considered, which is a likely scenario suggested by the Evanno analysis, the latter is close to *A. m. carnica*, as seen in figures 5B and 5C. Interestingly, a whole genome sequence study of Italian honey bees, also suggest that the Buckfast bees are closer to *A. m. carnica*, than to *A. m. ligustica* [64] and no proximity of the Buckfast bees with M lineage bees were found neither in this study nor in ours, despite the cross at the origin of the Buckfast including an old native variety. Further investigations including more Buckfast samples and additional honey bee subspecies will be needed to fully elucidate this question. The *A. m. carnica* samples from Slovenia, Germany, France, Switzerland and Poland all share the same genetic background, reflecting their identical origin, probably recent imports.

The population of bees from Corsica has the distinct characteristic of being homogenous in composition, despite being admixed, with all samples showing mean proportions of 75% and 25% of *A. m. mellifera* and *A. m. ligustica* backgrounds, respectively (figures 2 and 3). The introgression of Italian bees is confirmed by the TreeMix migration analysis and when this is accounted for, the Corsican samples group with *A. m. mellifera* bees from mainland France instead of being situated between the two main genetic subgroups of western and eastern European bees (figures 2B and 2C). This result likely reflects the fact that Italian bees may have been imported on the Island until the 1980’s, following which the import of foreign genetic material was prohibited. As beekeepers generally prefer the *A. m. ligustica* Italian bees over *A. m. mellifera*, it is very likely that the latter is the original population, as also suggested by Ruttner [1]. Although the hypothesis of the separation of the two subspecies on the mainland by the Alps seems appropriate [5], the situation of the Mediterranean islands in the region is not so clear. Based on physical geography alone, Corsica being at a closer distance to Italy than to France, the chances would have been greater to have originally M lineage rather than C lineage Italian bees. Moreover, Corsica was under the control of Pisa, then fell to Genoa in 1284 and was only purchased by France in 1768. Further studies including samples from Sardinia would certainly help defining the Mediterranean boundaries between the M lineage and C lineage honey bees and confirm observations based on morphology [1].

Apart from the subspecies references and the royal jelly populations, the honey bees provided by breeders are largely admixed, exhibiting high variability in background proportions - even for samples sourced from the same region. A typical example is that of the Tarn1 and Tarn2 populations, revealing that two breeders situated very close to one another (less than 100 km), have very different genetic management strategies. Tarn1 samples are mainly composed of Buckfast and *A. m. carnica* backgrounds, whereas in Tarn2 a large proportion of *A. m. mellifera* background is also present and the population is far less homogenous (figures 2, 3 and 4). This shows the great heterogeneity of the managed populations found in France and a question that needs further investigation is the influence of the mating strategies used by the breeders, such as artificial insemination, mating stations, with drone producing hives to saturate the environment with the desired genetic strains, or open mating. These strategies influence variable levels of control on the genetic makeup of a breeder’s stock. The higher proportion of *A.m. mellifera* background in the Tarn2 population could either be deliberate or due to a lower level of control over the mating of the queens, with a proportion of queens mating to *A. m. mellifera* drones from the environment.

The Royal Jelly population is the inverse: beekeepers from all over France exchange their genetic stock within a selection programme and practice controlled mating. As a result, a specific background with individuals presenting very little admixture, is found for this population at very distant locations. Most of the worldwide production of Royal Jelly comes from China, where high Royal Jelly-producing lineages of honey bees were developed from an imported *A. m. ligustica* lineage [65]. Interestingly, in our dataset, only three *A. m. ligustica* and all of the bees from China have some Royal Jelly genetic background.

### Large haplotype blocks in the honey bee genome, specific to the M and C lineages

When investigating the contribution of SNPs to variance in the PCA, we noted several large genomic regions, up to 3.5 Mb long, in which almost all markers contributed very strongly to the first principal component, separate bees from north-western (M lineage) and south-eastern Europe (C lineage). These regions were noted to coincide with haplotype blocks detected with Plink. To investigate the matter further, we performed local ancestry inference in the admixed samples with RFMix, using samples exhibiting 95% ancestry for the three main genetic backgrounds as references. A low recombination rate is confirmed by the observation of very few switches between the three main genetic backgrounds within these haplotype blocks. Interestingly, some of our regions, including the largest one detected on chromosome 11, coincide with regions of low recombination rate detected in other studies. These include a LD map produced with 30 diploid sequences from African worker bees [39], ancestry inference in an admixed population [35], low resolution genetic maps produced by Rad or ddRAD sequencing, with microsatellite or SNP markers, ddRAD sequencing [66, 67] or higher resolution genetic maps produced by whole genome sequencing of European [37] and African subspecies [38]. Most of these regions coincide with the position of the centromeres such as described in the reference genome assembly, which is primarily based on the combination of the location of *Ava*I repeats, that were previously assigned to centromeres by cytogenetic analysis, and of a low GC content [33, 68]. However, the *Ava*I repeats only represent a very small fraction of the centromeric regions described, with the largest one only covering 14 kb [33], whereas the estimation of the extent of the centromeres, based on a GC content lower than the genome average is much larger although imprecise and supposes a similar organization as for the AT-rich alpha-satellite repeats in vertebrates, such as human [69]. Whereas in some cases the boundaries of our regions of low recombination rate coincide with the actual positioning of the centromere on the genome assembly [33], such as in chromosomes 5 or 8, in other instances, such as in chromosome 12, the region defined is much narrower. Due to the difficulties in interpreting banding patterns in honey bee chromosomes, the position of the centromeres is not well defined. Some evidence based on G- and C- banding suggests there are four metacentric and 12 submetacentric or subtelocentric chromosomes [70], whereas other evidence based on the fluorescent *in situ* hybridization of a centromere probe suggests there are two metacentric, four submetacentric, two subtelocentric and eight telocentric chromosomes [71]. Our evidence suggests at least six chromosomes that could be telocentric or acrocentric: chromosomes 3, 5, 6, 9, 14 and 15.

Some of the haplotype blocks/regions of low recombination can seem very large, such as representing up to 21 % in the case of chromosome 11 (figure 6). This may seem a lot, but recent findings in a complete sequencing of the human genome give a similar proportion for chromosomes 9, in which 40 Mb of satellite arrays represent 20 % of the chromosome [72]. One important difference, however, is that the block on honey bee chromosome 11 contains some genes, except in the central region, whereas the satellite array described on human chromosome 9 does not. This reaffirms that our understanding of the centromere positions in the honey bee chromosomes requires refinement. The specific case of the acrocentric chromosomes in terms of gene content (supplementary figure 20) seems to compare better to the situation described in human, as the sequencing of the p-arm of the five human acrocentric chromosomes has allowed the discovery of novel genes within the satellite repeat-containing regions [69].

Some haplotype blocks may have another origin than centromeric DNA. For instance, some may have maintained genetic divergence by limiting recombination via the presence of structural variants such as inversions. Indeed, two of the blocks described here, between positions 4.0 - 5.1 Mb and 5.8 - 6.9 Mb on chromosome 7, seem to coincide at least partially with two regions of haplotype divergence possibly due to inversions, detected between positions 3.9 – 4.3 and 6.3 – 7.3 Mb on the same chromosome, in a highland versus lowland study of East African bees [36]. The slight differences in coordinates found between the two studies could be due to the fact that different version of the HAv3 assembly were used. However, if confirmed, this finding suggests that haplotype blocks differing between M lineage and C lineage bees such as found here, might coincide with blocks found in other subspecies in Africa. Another study identifying the thelytoky locus (*Th*) in the South African Cape honey bee *Apis mellifera capensis* showed it was in a non-recombining region over 100 kb long on chromosome 1, although long-read mapping failed to detect any inversion [73].

Given the current hypotheses on the colonization of Europe by honey bees via distinct western and eastern routes [1,24,62,63], it is not surprising that the haplotype blocks described here, whether or not representing centromeric regions, tend to separate mainly the M and C lineage bees. Further analyses will be necessary to define the centromeric regions more precisely and study their implication, together with the other haplotype blocks, in the sub species structure of the honey bee populations.

## Conclusion

The sequencing of close to 900 haploid honey bee drones, was shown here to be an invaluable approach for variant detection and for understanding the fine genetic makeup of a complex population having gone through multiple events of admixture. In addition, the extent of regions of extremely low recombination rate could be defined with a higher precision than previously. The dataset generated here is a solid base for future research involving other honey bee populations and for any analyses requiring a reference set for phasing or imputation.

## Supporting information

Scripts

## Acknowledgements

This work was performed in collaboration with the GeT platform, Toulouse (France), a partner of the National Infrastructure France Génomique, thanks to support by the Commissariat aux Grands Invetissements (ANR-10-INBS-0009). Bioinformatics analyses were performed on the GenoToul Bioinfo computer cluster. This work was funded by a grant from the INRA Département de Génétique Animale (INRA Animal Genetics division) and by the SeqApiPop programme, funded by the FranceAgriMer grant 14-21-AT. We thank Andrew Abrahams for providing honey bee samples from Colonsay (Scotland), the Association Conservatoire de l’Abeille Noire Bretonne (ACANB) for samples from Ouessant (France), CETA de Savoie for sample from Savoie, ADAPI for samples from Porquerolles and all beekeepers and bee breeders who kindly participated to this study by providing samples from their colonies.

## Data Accessibility

DNA Sequences for this project have been deposited in the Sequence Read Archive (SRA) at www.ncbi.nlm.nih.gov/sra under the BioProject accessions PRJNA311274 as part of the SeqApiPop French honey bee diversity project dataset and PRJEB16533 as part of the Swiss honey bee population and conservation genomics project dataset. Individual SRA run and BioSample accessions for all samples are given in supplementary table 1. A vcf file with the filtered 7 million SNP and 870 samples is available at https://doi.org/10.5281/zenodo.5592452 for download, together with the list of the 629 unique samples.

## Authors’ contributions

YLC, J-PB, BB and AV designed the experiment. BB, YLC, and AV coordinated colony selection and sampling and samples were provided by KB, MB, CC, AG, PK, MP and AP. KC-T, EL and OB performed DNA extraction, library preparation and sequencing. DW, AV, SE and BS performed the bioinformatic analyses and co-wrote the manuscript.

