## Supplementary figures for "Complex population structure and haplotype patterns in Western Europe honey bee from sequencing a large panel of haploid drones"

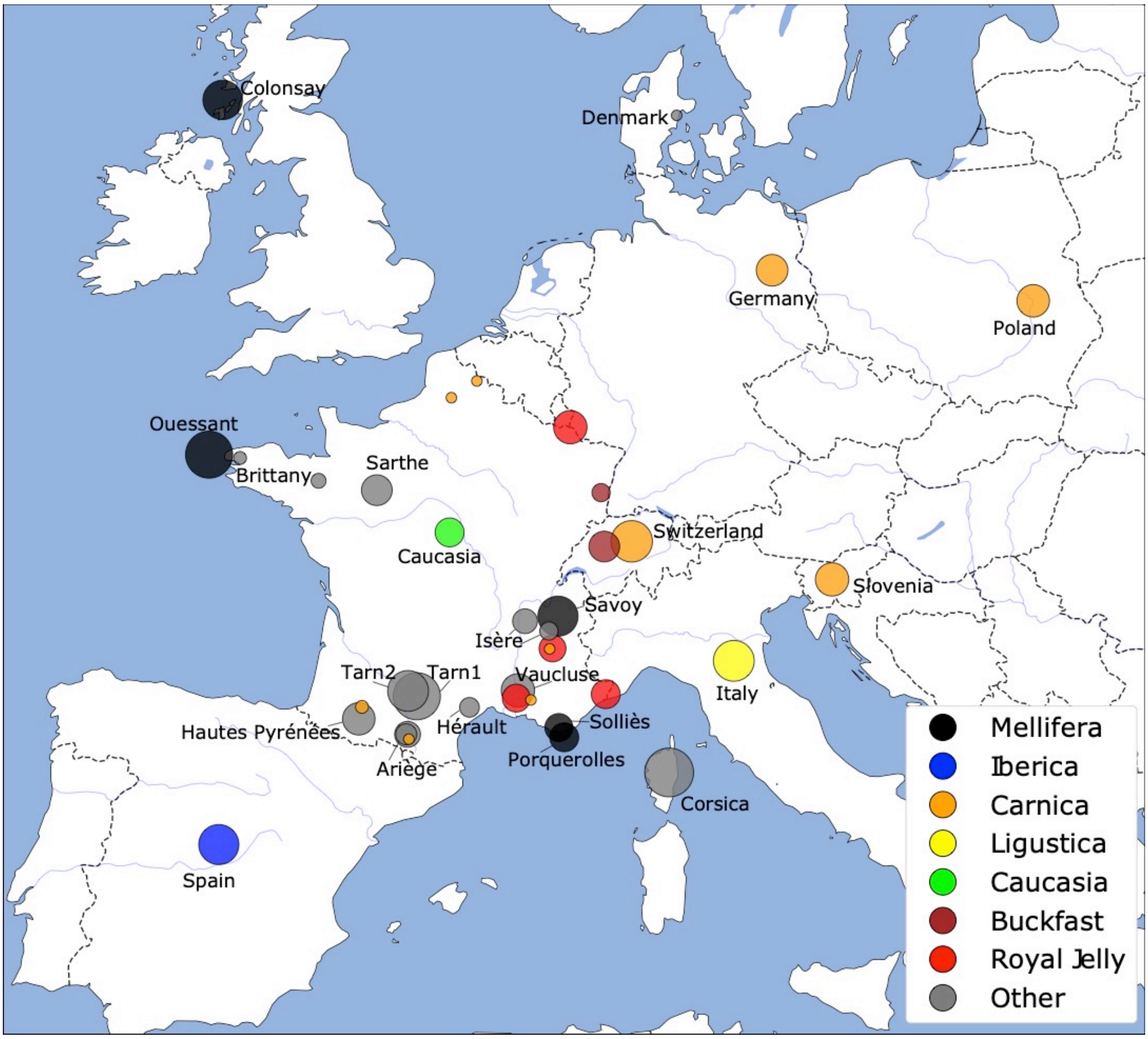


**Figure 1: Location of samples used in the study.** Colours indicates the presumed genetic type and the size of the circles the number of samples from a given location, with the number ranging from 2 samples (e.g. Denmark) to 43 samples (Corsica). Positions in France indicate the coordinates of the breeder or honey bee conservatory sampled. In other countries, reference samples are all grouped together, unless two genetic types were sampled (e.g. Switzerland).

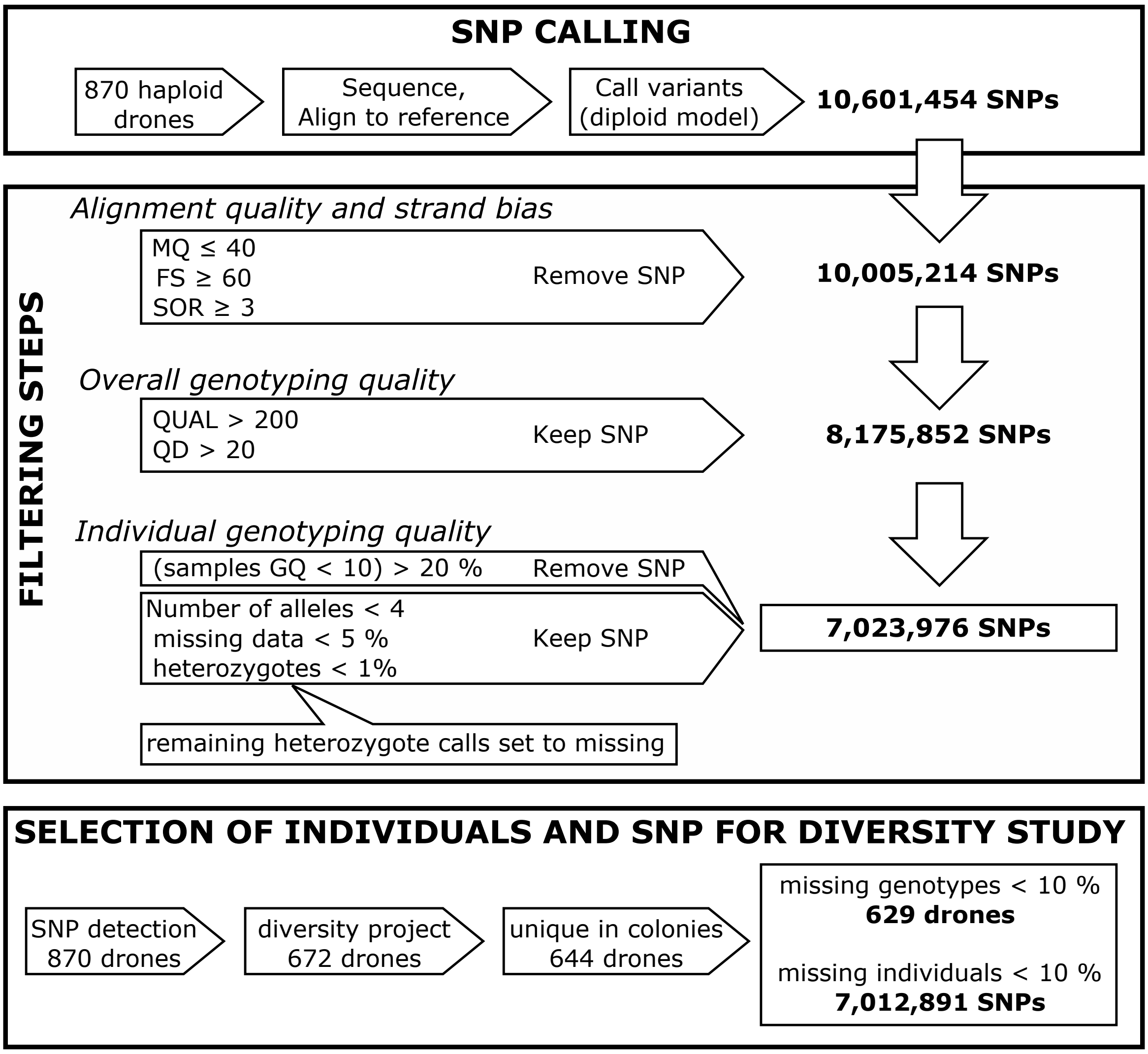


**Figure 2: Overall strategy for SNP calling and filtering.** Variant calling and technical filtering was done on a large dataset of 870 drone samples, to increase robustness. The final dataset for the diversity study is of 7,012,891 SNPs and 629 individuals.

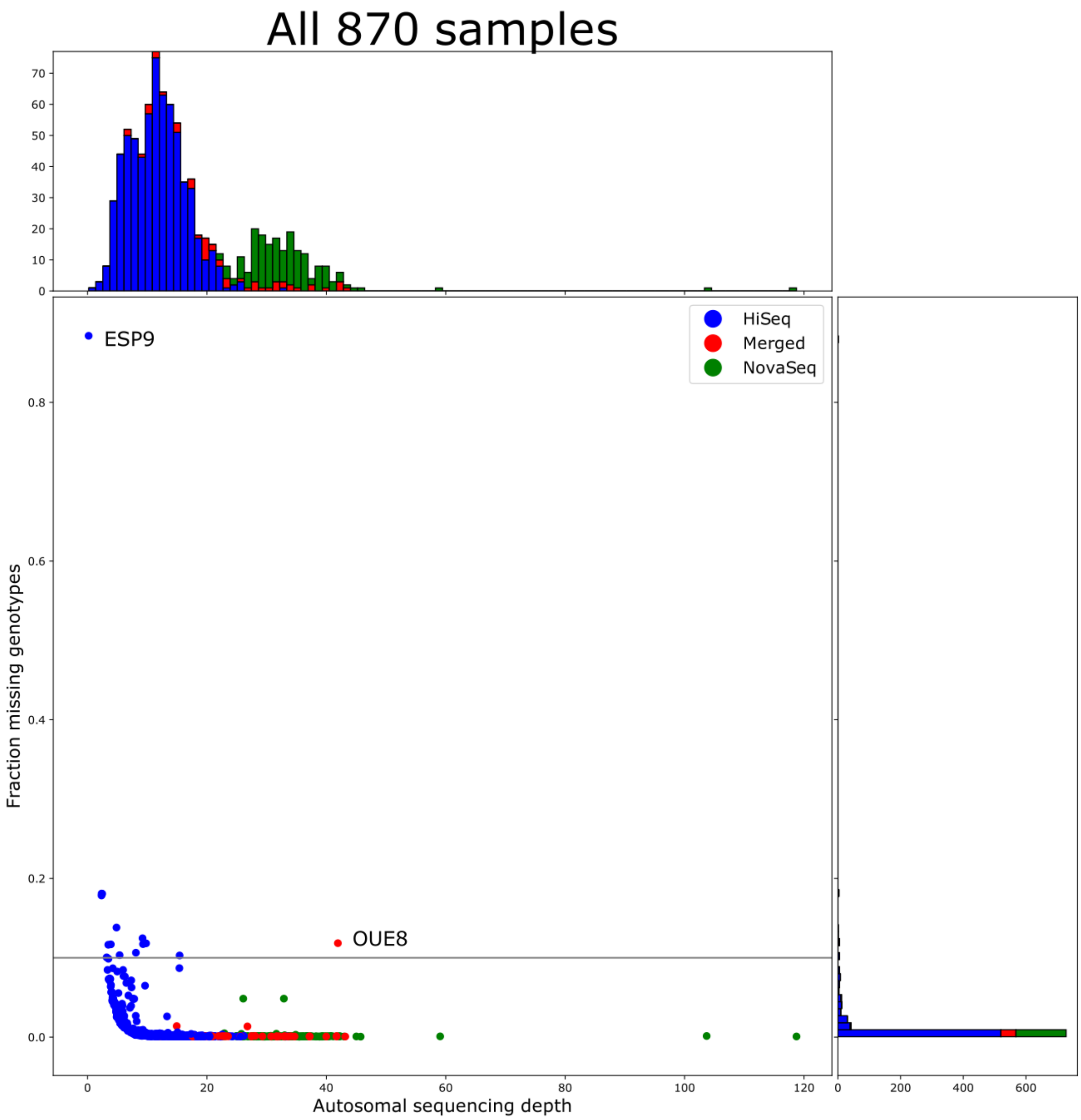


**Figure 3: Sequencing depth and fraction of missing genotypes for all the 870 sequenced samples.**

Blue: samples sequenced with the Illumina™ HiSeq instrument; green: samples sequenced with the Illumina™ NovaSeq instrument and red: samples sequenced with the Illumina™ HiSeq instrument in two or more runs. The horizontal line is the genotyping rate threshold applied to final dataset. Sequencing data could not be obtained from sample ESP9 and data for sample OUE8 was obtained from 3 sequencing runs.

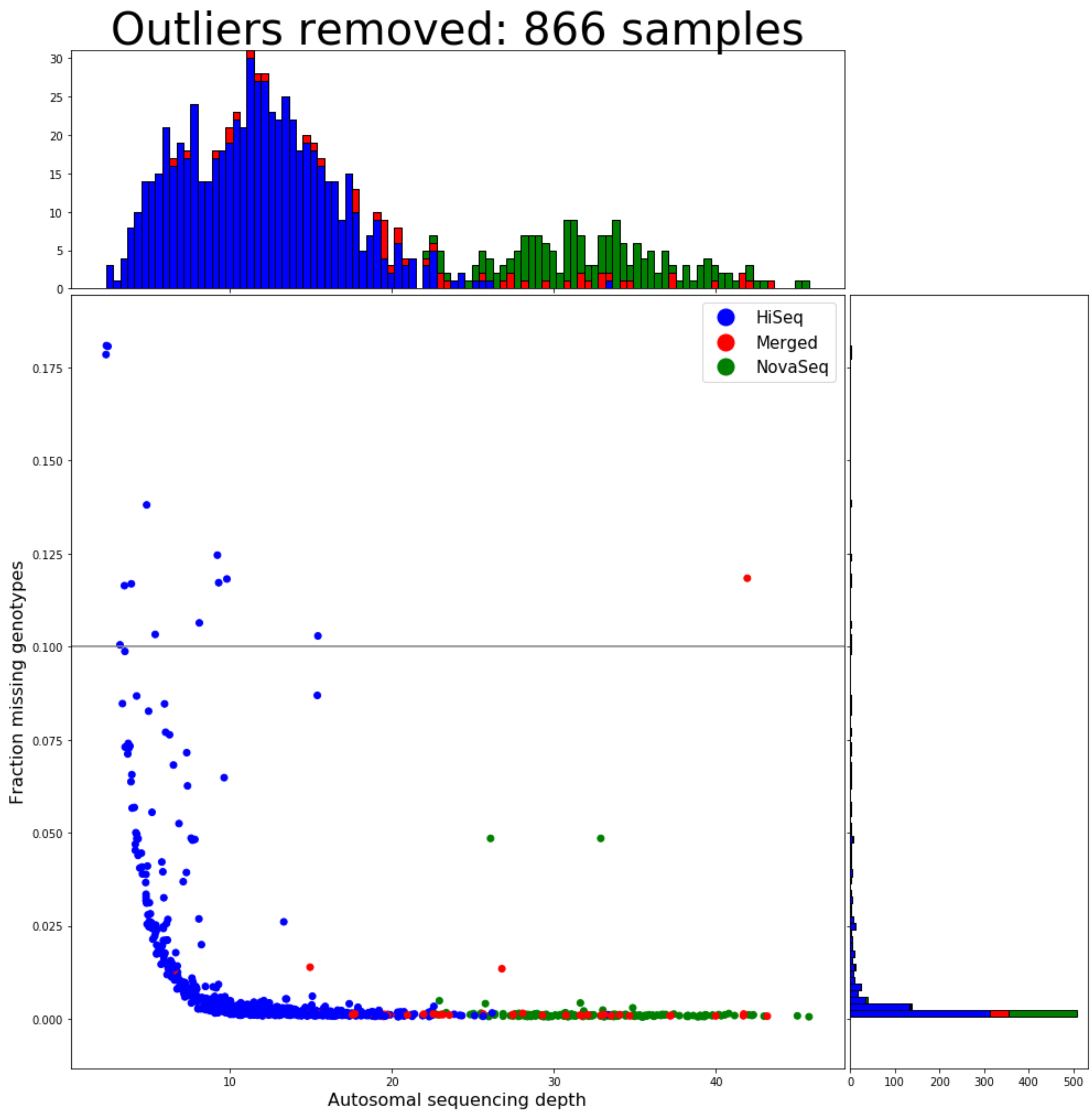


**Figure 4: Sequencing depth and fraction of missing genotypes with outliers removed.** Blue: samples sequenced with the Illumina$^{TM}$ HiSeq instrument; green: samples sequenced with the Illumina$^{TM}$ NovaSeq instrument and red: samples sequenced with the Illumina$^{TM}$ HiSeq instrument in two or more runs. The horizontal line is the genotyping rate threshold applied to final dataset.

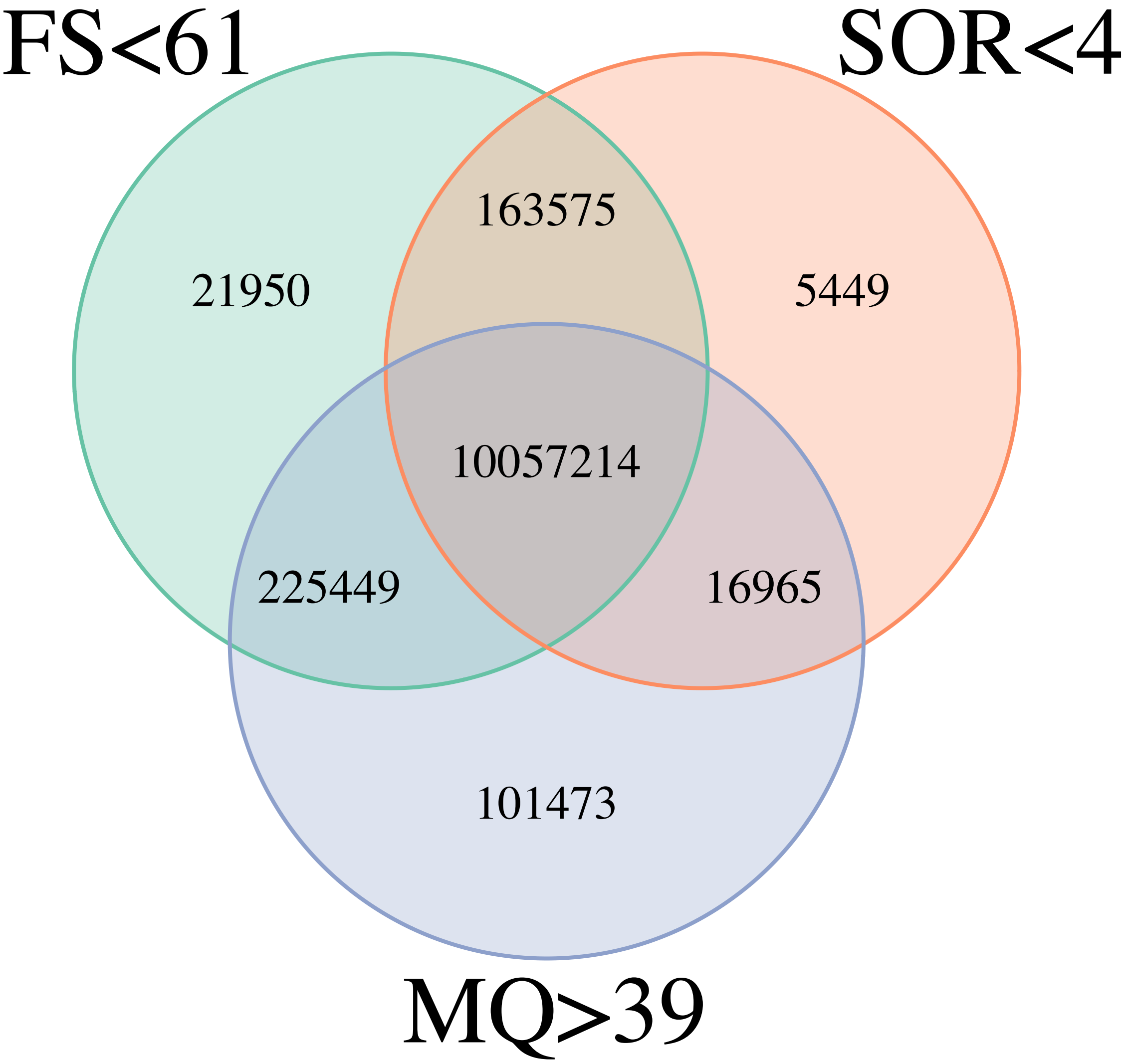


**Figure 5: Filters on alignment quality (MQ) and strand bias (FS and SOR) metrics.**
FS (FisherStrand): phred-scaled probability that there is strand mapping bias at the site; SOR (StrandOddsRatio): strand bias mapping estimate; MQ (RMSMappingQuality): root mean square mapping quality over all the reads at the site. The intercept of the 3 filters was used for further filtering (see figure 2).

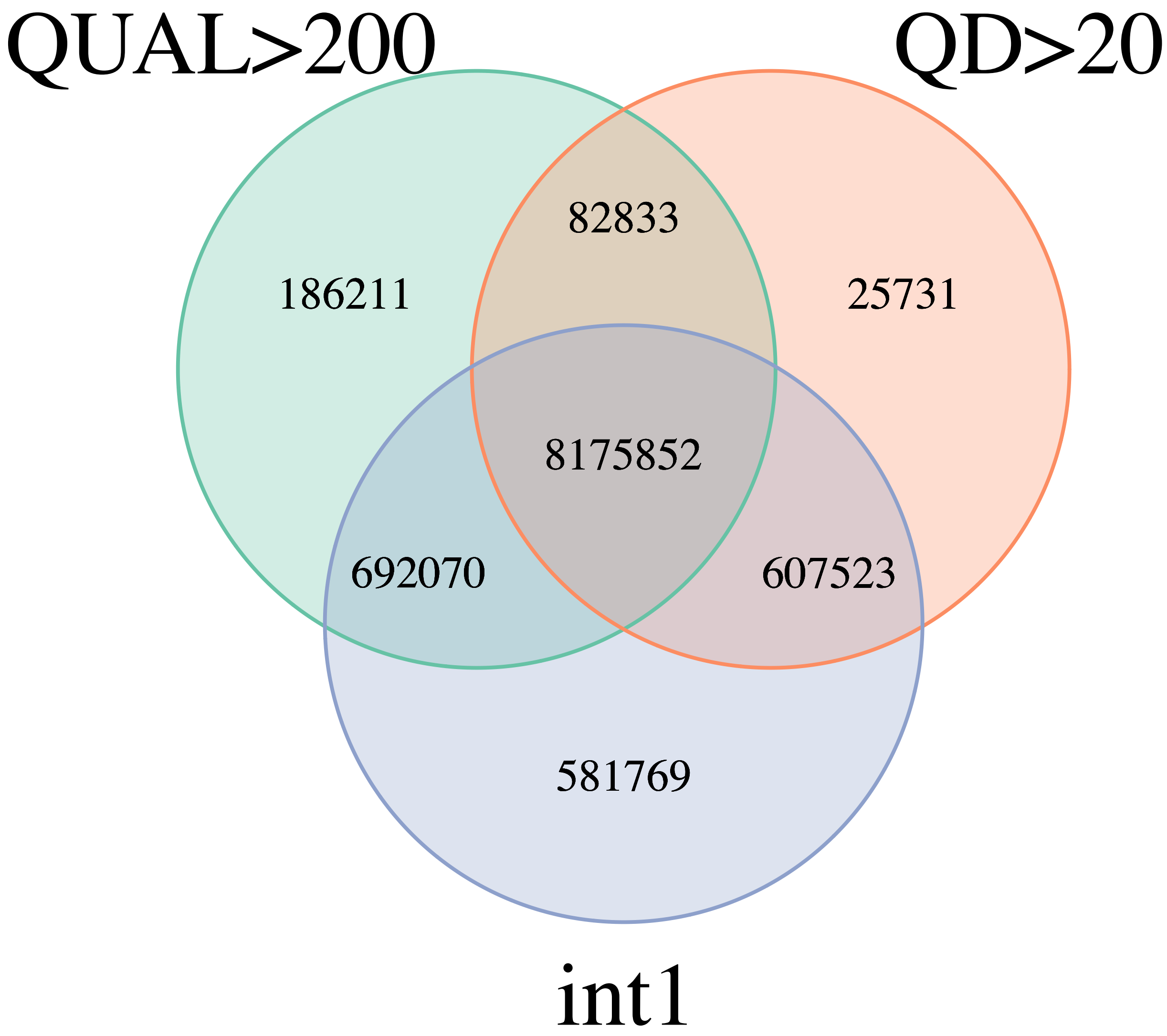


**Figure 6: filters on overall genotyping quality.**
Int1 is the intersect of the mapping quality filters. QUAL: Phred-scaled quality score for the assertion made in ALT: the more samples have the ATL allele, the higher the QUAL score. QD: quality score normalized by allele depth in which only informative reads are counted. The intercept of the 2 filters with the previous mapping quality filters was used for further filtering (see figure 3).

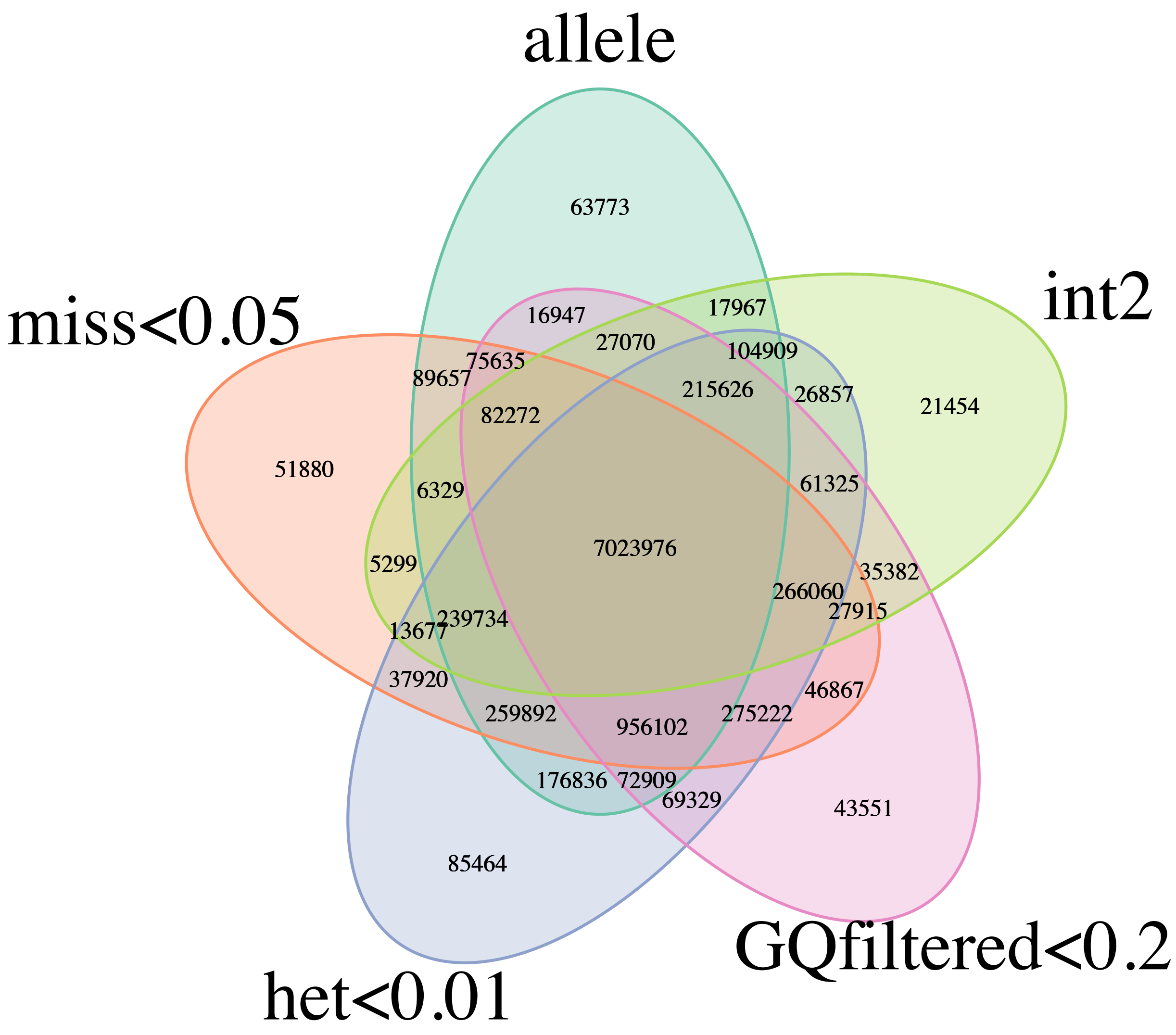


**Figure 7: filters on individual genotyping quality.**
Int2 is the intersect of the previous filters. Filters are (i) het: proportion of heterozygote calls less than 1% for a SNP, as haploid drones were sequenced, the remaining heterozygote calls were set to missing; (ii) allele: less than 4 alleles for a SNP; (iii) miss: less than 5% missing data; (iv) GCfiltered: SNPs are removed if more than 20% samples have a genotyping quality (GQ) under 10. Note: although SNPs with more than 5% of missing data were filtered out, some markers may have more than 5% missing data due to the heterozygote calls that were set to missing.

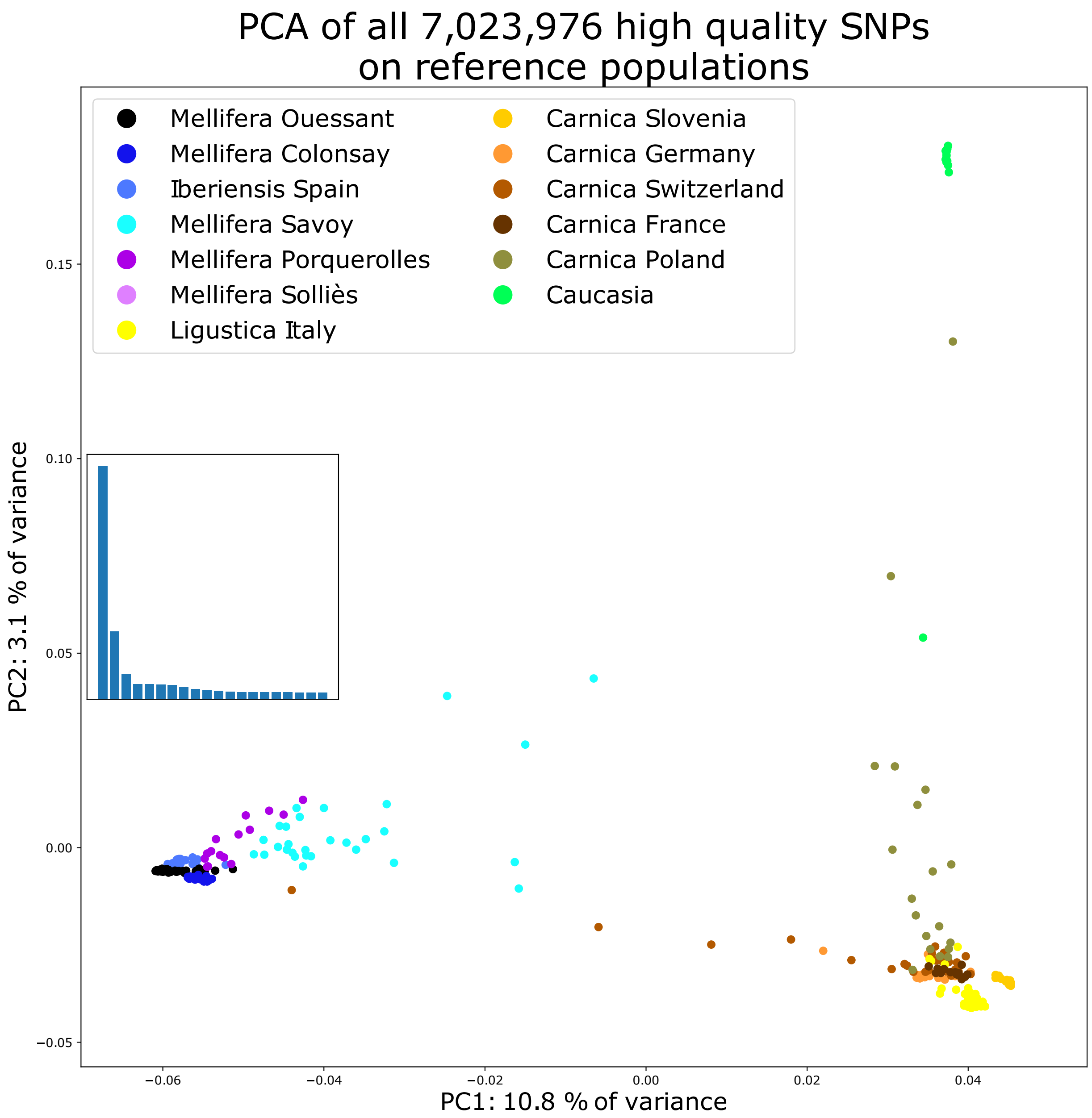


**Figure 8: Principal component analysis for all SNPs: reference populations.** The first component separates clearly the *A. m. mellifera* and *A. m. iberiensis* on one side and the *A. m. ligustica*, *A. m. carnica* and *A. m. caucasia* on the other. The second distinguishes the *A. m. caucasia* from the rest. The blue barplot in the inset represents the proportion of the variance represented by the first 20 components.

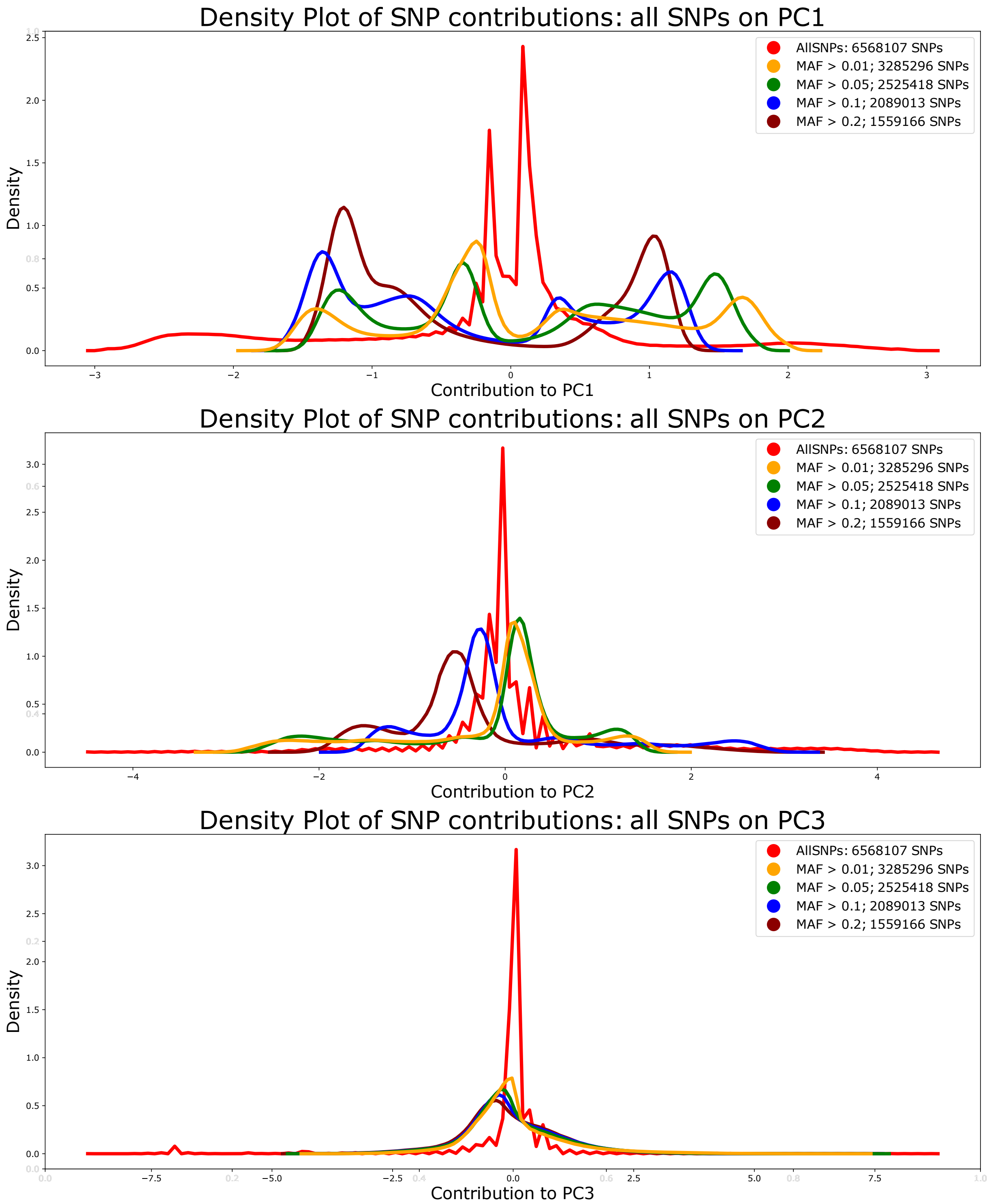


**Figure 9: Contributions of SNPs to PCs 1, 2 and 3, according to MAF filters.** With no MAF filtering (red lines), most SNPs contribute very little to PC1 and most SNPs do not contribute to PC2 and PC3 at all. With MAF filtering, the proportion of SNPs contributing to the PCs increases, including for PCs 2 and 3.

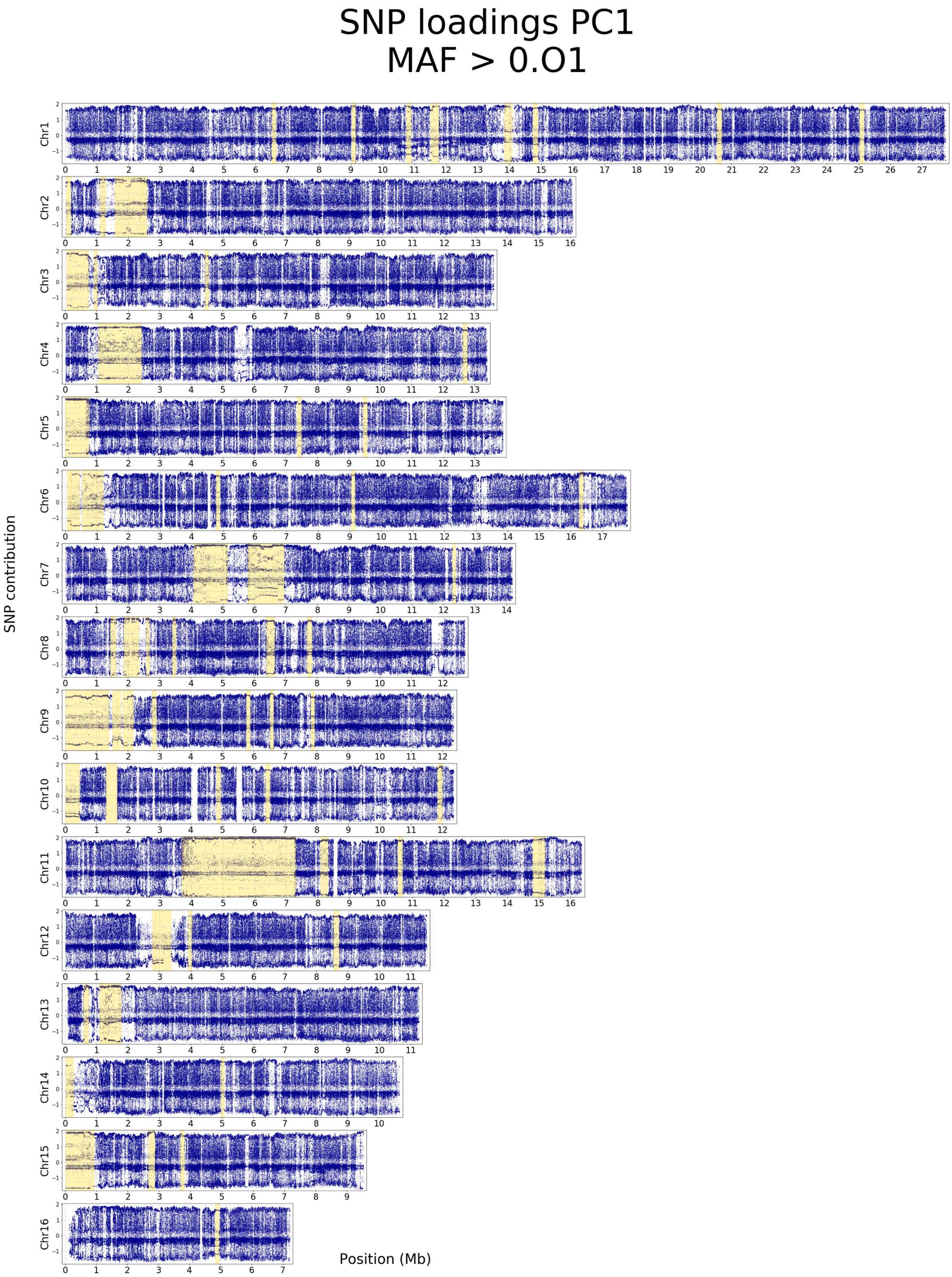


**Figure 10: Contribution of SNPs with MAF > 0.01 to PC1 on all 16 chromosomes.** Yellow backgrounds correspond to haplotype blocks detected with the plink blocks function, of size larger than 100 kb. SNP contributions were estimated with SMARTPCA.

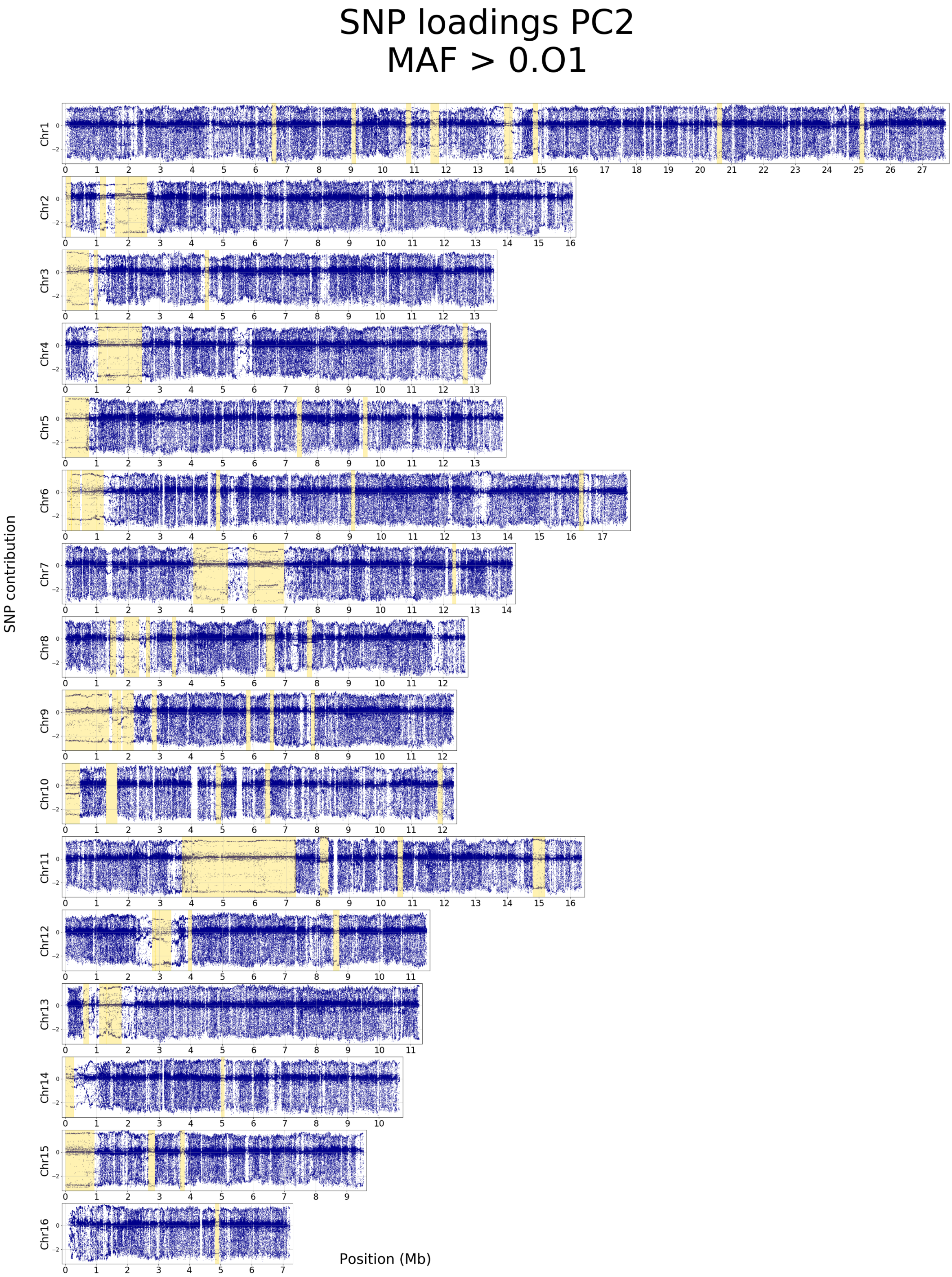


**Figure 11: Contribution of SNPs with MAF > 0.01 to PC2 on all 16 chromosomes.** Yellow backgrounds correspond to haplotype blocks detected with the plink blocks function, of size larger than 100 kb. SNP contributions were estimated with SMARTPCA.

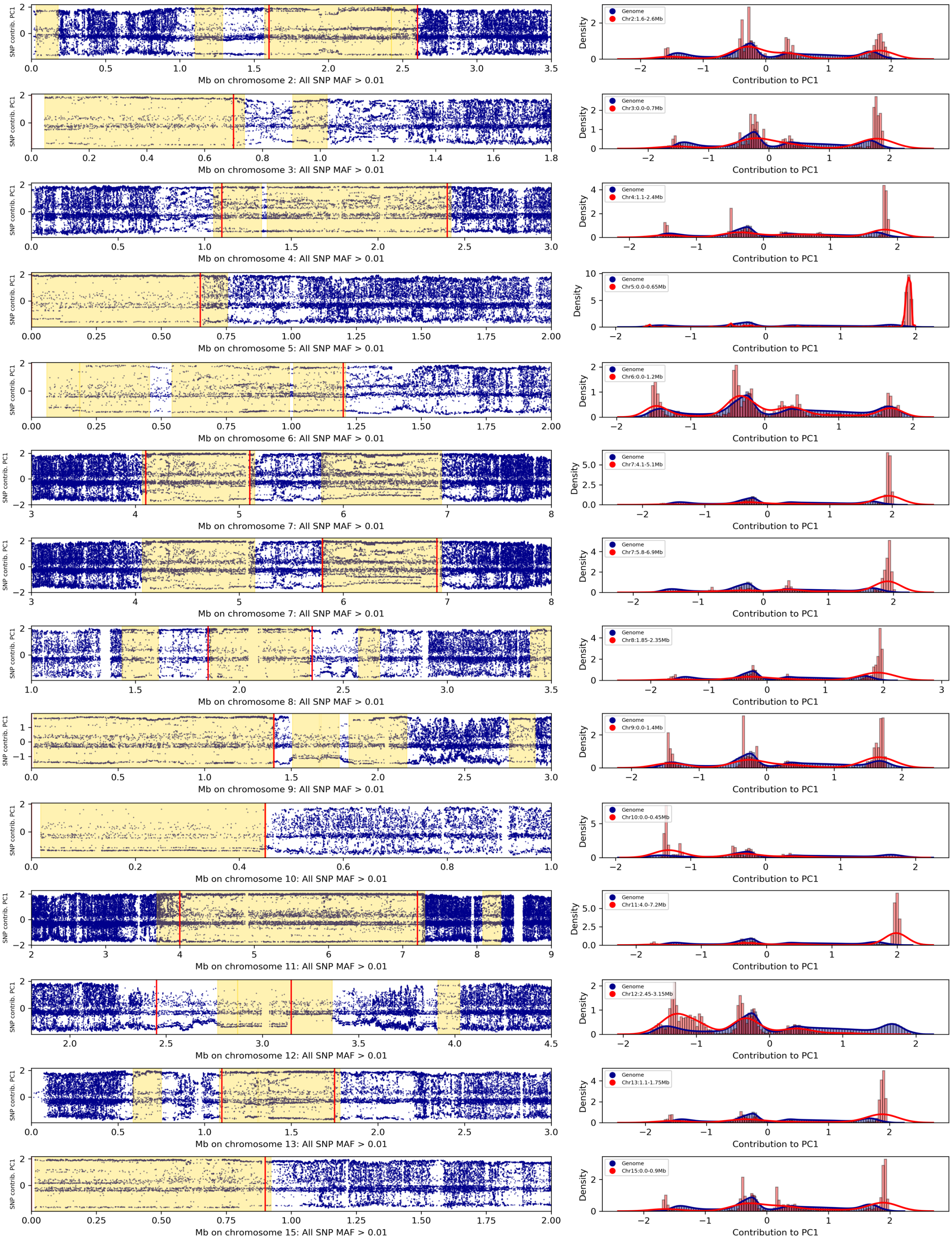


**Figure 12: Contribution of SNPs to PC1 in haplotype blocks.** Some of the most striking haplotype blocks are shown, showing their very strong contribution to PC1. Left: contribution of the individual SNPs to PC1. The yellow background indicates haplotype blocks of size larger than 100 kb, as detected by the block command of plink. The red vertical lines delimit the regions selected for plotting SNP contribution densities in the corresponding figures on the right. Right: density plots of SNP contributions to PC1; blue: SNPs from all the genome; red: SNPs from the selected region.

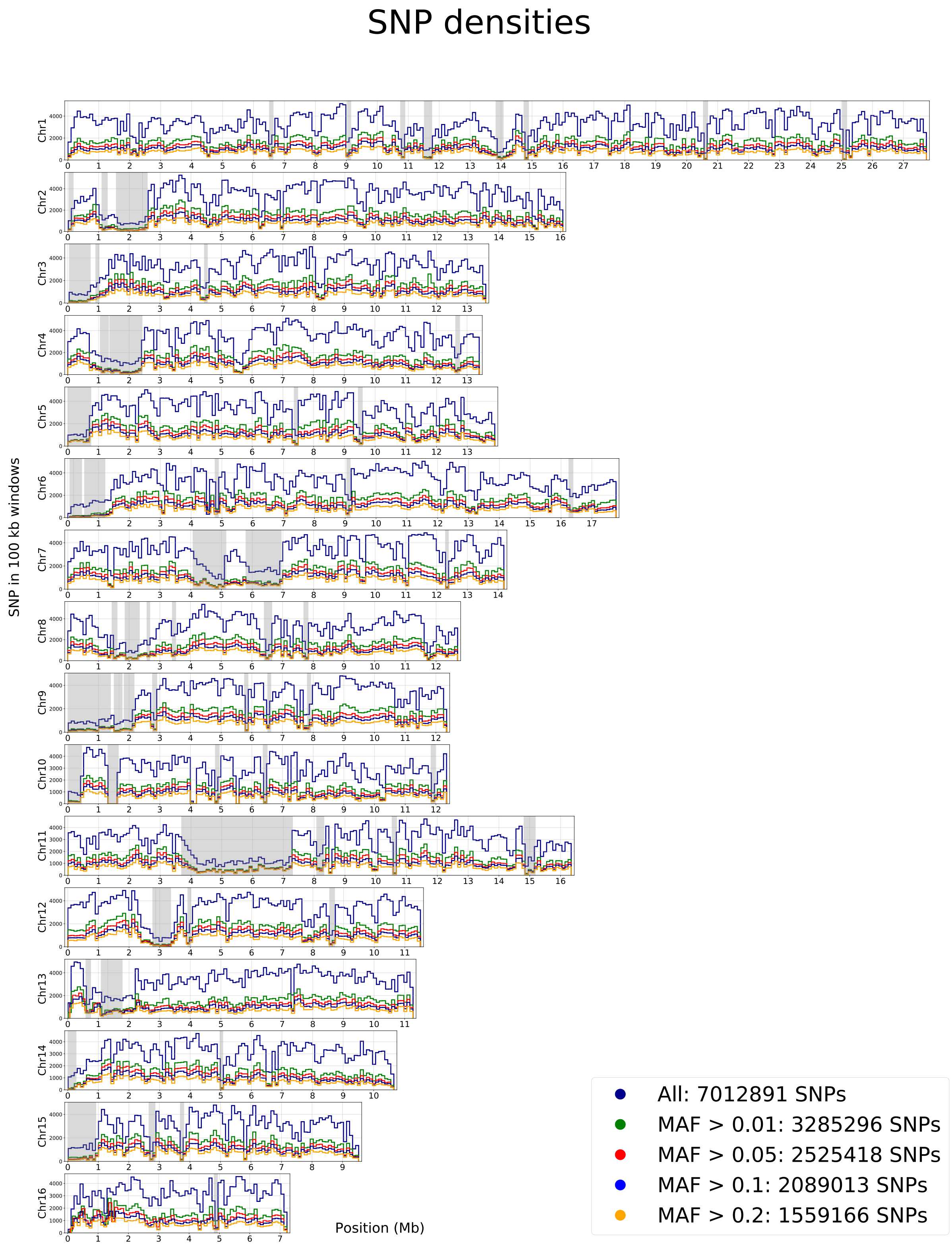


**Figure 13: Number of SNPs in 100 kb bins on all 16 chromosomes.** Plots represent SNP counts for all markers and after MAF filters. Grey backgrounds correspond to haplotype blocks detected with the plink blocks function, of size larger than 100 kb.

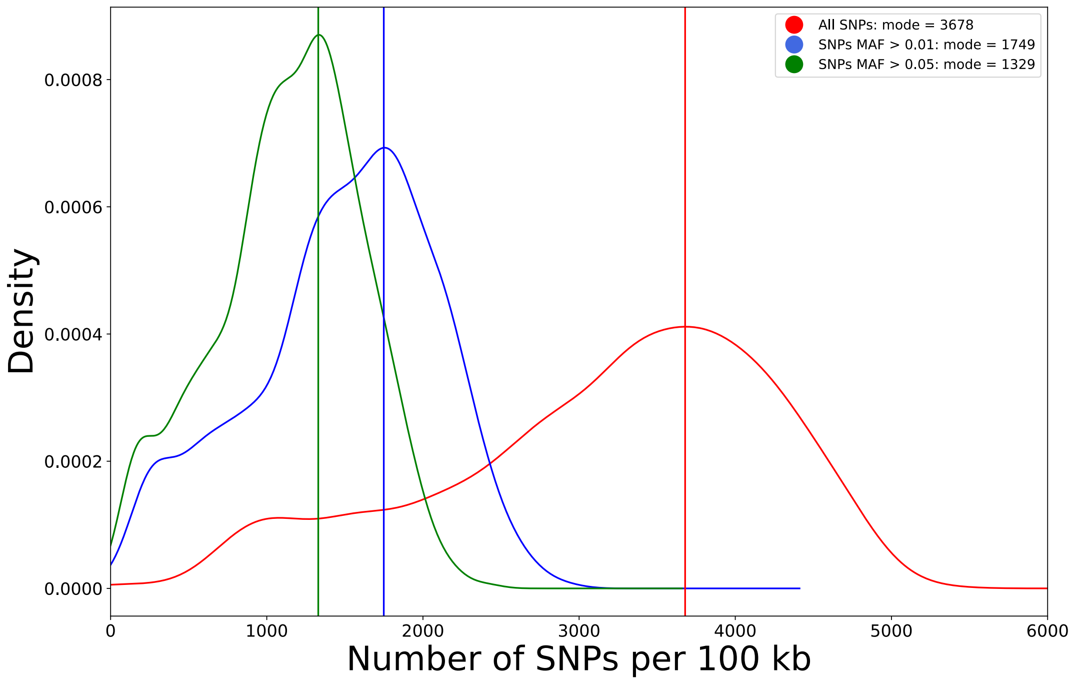


**Figure 14: Distribution of SNPs in 100 kb bins.**

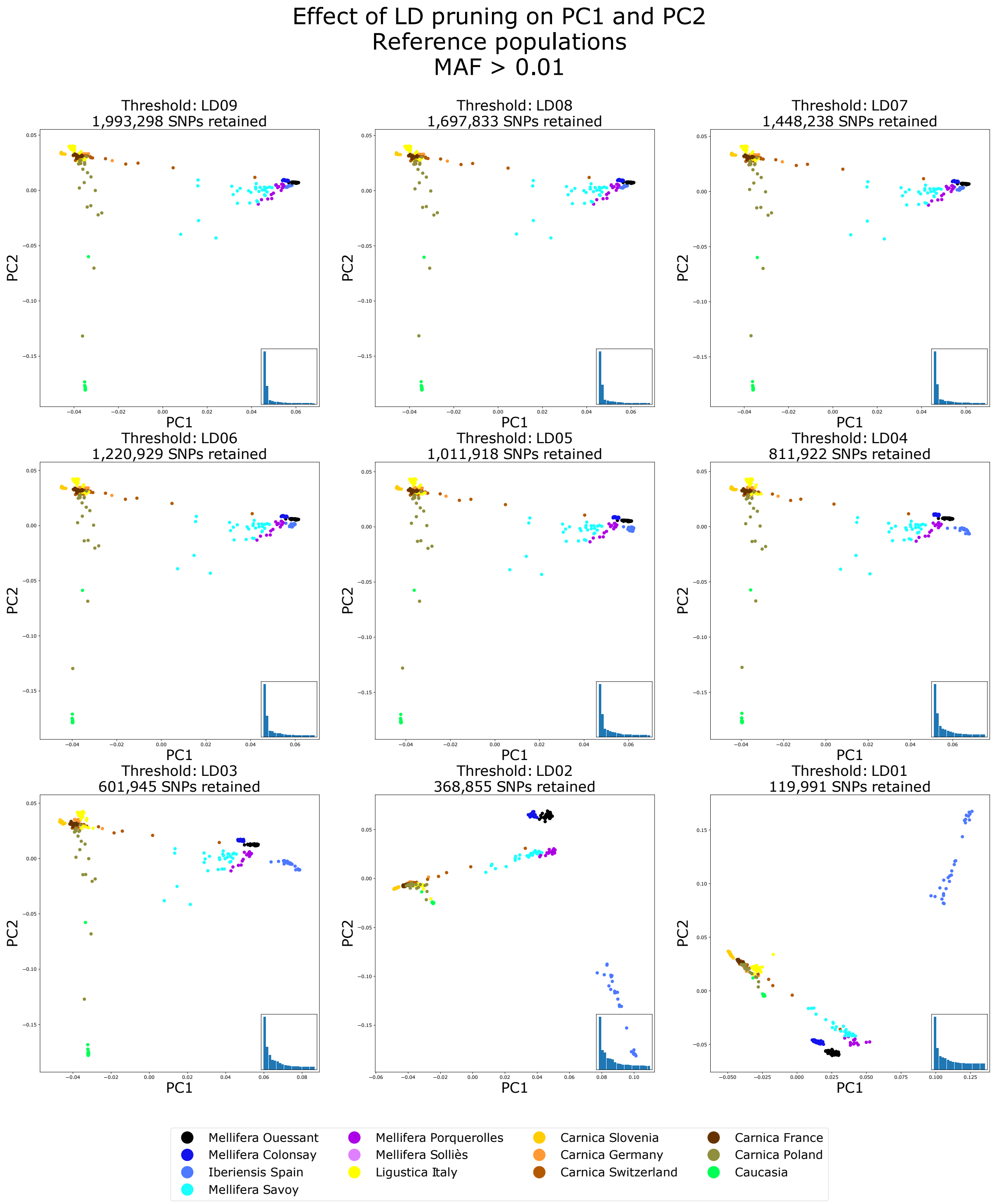


**Figure 15: Effect of LD pruning on PC1 and PC2.**
Only the reference populations are shown. PC1 and PC2 are plotted with datasets resulting from different LD pruning values. Down to LD = 0.3, the overall pattern is preserved, The blue barplots within the figures are the proportion of variance explained by the PCs 1 through 20.

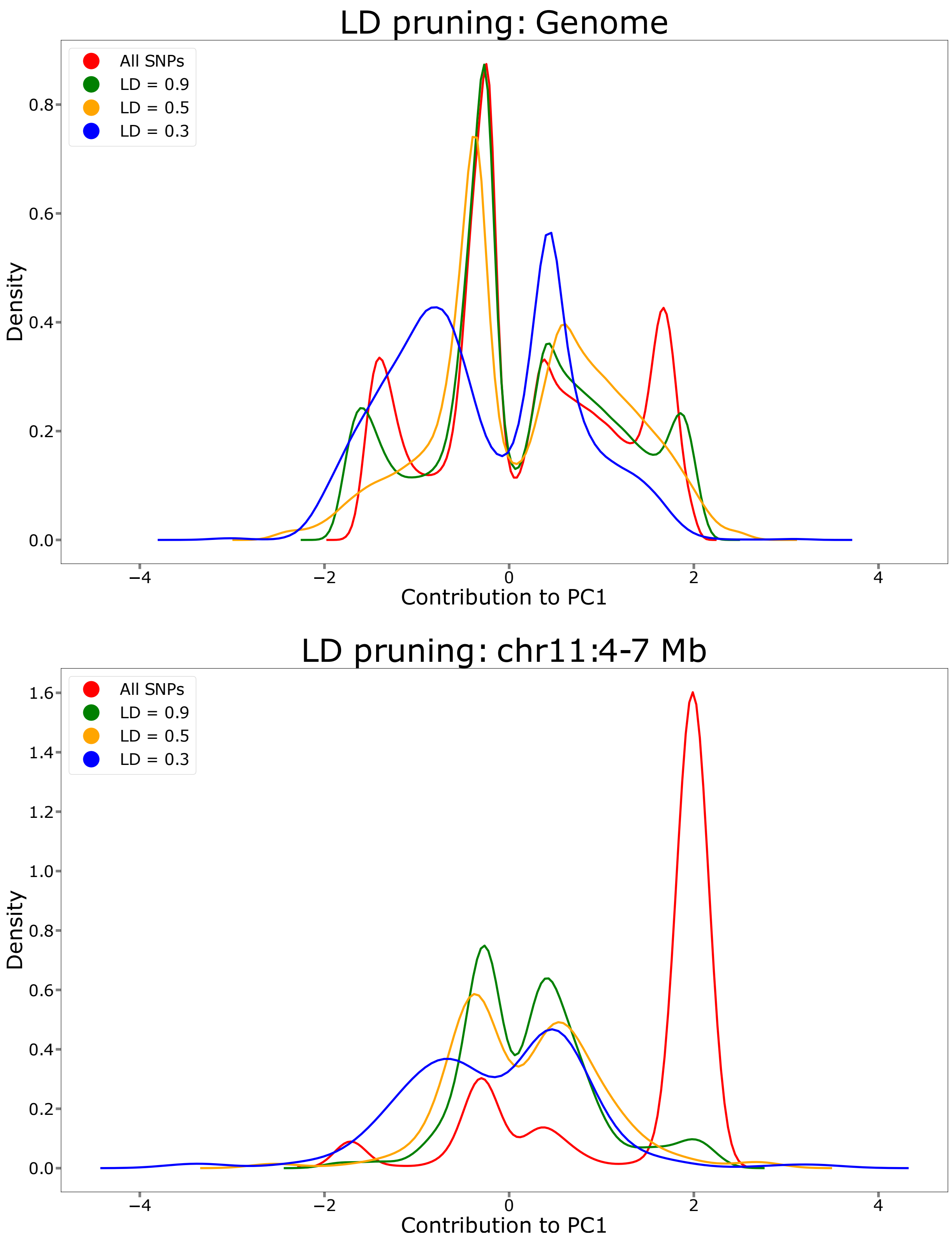


**Figure 16: Effect of LD pruning on SNP contribution to PC1** Top: on the whole genome; bottom: on the 3 Mb haplotype block on chromosome 11 having a very strong contribution to PC1. LD pruning allows to increase the genome wide proportion of markers contributing to the variance, while efficiently removing the excess of contributing markers in the haplotype block.

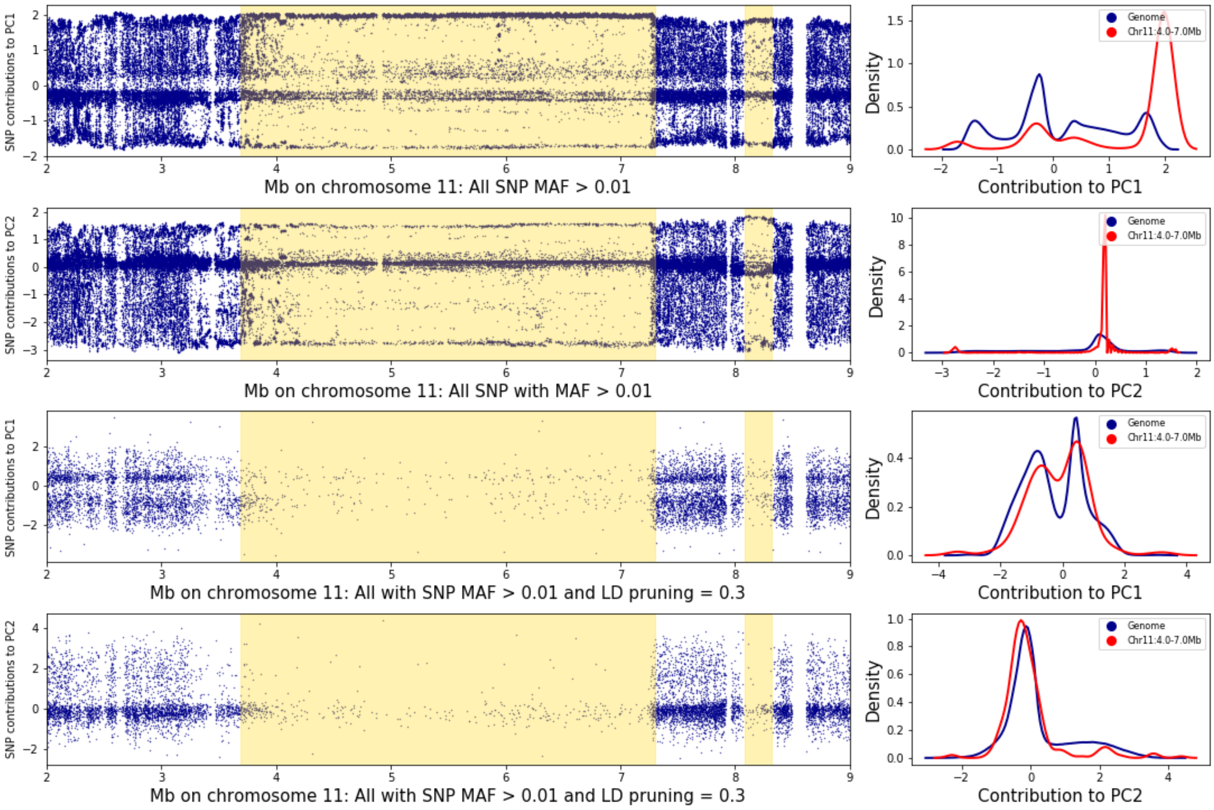


**Figure 17: Effect of LD pruning on SNP density in haplotype blocks.** Left: contribution of the individual SNPs to principal component 1 or 2 on chromosome 11, between positions 2 and 7 Mb before (top) or after LD pruning with LD = 0.3 (bottom); the yellow background indicates haplotype blocks of size larger than 100 kb, as detected by the block command of plink. Right: densities of the SNP contributions to PC1 and 2 with and without LD pruning; blue: SNPs from all the genome; red: SNPs from the haplotype block region detected with plink. The LD = 0.3 pruning value removes most markers from the haplotype block and the density distribution of SNP contributions within the haplotype block now matches that of the whole genome, for both PC1 and PC2.

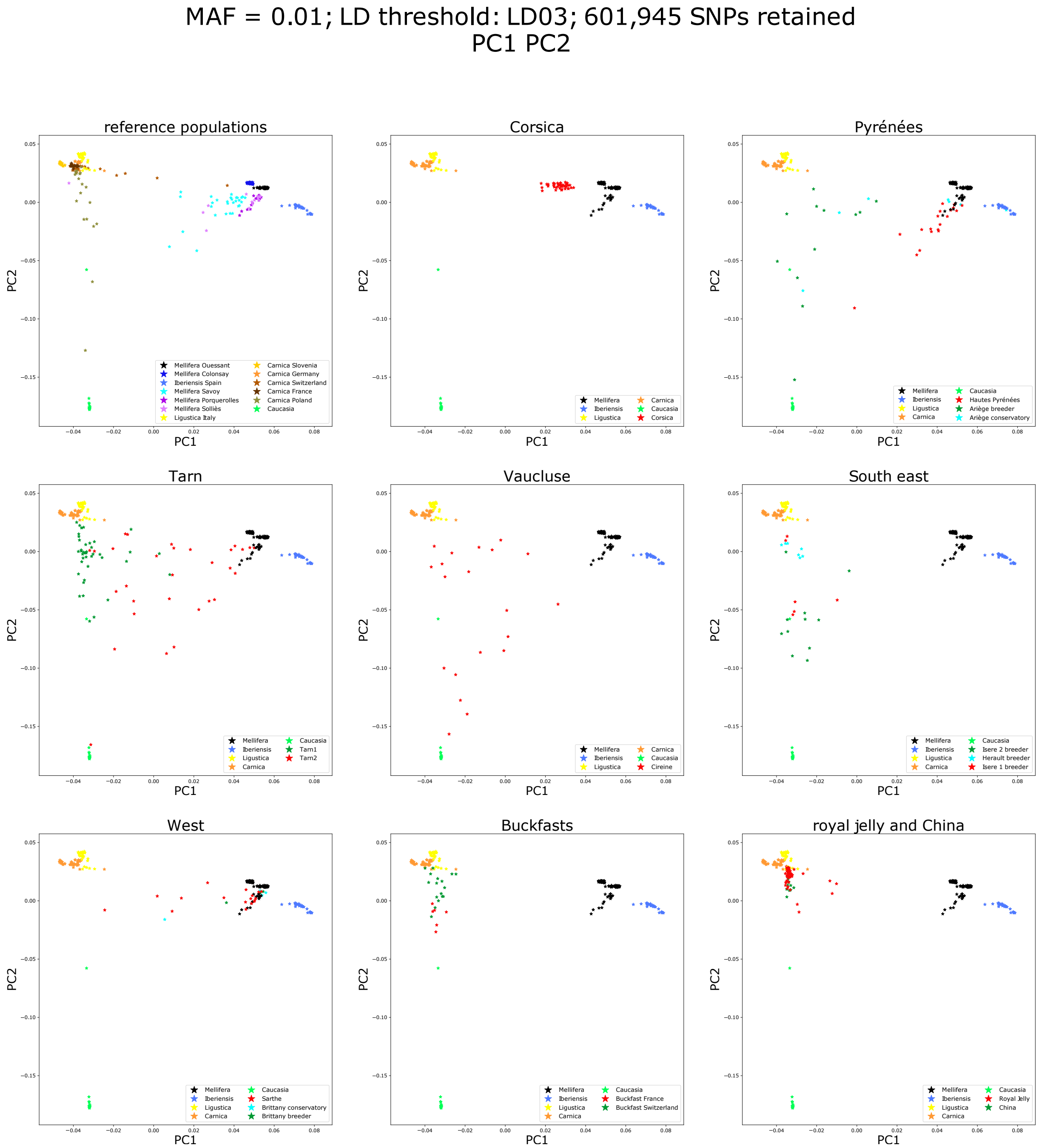


**Figure 18A: Principal component analysis with all populations – PC1 and PC2.**

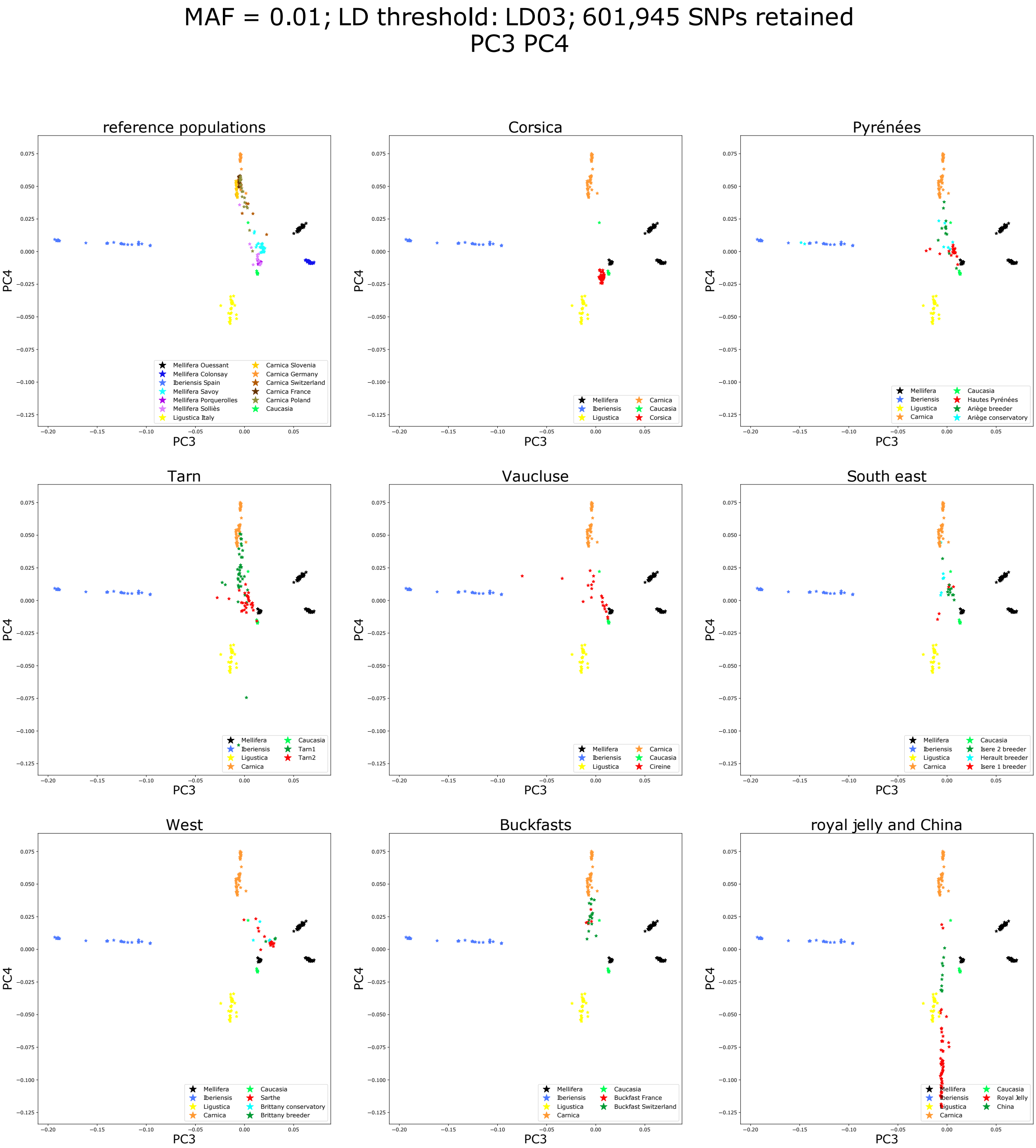


**Figure 18B: Principal component analysis with all populations – PC3 and PC4.**

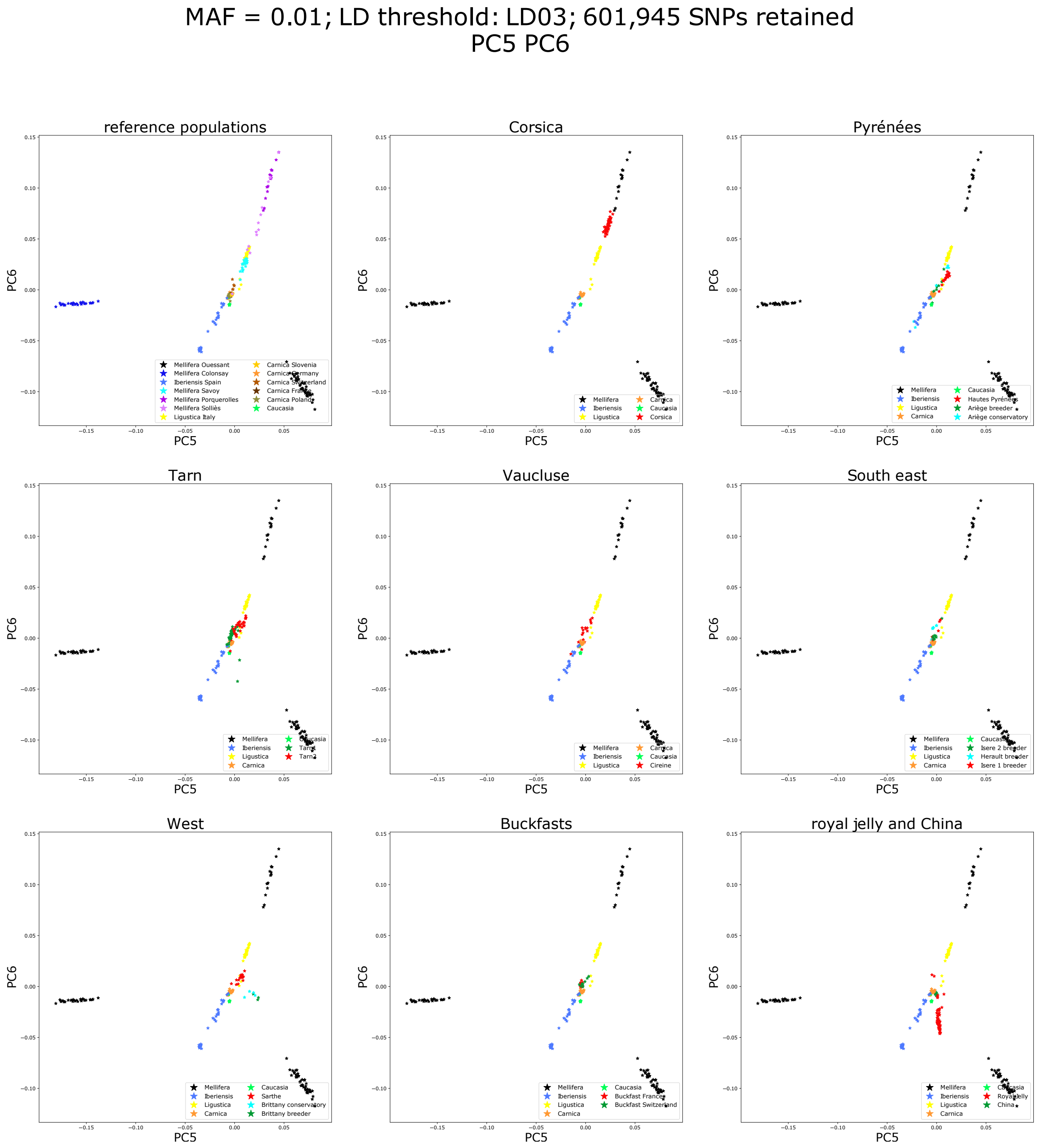


**Figure 18C: Principal component analysis with all populations – PC5 and PC6.**

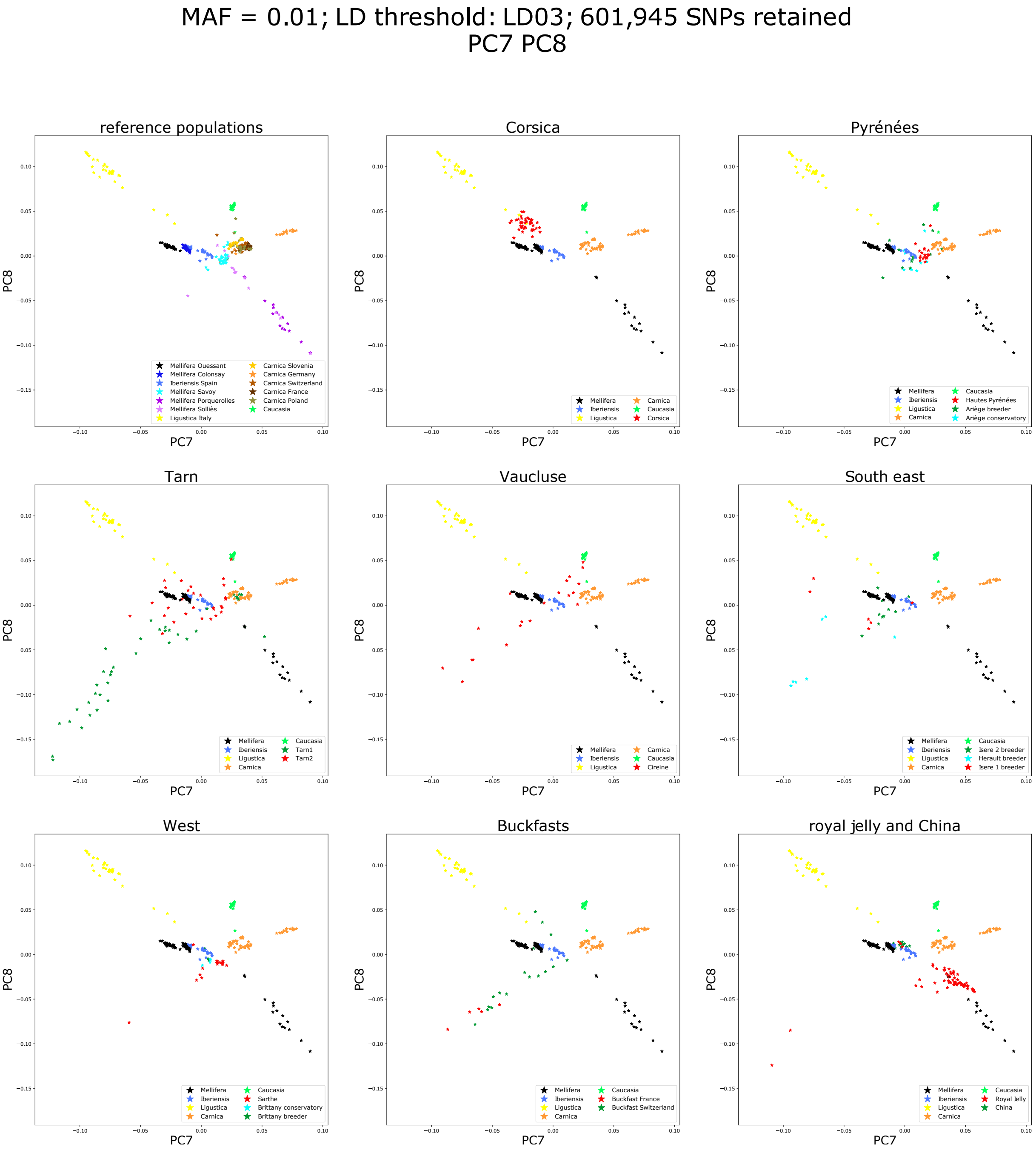


**Figure 18D: Principal component analysis with all populations – PC7 and PC8.**

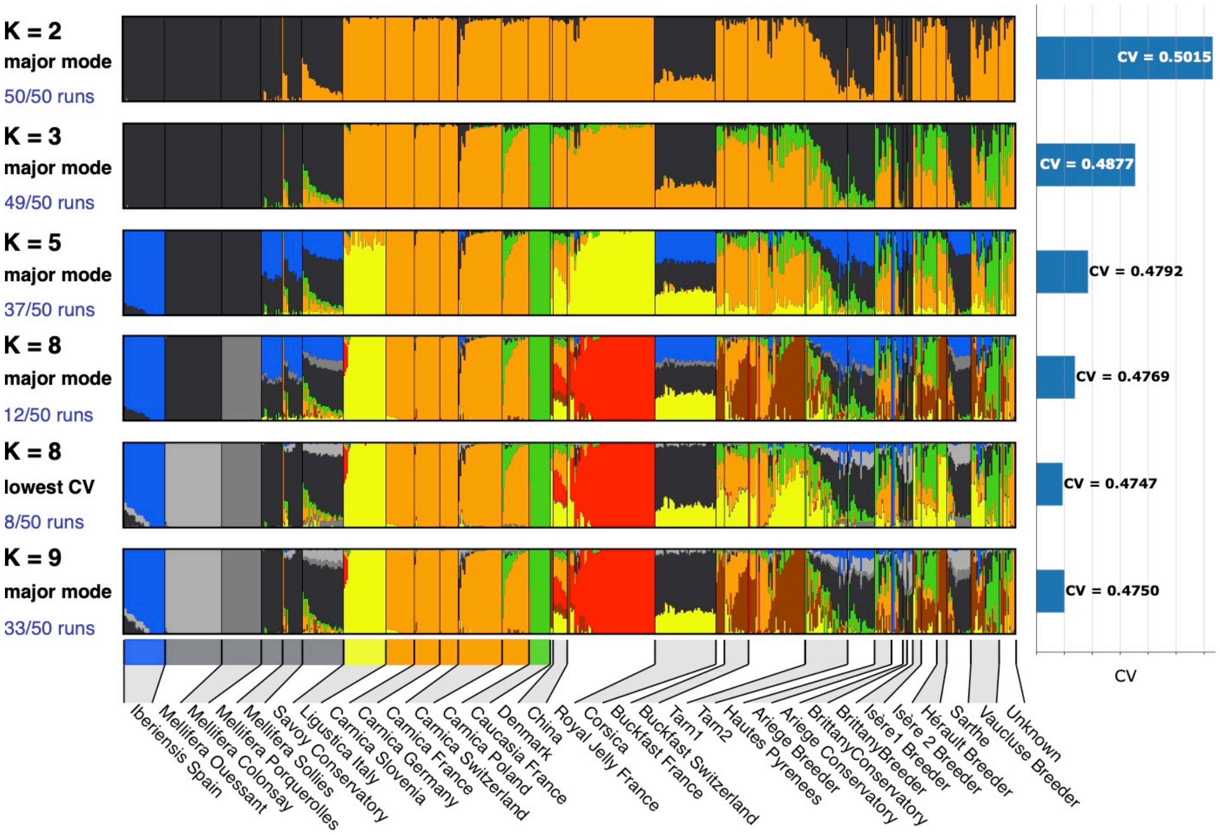


**Figure 19: remarkable results from Admixture runs.** Admixture patterns at K = 8 show how different runs of admixture can give slightly different results. The major mode (12 out of 50 runs) at K = 8 suggests the *A. m. mellifera* bees from mainland France in the black bee conservatories, as being hybrids between the populations from Ouessant and Spain, which does make any sense given the geography of Western Europe and our knowledge of the history of the bees in Ouessant. The 8 out of 50 runs at K = 8 are the ones with the lowest CV and are also the more likely based on prior knowledge. At K = 9, admixture runs converge better (33 out of 50 runs in the major mode) and a new background corresponding probably to the Buckfast bees appears.

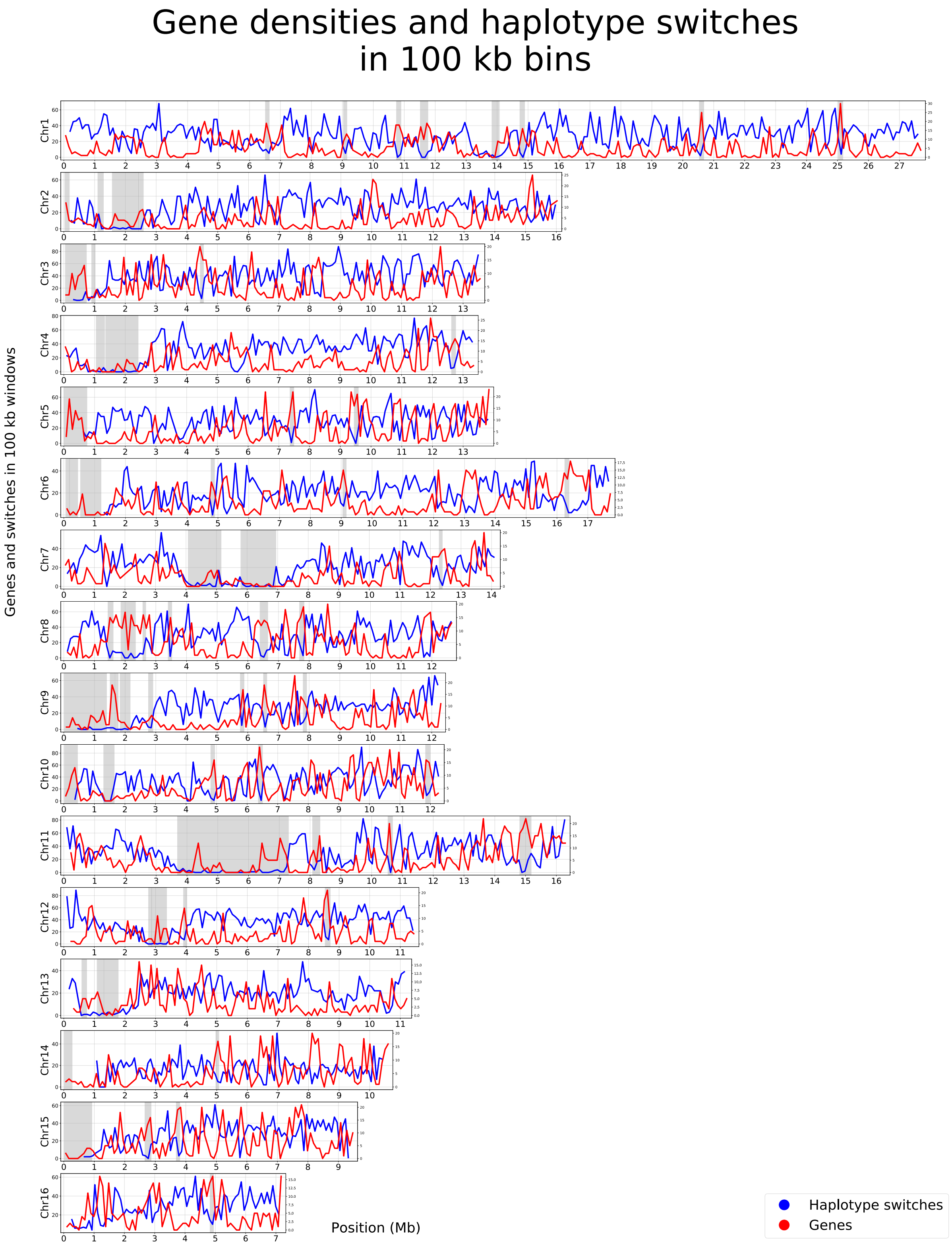


Figure 20: Haplotype switches and gene densities per 100 kb bins in all chromosomes. Grey backgrounds correspond to haplotype blocks detected with the plink blocks function, of size larger than 100 kb.
