## Supplementary methods 1 for "Complex population structure and haplotype patterns in Western Europe honey bee from sequencing a large panel of haploid drones"

### SeqApiPop analyses: from fastq files to vcf for 870 samples

The corresponding html document and scripts are also found in Github


- 1. Introduction

- [ ] (#)

- 2. Estimating mapped sequencing depth in BAM files
- 2. From fastq files to gvcf files
  - 2.1. Reference genome
  - 2.2. fastq files
    - 2.2.1. Availability
    - 2.2.2. Specific cases
      - 2.2.2.1. Sample BR1 has two library tags
      - 2.2.2.2. Samples sequenced in multiple lanes from one library
    - 2.2.3. Preparing for mapping
  - 2.3. Mapping: Calling script: Map_seqapipop_HAV3_1.bash
    - 2.3.1. Map_seqapipop_HAV3_1.bash
    - 2.3.2. PLOIDY variable
    - 2.3.3. N variable
    - 2.3.4. SAMPLE_FILE variable
    - 2.3.5. OUT variable
    - 2.3.6. REF variable
    - 2.3.7. CHROMOSOME variable
    - 2.3.8. UNKNOWN variable
    - 2.3.9. SCRIPT variable
    - 2.3.10. Output files
  - 2.4. Merging bams
    - 2.4.1. Preparing files and scripts
    - 2.4.2. List of samples to merge
    - 2.4.3. Merging: Calling script callMerging.sh
      - 2.4.3.1. callMerging.sh
      - 2.4.3.2. SAMPLE_FILE variable
      - 2.4.3.2. STARTPOINT variable
      - 2.4.3.3. Other variables
    - 2.4.3.4 Output files
    - 2.4.4. BQSR and calling on the merged bams
      - 2.4.4.1. Bootstrap_Call.bash script
      - 2.4.4.2. The SAMPLE_FILE variable
      - 2.4.4.2. Other variables
    - 2.4.4.3. Output files
    - 2.4.4.4. Same samples under different names.
  - 2.5 controlling that all gvcf files are complete
    - 2.5.1 Script controlVcfs.bash
    - 2.5.2. Check all went to the end
- 3. Estimating mapped sequencing depth in BAM files
- 4. Making the large vcf file with all 870 samples
  - 4.1 Combine gvcf files
  - 4.1.1. Generate list of all samples for the combine script
  - 4.1.2 Write the combining scripts
  - 4.1.3 Run the combine script with combineGVCFsHAV3_1_Lance_slurm.bash
  - 4.1.4. Output
  - 4.1.5. Check all went to the end
    - 4.1.5.1. Example for mitochondrial DNA

        - 4.1.5.2. For all chromosomes:
    - 4.2. Genotype across the Combined GVCF
        - 4.2.1. Genotype
        - 4.2.2. Check output
    - 4.3 Concatenate vcf files
    - 4.4. Count variants per chromosomes
    - 4.5 Retain only SNPs


### 1. Introduction

This document presents the analyses performed for obtaining the results presented in the paper *SeqApiPop* by Wragg et al (2020). Scripts and paths will have to be adapted to your local environment (SGE vs Slurm cluster; paths…)

The main scripts are in Scripts_1_MappingCalling.

Short scripts are directly described in the text.

Versions of the software used:

```
more program_module
#%Module1.0######################################################################
##

module load bioinfo/bwa-0.7.15
module load bioinfo/samtools-1.8
module load bioinfo/bedtools-2.27.1
module load bioinfo/tabix-0.2.5
module load bioinfo/vcftools-0.1.15
module load bioinfo/bcftools-1.6
module load bioinfo/picard-2.18.2
module load bioinfo/gatk-4.1.2.0
module load bioinfo/plink-v1.90b5.3
module load system/Python-3.6.3
module load system/Java8
module load system/R-3.4.3
```

### 2. Estimating mapped sequencing depth in BAM files

- The sequencing depth for all SeqApiPop samples was estimated with the Mosdepth software
- AllSeqApiPopBams.list is a list of paths to the bam files

```
#!/bin/bash

#DepthMosDepthSeqApiPop.bash

for i in `cat  ~/seqapipopOnHAV3_1/sequencingDepth/AllSeqApiPopBams.list`
do
        IFS='/' read -ra ARR1 <<< "$i"
        SAMPLE=${ARR1[8]}
        IFS="_" read -ra ARR2 <<< ${SAMPLE}
        NAME=${ARR2[0]}
        sbatch -J Depth --mem=2G --wrap="source /home/gencel/vignal/.bashrc;
                conda activate seqDepth;
                mosdepth -n  ${NAME} ${i};
                conda deactivate"
done
```

- for each bam, results are in a file BamName.mosdepth.summary.txt
- results were then aggregated. Two output files:

**allSummariesSeqApiPop.txt:**

- With Sample name, chromosome name, chromosome length, bases mapped, mean depth, minimum depth, maximum depth

| name | chrom | length | bases | mean | min | max |
|---|---:|---:|---:|---:|---:|---:|
| BS16-19-M2 | NC_037638.1 | 27754200 | 891809856 | 32.13 | 0 | 338224 |
| BS16-19-M2 | NC_037639.1 | 16089512 | 522408492 | 32.47 | 0 | 1864 |
| BS16-19-M2 | NC_037640.1 | 13619445 | 430268749 | 31.59 | 0 | 1481 |
| BS16-19-M2 | NC_037641.1 | 13404451 | 417610967 | 31.15 | 0 | 8049 |
| … | | | | | | |

**allSummariesSeqApiPop.txt:**

- With Sample name, mean chromosome (autosome) depth, mitochondrial DNA depth, mitochondrial / autosomal depths ratios.

| name | chrDepth | beeMitoDepth | beeMitoratio |
|---|---:|---:|---:|
| BS16-19-M2 | 31.29625 | 4500.92 | 143.8165914446619 |
| BS16-198-M1 | 33.228125 | 18661.06 | 561.6043637731591 |
| ITA7A | 17.368125 | 4261.76 | 245.37824318975137 |
| Sar21 | 19.418125 | 3597.02 | 185.24033602626412 |
| NCA35 | 12.55125 | 4693.06 | 373.91176177671554 |
| … | | | |

```
#!/usr/bin/env python3
# _*_ coding: Utf-8 _*_
# coding: utf-8

#aggregateSummariesSeqApiPop.py

import re
import glob
import csv

outFile = open("~/seqapipopOnHAV3_1/sequencingDepth/allSummariesSeqApiPop.txt",'w')
outFile2 = open("~/seqapipopOnHAV3_1/sequencingDepth/compareDepthsSeqApiPop.txt",'w')

titles = ["name","chrom", "length", "bases", "mean", "min", "max"]
print("\t".join(titles), file=outFile)
titles2 = ["name","chrDepth", "beeMitoDepth", "beeMitoratio"]
print("\t".join(titles2), file=outFile2)

for i in glob.glob("/work/project/cytogen/Alain/seqapipopOnHAV3_1/sequencingDepth/*summary*"):
    pathArray = re.split("/",i)
    nameLong = pathArray[7]
    nameArray = re.split("\.",nameLong)
    name = nameArray[0]
    with open(i) as csvFile:
        data=csv.reader(csvFile, delimiter = "\t")
        sumDepths = 0
        for row in data:
            if re.search('^NC',row[0]):
```

```
                out = [name, row[0],row[1], row[2], row[3], row[4],row[5]]
                print("\t".join(out), file=outFile)
                if row[0] == "NC_001566.1":
                    beeMitoDepth = float(row[3])
                    #print("BeeMito")
                else:
                    sumDepths = sumDepths + float(row[3])
                    #print("Chr")
        chrAverage = sumDepths / 16
        out2 = [name,str(chrAverage),str(beeMitoDepth),str(beeMitoDepth / chrAverage)]
        print("\t".join(out2), file=outFile2)
```

### 2. From fastq files to gvcf files

#### 2.1. Reference genome

The reference genome used is the Amel_HAv3.1, having Genbank and RefSeq assembly accessions GenBank assembly accessions
GCA_003254395.2 and GCF_003254395.2.

#### 2.2. fastq files

##### 2.2.1. Availability

The individual fastq files for the population analysis were produced by Illumlina sequencing machines. These raw sequence reads are available for download at NCBI within the SeqApiPop project PRJNA311274 and SRA runs of accession SRP069814.

Naming of sequence files is in the form "SampleId_LibraryTag_SequencingLane_Read1or2.fastq.gz". For instance:
OUE9_GTCCGC_L008_R1.fastq.gz

##### 2.2.2. Specific cases

###### 2.2.2.1. Sample BR1 has two library tags

- BR1_ATGTCA_L003
- BR1_CCGTCC_L001
  - These were considered as two different samples and renamed BR1A and BR1B

###### 2.2.2.2. Samples sequenced in multiple lanes from one library

For samples having low depth of coverage in a first run, the library was re-loaded for sequencing in a second run.
Samples sequenced in multiple lanes were on two lanes, except OUE8 that was on 3 lanes. Mapping was done separately and the bam files were merged after the mapping.

##### 2.2.3. Preparing for mapping

For each sample/run, there is one *_R1.fastq.gz and one *_R2.fastq.gz file with the forward and reverse reads. In our case, these are all in the directory *FastqFromNG6*.

#### 2.3. Mapping: Calling script: Map_seqapipop_HAV3_1.bash

The mapping scripts perform mapping with BWA, marking duplicate reads with Picard, GATK BQSR (Base Quality Score Recalibration) and variant calling with GATK HaplotypeCaller, thus producing individual *.gvcf files.

Map_seqapipop_HAV3_1.bash will take a list of paths to sample names and call:

mappingAV_2019_Dec.sh, that will perform the mapping and duplicate reads detection, then call:

bootstrapingAV_2019_Dec.sh, that will perform the Base Quality Score Recalibration on each chromosome separately, then call:

callingAV_2019_Dec.sh, that will merge chromosome bams into a genome vcf and do the genotyping.

##### 2.3.1. Map_seqapipop_HAV3_1.bash

```
#!/bin/bash

# Usage:        Map_seqapipop_HAV3_1.bash

N=1
PLOIDY=2

#--------------------------------------------------------------------
#EDIT FOR ACCESS TO SEQUENCE FILES
# First download links from NG6 to genotoul and update the path here
SAMPLE_FILE=/work/project/cytogen/Alain/seqapipopOnHAV3_1/O.list
OUT=/work/project/cytogen/Alain/seqapipopOnHAV3_1/Map_O
REF=/home/gencel/vignal/save/Genomes/Abeille/HAv3_1_indexes/GCF_003254395.2_Amel_HAv3.1_genomic.fna
CHROMOSOME=/home/gencel/vignal/save/Genomes/Abeille/HAv3_1_indexes/HAv3_1_Chromosomes.list
UNKNOWN=/home/gencel/vignal/save/Genomes/Abeille/HAv3_1_indexes/HAv3_1_Unknown.list
SCRIPT=/genphyse/cytogen/seqapipop/ScriptsHAV3Mapping/mapping_calling
#--------------------------------------------------------------------
mkdir -p ${OUT}/logs/mapping    # /work/ktabetaoul/Project_seqapipop/RESULTATS/logs/mapping
cd ${OUT}

#EDIT FOR mapping.sh
# This is the loop to send each sample to the cluster to be mapped
while read line
do
        ID=`basename ${line}`   # give the name of the sample-population to map
        IN=`dirname ${line}`    # give the path where find the sample fastq

        mkdir -p ${OUT}/${ID}

        sbatch --cpus-per-task=1 --mem-per-cpu=5G \
                -J ${ID}_mapping -o ${OUT}/logs/${ID}_mapping.o -e ${OUT}/logs/${ID}_mapping.e \
                ${SCRIPT}/mappingAV_2019_Dec.sh -s ${ID} -i ${IN} -o ${OUT}/${ID} \
                -p ${PLOIDY} -n ${N} -e ${OUT}/logs -R ${REF} -C ${CHROMOSOME} -U ${UNKNOWN}

done < ${SAMPLE_FILE}
# end of file
```

##### 2.3.2. PLOIDY variable

The variable PLOIDY used for specifying the HaplotypeCaller model to call variants is set here to POILDY=2 although the drones sequenced are haploid. This was done following first tests with the haploid model (PLOIDY=1), showing this will lead to false genotype calling in repeated sequences, especially VNTRs. Indeed, these will cause reads from different copies of a repeated sequence to map to a same position, creating false positive SNPs. In the haploid model, these will be given arbitrarily one of the possible alleles, with a high GQ (Genotype Quality), whereas in the diploid model, they will be caled as heterozygotes. These heterozygote positions can then easily be filterd out in subsequent steps.

##### 2.3.3. N variable

Was introduced for eventual sequencing of DNA pools of individuals. Here, we perform individual sequencing. Is thus set to N=1

##### 2.3.4. SAMPLE_FILE variable

Contains a list of samples to process, in the form:

```
$ head -3 O.list
/genphyse/cytogen/seqapipop/FastqFromNG6/OUE10_GTGAAA_L003
```

```
/genphyse/cytogen/seqapipop/FastqFromNG6/OUE10_GTGAAA_L008
/genphyse/cytogen/seqapipop/FastqFromNG6/OUE11_GTGGCC_L003
```

###### 2.3.5. OUT variable

Directory for the output.

###### 2.3.6. REF variable

Path to the reference genome, indexed for BWA

###### 2.3.7. CHROMOSOME variable

Path to the list of chromosomes and their length. List in the form:

```
$ more HAv3_1_Chromosomes.list
NC_037638.1     27754200
NC_037639.1     16089512
NC_037640.1     13619445
NC_037641.1     13404451
NC_037642.1     13896941
NC_037643.1     17789102
NC_037644.1     14198698
NC_037645.1     12717210
NC_037646.1     12354651
NC_037647.1     12360052
NC_037648.1     16352600
NC_037649.1     11514234
NC_037650.1     11279722
NC_037651.1     10670842
NC_037652.1     9534514
NC_037653.1     7238532
NC_001566.1     16343
```

###### 2.3.8. UNKNOWN variable

Path to the list of unknown and/or unplaced scaffolds and their length. List in the form:

```
head -3 HAv3_1_Unknown.list
NW_020555788.1  3988
NW_020555789.1  24043
NW_020555790.1  23881
```

###### 2.3.9. SCRIPT variable

Path to the directory containing the mapping, BQSR and calling scripts

- mappingAV_2019_Dec.sh : does the mapping and duplicate reads detection. Called by Map_seqapipop_HAV3_1.bash
- bootstrapingAV_2019_Dec.sh : does the BQSR. Called by mappingAV_2019_Dec.sh
- callingAV_2019_Dec.sh : does the HaplotypeCaller variant calling. Called by mappingAV_2019_Dec.sh

###### 2.3.10. Output files

For each sample, output files will be in a directory with the sample name (SAMPLE_NAME) in the form "SampleId_LibraryTag_SequencingLane", each containing 4 directories, with the bam and gvcf files:

- SAMPLE_NAME_sort.bam (before BQSR)
- SAMPLE_NAME.bam (after BQSR)

- SAMPLE_NAME.g.vcf.gz
- SAMPLE_NAME_dedup.metrics

For instance:

```
$ ls OUE9_GTCCGC_L008/*/
OUE9_GTCCGC_L008/bootstraping/:
OUE9_GTCCGC_L008.bam  OUE9_GTCCGC_L008.bam.bai  OUE9_GTCCGC_L008_bam.list

OUE9_GTCCGC_L008/calling/:
OUE9_GTCCGC_L008.g.vcf.gz  OUE9_GTCCGC_L008.g.vcf.gz.tbi

OUE9_GTCCGC_L008/mapping/:
OUE9_GTCCGC_L008_sort.bam  OUE9_GTCCGC_L008_sort.bam.bai

OUE9_GTCCGC_L008/metrics/:
OUE9_GTCCGC_L008_dedup.metrics
```

#### 2.4. Merging bams

##### 2.4.1. Preparing files and scripts

Some samples were sequenced in several runs. For each sample/run, there is one *_R1.fastq.gz and one *_R2.fastq.gz file with the forward and reverse reads. These are all in the directory *FastqFromNG6*.

##### 2.4.2. List of samples to merge

```
ls FastqFromNG6/*_R1.fastq.gz | awk 'BEGIN{FS="/"}{print $6}' | \
    awk 'BEGIN{FS=OFS="_"}{print $1,$2}' | sort | uniq -c | sort > ToMerge.list
awk '$1 > 1 {print $2}' ToMerge.list | awk 'BEGIN{FS="_"}{print $1}' > SamplesToMerge.list
```

The list is in the form:

```
$ head -3 SamplesToMerge.list
BER15
BER16
BER17
```

##### 2.4.3. Merging: Calling script callMerging.sh

The merging script will take a list of samples to merge, merge the corresponding bam files together and re-evaluate duplicate reads. Output will be merged bams in a directory structure identical to the standard mapping, except for the SequencinghLane identifier in "SampleId_LibraryTag_SequencingLane" replaced by "merged": ~/OUE9_GTCCGC_merged/mapping/OUE9_GTCCGC_merged_sort.bam. These will have to be run through the BQSR and calling processes with the script Bootstrap_Call.bash.

###### 2.4.3.1. callMerging.sh

The script callMerging.sh will call:

mergingGenologin.sh

```
#!/bin/bash

# Usage:        callMerging.sh
```

```
#-------------------------------------------------------------------
#EDIT FOR ACCESS TO BAM FILES
SAMPLE_FILE=/work/project/cytogen/Alain/seqapipopOnHAV3_1/Merging/SamplesToMerge.list
#Names in the form OUE8
STARTPOINT=/genphyse/cytogen/seqapipop/Data/Apis-mellifera/seqapipopOnHAV3_1/*/*/mapping
#Where the bam files are at the time : in sub-folders
OUT=/work/project/cytogen/Alain/seqapipopOnHAV3_1/Merging/MergedBams
REF=/home/gencel/vignal/save/Genomes/Abeille/HAv3_1_indexes/GCF_003254395.2_Amel_HAv3.1_genomic.fna
CHROMOSOME=/home/gencel/vignal/save/Genomes/Abeille/HAv3_1_indexes/HAv3_1_Chromosomes.list
UNKNOWN=/home/gencel/vignal/save/Genomes/Abeille/HAv3_1_indexes/HAv3_1_Unknown.list
SCRIPT=/genphyse/cytogen/seqapipop/ScriptsHAV3Mapping/mapping_calling
#-------------------------------------------------------------------

mkdir -p ${OUT}/logs/merging

while read sample
do

        BAM_LIST=(`ls ${STARTPOINT}/${sample}_*.bam`)
        DIR=`dirname ${BAM_LIST[0]}`
        IN=`basename ${BAM_LIST[0]}`
        SAMPLE=${IN%_*}
        SAMPLE_SHORT=${SAMPLE%_*}
        SAMPLE_MERGED=${SAMPLE_SHORT}_merged
        mkdir -p ${OUT}/${SAMPLE_MERGED}
        mkdir -p ${OUT}/${SAMPLE_MERGED}/mapping
        mkdir -p ${OUT}/${SAMPLE_MERGED}/metrics
        if [ -e ${OUT}/${SAMPLE_MERGED}/mapping/${sample}.list ]
        then
                rm ${OUT}/${SAMPLE_MERGED}/mapping/${sample}.list
        fi
        for i in ${BAM_LIST[@]:0:${#BAM_LIST[@]}}
        do
                echo ${i} >> ${OUT}/${SAMPLE_MERGED}/mapping/${sample}.list
        done
        mkdir -p ${OUT}/${SAMPLE_MERGED}
        mkdir -p ${OUT}/${SAMPLE_MERGED}/mapping
        mkdir -p ${OUT}/${SAMPLE_MERGED}/metrics
        sbatch --cpus-per-task=1 --mem-per-cpu=6G \
                -J ${sample}_merging \
                -o ${OUT}/logs/merging/${sample}_merging.o \
                -e ${OUT}/logs/merging/${sample}_merging.e \
                ${SCRIPT}/mergingGenologin.sh -b ${OUT}/${SAMPLE_MERGED}/mapping/ \
                ${sample}.list -o ${OUT} -s ${SAMPLE_MERGED} -t 1

done < ${SAMPLE_FILE}

#end of file
```

###### 2.4.3.2. SAMPLE_FILE variable

List of samples to merge, in the form:

```
$ head -3 SamplesToMerge.list
BER15
BER16
BER17
```

###### 2.4.3.2. STARTPOINT variable

Path to the bams to be merged. As bam files are each in ~/SAMPLE_NAME/mapping directories (see 2.3.10. Output FILES), the paths are here in the form ~/DIRECTORY/*/*/mapping

###### 2.4.3.3. Other variables

See 2.3.5. OUT variable and following

###### 2.4.3.4 Output files

For each sample, output files will be in a directory with the sample name (SAMPLE_NAME) in the form "SampleId_LibraryTag_merged" , each containing 1 directory, with the files:

- SAMPLE_NAME_sort.bam (before BQSR)

For instance:

```
$ ls OUE8_CCGTCC_merged/mapping
OUE8_CCGTCC_merged_sort.bam  OUE8_CCGTCC_merged_sort.bam.bai  OUE8.list
```

These will have to be run through the BQSR and calling processes with the script Bootstrap_Call.bash.

##### 2.4.4. BQSR and calling on the merged bams

###### 2.4.4.1. Bootstrap_Call.bash script

Bootstrap_Call.bash, takes a list of merged bam files and will call:

mappingAV_2019_Dec_RelanceBoots.sh, itself calling:

callingAV_2019_Dec.sh for the genotyping.

```
#!/bin/bash

# Usage:    Bootstrap_Call.bash

N=1
PLOIDY=2

#--------------------------------------------------------------------
#EDIT FOR ACCESS TO SEQUENCE FILES
SAMPLE_FILE=/work/project/cytogen/Alain/seqapipopOnHAV3_1/Merging/forBootstrapCalling.list
OUT=/work/project/cytogen/Alain/seqapipopOnHAV3_1/Merging/MergedBams
REF=/home/gencel/vignal/save/Genomes/Abeille/HAv3_1_indexes/GCF_003254395.2_Amel_HAv3.1_genomic.fna
CHROMOSOME=/home/gencel/vignal/save/Genomes/Abeille/HAv3_1_indexes/HAv3_1_Chromosomes.list
UNKNOWN=/home/gencel/vignal/save/Genomes/Abeille/HAv3_1_indexes/HAv3_1_Unknown.list
SCRIPT=/genphyse/cytogen/seqapipop/ScriptsHAV3Mapping/mapping_calling
#--------------------------------------------------------------------
mkdir -p ${OUT}/logs/mapping    # /work/ktabetaoul/Project_seqapipop/RESULTATS/logs/mapping
cd ${OUT}

#EDIT FOR mapping.sh
# This is the loop to send each sample to the cluster to be mapped
while read line
do
    ID=`basename ${line}`  # give the name of the sample-population to map
    IN=`dirname ${line}`   # give the path where find the sample fastq

    mkdir -p ${OUT}/${ID}

    sbatch --cpus-per-task=1 --mem-per-cpu=6G \
        -J ${ID}_mapping -o ${OUT}/logs/${ID}_mapping.o -e ${OUT}/logs/${ID}_mapping.e \
        ${SCRIPT}/mappingAV_2019_Dec_RelanceBoots.sh -s ${ID} -i ${IN} -o ${OUT}/${ID} \
        -p ${PLOIDY} -n ${N} -e ${OUT}/logs -R ${REF} -C ${CHROMOSOME} -U ${UNKNOWN}

done < ${SAMPLE_FILE}


# end of file
```

###### 2.4.4.2. The SAMPLE_FILE variable

A list in the form:

```
$ head -3 forBootstrapCalling.list
/work/project/cytogen/Alain/seqapipopOnHAV3_1/Merging/MergedBams/BER15_ATCACG_merged
/work/project/cytogen/Alain/seqapipopOnHAV3_1/Merging/MergedBams/BER16_CGATGT_merged
/work/project/cytogen/Alain/seqapipopOnHAV3_1/Merging/MergedBams/BER17_TTAGGC_merged
```

###### 2.4.4.2. Other variables

See 2.3.5. OUT variable and following

##### 2.4.4.3. Output files

For each sample, output files will be in a directory with the sample name (SAMPLE_NAME) in the form "SampleId_LibraryTag_merged", each containing 4 directories, with the bam and gvcf files:

- SAMPLE_NAME_sort.bam (before BQSR)
- SAMPLE_NAME.bam (after BQSR)
- SAMPLE_NAME.g.vcf.gz
- SAMPLE_NAME_dedup.metrics

For instance:

```
$ ls OUE8_CCGTCC_merged/*/
OUE8_CCGTCC_merged/bootstraping/:
OUE8_CCGTCC_merged.bam  OUE8_CCGTCC_merged.bam.bai  OUE8_CCGTCC_merged_bam.list

OUE8_CCGTCC_merged/calling/:
OUE8_CCGTCC_merged.g.vcf.gz  OUE8_CCGTCC_merged.g.vcf.gz.tbi

OUE8_CCGTCC_merged/mapping/:
OUE8_CCGTCC_merged_sort.bam  OUE8_CCGTCC_merged_sort.bam.bai  OUE8.list

OUE8_CCGTCC_merged/metrics/:
OUE8_CCGTCC_merged_dedup.metrics
```

##### 2.4.4.4. Same samples under different names.

JFM8 and JFM11 were used for testing a PCR-free library preparation protocol and are sequenced under different names:

- JFM8_ACTTGA_L001
- JFM8-PCRfree_TCGAAG_L001
- JFM11_GGCTAC_L001
- JFM11-PCRfree_TCGGCA_L001

This causes problems in the calling step, as the sample names are different (merging works, but message at the calling step saying that there are two samples in the bam).

Two specific scripts had to be used: mergingJFM8.sh and mergingJFM11.sh, followed by Bootstrap_Call.bash

#### 2.5 controlling that all gvcf files are complete

##### 2.5.1 Script controlVcfs.bash

controlVcfs.bash

- Counts the number of variants per chromosomes for each sample

- All directories for all 870 samples are in ~/Haploid

```
#!/bin/bash

#Usage: controlVcfs.bash

module load -f  /home/gencel/vignal/save/000_ProgramModules/program_module

for i in `ls /genphyse/cytogen/seqapipop/Data/Apis-mellifera/seqapipopOnHAV3_1/Haploid/*/calling/*.gz`
do
        ID=`basename ${i}`      # give the name of the sample-population to map
        IN=`dirname ${i}`
        sbatch -J test --mem=1G \
        --wrap="module load -f /home/gencel/vignal/save/000_ProgramModules/program_module; \
        bcftools query -f '%CHROM\n' ${i} | \
        grep ^NC | uniq -c > \
        /work/project/cytogen/Alain/seqapipopOnHAV3_1/controlVcfs/${ID}.count"
done
```

##### 2.5.2. Check all went to the end

- The last chromosome in the files is the mitochondrial DNA: NC_001566.1

```
$ grep NC_001566.1 *.count | wc -l
870
```

### 3. Estimating mapped sequencing depth in BAM files

- The sequencing depth for all SeqApiPop samples was estimated with the Mosdepth software
- AllSeqApiPopBams.list is a list of paths to the bam files

```
#!/bin/bash

#DepthMosDepthSeqApiPop.bash

for i in `cat  ~/seqapipopOnHAV3_1/sequencingDepth/AllSeqApiPopBams.list`
do
        IFS='/' read -ra ARR1 <<< "$i"
        SAMPLE=${ARR1[8]}
        IFS="_" read -ra ARR2 <<< ${SAMPLE}
        NAME=${ARR2[0]}
        sbatch -J Depth --mem=2G --wrap="source /home/gencel/vignal/.bashrc;
                conda activate seqDepth;
                mosdepth -n  ${NAME} ${i};
                conda deactivate"
done
```

- for each bam, results are in a file BamName.mosdepth.summary.txt
- results were then aggregated. Two output files:

**allSummariesSeqApiPop.txt:**

- With Sample name, chromosome name, chromosome length, bases mapped, mean depth, minimum depth, maximum depth

| name | chrom | length | bases | mean | min | max |
|---|---|---|---|---|---|---|
| BS16-19-M2 | NC_037638.1 | 27754200 | 891809856 | 32.13 | 0 | 338224 |

| name | chrom | length | bases | mean | min | max |
|---|---|---|---|---|---|---|
| BS16-19-M2 | NC_037639.1 | 16089512 | 522408492 | 32.47 | 0 | 1864 |
| BS16-19-M2 | NC_037640.1 | 13619445 | 430268749 | 31.59 | 0 | 1481 |
| BS16-19-M2 | NC_037641.1 | 13404451 | 417610967 | 31.15 | 0 | 8049 |
| … | | | | | | |

**allSummariesSeqApiPop.txt:**

- With Sample name, mean chromosome (autosome) depth, mitochondrial DNA depth, mitochondrial / autosomal depths ratios.

| name | chrDepth | beeMitoDepth | beeMitoratio |
|---|---|---|---|
| BS16-19-M2 | 31.29625 | 4500.92 | 143.8165914446619 |
| BS16-198-M1 | 33.228125 | 18661.06 | 561.6043637731591 |
| ITA7A | 17.368125 | 4261.76 | 245.37824318975137 |
| Sar21 | 19.418125 | 3597.02 | 185.24033602626412 |
| NCA35 | 12.55125 | 4693.06 | 373.91176177671554 |
| … | | | |

```
#!/usr/bin/env python3
# _*_ coding: Utf-8 _*_
# coding: utf-8

#aggregateSummariesSeqApiPop.py

import re
import glob
import csv

outFile = open("~/seqapipopOnHAV3_1/sequencingDepth/allSummariesSeqApiPop.txt",'w')
outFile2 = open("~/seqapipopOnHAV3_1/sequencingDepth/compareDepthsSeqApiPop.txt",'w')

titles = ["name","chrom", "length", "bases", "mean", "min", "max"]
print("\t".join(titles), file=outFile)
titles2 = ["name","chrDepth", "beeMitoDepth", "beeMitoratio"]
print("\t".join(titles2), file=outFile2)

for i in glob.glob("/work/project/cytogen/Alain/seqapipopOnHAV3_1/sequencingDepth/*summary*"):
    pathArray = re.split("/",i)
    nameLong = pathArray[7]
    nameArray = re.split("\.",nameLong)
    name = nameArray[0]
    with open(i) as csvFile:
        data=csv.reader(csvFile, delimiter = "\t")
        sumDepths = 0
        for row in data:
            if re.search('^NC',row[0]):
                out = [name, row[0],row[1], row[2], row[3], row[4],row[5]]
                print("\t".join(out), file=outFile)
                if row[0] == "NC_001566.1":
                    beeMitoDepth = float(row[3])
                    #print("BeeMito")
                else:
                    sumDepths = sumDepths + float(row[3])
                    #print("Chr")
```

```
        chrAverage = sumDepths / 16
        out2 = [name,str(chrAverage),str(beeMitoDepth),str(beeMitoDepth / chrAverage)]
        print("\t".join(out2), file=outFile2)
```

#### 4. Making the large vcf file with all 870 samples

In theory, the GATK CombineGVCFs accepts a list of samples, but I couldn't get it to work. Terefore, the bash script for combining all gvcf files will have all files listed in the form:

```
--variant ~/Haploid/AOC16_GTTTCG_L002/calling/AOC16_GTTTCG_L002.g.vcf.gz \
--variant ~/Haploid/AOC17_CGTACG_L002/calling/AOC17_CGTACG_L002.g.vcf.gz \
--variant ~/Haploid/AOC18_GAGTGG_L002/calling/AOC18_GAGTGG_L002.g.vcf.gz \
...
```

##### 4.1 Combine gvcf files

##### 4.1.1. Generate list of all samples for the combine script

- All directories for all 870 samples are in ~/Haploid

```
ls ~/Haploid/*/calling/* | grep vcf.gz$ | awk '{print "    --variant",$1,"\\"}' > temp.list
```

```
$ head -3 temp.list
--variant ~/Haploid/AOC16_GTTTCG_L002/calling/AOC16_GTTTCG_L002.g.vcf.gz \
--variant ~/Haploid/AOC17_CGTACG_L002/calling/AOC17_CGTACG_L002.g.vcf.gz \
--variant ~/Haploid/AOC18_GAGTGG_L002/calling/AOC18_GAGTGG_L002.g.vcf.gz \
```

##### 4.1.2 Write the combining scripts

> Done by combining Head (combineScriptHead) and Tail (combineScriptTail) files to temp.list

- Head

```
$ more combineScriptHead
#!/bin/bash

#combineGVCFsHAV3Called_slurm.bash

module load -f  /home/gencel/vignal/save/000_ProgramModules/program_module

while getopts ":c:" opt; do
  case $opt in
    c) CHR=${OPTARG};;
  esac
done

gatk --java-options "-Xmx80g" CombineGVCFs  \
        -R /home/gencel/vignal/save/Genomes/Abeille/HAv3_1_indexes/GCF_003254395.2_Amel_HAv3.1_genomic.fna \
        -L ${CHR} \
```

- Tail

```
$ more combineScriptTail
        -O /work/project/cytogen/Alain/seqapipopOnHAV3_1/combineGVCFs/The870vcf/MetaGenotypes${CHR}.g.vcf.gz

echo "Finnished: "`date`

#end of file
```

- Combining together

```
cat combineScriptHead ../temp.list combineScriptTail > combineGVCFsHAV3_1_Called_slurm.bash
```

##### 4.1.3 Run the combine script with combineGVCFsHAV3_1_Lance_slurm.bash

- The script will work in parallel by combining the gvcf files for each chromosome separately

```
#!/bin/bash

#combineGVCFsHAV3_1_Lance_slurm.bash

for i in `cat /home/gencel/vignal/save/Genomes/Abeille/HAv3_1_indexes/HAv3_1_Chromosomes.list | cut -f 1`
do
mkdir -p ~/seqapipopOnHAV3_1/combineGVCFs/The870vcf/logs

sbatch --cpus-per-task=1 --mem-per-cpu=100G \
        -J ${i}_genotAll \
        -o ~/seqapipopOnHAV3_1/combineGVCFs/The870vcf/logs/${i}_combine.o \
        -e ~/seqapipopOnHAV3_1/combineGVCFs/The870vcf/logs/${i}_combine.e \
        /seqapipopOnHAV3_1/combineGVCFs/The870vcf/combineGVCFsHAV3_1_Called_slurm.bash -c ${i}
done
```

##### 4.1.4. Output

- One file per chromosome: 16 autosomes, plus mitochondrial DNA, plus Unknown, in the form:
    - MetaGenotypesNC_001566.1.g.vcf.gz

##### 4.1.5. Check all went to the end

- The idea is to printout the last positions for each chromosome's gvcf file.

###### 4.1.5.1. Example for mitochondrial DNA

```
~/combineGVCFs/The870vcf $ bcftools query -f '%CHROM\t%POS\n' MetaGenotypesNC_001566.1.g.vcf.gz | tail
NC_001566.1     16331
NC_001566.1     16332
NC_001566.1     16333
NC_001566.1     16334
NC_001566.1     16335
NC_001566.1     16337
NC_001566.1     16338
NC_001566.1     16339
NC_001566.1     16342
NC_001566.1     16343
```

- OK, mitochondrial chromosome is 16471 bp long

###### 4.1.5.2. For all chromosomes:

- script combineCheck.bash

```
#!/bin/bash

#combineCheck.bash

module load -f  /home/gencel/vignal/save/000_ProgramModules/program_module

for i in `ls ~/seqapipopOnHAV3_1/combineGVCFs/The870vcf/*.gz`
do
        ID=`basename ${i}`
        OUT=${ID/gz}check

        echo  ${ID}
        echo ${ID/gz}check
        echo ${i}

sbatch --cpus-per-task=1 --mem-per-cpu=1G \
        --wrap="module load -f /home/gencel/vignal/save/000_ProgramModules/program_module; \
        bcftools query -f '%CHROM\t%POS\n' ${i} | tail  > \
        ~/Alain/seqapipopOnHAV3_1/combineGVCFs/The870vcf/checkCombines/${OUT}"

done
```

- Outputs the last 10 positions for each chromosome in a file with names in the form: MetaGenotypesNC_001566.1.g.vcf.check
- Positions of last variant for each chromosome:

```
$ for i in `ls *.check`; do tail -1 ${i}; done
NC_001566.1     16343
NC_037638.1     27752935
NC_037639.1     16088822
NC_037640.1     13618309
NC_037641.1     13404430
NC_037642.1     13896581
NC_037643.1     17789025
NC_037644.1     14198576
NC_037645.1     12716507
NC_037646.1     12354641
NC_037647.1     12360026
NC_037648.1     16352587
NC_037649.1     11514127
NC_037650.1     11279598
NC_037651.1     10669571
NC_037652.1     9533787
NC_037653.1     7237125
```

- Compare to chromosome lengths:

```
$ cat /home/gencel/vignal/save/Genomes/Abeille/HAv3_1_indexes/HAv3_1_Chromosomes.list
NC_037638.1     27754200
NC_037639.1     16089512
NC_037640.1     13619445
NC_037641.1     13404451
NC_037642.1     13896941
NC_037643.1     17789102
NC_037644.1     14198698
NC_037645.1     12717210
NC_037646.1     12354651
NC_037647.1     12360052
NC_037648.1     16352600
NC_037649.1     11514234
NC_037650.1     11279722
NC_037651.1     10670842
NC_037652.1     9534514
```

```
NC_037653.1      7238532
NC_001566.1      16343
```

#### 4.2. Genotype across the Combined GVCF

##### 4.2.1. Genotype

- Genotyping script genotypeGVCFsHAV3_1_Called_slurm.bash

```
#!/bin/bash

#genotypeGVCFsHAV3_1_Called_slurm.bash

module load -f  /home/gencel/vignal/save/000_ProgramModules/program_module

while getopts ":c:" opt; do
  case $opt in
    c) CHR=${OPTARG};;
  esac
done

gatk --java-options "-Xmx80g" GenotypeGVCFs  \
    -R /home/gencel/vignal/save/Genomes/Abeille/HAv3_1_indexes/GCF_003254395.2_Amel_HAv3.1_genomic.fna \
    -V /work/project/cytogen/Alain/seqapipopOnHAV3_1/combineGVCFs/The870vcf/MetaGenotypes${CHR}.g.vcf.gz \
    --use-new-qual-calculator \
    -O /work/project/cytogen/Alain/seqapipopOnHAV3_1/combineGVCFs/The870vcf/MetaGenotypesCalled${CHR}.vcf.gz

echo "Finnished: "`date`

#end of file
```

- Called by combineGVCFsHAV3_1_Lance_slurm.bash

```
#!/bin/bash

#genotypeGVCFsHAV3_1_Lance_slurm.bash

for i in `cat /home/gencel/vignal/save/Genomes/Abeille/HAv3_1_indexes/HAv3_1_Chromosomes.list | cut -f 1`

do
mkdir -p /work/project/cytogen/Alain/seqapipopOnHAV3_1/combineGVCFs/The870vcf/logsCalled

sbatch --cpus-per-task=1 --mem-per-cpu=100G \
    -J ${i}_genotAll \
    -o /work/project/cytogen/Alain/seqapipopOnHAV3_1/combineGVCFs/The870vcf/logsCalled/${i}_genotype.o \
    -e /work/project/cytogen/Alain/seqapipopOnHAV3_1/combineGVCFs/The870vcf/logsCalled/${i}_genotype.e \
    /work/project/cytogen/Alain/seqapipopOnHAV3_1/combineGVCFs/The870vcf/\
    genotypeGVCFsHAV3_1_Called_slurm.bash -c ${i}
done
```

##### 4.2.2. Check output

`$ grep complete *_genotype.e`

Should find a line for each chromosome

#### 4.3 Concatenate vcf files

```
#!/bin/bash

#concatenateVCFs.bash
```

```
module load -f  /home/gencel/vignal/save/000_ProgramModules/program_module

bcftools concat -f vcf.list -o MetaGenotypesCalled870.vcf.gz -O z

tabix MetaGenotypesCalled870.vcf.gz
```

#### 4.4. Count variants per chromosomes

```
#!/bin/bash
#statsVcfsRaw.bash
zcat MetaGenotypesCalled870.vcf.gz  | grep -v '#' | cut -f 1 | sort | uniq -c | \
    awk 'BEGIN {OFS="\t";sum=0}{print $2, $1; sum += $1} END {print "Sum", sum}' > countVcfSumRaw
```

```
$ more countVcfSumRaw
NC_001566.1     1510
NC_037638.1     1908476
NC_037639.1     1057375
NC_037640.1     982972
NC_037641.1     889447
NC_037642.1     953841
NC_037643.1     1203559
NC_037644.1     1032802
NC_037645.1     898131
NC_037646.1     808198
NC_037647.1     759176
NC_037648.1     976036
NC_037649.1     822893
NC_037650.1     808625
NC_037651.1     706809
NC_037652.1     627851
NC_037653.1     552873
Sum     14990574
```

- Almost 15 million variants!

#### 4.5 Retain only SNPs

```
#!/bin/bash

#retainSNPs.bash

module load -f  /home/gencel/vignal/save/000_ProgramModules/program_module

OUT=~/seqapipopOnHAV3_1/combineGVCFs/The870vcf

gatk --java-options "-Xmx64g" SelectVariants  \
    -R /home/gencel/vignal/save/Genomes/Abeille/HAv3_1_indexes/GCF_003254395.2_Amel_HAv3.1_genomic.fna \
    -V ${OUT}/MetaGenotypesCalled870.vcf.gz \
    --select-type-to-include SNP \
    -O ${OUT}/MetaGenotypesCalled870_raw_snps.vcf.gz

#Count the SNPs
zcat MetaGenotypesCalled870_raw_snps.vcf.gz  | grep -v '#' | cut -f 1 | sort | uniq -c | \
    awk 'BEGIN {OFS="\t";sum=0}{print $2, $1; sum += $1} END {print "Sum", sum}' > countVcfSumRawSNPs
```

```
$ more countVcfSumRawSNPs
NC_001566.1     1057
NC_037638.1     1351213
NC_037639.1     755896
NC_037640.1     685281
```

```
NC_037641.1     631431
NC_037642.1     676224
NC_037643.1     854365
NC_037644.1     725683
NC_037645.1     626552
NC_037646.1     572812
NC_037647.1     545884
NC_037648.1     689572
NC_037649.1     583414
NC_037650.1     580887
NC_037651.1     496859
NC_037652.1     441923
NC_037653.1     382401
Sum     10601454
```

- More than 10 million SNPs!
- The bee genome is 250 Gb long => ~one SNP per 25 bp!
