## Supplementary methods 2 for "Complex population structure and haplotype patterns in Western Europe honey bee from sequencing a large panel of haploid drones"

### SeqApiPop analyses: filter vcf file

The corresponding html document and scripts are also found in Github


- 1. introduction
- 2. Filters on annotations in the vcf file
    - 2.1. In the INFO field: general SNP quality estimations
    - 2.2. Sample level annotations: genotype quality estimations
- 3. Other filters
- 4. SCRIPTS for filtering
    - 4.1. General variables to edit in the calling script run_vcfcleanup.sh
    - 4.2. Variables to edit for quality filter threshold values :
    - 4.3. Variables to edit for type of run
    - 4.4. Parameters used in the study
        - 4.4.1 Plotting the diagnostic histograms
            - 4.4.1.1 Mapping quality metrics: Stand Odds Ratio (SOR)
            - 4.4.1.2 Mapping quality metrics: Fisher Strand (FS)
            - 4.4.1.3 Mapping quality metrics: Mapping Quality (MQ)
            - 4.4.1.4 Genotyping quality metrics: SNP quality (QUAL)
            - 4.4.1.5 Genotyping quality metrics: quality by depth (QD)
            - 4.4.1.6. Individual genotyping metrics: heterozygote calls
            - 4.4.1.7. Individual genotyping metrics: missing genotypes
            - 4.4.1.8. Individual genotyping metrics: genotype quality (QD)
            - 4.4.1.9 Notice on resulting missing data.
        - 4.4.2 Running the filters: generate Venn diagrams and filtered vcf
- 5. results:
- 6. Prepare bed, bim, fam files for plink, admixture, etc.
    - 6.1. change chromosome names to numbers
    - 6.2 Prepare bed, bim, fam files and filter on missing data for SNPs and samples
- 7. Plots on sequencing depth and missing genotypes


#### 1. introduction

On hard filtering variants

Variants were filtered on the INFO field and on samples-level annotations of the vcf. As we genotyped under a diploid model whereas sequenced haploid drones, SNPs with a high proportion of heterozygote calls were also filtered out. Finally, we removed the SNP markers having an additional allele noted *, which are indels (InDel), that can't be managed by subsequent plink analyses.

#### 2. Filters on annotations in the vcf file

The general marker INFO fields (FS, SOR, MQ…) and the sample level annotations that were analysed by plotting their distribution of values in the dataset and/or used for filtering are indicated in bold in the lists 2.1 and 2.2. Other annotations (DP, AC, AF) are indicated for reference.

MQRankSum and ReadPosRankSum (italics): were finally not used in the filters, as suggested by the distribution of their values on the plots (see histograms and ECDF in Figures_S1_VcfCleanup and below).

##### 2.1. In the INFO field: general SNP quality estimations

- **FS** = FisherStrand; phred-scaled probability that there is strand bias at the site.

- **SOR** = StrandOddsRatio: another way to estimate strand bias using a test similar to the symmetric odds ratio test.

- **MQ** = RMSMappingQuality: root mean square mapping quality over all the reads at the site.

- ***MQRankSum*** = MappingQualityRankSumTest: compares the mapping qualities of the reads supporting the reference allele and the alternate allele.

- ***ReadPosRankSum*** = ReadPosRankSumTest: compares whether the positions of the reference and alternate alleles are different within the reads.

- **QUAL** = Phred-scaled quality score for the assertion made in ALT. The more samples have the ATL allele, the higher the QUAL score

- DP = In the INFO field: combined depth across samples

- **QD** = QUAL score normalized by allele depth (AD). For a single sample, the HaplotypeCaller calculates the QD by taking QUAL/AD. For multiple samples, HaplotypeCaller and GenotypeGVCFs calculate the QD by taking QUAL/AD of samples with a non hom-ref genotype call. The reason we leave out the samples with a hom-ref call is to not penalize the QUAL for the other samples with the variant call. QD is roughly speaking QUAL / Sum(Sum(AD)). More complex for InDels.

- AC = Allele count in genotypes, for each ALT allele, in the same order as listed

- AF = allele frequency for each ALT allele in the same order as listed: use this when estimated from primary data, not called genotypes.

##### 2.2. Sample level annotations: genotype quality estimations

- GT = Genotype
- AD = Allele Depth. Only informative reads counted. Used for calculating QD (see INFO fields). Sum of AD can be inferior to DP.
- DP = depth for the given sample.
- PGT = Phased Genotype.
- PID = The PID contains the first site in the phased sites. For example, if sites 1,2,3 are phased, the PIDs for all the 3 sites will contain 1. See discussion about read-backed phasing.
- **GQ** = Genotyping Quality: difference between the second lowest PL and the lowest PL (which is always 0.
- PL = : normalized Phred-scaled likelihoods of the genotypes considered in the variant record for each sample.
- PS : phase set. A phase set is defined as a set of phased genotypes to which this genotype belongs.

#### 3. Other filters

- SNPs with more than 3 alleles were filtered out (**variable limit_allele=3**)
- As we sequenced haploid drones, SNPs with a high proportion of heterozygote calls (**variable limit_het=0.01**) were filtered out. Some heterozygote calls (< 1%) had to be retained to avoid loosing too many markers. These were probably genotypiong errors and were set to missing.

#### 4. SCRIPTS for filtering

- run_vcfcleanup.sh, will call the script:

  - vcf_cleanup.sh, which will call the scripts:
  - diagnostic.r. Will output:
    - histogram and ecdf plots for the distribution of the various quality estimators in the input vcf.
    - Values set for filtering in run_vcfcleanup.sh will be indicated on the plots
  - filter.r. Will output:
    - Venn diagrams

        - the list of SNPs to keep: list_kept.txt.
            - Any SNP marker having > 3 alleles or one of the alleles being an indel (noted '*' in the ALT field ofthe vcf) will be removed by the script.
            - list_kept.txt will be used by vcf_cleanup.sh to produce the filtered vcf.
        - the number of SNPs in the input and the output vcfs will be counted.
    - filter_list.r. Will output:
        - Filters will be run, producing intermediate vcf files and the number of SNPs will be counted counted for each vcf file.
        - Will remove any SNP marker having > 3 alleles or one of the alleles being an indel (noted '*' in the ALT field of the vcf)
    - count_phased_geno.py. Will output:
        - count_phased_geno.txt : number of unphased, phased and missing genotype calls for each variant of the input vcf

- According to the type of run: 'diagnostic', 'filter_all' or 'filter_sequential', only the diagnostic plots will be produced or the plots plus the filtering.

#### 4.1. General variables to edit in the calling script run_vcfcleanup.sh

- paths, number of authorised alleles, etc.
- All editing of paths and values for filters are done in this script
    - username=avignal # Deprecated
    - SCRIPTS='~/seqapipopOnHAV3_1/vcf_cleanup_scripts' #path to the other scripts called
    - DIRIN='~/seqapipopOnHAV3_1/combineGVCFs/The870vcf' #path to directory containing the input vcf
    - DIROUT='~/seqapipopOnHAV3_1/vcf_cleanup' #path to output directory
    - VCFIN='MetaGenotypesCalled870_raw_snps.vcf.gz' #name of the vcf file to filter
    - limit_allele=3 #accept up to three alleles (edit to 2 or 4)

#### 4.2. Variables to edit for quality filter threshold values :

The variables limit_FS, limit_SOR … limit_het can either be set to a specified value, or set to -999, in which case each filter threshold will be calculated such as a percentage of the data, specified in the variables quantile_prob_above_threshold and quantile_prob_below_threshold, will be kept.

- limit_FS=61
- limit_SOR=4
- limit_MQ=39
- limit_MQRankSum=-12.5
- limit_ReadPosRankSum=-8
- limit_QUAL=200
- limit_QD=20
- limit_GQ=10
- limit_miss=0.05
- limit_het=0.01
- limit_GQfiltered=0.2
- quantile_prob_above_threshold=0.1
    - In this example, any of the variables above for which variants are kept above a threshold is set to -999, will be adjusted to eliminate 10 % of the data.
- quantile_prob_below_threshold=0.9
    - In this example, any of the variables above for which variants are kept below a threshold is set to -999, will be adjusted to eliminate 10 % of the data.
- kept_above_threshold="MQ_QUAL_QD~GQ~GQ"
- kept_below_threshold="FS_SOR_allele~miss_het~GQfiltered"
    - kept_above_threshold and kept_below_threshold: variables are separated by "_" and groups of variables by "~". The number of groups of variables must be the same, hence GQ in two groups for kept_above_threshold

#### 4.3. Variables to edit for type of run

- The variable #run can take the values: 'diagnostic', 'filter_all' or 'filter_sequential'
  - diagnostic: will only output the distribution plots for each quality parameter, to help decide on threshold values setting.
  - filter_all: will filter the vcf file using all parameters set by the variables simultaneously, as specified in the kept_above_threshold and kept_below_threshold variables
  - filter_sequential: filter on each parameter set by the variables, but sequencially.

#### 4.4. Parameters used in the study

##### 4.4.1 Plotting the diagnostic histograms

A run with run='diagnostic' will plot histograms and empirical cumulative distribution functions (ECDF), without performing the actual filtering. Once satisfactory filtering values are obtained, a second run='filter_all' will perform the filtering and produce the Venn diagrams.

```
#! /bin/bash

#run_vcf_cleanup.sh

#vcf filter (only done once to prepare list SNP positions)

# modules #####################################################
module load system/R-3.5.1
module load bioinfo/bcftools-1.6
module load bioinfo/tabix-0.2.5
module load bioinfo/vcftools-0.1.15
module load bioinfo/samtools-1.8
# end modules #################################################

# parameters ##################################################
username=avignal
SCRIPTS='~/seqapipopOnHAV3_1/vcf_cleanup_scripts' #path to scripts
DIRIN='~/combineGVCFs/The870vcf' #path to directory containing the input vcf
DIROUT='~/seqapipopOnHAV3_1/vcf_cleanup' #path to output directory
VCFIN='MetaGenotypesCalled870_raw_snps.vcf.gz' #inputvcf before filters
limit_FS=61
limit_SOR=4
limit_MQ=39
limit_MQRankSum=-12.5
limit_ReadPosRankSum=-8
limit_QUAL=200
limit_QD=20
limit_GQ=10
limit_miss=0.05
limit_het=0.01
limit_GQfiltered=0.2
quantile_prob_above_threshold=0.1
quantile_prob_below_threshold=0.9
kept_above_threshold="MQ_QUAL_QD~GQ~GQ"
kept_below_threshold="FS_SOR_allele~miss_het~GQfiltered"
run='diagnostic'
# end parameters ##############################################
sbatch -W -J vcf_cleanup -o ${DIROUT}/log/vcf_cleanup.o -e ${DIROUT}/log/vcf_cleanup.e \
        --wrap="${SCRIPTS}/vcf_cleanup.sh ${username} ${SCRIPTS} ${DIRIN} ${DIROUT} ${VCFIN} \
        ${limit_allele} ${limit_FS} ${limit_SOR} ${limit_MQ} ${limit_MQRankSum} ${limit_ReadPosRankSum} \
        ${limit_QUAL} ${limit_QD} ${limit_GQ} ${limit_miss} ${limit_het} ${limit_GQfiltered} \
        ${quantile_prob_above_threshold} ${quantile_prob_below_threshold} \
        ${kept_above_threshold} ${kept_below_threshold} ${run}"

# end of file
```

###### 4.4.1.1 Mapping quality metrics: Stand Odds Ratio (SOR)

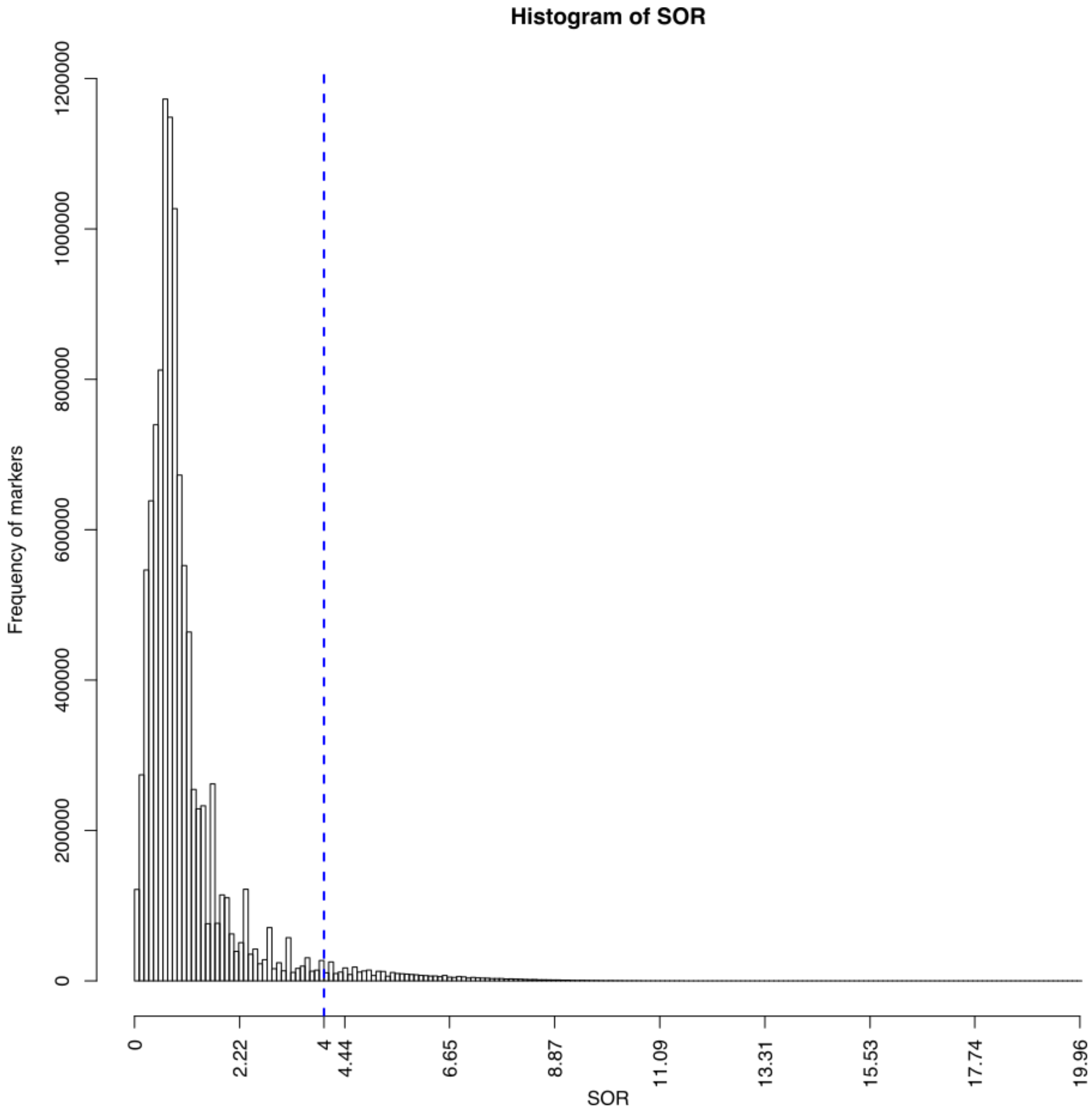


Counts of SNPs according to SOR values. The blue dotted line indicates the threshold retained for filtering the vcf: SOR > 4.

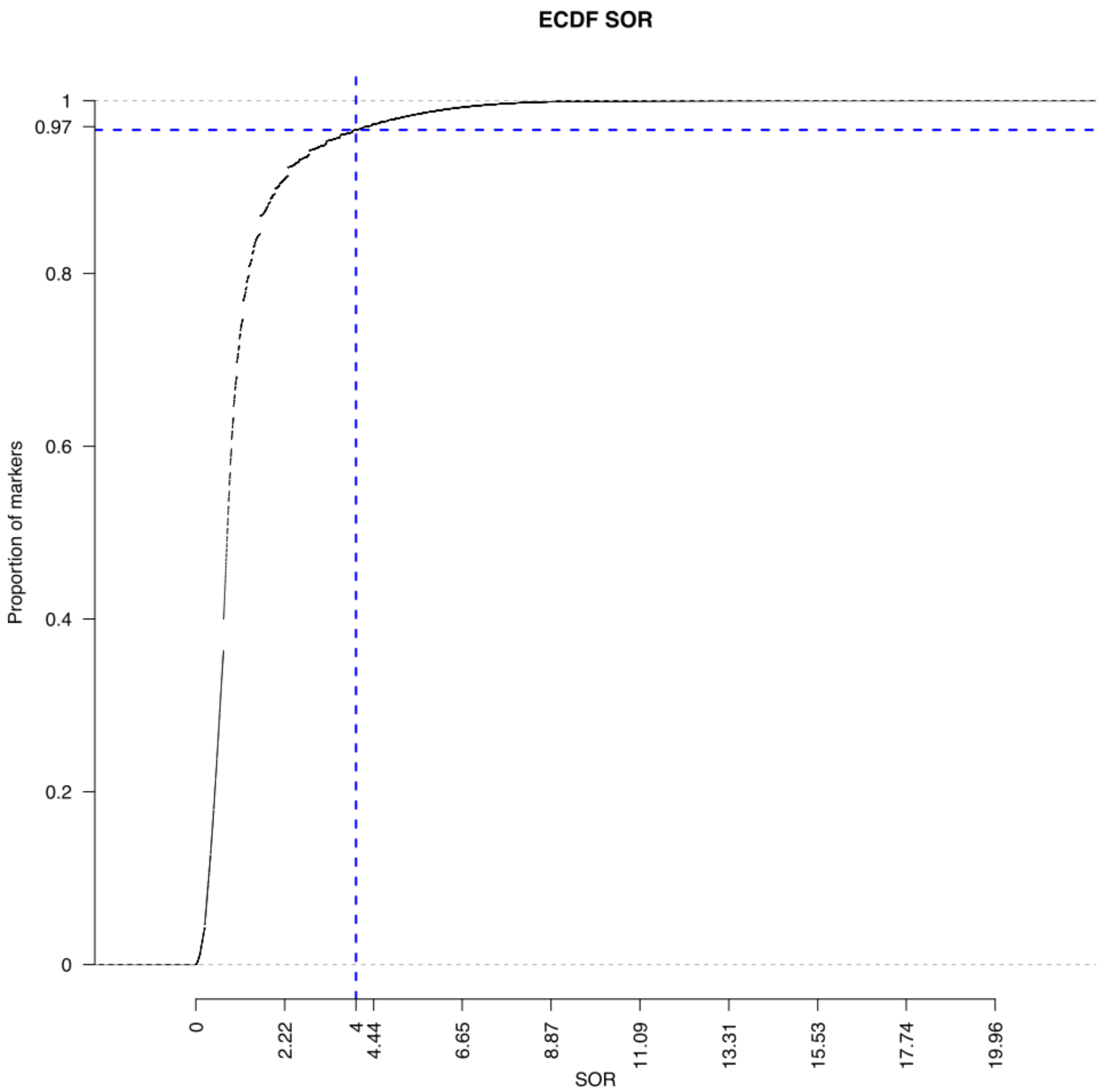

Empirical cumulative distribution function of SOR values. The blue dotted line indicates the threshold used for filtering the vcf: SOR > 4.

###### 4.4.1.2 Mapping quality metrics: Fisher Strand (FS)

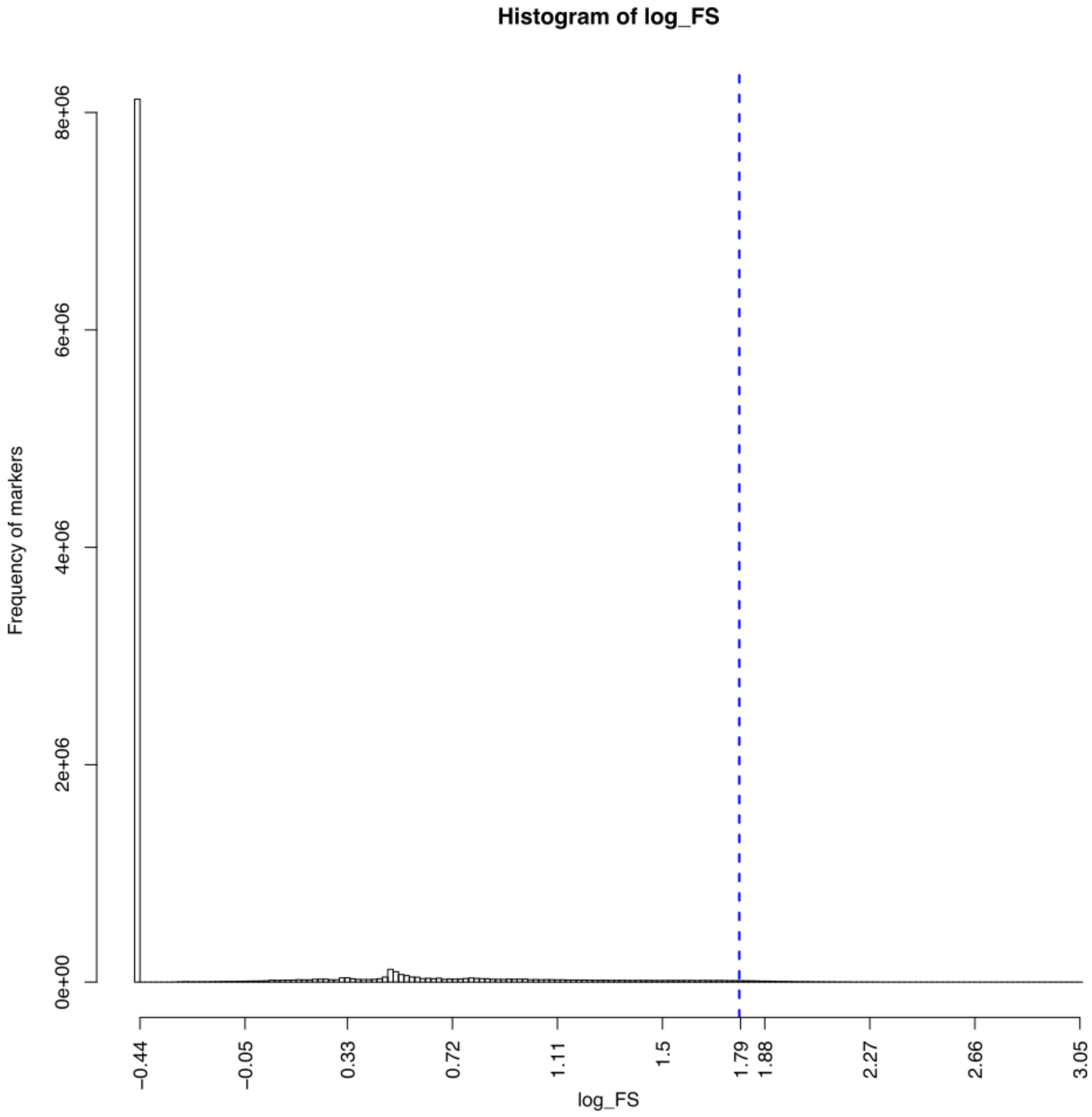


Counts of SNPs according to FS values. The blue dotted line indicates the threshold used for filtering the vcf: FS < 61. X axis is on a log scale log(61)=1.785.

###### 4.4.1.3 Mapping quality metrics: Mapping Quality (MQ)

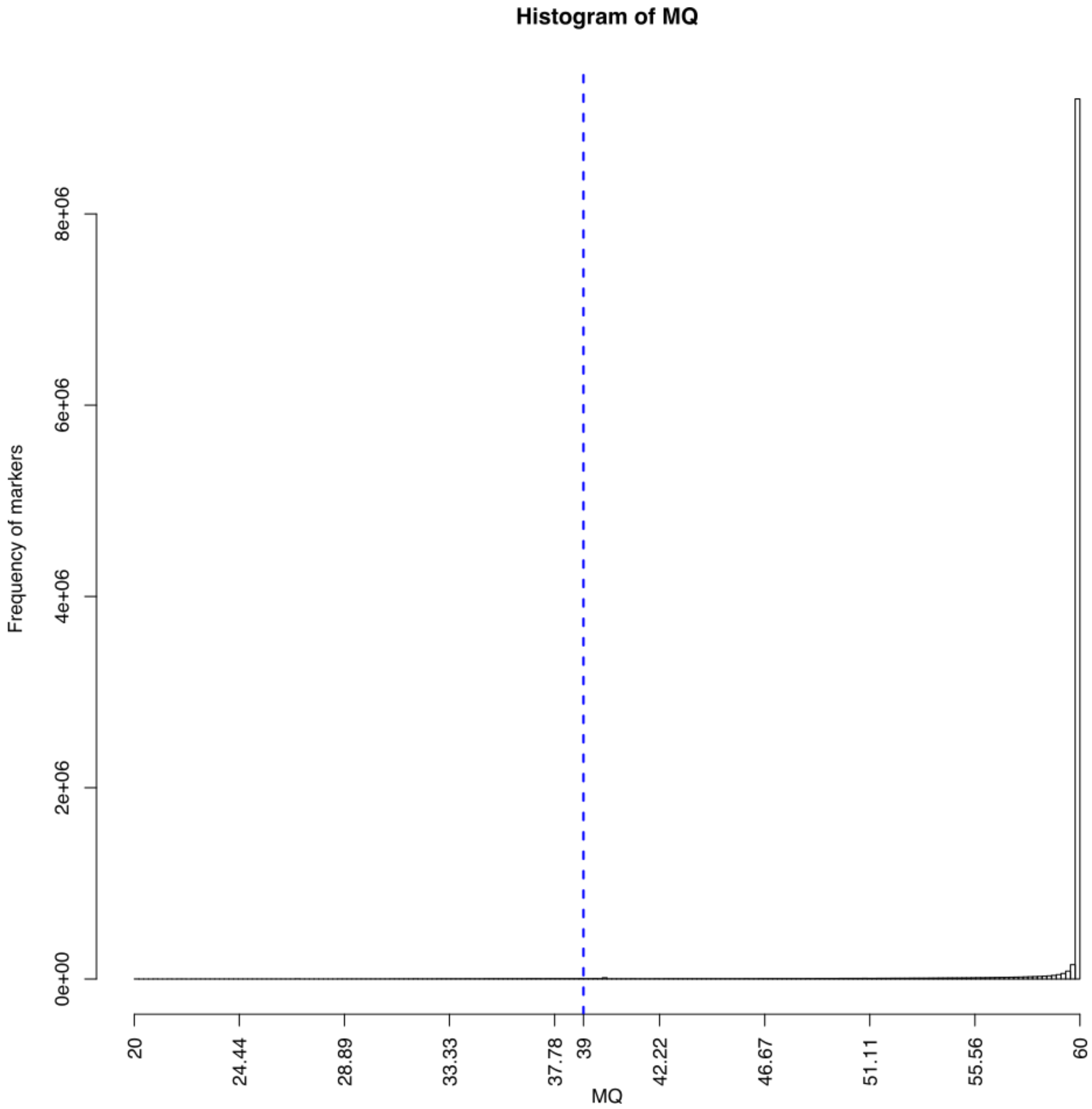


Counts of SNPs according to MQ values. The blue dotted line indicates the threshold used for filtering the vcf: MQ > 40.

###### 4.4.1.4 Genotyping quality metrics: SNP quality (QUAL)

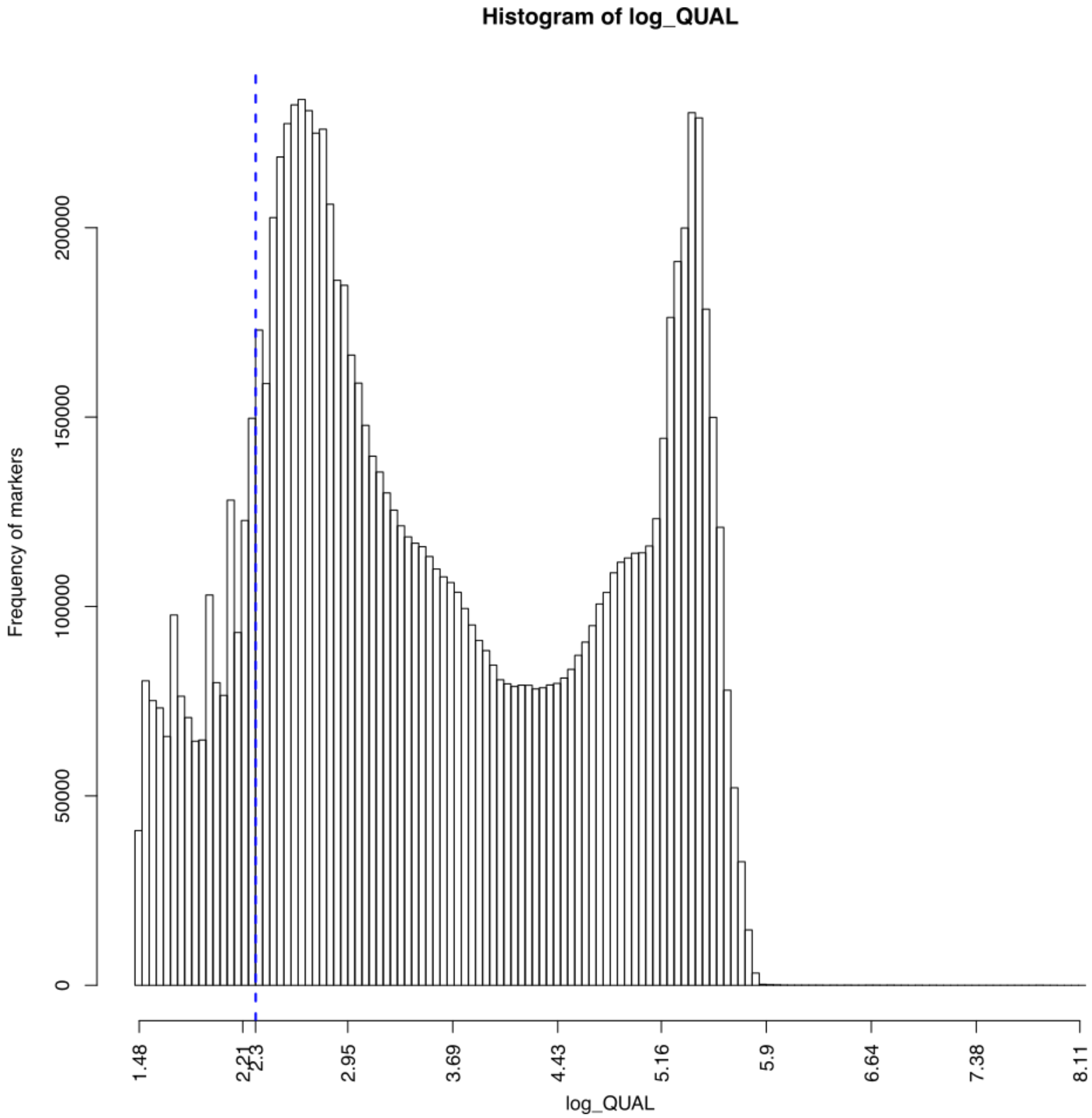


Counts of SNPs according to QUAL values. The blue dotted line indicates the threshold used for filtering the vcf: QUAL > 200. X axis is on a log scale log(200)=2.3.

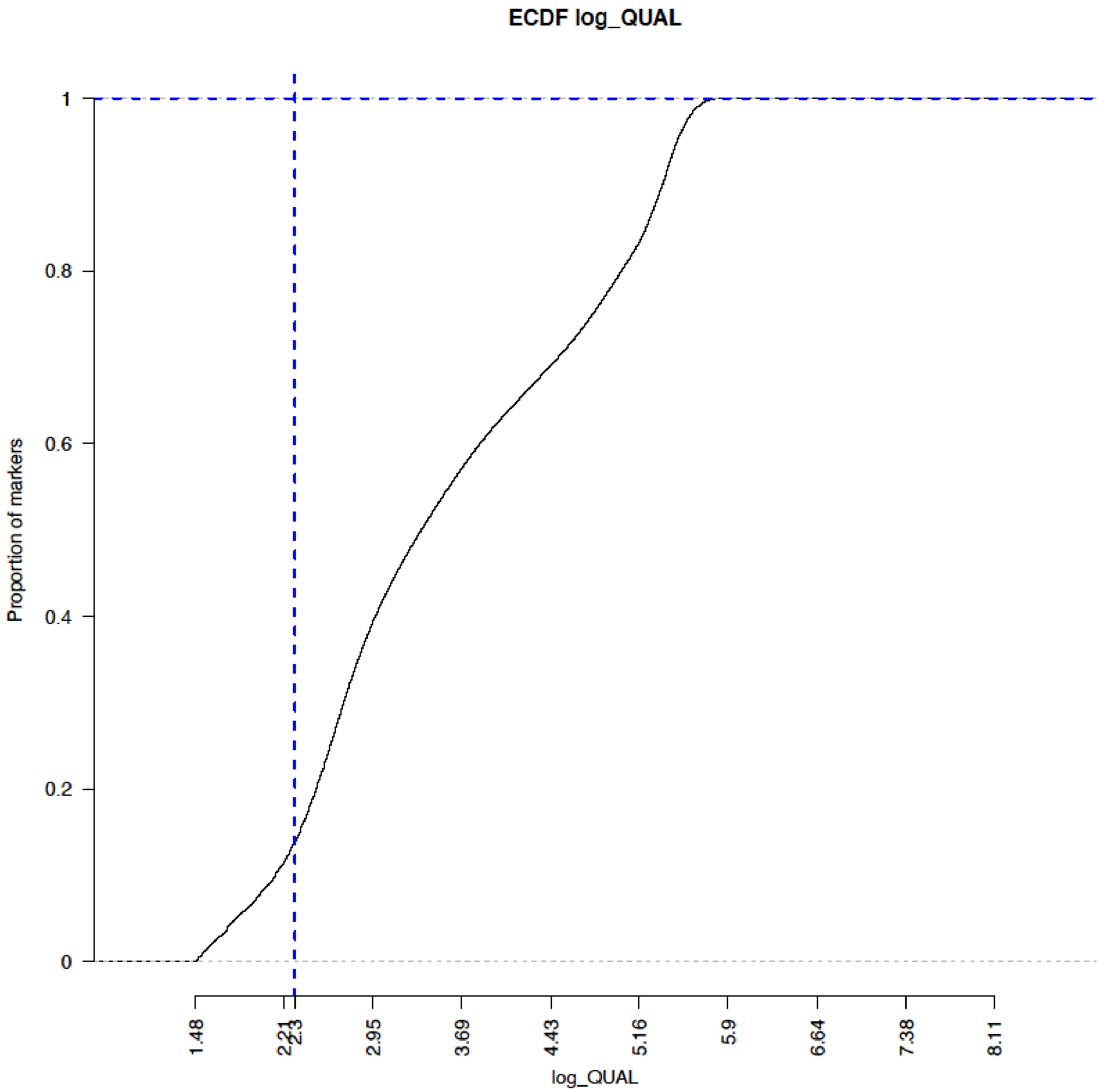


Empirical cumulative distribution function of QUAL values. The blue dotted line indicates the threshold used for filtering the vcf: QUAL > 200. X axis is on a log scale log(200)=2.3.

###### 4.4.1.5 Genotyping quality metrics: quality by depth (QD)

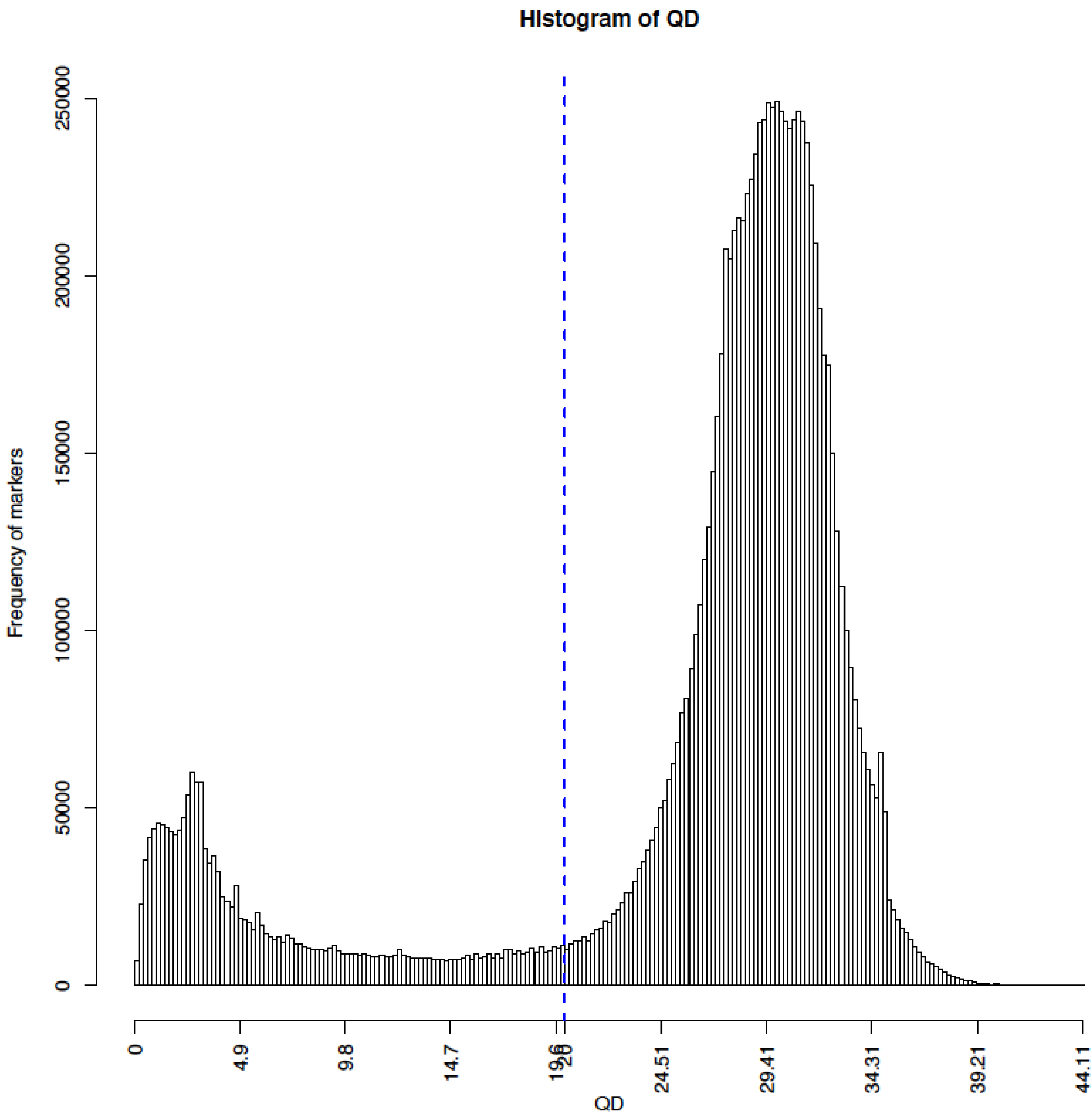


Counts of SNPs according to QUAL values. The blue dotted line indicates the threshold used for filtering the vcf: QD < 20.

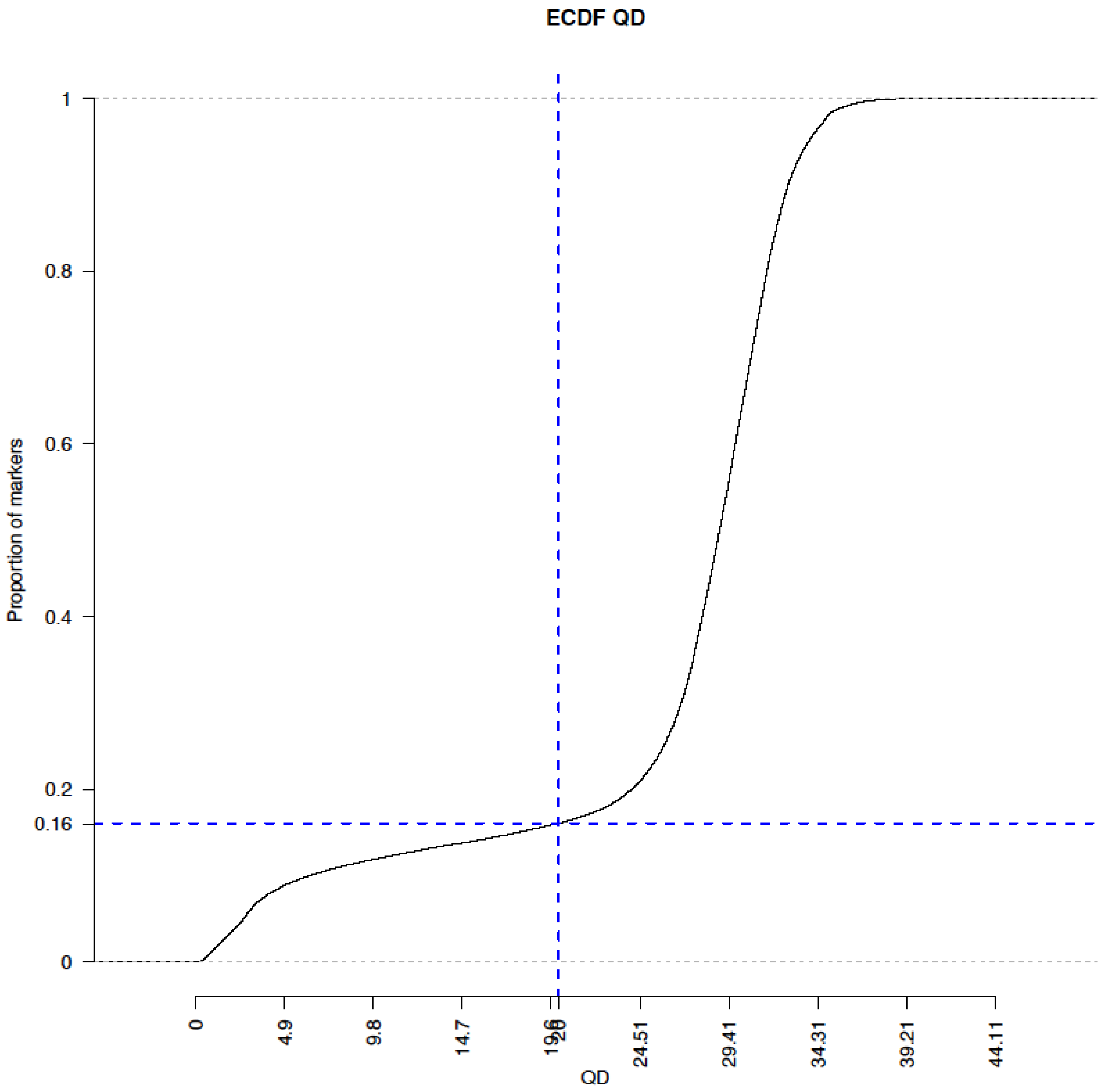


Empirical cumulative distribution function of QUAL. values. The blue dotted line indicates the threshold used for filtering the vcf: QD < 20.

###### 4.4.1.6. Individual genotyping metrics: heterozygote calls

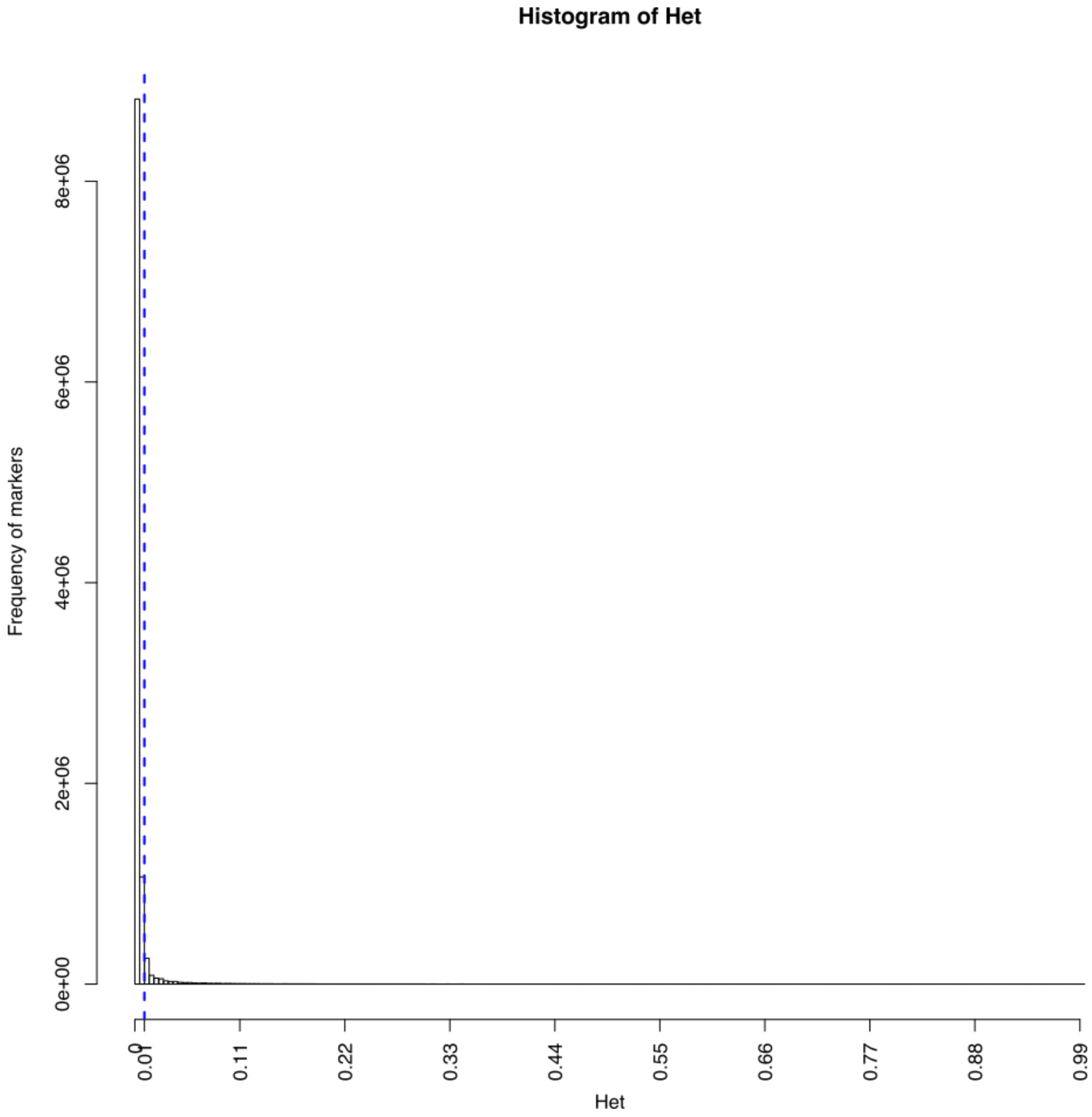


Counts of SNPs according to proportion of heterozygote genotypes. The blue dotted line indicates the threshold used for filtering the vcf: heterozygote genotypes for a SNP must be < 1%.

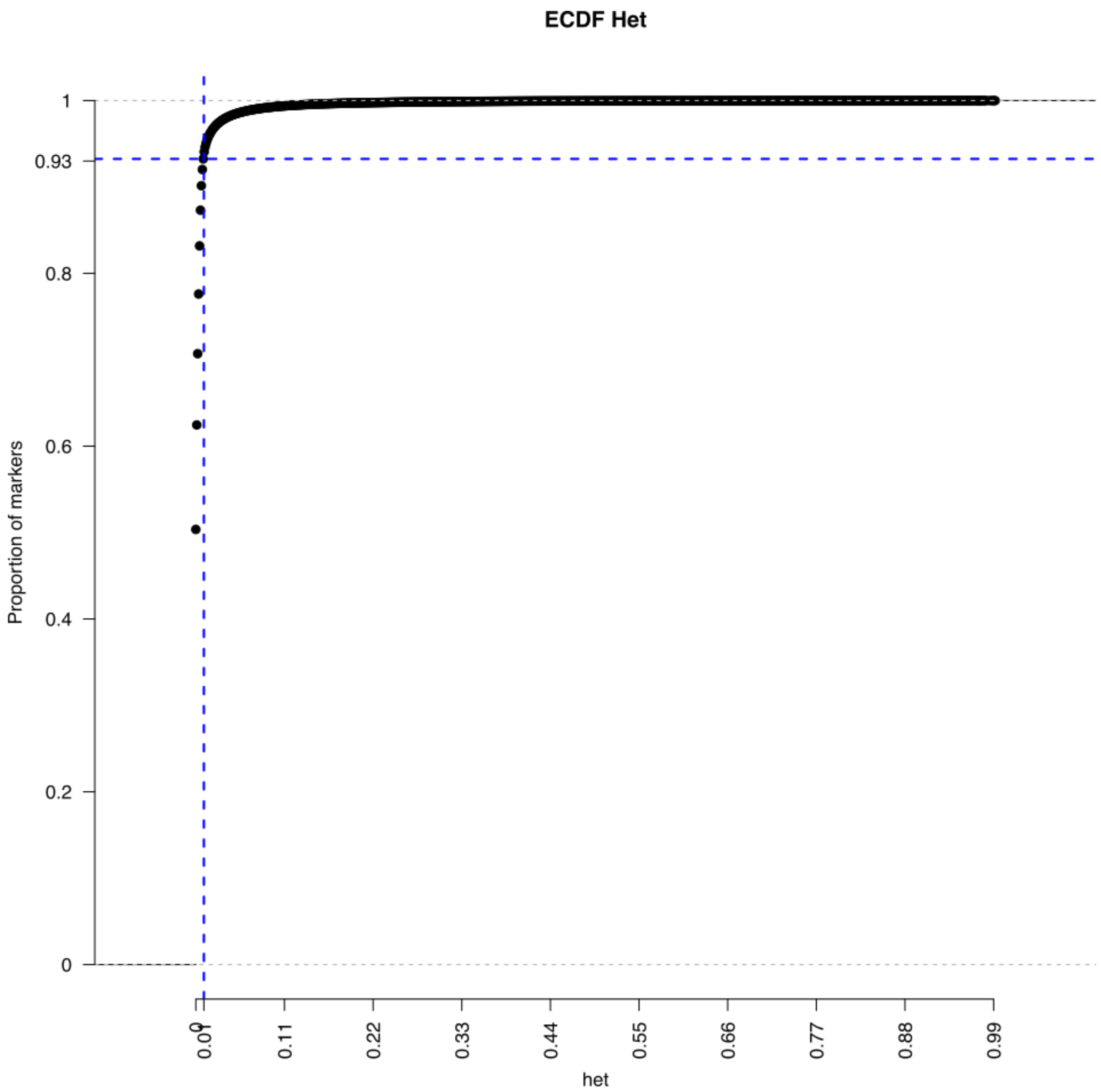


Empirical cumulative distribution function of heterozygote genotypes. The blue dotted line indicates the threshold used for filtering the vcf: heterozygote genotypes for a SNP must be < 1%.

###### 4.4.1.7. Individual genotyping metrics: missing genotypes

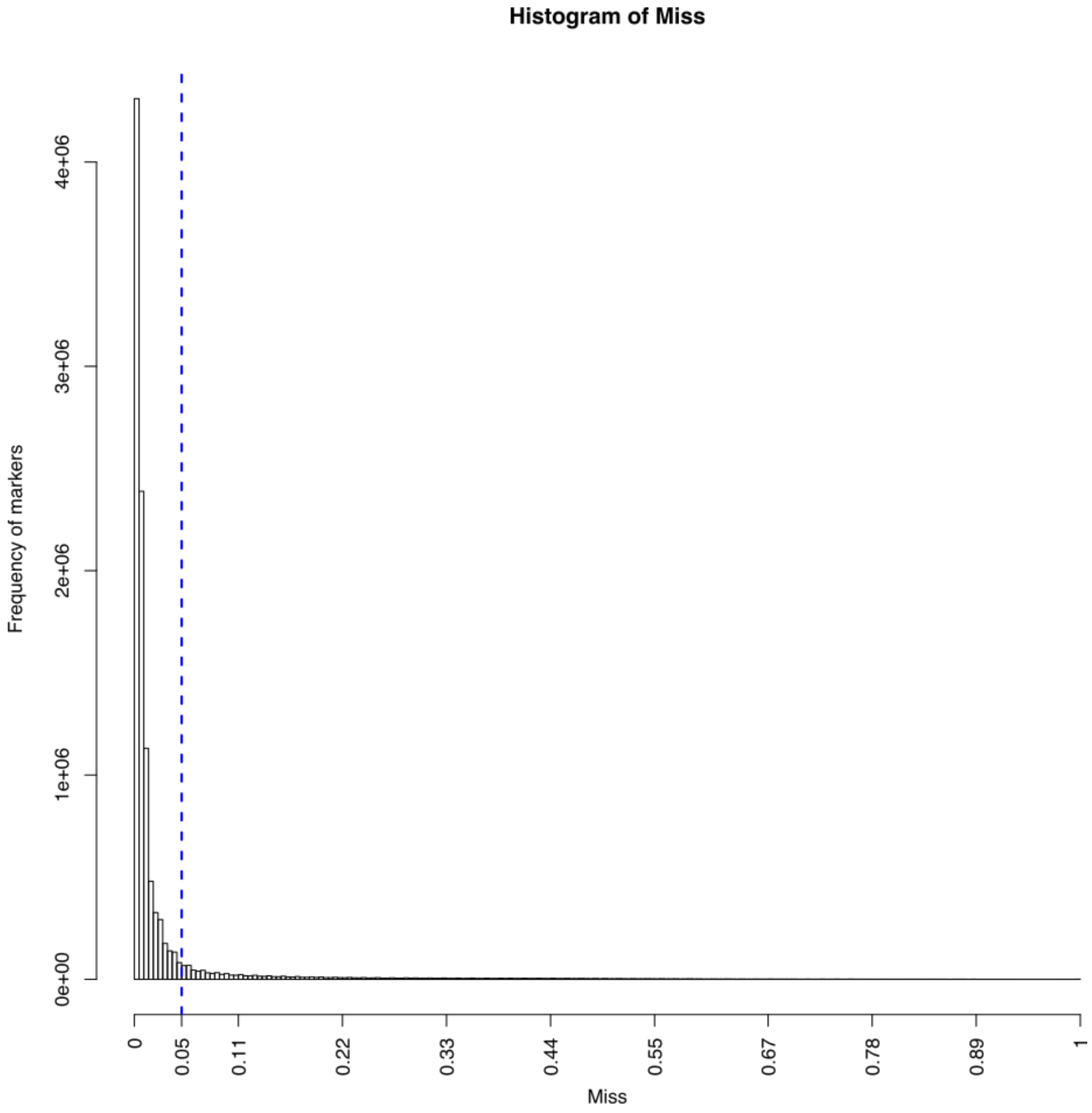


Counts of SNPs according to proportion of missing genotypes. The blue dotted line indicates the threshold used for filtering the vcf: missing genotypes for a SNP must be < 5%.

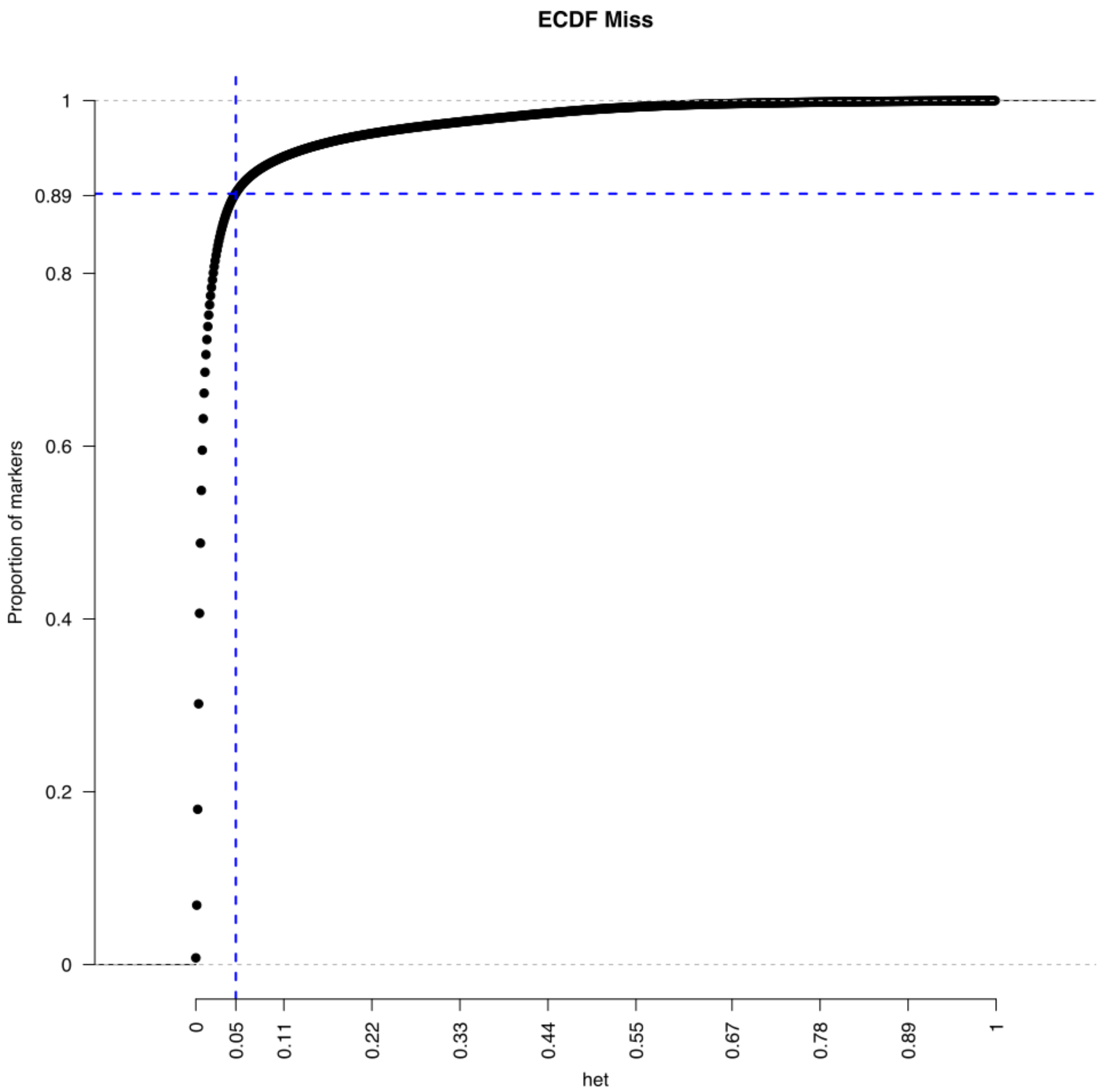


Empirical cumulative distribution function of missing genotypes. The blue dotted line indicates the threshold used for filtering the vcf: missing genotypes for a SNP must be < 5%.

###### 4.4.1.8. Individual genotyping metrics: genotype quality (QD)

QD is a quality measure of each individual genotype. Our filter removes all SNPs having more than 20 % of genotypes having QD values < 10. In the remaining SNPs, genotypes with QD values < 10 are retained.

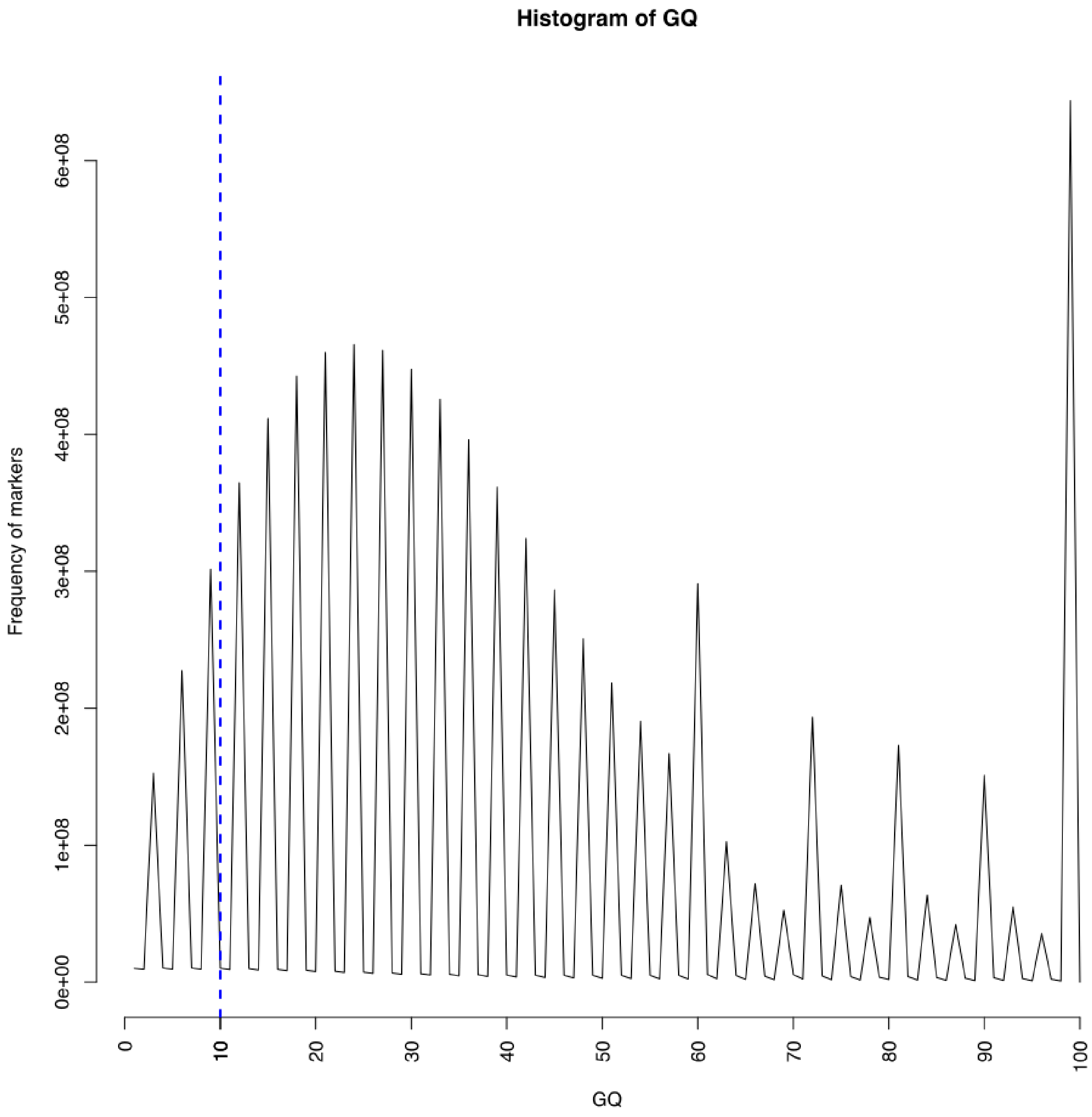


Counts of all individual GQ values for all SNPs. The blue dotted line indicates the threshold used: GQ < 10.

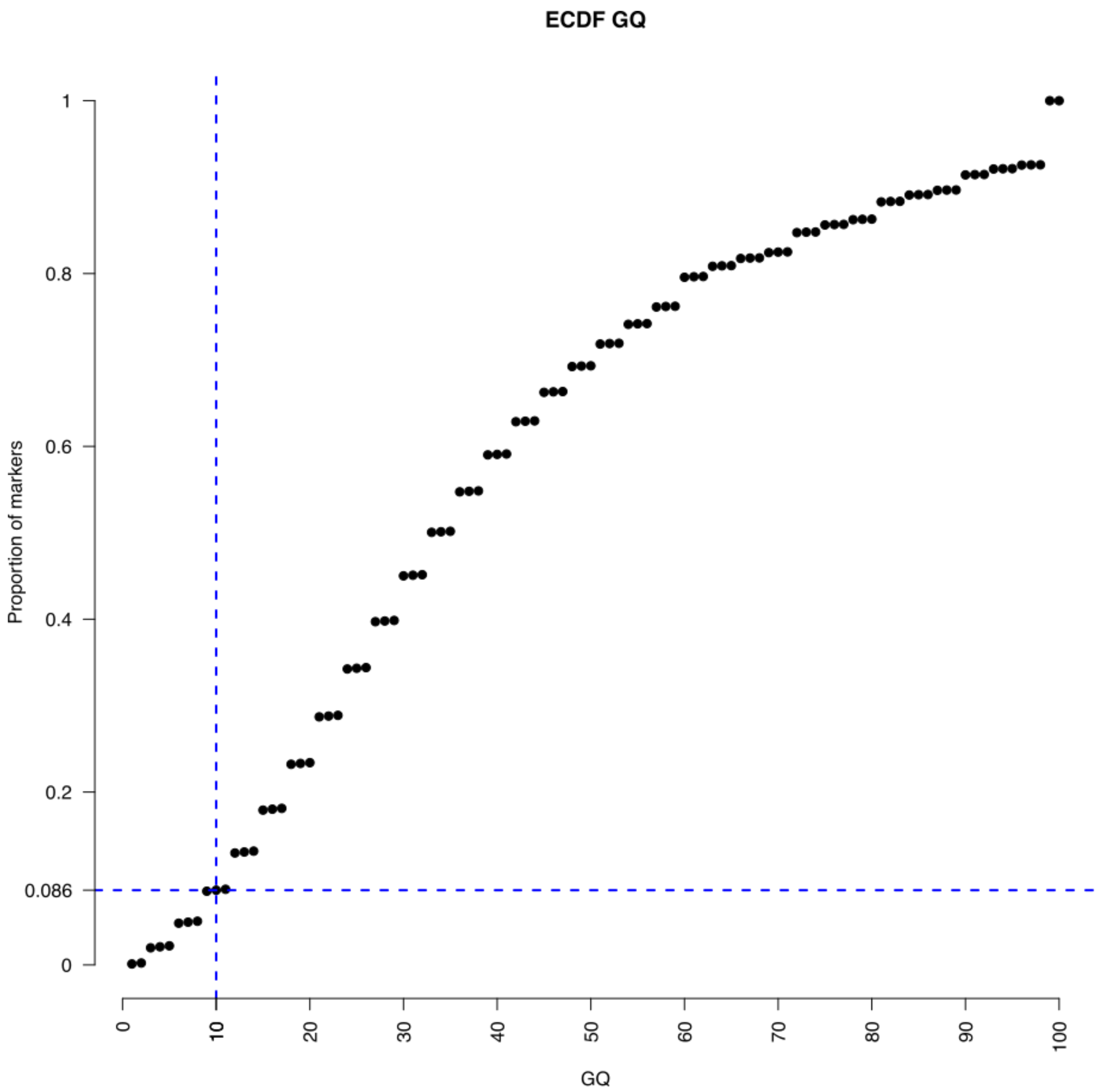

Empirical cumulative distribution function of GQ values for all SNPs. The blue dotted line indicates the threshold used: GQ < 10.

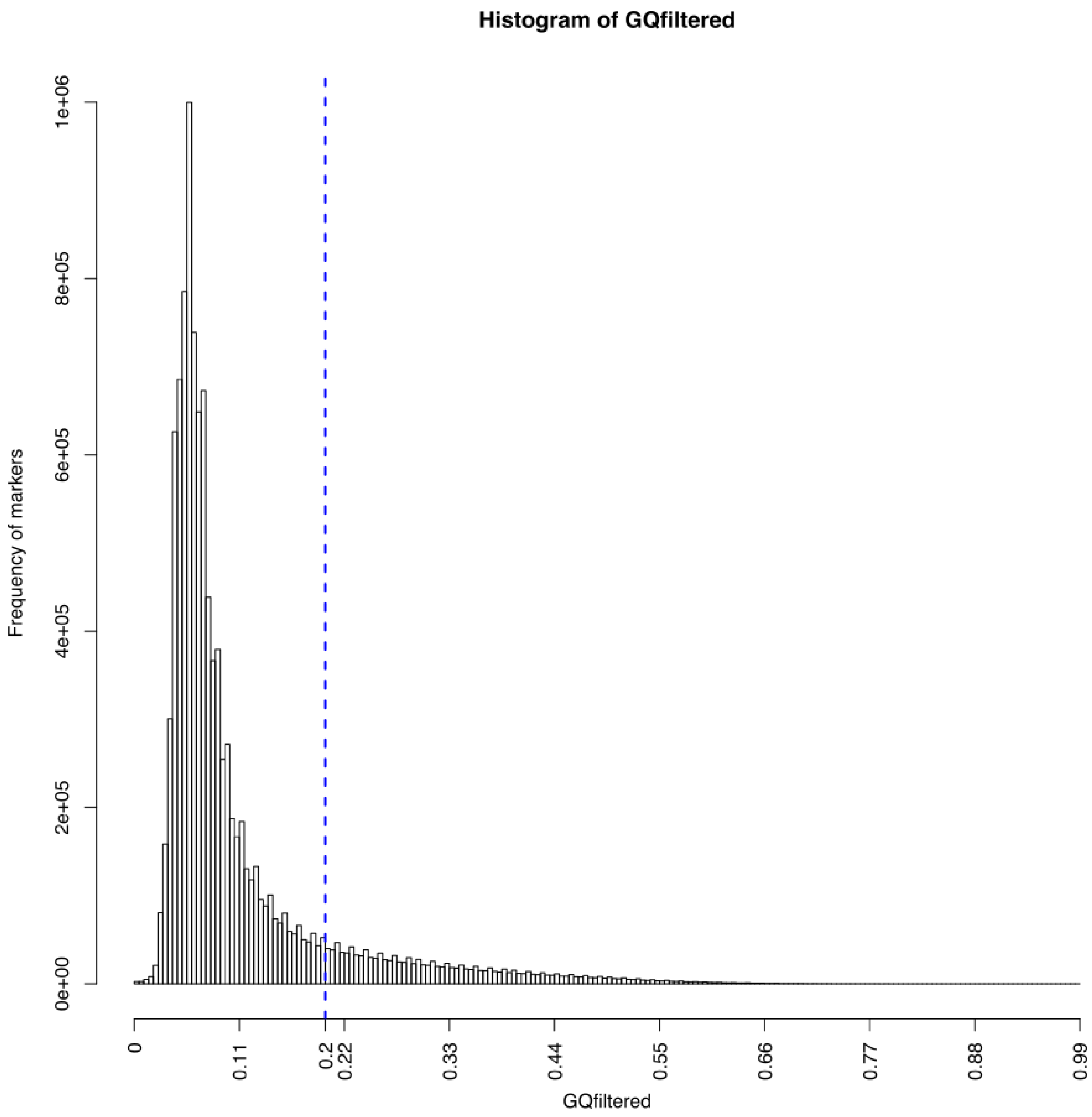


SNP counts according to proportion of genotypes with GQ < 10. The blue dotted line indicates the threshold used for filtering the vcf: proportion of genotypes for a SNP with GQ < 10%, must be < 20%.

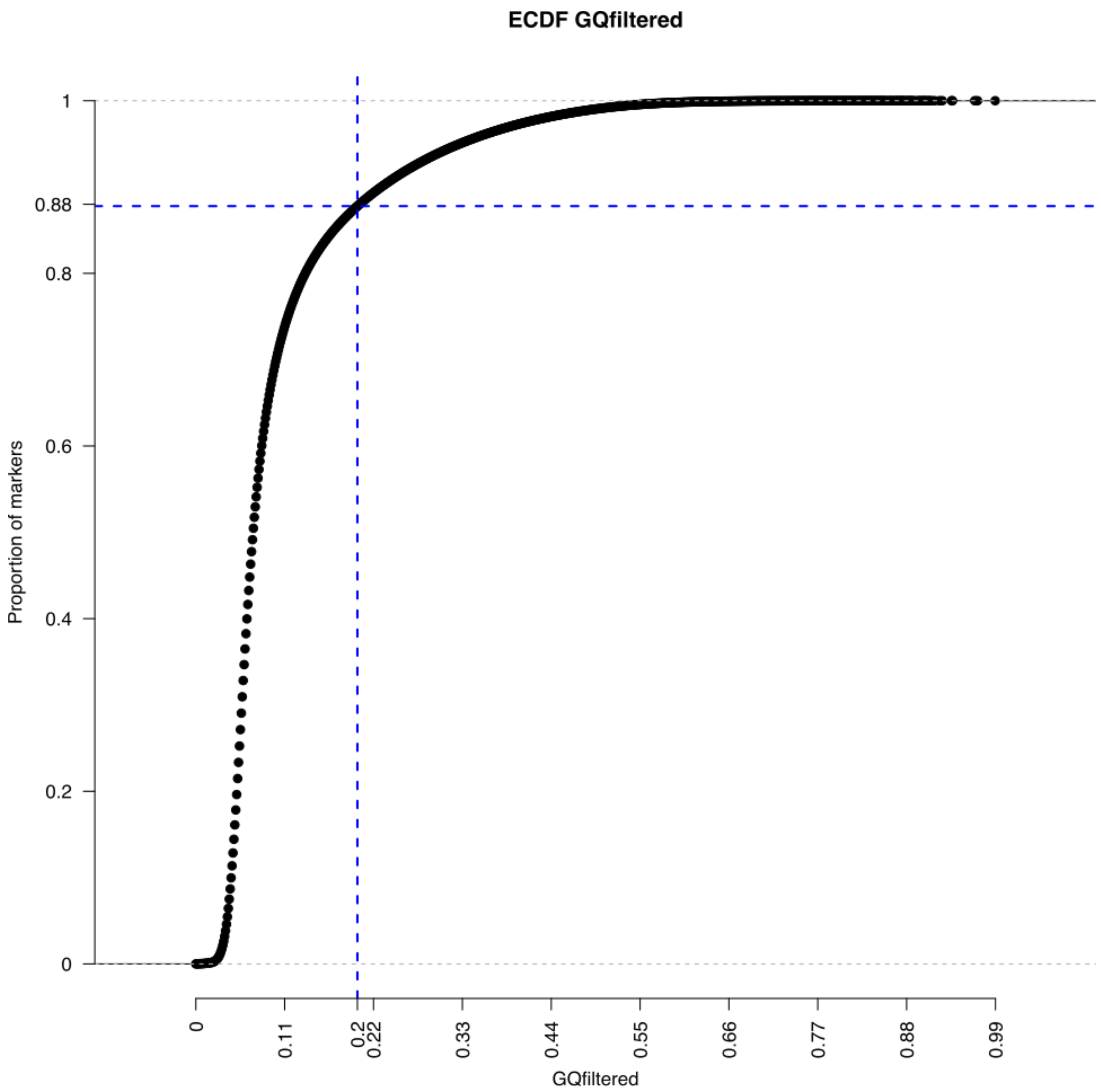

Empirical cumulative distribution function of SNP according to their proportion of genotypes with GQ < 10. The blue dotted line indicates the threshold used for filtering the vcf: proportion of genotypes for a SNP with GQ < 10%, must be < 20%.

---

###### 4.4.1.9 Notice on resulting missing data.

although SNPs with more than 5% of missing data were filtered out, some markers may have more than 5% missing data due to the heterozygote calls still remaining after the heterozygote filter, that were set to missing. These can be removed if needed during further analyses.

##### 4.4.2 Running the filters: generate Venn diagrams and filtered vcf

Running the following script with run='filter_all', will perform the filtering and produce the Venn diagrams and filtered vcf file.

```
#! /bin/bash

#run_vcf_cleanup.sh

#vcf filter (only done once to prepare list SNP positions)

# modules #####################################################
```

```
module load system/R-3.5.1
module load bioinfo/bcftools-1.6
module load bioinfo/tabix-0.2.5
module load bioinfo/vcftools-0.1.15
module load bioinfo/samtools-1.8
# end modules #################################################

# parameters ##################################################
username=avignal #Deprecated
SCRIPTS='~/seqapipopOnHAV3_1/vcf_cleanup_scripts' #path to scripts
DIRIN='~/combineGVCFs/The870vcf' #path to directory containing the input vcf
DIROUT='~/seqapipopOnHAV3_1/vcf_cleanup' #path to output directory
VCFIN='MetaGenotypesCalled870_raw_snps.vcf.gz' #inputvcf before filters
limit_FS=61
limit_SOR=4
limit_MQ=39
limit_MQRankSum=-12.5
limit_ReadPosRankSum=-8
limit_QUAL=200
limit_QD=20
limit_GQ=10
limit_miss=0.05
limit_het=0.01
limit_GQfiltered=0.2
quantile_prob_above_threshold=0.1
quantile_prob_below_threshold=0.9
kept_above_threshold="MQ_QUAL_QD~GQ~GQ"
kept_below_threshold="FS_SOR_allele~miss_het~GQfiltered"
run='filter_all'
# end parameters ##############################################
sbatch -W -J vcf_cleanup -o ${DIROUT}/log/vcf_cleanup.o -e ${DIROUT}/log/vcf_cleanup.e \
        --wrap="${SCRIPTS}/vcf_cleanup.sh ${username} ${SCRIPTS} ${DIRIN} ${DIROUT} ${VCFIN} \
        ${limit_allele} ${limit_FS} ${limit_SOR} ${limit_MQ} ${limit_MQRankSum} ${limit_ReadPosRankSum} \
        ${limit_QUAL} ${limit_QD} ${limit_GQ} ${limit_miss} ${limit_het} ${limit_GQfiltered} \
        ${quantile_prob_above_threshold} ${quantile_prob_below_threshold} \
        ${kept_above_threshold} ${kept_below_threshold} ${run}"

# end of file
```

#### 5. results:

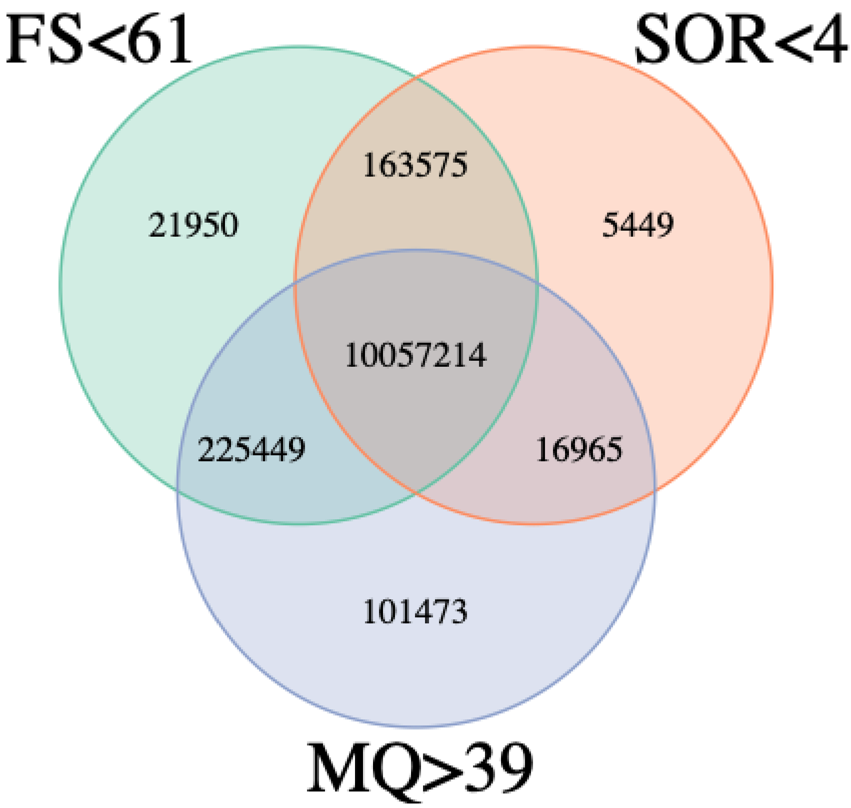


**Filters on mapping quality (MQ) and strand bias (FS and SOR) metrics.**

FS (FisherStrand): phred-scaled probability that there is strand mapping bias at the site; SOR (StrandOddsRatio): strand bias mapping estimate; MQ (RMSMappingQuality): root mean square mapping quality over all the reads at the site. The intercept of the 3 filters gives 10,057,214 SNPs, used for further filtering

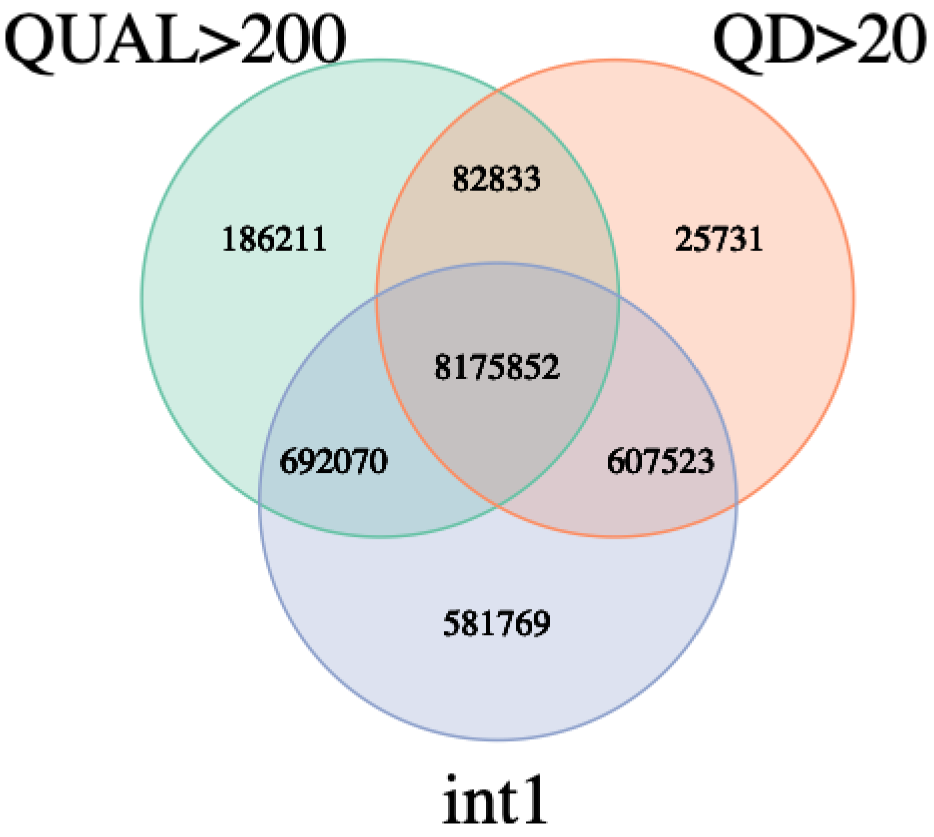


**Filters on genotyping quality.**

Int1 is the intersect of the mapping quality filters. QUAL: Phred-scaled quality score for the assertion made in ALT: the more samples have the ATL allele, the higher the QUAL score. QD: quality score normalized by allele depth in which only informative reads are counted. The intercept of the 2 filters with the previous mapping quality filters gives 8,175,852 SNPs, used for further filtering (see figure 3).

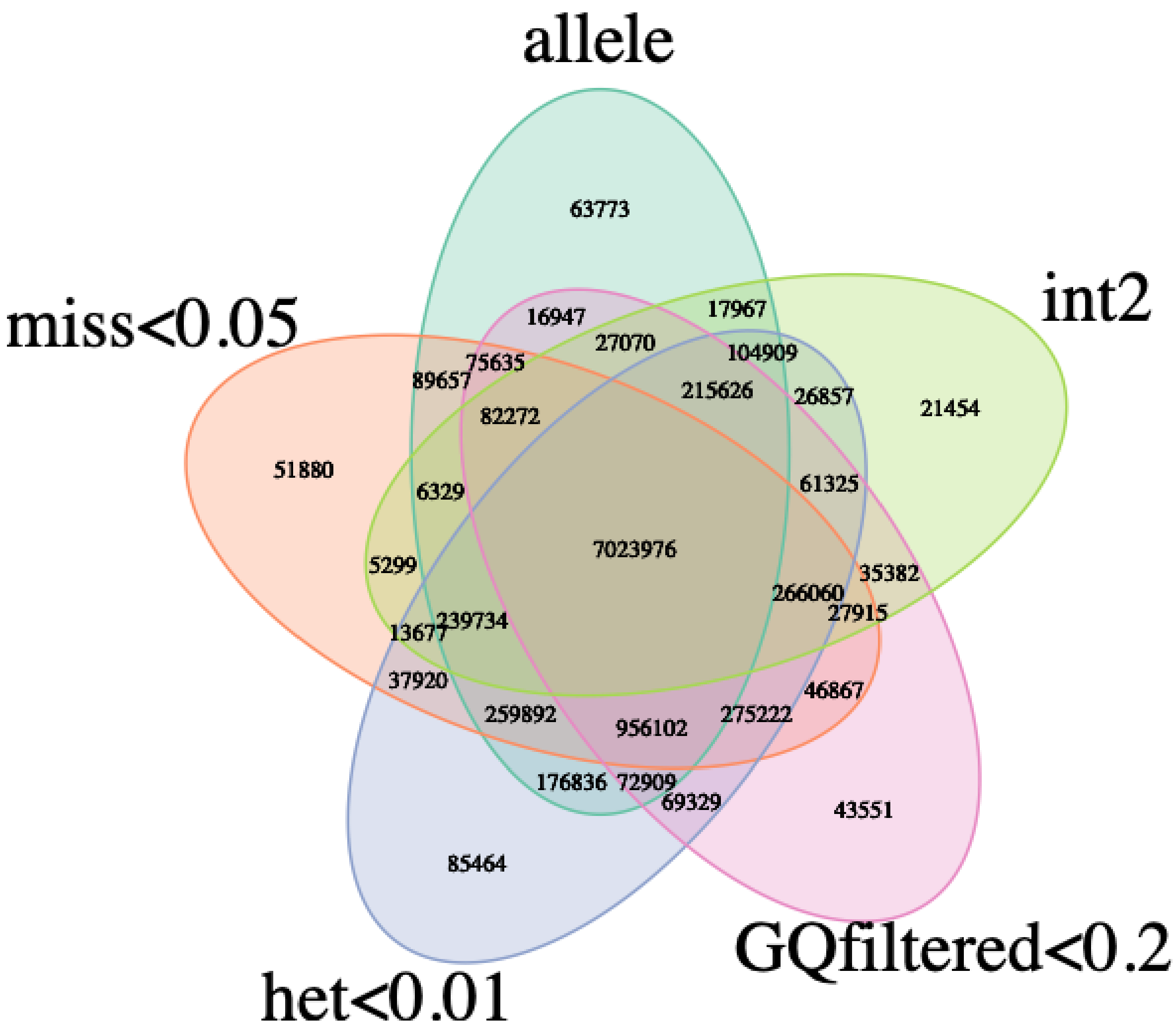


**Filters individual genotyping quality.**

Int2 is the intersect of the previous filters. Filters are (i) het: proportion of heterozygote calls less than 1% for a SNP, as haploid drones were sequenced, the remaining heterozygote calls were set to missing; (ii) allele: less than 4 alleles for a SNP; (iii) miss: less than 5% missing data; (iv) GCfiltered: SNPs are removed if more than 20% samples have a genotyping quality (GQ) under 10. Note: although SNPs with more than 5% of missing data were filtered out, some markers may have more than 5% missing data due to the heterozygote calls that were set to missing.

---

- **Final vcf file:**
    - **MetaGenotypesCalled870_raw_snps_allfilter.vcf**
- **The final vcf file has just over 7 million SNPs : 7,023,976 in total.**
- **However, although SNPs with more than 5% of missing data were filtered out, some markers may have more than 5% missing data due to the hetozygote calls that were set to missing.**
- **Markers can have up to 3 alleles**

**Number of SNPs per chromosome:**

| Accession | Chr | Nb. SNP |
|---|---|---|
| NC_001566.1 | MT | 287 |

| Accession | Chr | Nb. SNP |
|---|---|---|
| NC_037638.1 | 1 | 913023 |
| NC_037639.1 | 2 | 534733 |
| NC_037640.1 | 3 | 442882 |
| NC_037641.1 | 4 | 440141 |
| NC_037642.1 | 5 | 462122 |
| NC_037643.1 | 6 | 577596 |
| NC_037644.1 | 7 | 463575 |
| NC_037645.1 | 8 | 397891 |
| NC_037646.1 | 9 | 378566 |
| NC_037647.1 | 10 | 355296 |
| NC_037648.1 | 11 | 441395 |
| NC_037649.1 | 12 | 388488 |
| NC_037650.1 | 13 | 378466 |
| NC_037651.1 | 14 | 330298 |
| NC_037652.1 | 15 | 285750 |
| NC_037653.1 | 16 | 233467 |
| Sum | Sum | 7023976 |

### 6. Prepare bed, bim, fam files for plink, admixture, etc.

#### 6.1. change chromosome names to numbers

- The chromosome names have to be numbers
- Script to generate sed commands for all chromosomes: substForPlinkWrite.bash
- vcf file with chromosomes as numbers:
  - MetaGenotypesCalled870_raw_snps_allfilter_plink.vcf

```
#!/bin/bash

#substForPlinkWrite.bash

printf "#!/bin/bash\n\n"

printf "cp ~/seqapipopOnHAV3_1/seqApiPopVcfFilteredSonia/vcf_cleanup/
MetaGenotypesCalled870_raw_snps_allfilter.vcf ~/seqapipopOnHAV3_1/seqApiPopVcfFilteredSonia/plinkAnalyses
/MetaGenotypesCalled870_raw_snps_allfilter_plink.vcf\n\n"

j=0
for i in `cut -f1 /home/gencel/vignal/save/Genomes/Abeille/HAv3_1_indexes/HAv3_1_Chromosomes.list`
do
j=$((j+1))
printf "sed -i \'s/${i}/${j}/g\' ~/seqapipopOnHAV3_1/seqApiPopVcfFilteredSonia/
plinkAnalyses/MetaGenotypesCalled870_raw_snps_allfilter_plink.vcf\n"
done
```

```
substForPlinkWrite.bash > substForPlink.bash
```

```
sbatch substForPlink.bash
```

#### 6.2 Prepare bed, bim, fam files and filter on missing data for SNPs and samples

- **Final bed, bim, bam files:**
  - **MetaGenotypesCalled870_raw_snps_allfilter_plink_missIndGeno.bim**
  - **MetaGenotypesCalled870_raw_snps_allfilter_plink_missIndGeno.bam**
  - **MetaGenotypesCalled870_raw_snps_allfilter_plink_missIndGeno.fam**
- --maf filters out all variants with minor allele frequency below the provided threshold (default 0.01)
- --geno filters out all variants with missing call rates exceeding the provided value (default 0.1) to be removed
- --mind does the same for samples.
- --indep-pairwise 500000 50000 0.9 : LD filters out variants with LD > 0.9, on a window of 500000 SNPs, a sliding window of 50000.

File with the 7023689 variants: /work/project/cytogen/Alain/seqapipopOnHAV3_1/seqApiPopVcfFilteredSonia/plinkAnalyses/MetaGenotypesCalled870_raw_snps_allfilter_plink

```
#! /bin/bash

#convertToBed.bash

module load -f /work/project/cytogen/Alain/seqapipopOnHAV3_AV/program_module

VCFin=/work/project/cytogen/Alain/seqapipopOnHAV3_1/seqApiPopVcfFilteredSonia/plinkAnalyses/MetaGenotypesCall
VCFout=/work/project/cytogen/Alain/seqapipopOnHAV3_1/seqApiPopVcfFilteredSonia/plinkAnalyses/MetaGenotypesCal
plink --vcf ${VCFin} \
  --keep-allele-order \
  --a2-allele ${VCFin} 4 3 '#' \
  --allow-no-sex \
  --allow-extra-chr \
  --chr-set 16 \
  --set-missing-var-ids @:#[HAV3.1]\$1\$2 \
  --chr 1-16 \
  --mind 0.1 \
  --geno 0.1 \
  --out ${VCFout} \
  --make-bed \
  --missing
```

- 11075 variants removed due to missing genotype data (--geno).
- 15 samples removed due to missing genotype data (--mind).
- 7012614 variants and 855 samples pass filters and QC.

**Samples removed: frequency of missing genotypes from the MetaGenotypesCalled870_raw_snps_allfilter_plink_missIndGeno.imiss plink file:**

| ID | N_MISS | N_GENO | F_MISS |
|---|---|---|---|
| AOC4 | 1270404 | 7023689 | 0.1809 |
| BR12 | 1269182 | 7023689 | 0.1807 |
| BR1A | 1253619 | 7023689 | 0.1785 |
| ESP9 | 6208279 | 7023689 | 0.8839 |
| JFM21 | 725846 | 7023689 | 0.1033 |
| JFM24 | 817509 | 7023689 | 0.1164 |
| JFM3 | 875208 | 7023689 | 0.1246 |
| JFM5 | 830181 | 7023689 | 0.1182 |
| KF21 | 722607 | 7023689 | 0.1029 |
| OUE8 | 831427 | 7023689 | 0.1184 |
| PM1 | 969888 | 7023689 | 0.1381 |
| SavB1 | 823422 | 7023689 | 0.1172 |
| SavB3 | 706024 | 7023689 | 0.1005 |
| XC3 | 821334 | 7023689 | 0.1169 |
| XC4 | 747325 | 7023689 | 0.1064 |

#### 7. Plots on sequencing depth and missing genotypes

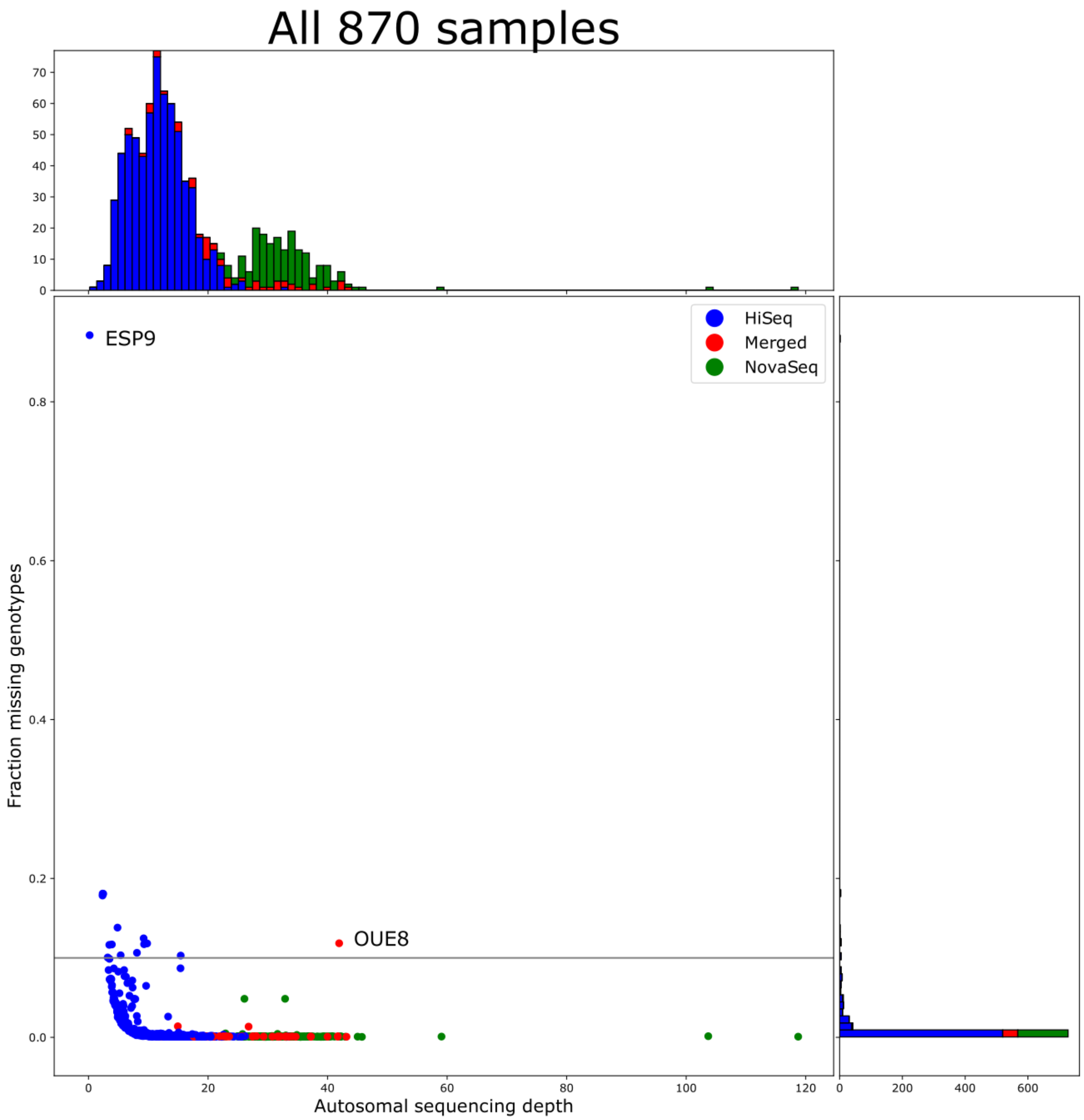


**Sequencing depth and fraction of missing genotypes for all the 870 sequenced samples.**

Blue: samples sequenced with the Illumina™ HiSeq instrument; green: samples sequenced with the Illumina™ NovaSeq instrument and red: samples sequenced with the Illumina™ HiSeq instrument in two or more runs. The horizontal line is the genotyping rate threshold applied to final dataset. Sequencing data could not be obtained from sample ESP9 and data for sample OUE8 was obtained from 3 sequencing runs.

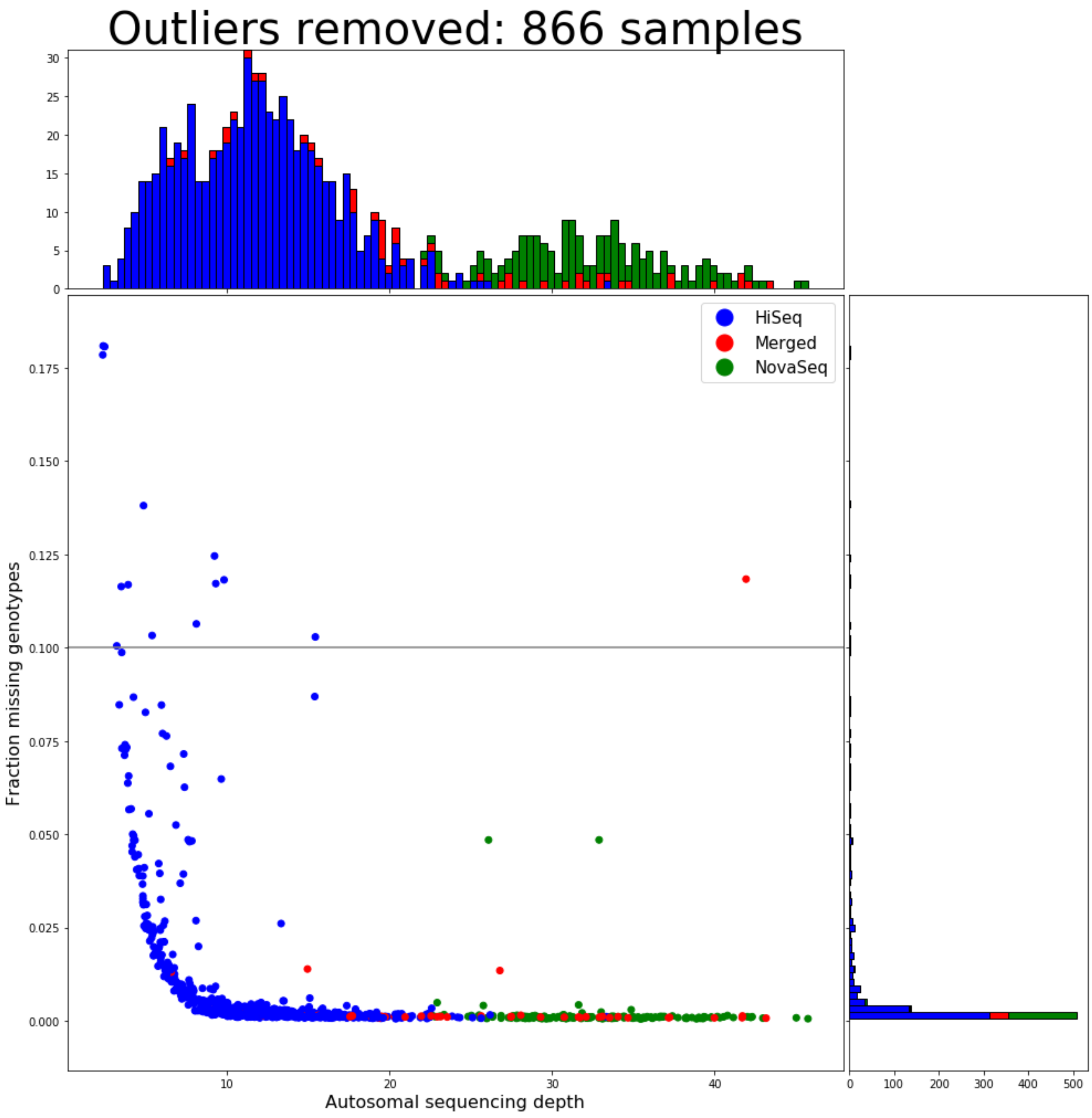


**Sequencing depth and fraction of missing genotypes with outliers removed.**

Blue: samples sequenced with the IlluminaTM HiSeq instrument; green: samples sequenced with the IlluminaTM NovaSeq instrument and red: samples sequenced with the IlluminaTM HiSeq instrument in two or more runs. The horizontal line is the genotyping rate threshold applied to final dataset.
