## Supplementary methods 3 for "Complex population structure and haplotype patterns in Western Europe honey bee from sequencing a large panel of haploid drones"

### SeqApiPop analyses: filtering on LD and PCA


- 1. Defining haplotype blocks
  - 1.1 Obtaining 5 first columns for further Analyses
- 2. Select the 629 samples for the plinkAnalyses
- 3. PCA with smartpca
- 3.1. plot of PC1 and PC2 using all markers
- 3.2. Influence of MAF filtering thresholds on dataset's global SNP contributions
- 3.3. SNP contributions along the genome
  - 3.3.1. Genome-wide plots
  - 3.3.2. Plots in haplotype haploBlocks
- 4. LD pruning
  - 4.1. SNP density plots and window size selection
  - 4.2. LD pruning and effects on PCA
    - 4.2.1. Generating datasets (plink)
    - 4.2.2. Effect of LD pruning on SNP contributions to the PCs
    - 4.2.3. Effect of LD pruning on PCA


#### 1. Defining haplotype blocks

The plink blocks command was used to detect haplotype blocks in the genome, based on the complete dataset of 870 samples.

```
#! /bin/bash

#haplotypeBlocks5000.sh

module load -f /work/project/cytogen/Alain/seqapipopOnHAV3_AV/program_module

NAME=haplotypeBlocks5000

plink --bfile ../MetaGenotypesCalled870_raw_snps_allfilter_plink_missIndGeno \
  --out ${NAME} \
  --blocks no-pheno-req no-small-max-span \
  --blocks-max-kb 5000
```

##### 1.1 Obtaining 5 first columns for further Analyses

```
awk '{print $1,$2,$3,$4,$5}' haplotypeBlocks5000.blocks.det > haplotypeBlocks5000.blocks.cols
```

#### 2. Select the 629 samples for the plinkAnalyses

- The vcf file contains a total of 870 samples
  - All 870 samples are of the genetic types found in western Europe and therefore were included for SNP detection based on technical metrics.
  - A list of 629 samples will be selected for the genetic analyses, by removing:
  - Duplicate samples from a same hive
  - Samples from experimental subpopulations
  - Samples from another study
  - The 15 samples with > 0.1 missing data
- 3 set of files formatted for plink will be produced, with prefixes:

- SeqApiPop_629
- All SNPs for the 629 samples
- SeqApiPop_629_maf001
- Only marlers with MAF > 0.01
- SeqApiPop_629_maf005
- Only marlers with MAF > 0.05
- SeqApiPop_629_maf01
- Only marlers with MAF > 0.1
- SeqApiPop_629_maf02
- Only marlers with MAF > 0.2

```
#! /bin/bash

#select629.bash

module load -f /work/project/cytogen/Alain/seqapipopOnHAV3_AV/program_module

NAME=SeqApiPop_629

VCFin=~/plinkAnalyses/MetaGenotypesCalled870_raw_snps_allfilter_plink.vcf
VCFout=~/plinkAnalyses/WindowSNPs/${NAME}
plink --vcf ${VCFin} \
  --keep-allele-order \
  --keep ~/seqapipopOnHAV3_1/seqApiPopVcfFilteredSonia/plinkAnalyses/WindowSNPs/Unique629.list \
  --a2-allele ${VCFin} 4 3 '#' \
  --allow-no-sex \
  --allow-extra-chr \
  --chr-set 16 \
  --set-missing-var-ids @:#[HAV3.1]\$1\$2 \
  --chr 1-16 \
  --mind 0.1 \
  --geno 0.1 \
  --out ${VCFout} \
  --make-bed \
  --missing

plink --bfile ${VCFout} \
  --maf 0.01 \
  --out ${VCFout}_maf001 \
  --make-bed

plink --bfile ${VCFout} \
  --out ${VCFout}_maf005 \
  --maf 0.05 \
  --make-bed
```

- 7012891 variants and 629 samples pass filters and QC
- This number is slightly different than when using all 870 samples, due to variations in the number of variants removed due to missing genotypind data.

### 3. PCA with smartpca

- script paramFile.bash to generate smartpca parameter files:
- PREFIX = prefix of the plink bed file to read

```
#!/bin/bash
PREFIX=SeqApiPop_629
#PREFIX=SeqApiPop_629_maf005
#PREFIX=SeqApiPop_629_maf001
#PREFIX=SeqApiPop_629_maf001_LD03_pruned
echo genotypename: ../${PREFIX}.bed > ${PREFIX}.par
echo snpname: ../${PREFIX}.bim >> ${PREFIX}.par
echo indivname: ${PREFIX}.PCA.fam >> ${PREFIX}.par
```

```
echo snpweightoutname: ${PREFIX}.snpeigs >> ${PREFIX}.par
echo evecoutname: ${PREFIX}.eigs >> ${PREFIX}.par
echo evaloutname: ${PREFIX}.eval >> ${PREFIX}.par
echo phylipoutname: ${PREFIX}.fst >> ${PREFIX}.par
echo numoutevec: 20 >> ${PREFIX}.par
echo numoutlieriter: 0 >> ${PREFIX}.par
echo altnormstyle: NO >> ${PREFIX}.par
echo missingmode: NO >> ${PREFIX}.par
echo nsnpldregress: 0 >> ${PREFIX}.par
echo noxdata: YES >> ${PREFIX}.par
echo nomalexhet: YES >> ${PREFIX}.par
```

- The fam file needs a specific format with an additional status column:

```
awk '{print $1,$2,$3,$4,$5,1}' ../SeqApiPop_629.fam > SeqApiPop_629.PCA.fam
```

- Then run smartpca:

```
sbatch --wrap="smartpca -p SeqApiPop_629.par"
```

#### 3.1. plot of PC1 and PC2 using all markers

- From the log of smartpca, with SeqApiPop_629.* only:
  - number of samples used: 629 number of snps used: 7012891
  - total number of snps killed in pass: 444784 used: 6568107
- As soon as there is the slightest amount of selection on MAF, all remaining SNPs are used.
  - This could be an effect of SNPs only present in the individuals that were removed from the original 870 sample dataset.
- Plot of PC1 and PC2 with all markers and samples from reference populations (file SeqApiPop_629.eigs):

**Reference populations:**

| Population | Reference for |
|---|---|
| Mellifera Ouessant | *Apis mellifera mellifera* |
| Mellifera Ouessant | *Apis mellifera mellifera* |
| Mellifera Ouessant | *Apis mellifera mellifera* |
| Mellifera Ouessant | *Apis mellifera mellifera* |
| Mellifera Ouessant | *Apis mellifera mellifera* |
| Iberica Spain | *Apis mellifera iberica* |
| Ligustica Italy | *Apis mellifera ligustica* |
| Carnica Slovenia | *Apis mellifera carnica* |
| Carnica Germany | *Apis mellifera carnica* |
| Carnica Switzerland | *Apis mellifera carnica* |
| Carnica France | *Apis mellifera carnica* |
| Carnica Poland | *Apis mellifera carnica* |
| Caucasica | *Apis mellifera caucasica* |

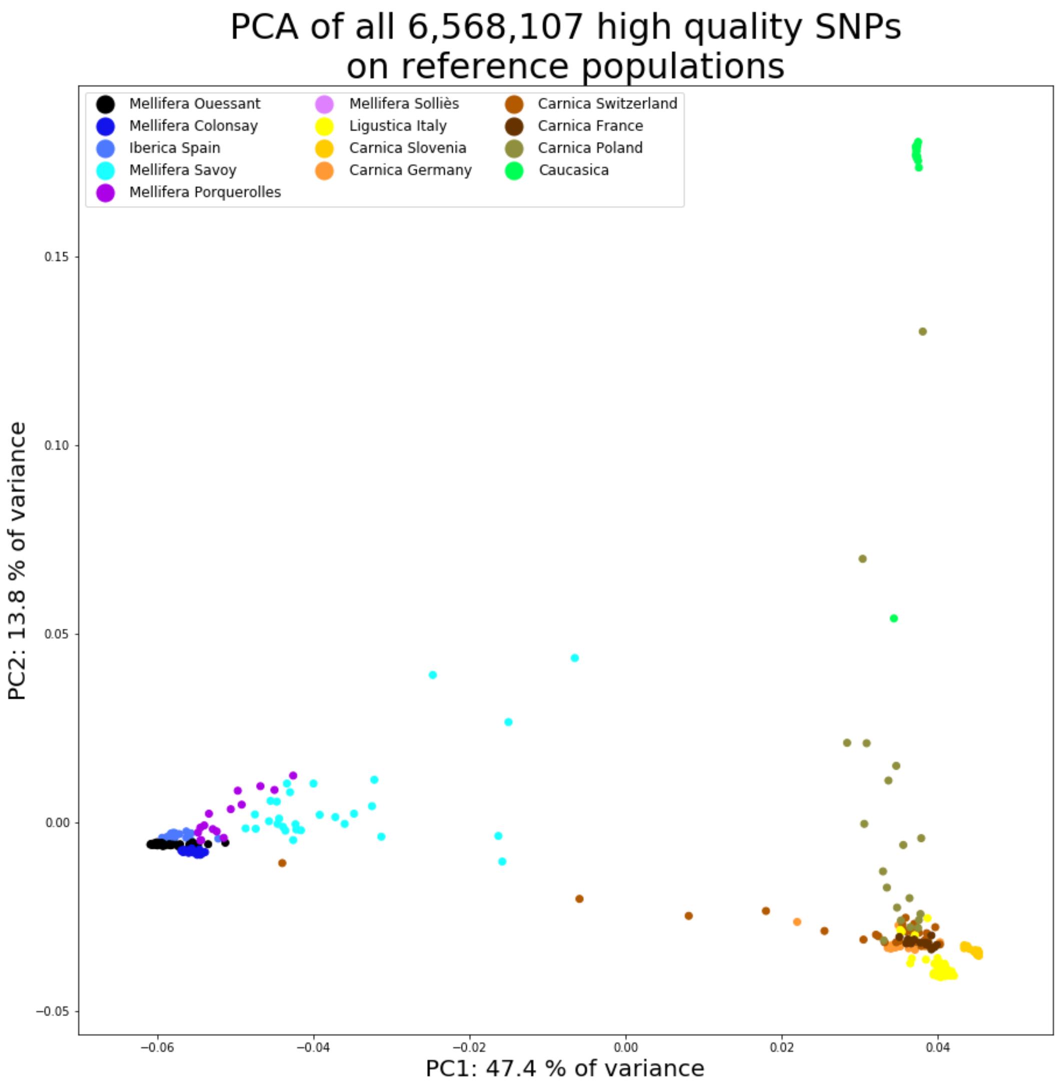


**Principal component analysis for all SNPs: reference populations.**

The first component representing 47.4 % of the variance separates clearly the A. m. mellifera and A. m. iberica on one side and the A. m. ligustica, A. m. carnica and A. m. caucasica on the other. The second component representing 13.8 % of the variance distinguishes the A. m. caucasica from the rest. The other dimensions represent each 5.2 % or less of the variance (see table below) and the separation of other populations or groups of populations on these subsequent principal component dimensions is not clear (data not shown)

**Contribution to the variance of PC1 to PC20**

| PC | percentVar |
|---|---|
| PC1 | 47.417834 |
| PC2 | 13.826587 |
| PC3 | 5.212604 |
| PC4 | 3.096361 |
| PC5 | 3.075373 |
| PC6 | 2.996320 |
| PC7 | 2.943851 |
| PC8 | 2.505911 |
| PC9 | 2.098054 |
| PC10 | 1.807726 |
| PC11 | 1.719579 |
| PC12 | 1.613242 |
| PC13 | 1.512502 |
| PC14 | 1.488016 |
| PC15 | 1.483119 |
| PC16 | 1.481020 |
| PC17 | 1.454436 |
| PC18 | 1.430650 |
| PC19 | 1.422255 |
| PC20 | 1.414560 |

#### 3.2. Influence of MAF filtering thresholds on dataset's global SNP contributions

- The output files *.SeqApiPop_629.snpeigs from the smartpca analysis are used to plot the SNP contributions to the principal components.
    - SeqApiPop_629.snpeigs
    - SeqApiPop_629_maf001.snpeigs
    - SeqApiPop_629_maf005.snpeigs
    - SeqApiPop_629_maf01.snpeigs
    - SeqApiPop_629_maf02.snpeigs

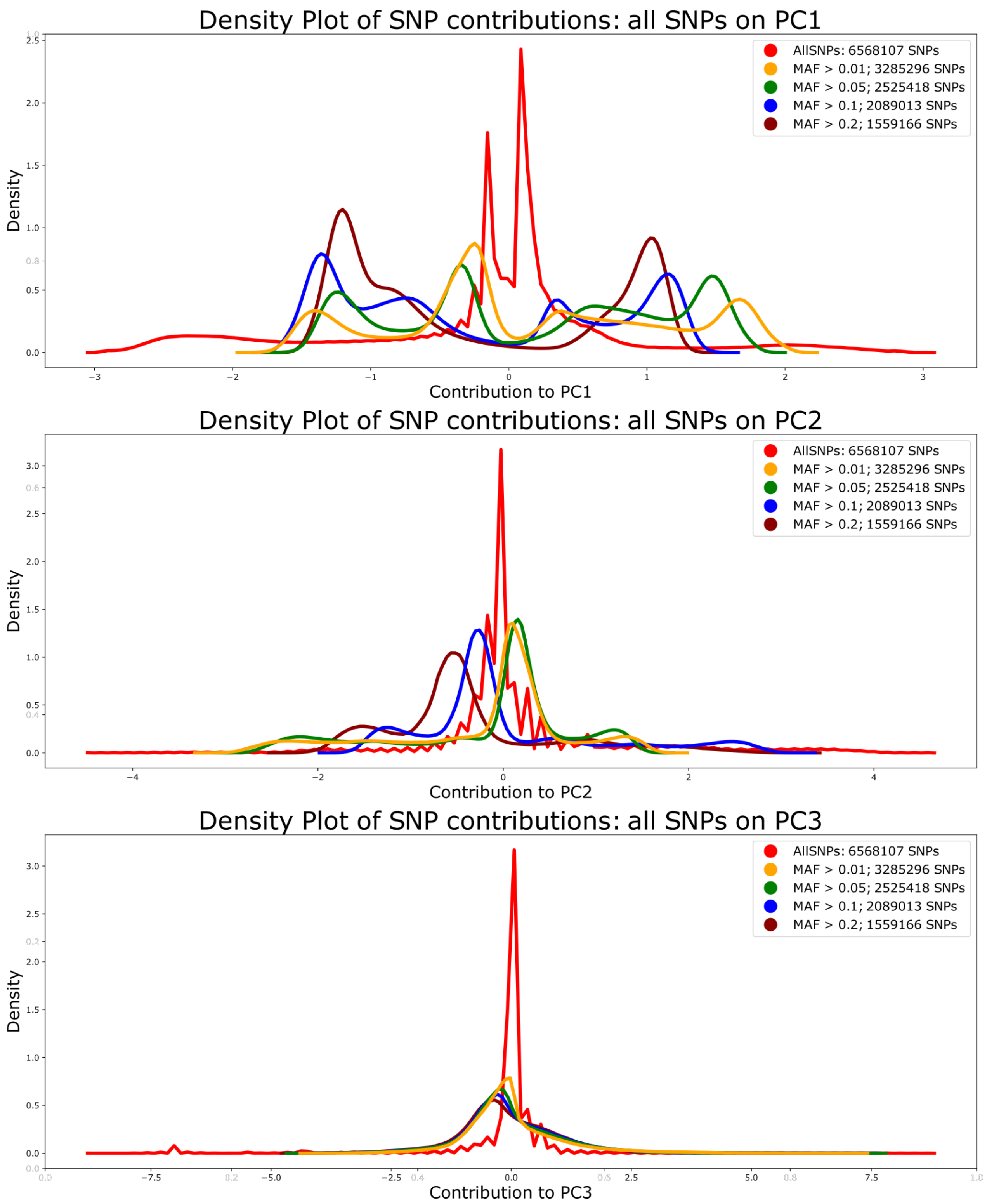


**Contributions of SNPs to PCs 1, 2 and 3, according to MAF filters.**

With no MAF filtering (red lines), there is a high proportion of SNPs contributing very little to PC1 and most SNPs do not contribute to PC2 and PC3. With MAF filtering, the proportion of SNPs contributing to the PCs increases, including for PCs 2 and 3.

---

A large majority of the ~7 million SNPs contribute only very little to PC1 and
the proportion of markers contributing to PCs 2 and 3 is even much smaller.
only a very small proportion contribute slightly to PCs 2 and 3. Clearly, datasets of SNPs with MAF > 0.01 or MAF > 0.05 are sufficient to allow a higher proportion of markers contributing to the PCs, with a notable increase of SNPs contributing to PC2

and PC3. To avoid losing too many potential population-specific markers present at low frequency in the data, we chose to use the dataset of 3,285,296 SNPs having MAF > 0.01 for subsequent analyses.

#### 3.3. SNP contributions along the genome

##### 3.3.1. Genome-wide plots

Inspection of the contributions of SNPs along the genome revealed a striking feature of our dataset, with regions whose size can be as large as one or more Mb, in which a very large majority of the SNPs have a contribution to PC1 superior to the average maximum contribution of all markers. These regions coincide with large haplotype blocks detected with plink. The largest on these blocks spans 3.6 Mb on chromosome 11, which is more than 1.5 % of the honeybee genome size, and four others on chromosomes 4, 7 and 9 are larger than 1 Mb

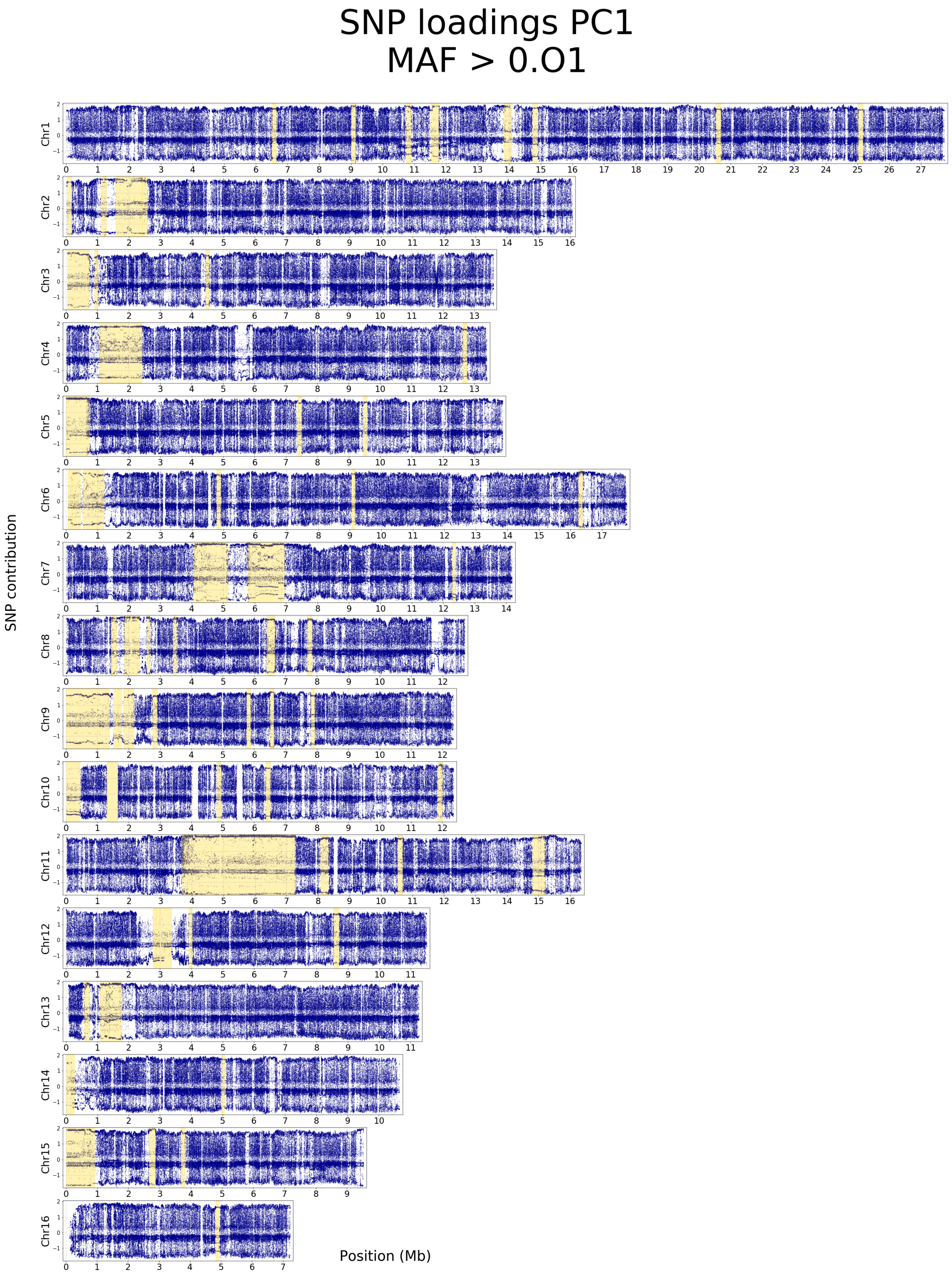


**Contribution of SNPs with MAF > 0.01 to PC1.** Yellow backgrounds correspond to haplotype blocks detected with the plink blocks function, of size larger than 100 kb. SNP contributions were estimated with SMARTPCA.

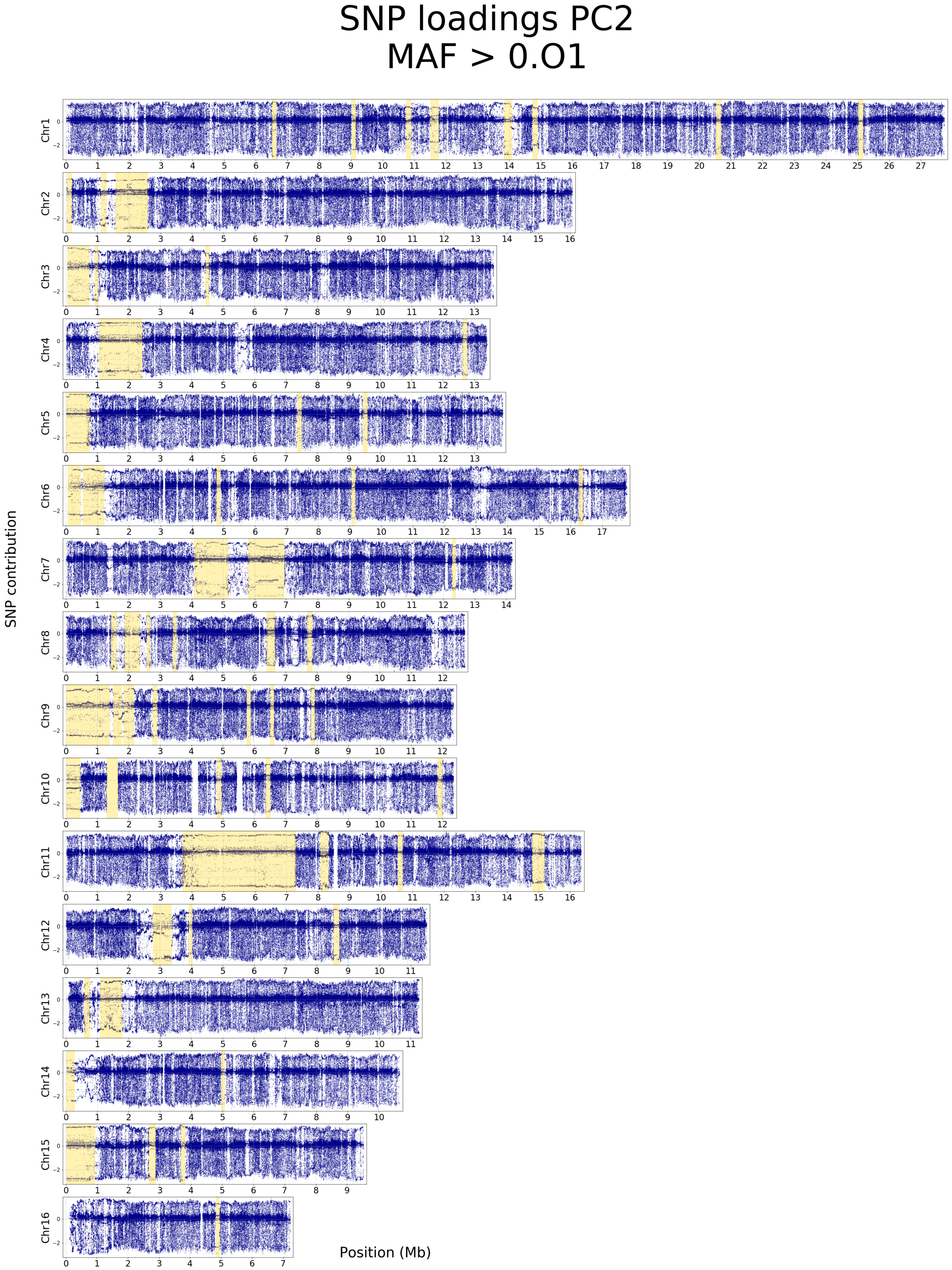


**Contribution of SNPs with MAF > 0.01 to PC2.** Yellow backgrounds correspond to haplotype blocks detected with the plink blocks function, of size larger than 100 kb. SNP contributions were estimated with SMARTPCA.

##### 3.3.2. Plots in haplotype haploBlocks

In most haplotype blocks, almost all SNPs have a very strong contribution to PC1, thus suggesting that these blocks are characterized by long haplotypes specific to either the *A. m. mellifera* and *A. m. iberica* on one side and to all other subspecies on the other. In one region on chromosome 12, representing a part of a haplotype block, almost all SNPs have a very strong contributin to PC2, suggesting that this region is characterized by a long haplotype specific to the *A. m. caucasica* subspecies.

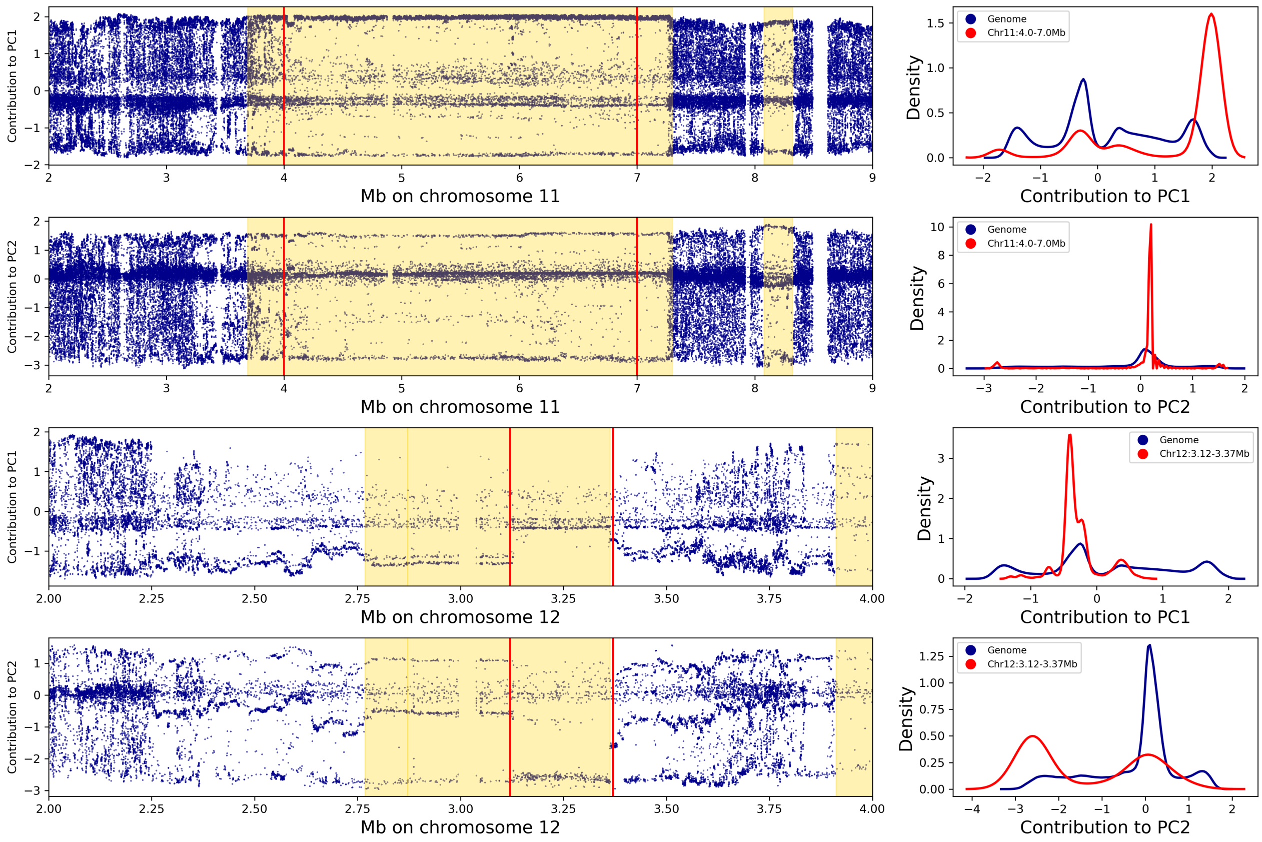


**Contribution of SNPs to PC1 and PC2 in haplotype blocks on chromosomes 11 and 12.** Left: contribution of the individual SNPs to PC1 and PC2. The yellow background indicates haplotype blocks of size larger than 100 kb, as detected by the block command of plink. The red vertical lines delimit the regions selected for plotting SNP contribution densities in the corresponding figures on the right. Right: density plots of SNP contributions to PC1 and 2; blue line: SNPs from all the genome; red line: SNPs from the selected region. In the haplotype block larger than 3 Mb found on chromosome 11, the vast majority of SNPs contribute strongly to PC1, while the contributions to PC2 is virtually inexistent. In a portion of the haplotype block found on chromosome 12, the proportion of SNPs contributing to PC1 is contrarywise lower than average, whereas a majority of SNPs contribution very strongly to PC2.

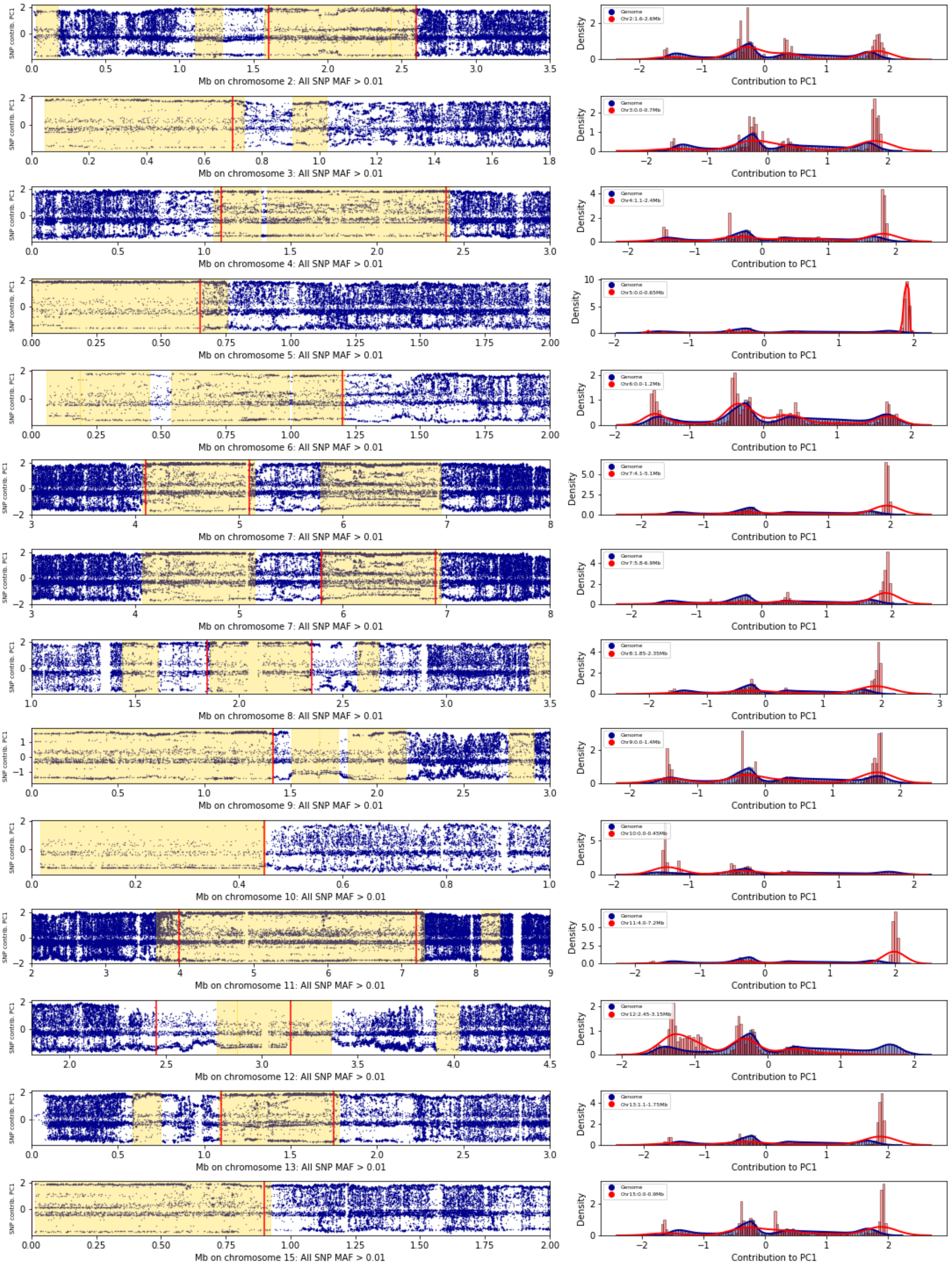


**Contribution of SNPs to PC1 in haplotype blocks.** Some of the other most striking haplotype blocks are shown, showing their very strong contribution to PC1. Left: contribution of the individual SNPs to PC1. The yellow background indicates haplotype blocks of size larger than 100 kb, as detected by the block command of plink. The red vertical lines delimit the regions selected for plotting SNP contribution densities in the corresponding figures on the right. Right: density plots of SNP contributions to PC1; blue: SNPs from all the genome; red: SNPs from the selected region.

### 4. LD pruning

Due to the presence of large haplotype blocks with almost all SNPs having exceptionally high contributions to PC1, we carefully monitored these regions while selecting a suitable threshold for the LD pruning step.

The number of SNPs used in a window for LD pruning was determined such as most windows would correspond to a physical size of 100 kb. The number of SNPs in 100 kb windows vary along the genome and is lower in the haplotype blocks.

#### 4.1. SNP density plots and window size selection

### SNP densities

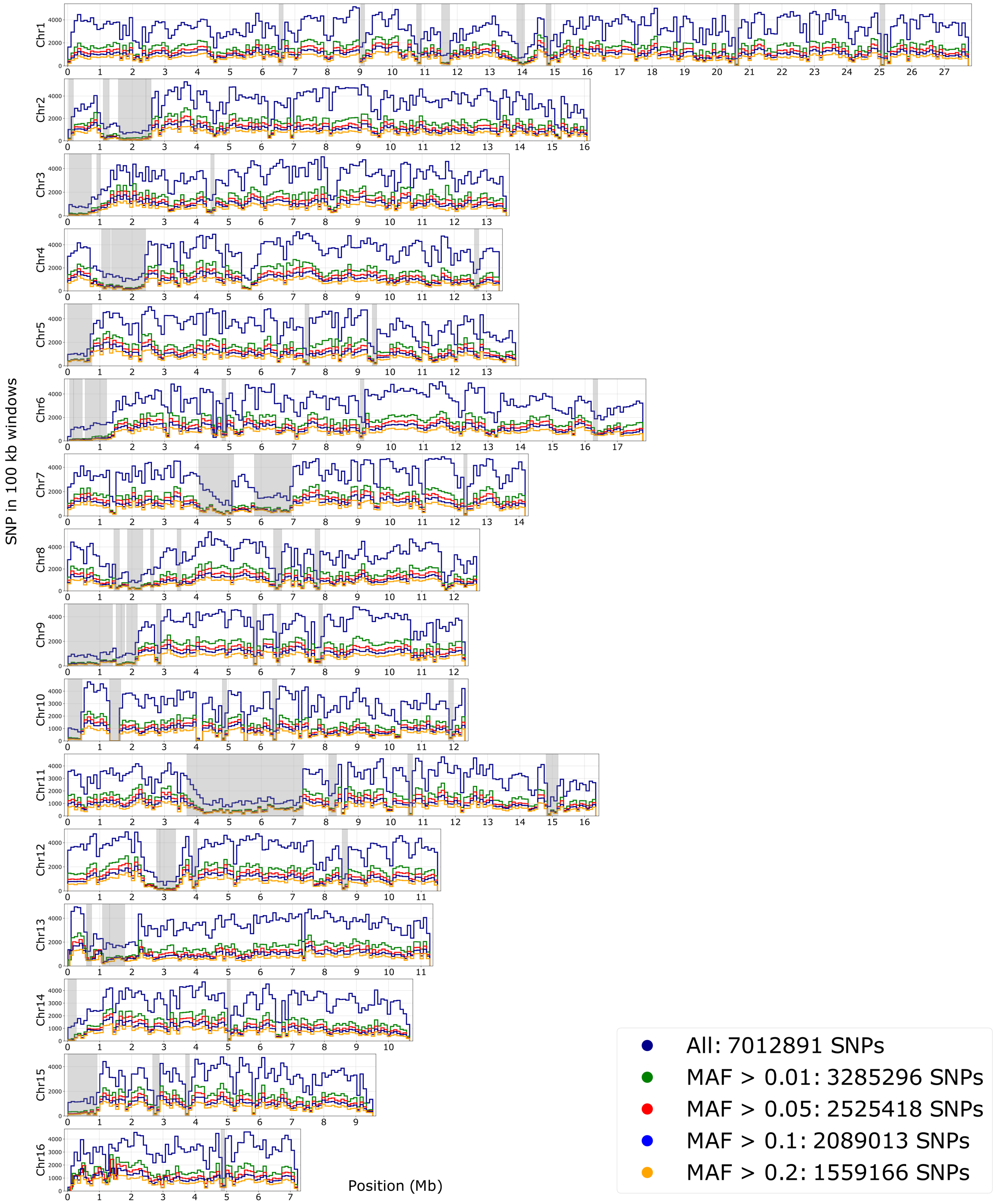


**Plot of SNP densities along the genome in 100 kb windows.** Plots represent SNP counts for all markers and after MAF filters. Grey backgrounds correspond to haplotype blocks detected with the plink blocks function, of size larger than 100 kb.

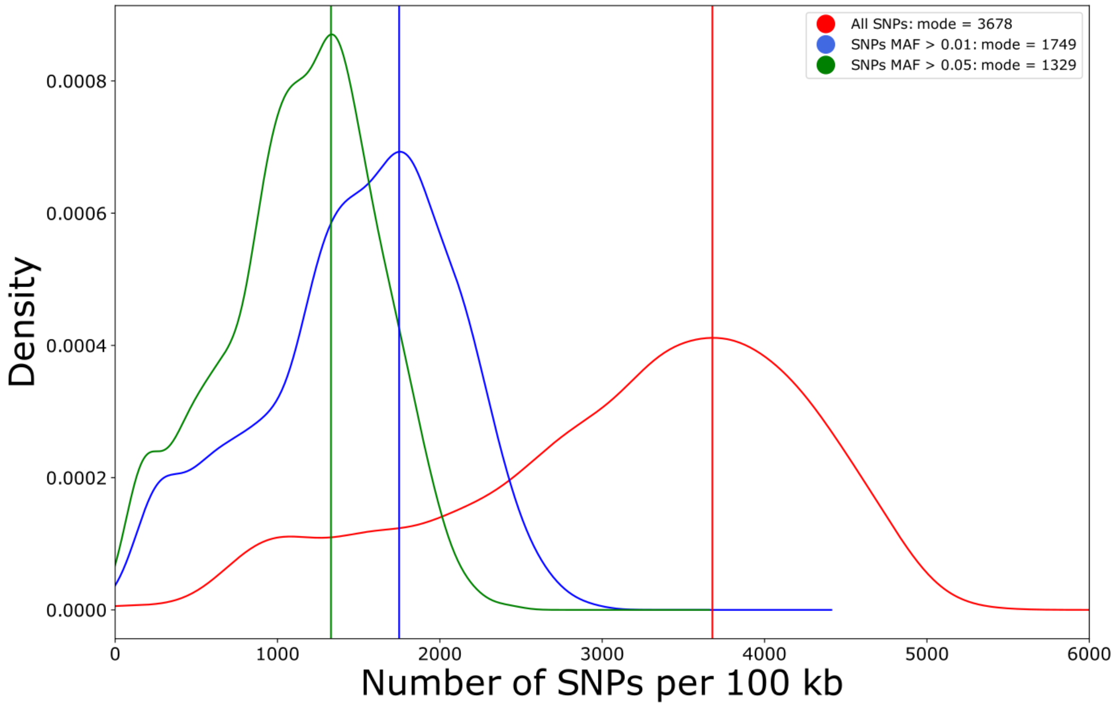


**Density plots of the number of SNPs per 100 kb windows.** The vertical lines represent the value of the mode for the distribution. The mode of the number of SNPs per 100 kb for the dataset of 3,285,296 SNPs with MAF > 0.01 is 1749 (supplementary figures 11 and 12), so pruning was done with a window size of 1749 SNPs and 175 bp (10 %) overlap.

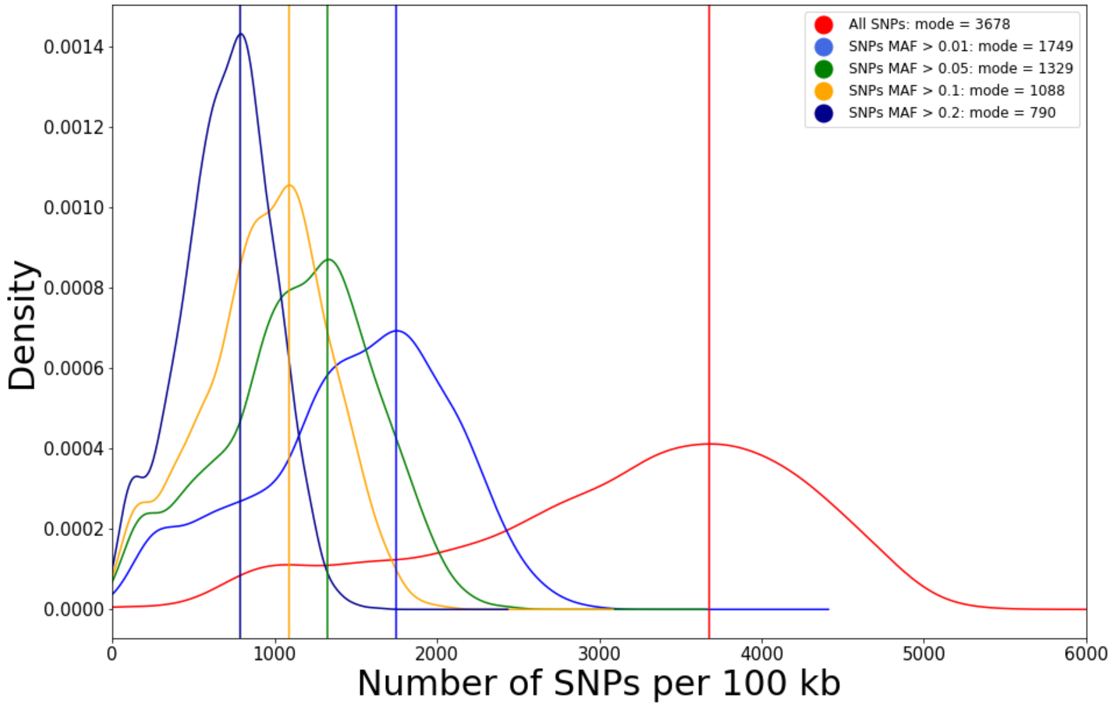


**Density plots of the number of SNPs per 100 kb windows.** The vertical lines represent the value of the mode for the distribution. The mode of the number of SNPs per 100 kb for the dataset of 3,285,296 SNPs with MAF > 0.01 is 1749 (supplementary figures 11 and 12), so pruning was done with a window size of 1749 SNPs and 175 bp (10 %) overlap.

#### 4.2. LD pruning and effects on PCA

##### 4.2.1. Generating datasets (plink)

```
#! /bin/bash

#select629_LD.bash

module load -f /work/project/cytogen/Alain/seqapipopOnHAV3_AV/program_module

NAME1=SeqApiPop_629_maf001
NAME2=SeqApiPop_629_maf001_LD03

plink --bfile ${NAME1} \
  --out ${NAME2} \
  --indep-pairwise 1749 175 0.3

plink --bfile ${NAME1} \
  --out ${NAME2}_pruned \
  --extract ${NAME2}.prune.in \
  --make-bed

plink --bfile ${NAME2}_pruned \
  --out PCA_${NAME2} \
  --pca
```

##### 4.2.2. Effect of LD pruning on SNP contributions to the PCs

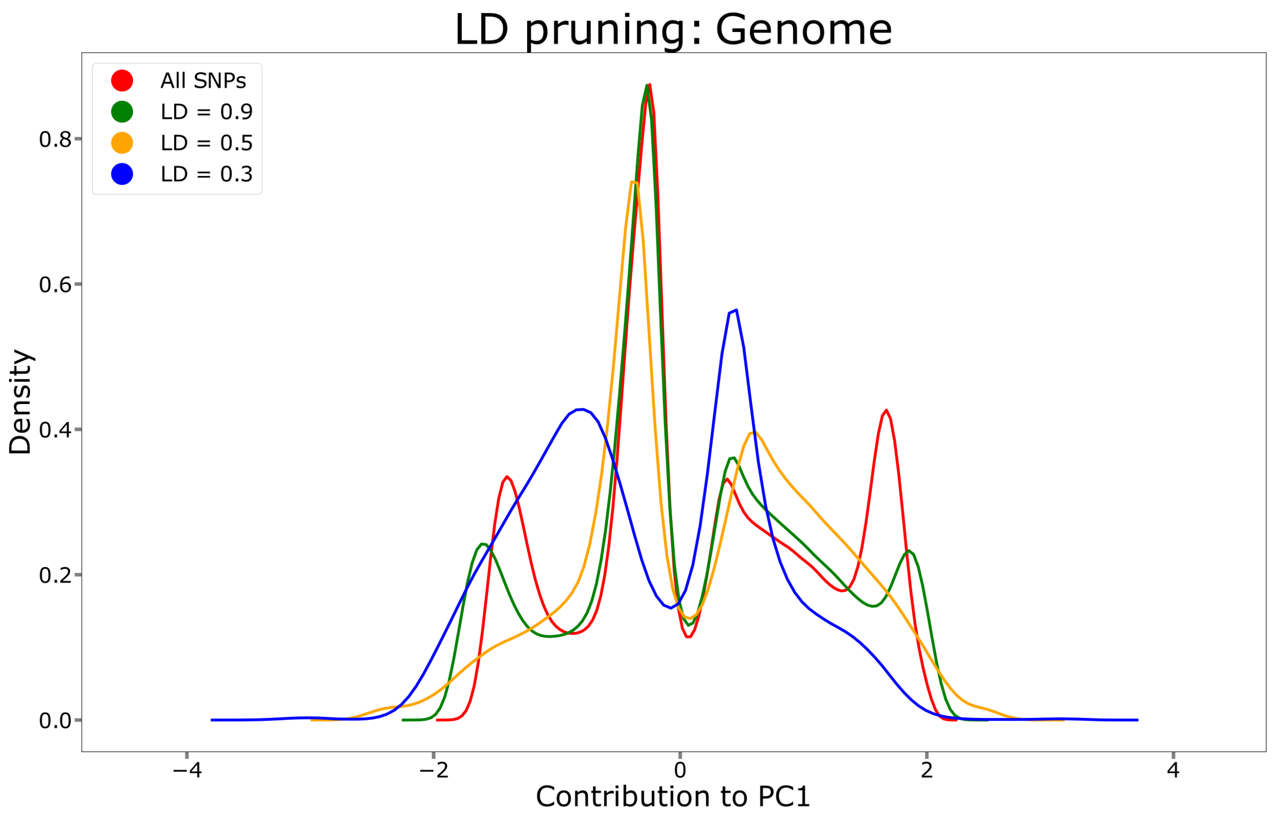


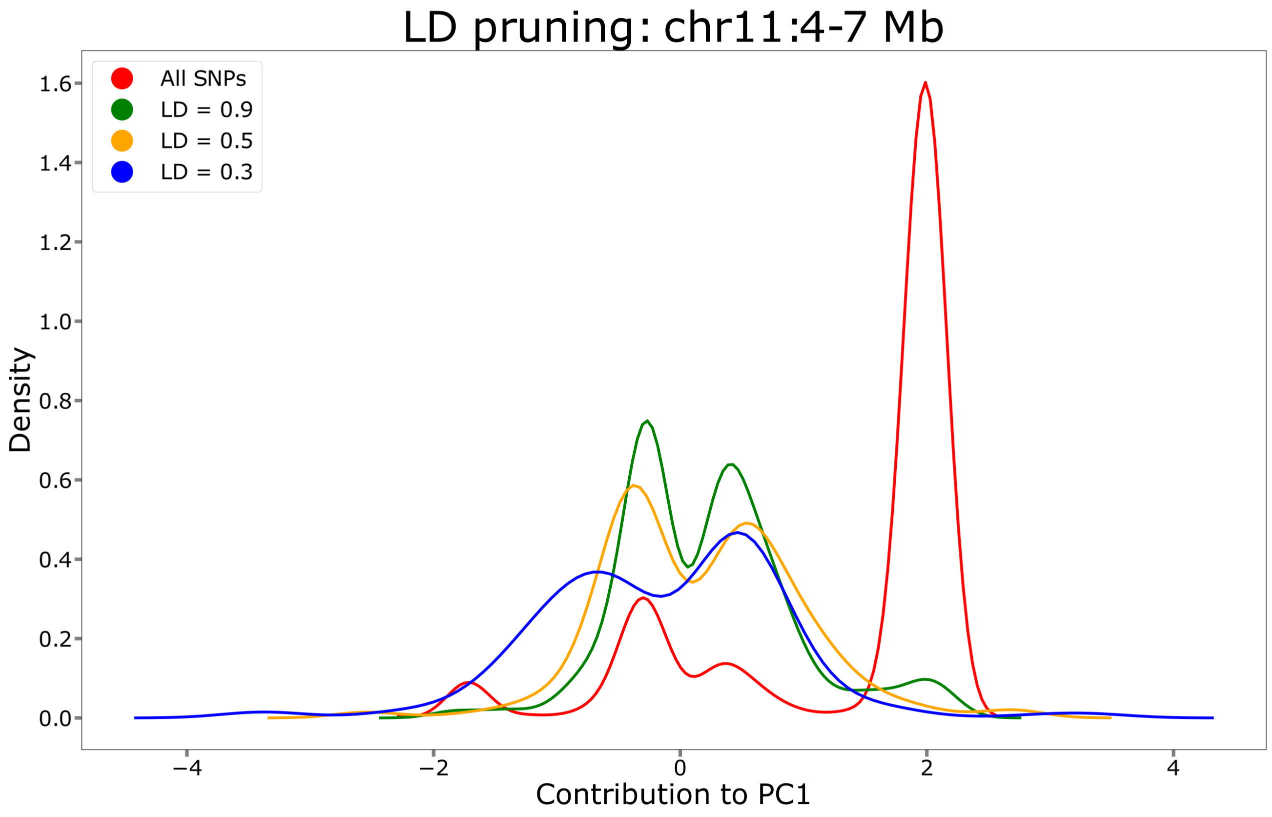


**Effect of LD pruning on SNP contribution to PC1.** Top: on the whole genome; bottom: on the 3 Mb haplotype block on chromosome 11 having a very strong contribution to PC1. LD pruning allows to increase the proportion of markers genome wide contributing to the variance, while efficiently removing the excess of contributing markers in the haplotype block.

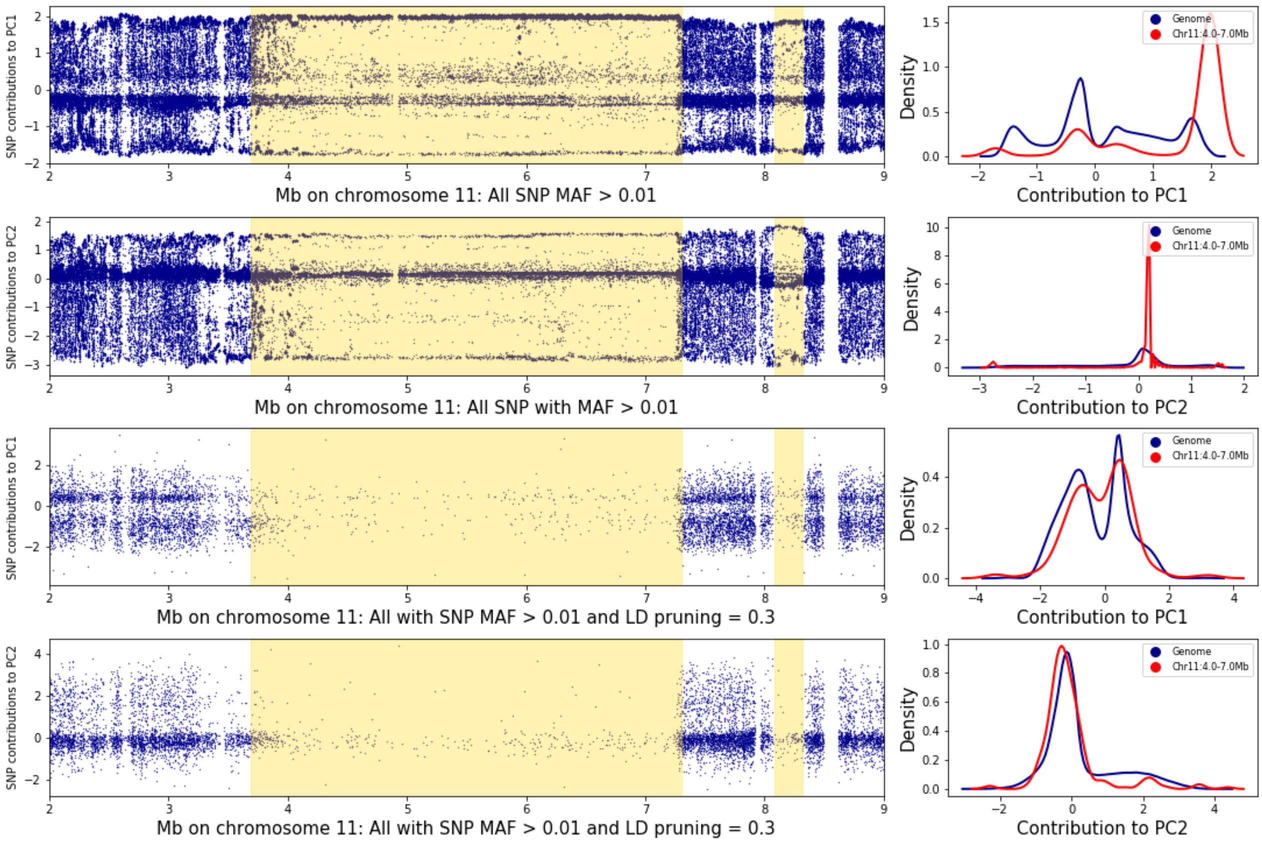


**Figure 14: Effect of LD pruning on SNP density in haplotype blocks.** Left: contribution of the individual SNPs to principal component 1 or 2 on chromosome 11, between positions 2 and 7 Mb before (top) or after LD pruning with LD = 0.3 (bottom); the yellow background indicates haplotype blocks of size larger than 100 kb, as detected by the block command of plink. Right: densities of the SNP contributions to PC1 and 2 with and without LD pruning; blue: SNPs from all the genome; red: SNPs from the haplotype block region detected with plink. The LD = 0.3 pruning value removes most markers from the haplotype block and the density distribution of SNP contributions within the haplotype block now matches that of the whole genome, for both PC1 and PC2..

##### 4.2.3. Effect of LD pruning on PCA

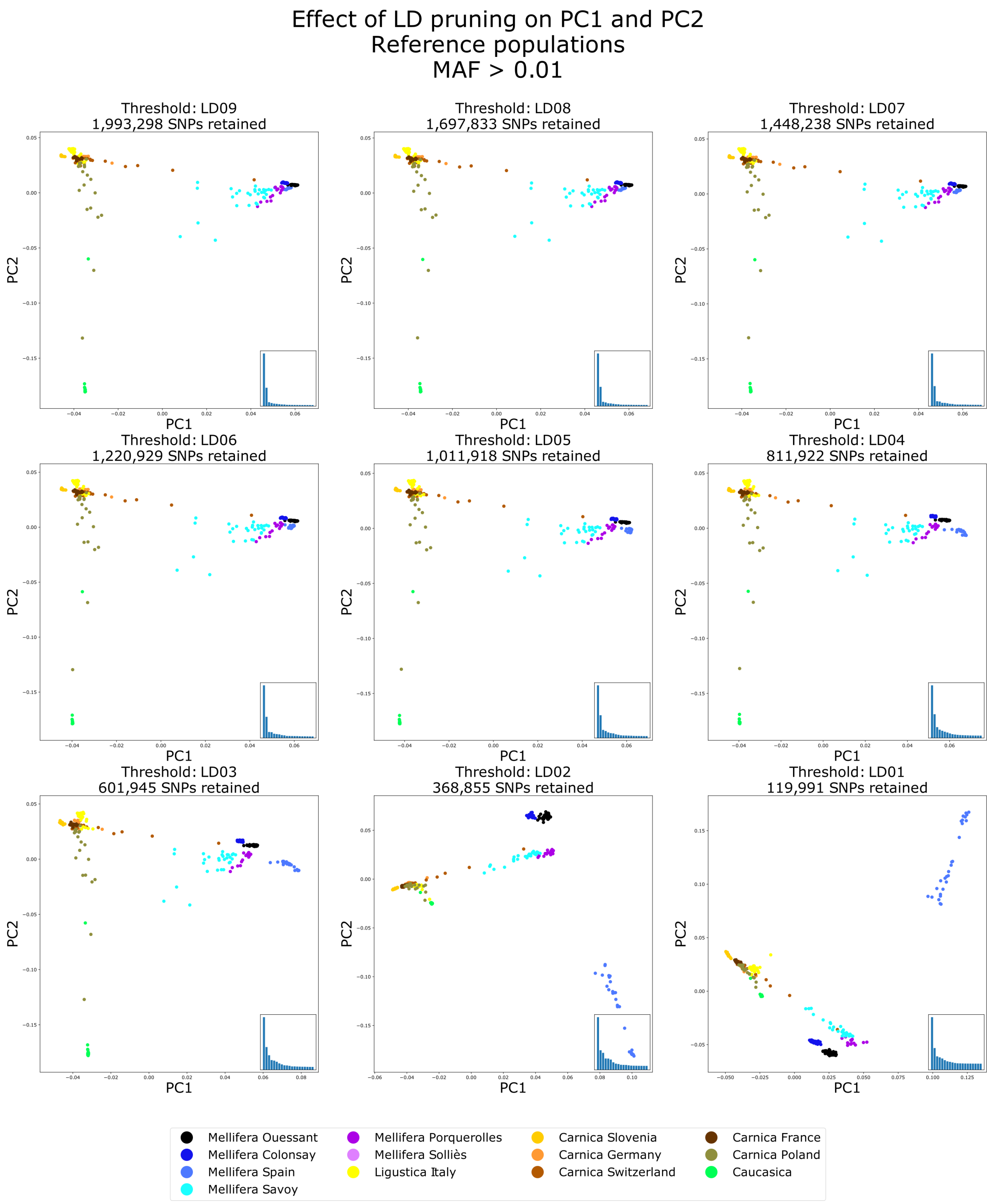


**Effect of LD pruning on PC1 and PC2.**

Only the reference populations are shown. PC1 and PC2 are plotted with datasets resulting from different LD pruning settings. Down to LD = 3, the overall pattern of variance as observed when using all markers is preserved, while it is lost when using lower values. At LD = 0.3, the separation of A. m. mellifera and A. m. iberica populations is improved.
