## Supplementary methods 4 for "Complex population structure and haplotype patterns in Western Europe honey bee from sequencing a large panel of haploid drones"

### SeqApiPop analyses: admixture


- Admixture runs
- Obtain the CV values for plotting:
- Collect and rename all Q matrix Files, for Pong analysis
- Pong analysis: on PC/MAC
  - From Admixture Qfiles to Pong format


#### Admixture runs

Following the MAF and LD pruning analysis, the 601,945 SNPs from the plink files SeqApiPop_629_maf001_LD03_pruned.* were used for the Admixture analysis.

Fifty runs of admixture were done, using a random seed. The 50 folders and the corresponding scripts for running admixture were generated by the script launchAdmixtureRunsWriteScriptsMAF001.bash

```
#!/bin/bash

#launchAdmixtureRunsWriteScriptsMAF001.bash

LD=LD03

for i in $(seq 00 49)
do
mkdir SeqApiPop_629_MAF001_${LD}rep${i}

echo \#!/bin/bash > SeqApiPop_629_MAF001_${LD}rep${i}/admixtureAnalysis.sh
echo
echo \#admixtureAnalysis_multiThread.sh >> SeqApiPop_629_MAF001_${LD}rep${i}/admixtureAnalysis.sh
echo >> SeqApiPop_629_MAF001_${LD}rep${i}/admixtureAnalysis.sh
echo module load -f  /home/gencel/vignal/save/000_ProgramModules/program_module  >> SeqApiPop_629_MAF001_${LD
echo >> SeqApiPop_629_MAF001_${LD}rep${i}/admixtureAnalysis.sh
echo IN=SeqApiPop_629_maf001_${LD}_pruned.bed >> SeqApiPop_629_MAF001_${LD}rep${i}/admixtureAnalysis.sh
echo >> SeqApiPop_629_MAF001_${LD}rep${i}/admixtureAnalysis.sh
echo "for K in 2 3 4 5 6 7 8 9 10 11 12 13 14 15 16;" >> SeqApiPop_629_MAF001_${LD}rep${i}/admixtureAnalysis.
echo do >> SeqApiPop_629_MAF001_${LD}rep${i}/admixtureAnalysis.sh
echo -e sbatch --cpus-per-task=4 --mem-per-cpu=4G \\ >> SeqApiPop_629_MAF001_${LD}rep${i}/admixtureAnalysis.s
echo -e "    -J \${K}admixt -o \${IN}.\${K}.o -e \${IN}.\${K}.e" \\ >> SeqApiPop_629_MAF001_${LD}rep${i}/admi
echo -en '    --wrap="admixture --cv'  >> SeqApiPop_629_MAF001_${LD}rep${i}/admixtureAnalysis.sh
echo -en " -s ${RANDOM}"  >> SeqApiPop_629_MAF001_${LD}rep${i}/admixtureAnalysis.sh
echo  -e ' ../../${IN} ${K} -j4 | tee ${IN}.log${K}"' >> SeqApiPop_629_MAF001_${LD}rep${i}/admixtureAnalysis.
echo done >> SeqApiPop_629_MAF001_${LD}rep${i}/admixtureAnalysis.sh

done
```

Example of output, script written in the folder ~/SeqApiPop_629_MAF001_LD03rep44/admixtureAnalysis.sh:

```
#!/bin/bash
#admixtureAnalysis_multiThread.sh

module load -f /home/gencel/vignal/save/000_ProgramModules/program_module

IN=SeqApiPop_629_maf001_LD03_pruned.bed

for K in 2 3 4 5 6 7 8 9 10 11 12 13 14 15 16;
do
sbatch --cpus-per-task=4 --mem-per-cpu=4G \
    -J ${K}admixt -o ${IN}.${K}.o -e ${IN}.${K}.e \
    --wrap="admixture --cv -s 20145 ../../${IN} ${K} -j4 | tee ${IN}.log${K}"
done
```

### Obtain the CV values for plotting:

```
#!/bin/bash

#obtainCVerrorAllSeqApiPop.bash

for i in `ls | grep ^SeqApiPop`
do
grep CV ${i}/*log* | \
    awk -v var="$i" 'BEGIN{OFS="\t"}{print $3,$4, var}' | \
    sed 's/(K=//' | \
    sed 's/)://' | \
    awk 'BEGIN{FS="_";OFS="\t"}{print $1,$4,$3}' | \
    awk 'BEGIN{OFS="\t"}{print $1,$2,$4,$5}'
done
```

### Collect and rename all Q matrix Files, for Pong analysis

```
#!/bin/bash

#renameQmatrixes.bash

#copies and renames the Q matrix outputs in the directory Qfiles

#Edit

MAF=MAF001

for h in $(seq 3 3)
do
        LD=LD0${h}
        for i in $(seq 30 49)
        do
                for j in `ls SeqApiPop_629_${MAF}_${LD}rep${i}/ | grep Q$`
                do
                        cp SeqApiPop_629_${MAF}_${LD}rep${i}/${j}  Qfiles/${j%.*}.r${i}.Q
                done
                #echo ${i}
        done
done
```

### Pong analysis: on PC/MAC

#### From Admixture Qfiles to Pong format

```
ls  ../Qfiles/* | \
    grep LD0 | \
    awk 'BEGIN{FS="/";OFS="\t"}{print $3, $0}' | \
    awk 'BEGIN{FS=".";OFS="\t"}{print $1"_"$2"_"$3"_"$4, $2, ".."$6"."$7"."$8"."$9}' | \
    awk 'BEGIN{OFS="\t"}{print $1,$2,$3}'
```

Pong couldn't run on All K values computed. As the CV values suggest K=8 or K=9 as the optimum, results for K = 2 to 12 were analysed.

```
conda activate Py27Pong
pong -m Q_fileMaps_MAF001LD03_K2K12_50samples \
    -n popOrder.list \
```

```
    -i ind2pop.list \
    -s 0.98 # Similarity threshold. Default = 0.97
```

With:

```
$ head popOrder.list
Spain   IbericaSpain
Ouessant        MelliferaOuessant
UK      MelliferaColonsay
Porquerolles    MelliferaPorquerolles
Sollies MelliferaSollies
Savoy   SavoyConservatory
Italy   LigusticaItaly
Slovenia        CarnicaSlovenia
CarGermany      CaricaGermany
CarFrance       CaricaFrance
```

```
>head ind2pop.list
Corsica
Corsica
Corsica
Corsica
Corsica
Corsica
Corsica
Corsica
Corsica
Corsica
```
