## Supplementary methods 5 for "Complex population structure and haplotype patterns in Western Europe honey bee from sequencing a large panel of haploid drones"

### SeqApiPop analyses: Treemix


- 1. Files
  - Lists for selection:
  - All samples with > 80 pure backgrounds, plus Corsica:
  - Select samples
  - Calculate frequencies and format for TreeMix
  - Convert to TreeMix information
- Run Treemix
- Estimate the number of migrations with R package OptM
- Plot Treemix trees
  - Done for all Treemix outputs:
  - Estimate mean trees:
    - Plot with figtree v1.4.4


#### 1. Files

##### Lists for selection:

```
Selection from the 629 samples, after Admixture analysis
```

##### All samples with > 80 pure backgrounds, plus Corsica:

```
$ head SeqApiPop_390_9pops.list
AOC10 AOC10
AOC11 AOC11
AOC12 AOC12
AOC14 AOC14

$ head SeqApiPop_390_9pops.fam
Corsica AOC10 0 0 0 -9
Corsica AOC11 0 0 0 -9
Corsica AOC12 0 0 0 -9
Corsica AOC14 0 0 0 -9
```

| Background > 80 | Nb. Samples |
|---|---|
| None | 239 |
| Carnica | 97 |
| RoyalJelly | 54 |
| Corsica | 43 |
| Ouessant | 40 |
| Buckfast | 34 |
| Colonsay | 28 |
| Ligustica | 27 |
| Mellifera | 25 |

| Background > 80 | Nb. Samples |
| --- | --- |
| Iberiensis | 23 |
| Caucasia | 19 |

#### Select samples

selectRefPopInds90.bash

- All SNPs

```
#! /bin/bash
module load -f /work/project/cytogen/Alain/seqapipopOnHAV3_AV/program_module
plink --bfile ../SeqApiPop_629 \
        --out SeqApiPop_324_9pops_AllSNPs \
        --keep SeqApiPop_324_9pops.list \
        --make-bed
```

- SNPs selected on MAF and LD

```
#! /bin/bash
module load -f /work/project/cytogen/Alain/seqapipopOnHAV3_AV/program_module
plink --bfile ../SeqApiPop_629_maf001_LD03_pruned \
        --out SeqApiPop_324_9pops \
        --keep SeqApiPop_324_9pops.list \
        --make-bed
```

Once the selections done, the fam files are over written, so the information on the populations is lost. Must generate them again!

#### Calculate frequencies and format for TreeMix

calculateFrequencies.bash

```
#! /bin/bash

module load -f /work/project/cytogen/Alain/seqapipopOnHAV3_AV/program_module

plink --bfile SeqApiPop_324_9pops \
        --freq \
        --missing \
        --family \
        --out SeqApiPop_324_9pops
gzip SeqApiPop_324_9pops.frq.strat
```

#### Convert to TreeMix information

plink2treemix.py script from: https://github.com/ekirving/ctvt/blob/master/plink2treemix.py

```
sbatch --mem=8G --wrap="..plink2treemix.py SeqApiPop_324_9pops.frq.strat.gz SeqApiPop_324_9pops.frq.gz"
```

### Run Treemix

launchTreemix.bash

```
#!/bin/bash

module load bioinfo/treemix-1.13

for i in $(seq 0 9)
do
    for j in $(seq 0 99)
    do
        sbatch --wrap="treemix -i ../SeqApiPop_324_9pops.frq.gz -bootstrap -seed ${RANDOM} \
                    -k 500 \
                    -m ${i} \
                    -o outstemM${i}_rep${j}"
    done
done
```

#### Estimate the number of migrations with R package OptM

```
Bootstraps90.optm = optM("~/plinkAnalyses/WindowSNPs/TreeMix/bootstraps90")

plot_optM(Bootstraps90.optm, method = "Evanno")
```

#### Plot Treemix trees

```
library(RColorBrewer)
#library(R.utils) Curiosly, seems not to work when R.utils loaded
plot_tree("~/plinkAnalyses/WindowSNPs/TreeMix/bootstraps80/outstemM1_rep69")
```

##### Done for all Treemix outputs:

```
#!/bin/bash

#plotTrees.bash

Rscript plotAll.R
```

```
#plotAll.R

library(RColorBrewer)
#library(R.utils)
source("../plotting_funcs.R")

path = "~/plinkAnalyses/WindowSNPs/TreeMix/bootstraps80/"


for (i in 0:9){
    for (j in 0:99){
        pdf(paste(path,"treePlotM",i,"rep",j,".pdf",sep=""))
        plot_tree(paste(path,"outstemM",i,"_rep",j, sep = ""))
        dev.off()
    }
}
```

##### Estimate mean trees:

SumTrees: Phylogenetic Tree Summarization and Annotation, from the DendroPy plylogenetic package
Sukumaran, J and MT Holder. 2010. DendroPy: a Python library for phylogenetic computing. Bioinformatics 26: 1569-1571.

sumtreeStats.bash

```
#!/bin/bash

module load system/Python-3.6.3

for i in $(seq 0 9)
do
    rm -f ~/WindowSNPs/TreeMix/bootstraps80/outMeanM${i}.tre
    touch ~/WindowSNPs/TreeMix/bootstraps80/outMeanM${i}.tre
    for j in `ls  ~/WindowSNPs/TreeMix/bootstraps90/outstemM${i}_rep*.treeout.gz`
    do
    gunzip -c ${j} | head --l 1 >> ~/WindowSNPs/TreeMix/bootstraps90/outMeanM${i}.tre
    done
    sumtrees.py ~/WindowSNPs/TreeMix/bootstraps80/outMeanM${i}.tre \
        > ~/WindowSNPs/TreeMix/bootstraps80/summaryM${i}.tre
done
```

##### Plot with figtree v1.4.4
