## Supplementary methods 6 for "Complex population structure and haplotype patterns in Western Europe honey bee from sequencing a large panel of haploid drones"

### SeqApiPop analyses: RFMix


- Phasing with shapeit
  - Select SNPs
  - VCFs per chromosome
  - Phasing with Shapeit
    - Phasing
    - convert phased genotypes back to vcf
    - Concatenate VCFs
- RFMix
- Select reference and query Samples
  - Make bcf files
- Make a genetic maps
  - Find crossing-overs in data from Liu et al., 2015
- Run RFMix
  - Couldn't get chromosome 1 to work


#### Phasing with shapeit

##### Select SNPs

- MAF > 1
- Max alleles = 2
- Chromosomes as numbers

```
#!/bin/bash

#makeVCF.bash

#select from the vcf with chromosomes indicated as numbers, for plinkAnalyses
#filter on MAF > 0.01
#nb alleles max = 2

module load bioinfo/samtools-1.10
module load bioinfo/vcftools-0.1.15
module load bioinfo/tabix-0.2.5
module load bioinfo/bcftools-1.9

#Select the 629 samples and filter on MAF001

bcftools view  --samples-file ~/plinkAnalyses/WindowSNPs/RFMix/in/IndsAll629.list \
            --min-af 0.01:minor \
            --max-alleles 2 \
            --output-file ~/plinkAnalyses/WindowSNPs/RFMix/out/SeqApiPop_629_MAF001_diAllelic_plink.vcf.gz \
            --output-type z \
            ~/plinkAnalyses/MetaGenotypesCalled870_raw_snps_allfilter_plink.vcf
bcftools index SeqApiPop_629_MAF001_diAllelic_plink.vcf.gz
```

##### VCFs per chromosome

- Shapeit requires one vcf per chromosome

```
#!/bin/bash

#separateChromosomes.bash

module load bioinfo/samtools-1.10
module load bioinfo/vcftools-0.1.15
```

```
module load bioinfo/tabix-0.2.5
module load bioinfo/bcftools-1.9

#Select the 629 samples and filter on MAF001

for i in  $(seq 1 16)

do

bcftools view  --regions ${i} \
        --output-file ~/plinkAnalyses/WindowSNPs/RFMix/out/SeqApiPop_629_MAF001_diAllelic_plink_chr${i}.vcf.g
        ~/plinkAnalyses/WindowSNPs/RFMix/out/SeqApiPop_629_MAF001_diAllelic_plink.vcf.gz
bcftools index SeqApiPop_629_MAF001_diAllelic_plink_chr${i}.vcf.gz
done

for i in  $(seq 1 16)
do
bcftools index SeqApiPop_629_MAF001_diAllelic_plink_chr${i}.vcf.gz
done
```

### Phasing with Shapeit

#### Phasing

```
#!/bin/bash

#phasingShapeit.bash

module load bioinfo/shapeit.v2.904

for i in  $(seq 3 16)

do

sbatch --mem=50g --wrap="shapeit --input-vcf --force -O SeqApiPop_629_MAF001_diAllelic_phased_chr${i}.vcf"

done
```

#### convert phased genotypes back to vcf

```
#!/bin/bash

#convertToVcf.bash

module load bioinfo/shapeit.v2.904

for i in  $(seq 1 16)

do

sbatch --wrap="shapeit -convert --input-haps SeqApiPop_629_MAF001_diAllelic_phased_chr${i}.vcf \
                --output-vcf SeqApiPop_629_MAF001_diAllelic_phased_chr${i}.vcf"

done

for i in  $(seq 1 16)

do

sbatch --wrap="bgzip SeqApiPop_629_MAF001_diAllelic_phased_chr${i}.vcf"

done

for i in  $(seq 1 16)

do
```

```
sbatch --wrap="bcftools index SeqApiPop_629_MAF001_diAllelic_phased_chr${i}.vcf.gz"

done
```

##### Concatenate VCFs

- make list

```
ls ~/plinkAnalyses/WindowSNPs/RFMix/out/*phased*.vcf.gz > vcf.list

$ head vcf.list
~/plinkAnalyses/WindowSNPs/RFMix/out/SeqApiPop_629_MAF001_diAllelic_phased_chr1.vcf.gz
~/plinkAnalyses/WindowSNPs/RFMix/out/SeqApiPop_629_MAF001_diAllelic_phased_chr2.vcf.gz
~/plinkAnalyses/WindowSNPs/RFMix/out/SeqApiPop_629_MAF001_diAllelic_phased_chr3.vcf.gz
```

The chromosomes were out in the correct order by cut and paste in nano

- Concatenate

```
#!/bin/bash

#concatenateVCFs.bash

module load -f  /home/gencel/vignal/save/000_ProgramModules/program_module

bcftools concat -f ~/plinkAnalyses/WindowSNPs/RFMix/out/vcf.list \
                -o ~/plinkAnalyses/WindowSNPs/RFMix/out/SeqApiPop_629_MAF001_diAllelic_phased_All.vcf.gz \
                -O z

tabix SeqApiPop_629_MAF001_diAllelic_phased_All.vcf.gz
```

- Remove unnecessary Files

```
rm shapeit*
rm *.haps
rm *.sample
rm *chr*
rm SeqApiPop_629_MAF001_diAllelic_plink.vcf.gz*
```

### RFMix

### Select reference and query Samples

Select from an Admixture Q matrix with K = 3 the individuals with > 0.95 pure backgrounds as reference => IndsReference.list
The other samples => IndsQuery.list

##### Make bcf files

- Reference

```
#!/bin/bash

#selectBcfRef.bash

module load bioinfo/bcftools-1.9
```

```
#Select the 629 samples and filter on MAF001

bcftools view  --samples-file ~/plinkAnalyses/WindowSNPs/RFMix/in/IndsReference.list \
        --output-file ~/plinkAnalyses/WindowSNPs/RFMix/out/SeqApiPop_320_MAF001_diAllelic_phased_Ref.bcf \
        --output-type b \
        ~/plinkAnalyses/WindowSNPs/RFMix/out/SeqApiPop_629_MAF001_diAllelic_phased_All.vcf.gz
bcftools index SeqApiPop_320_MAF001_diAllelic_phased_Ref.bcf
```

- Query

```
#!/bin/bash

#selectBcfQuery.bash

module load bioinfo/bcftools-1.9

bcftools view  --samples-file ~/plinkAnalyses/WindowSNPs/RFMix/in/IndsQuery.list \
            --output-file ~/plinkAnalyses/WindowSNPs/RFMix/out/SeqApiPop_309_MAF001_diAllelic_phased_Query.bc
            --output-type b \
            ~/plinkAnalyses/WindowSNPs/RFMix/out/SeqApiPop_629_MAF001_diAllelic_phased_All.vcf.gz
bcftools index SeqApiPop_309_MAF001_diAllelic_phased_Query.bcf
```

### Make a genetic maps

#### Find crossing-overs in data from Liu et al., 2015

Liu H, Zhang X, Huang J, Chen J-Q, Tian D, Hurst LD, et al. Causes and consequences of crossing-over evidenced via a high-resolution recombinational landscape of the honey bee. Genome Biol. 2015 Jan 1;16:15.

Reads from the project SRP043350 (Liu et al., 2015) were retrieved from Short Read Archive (SRA) (https://www.ncbi.nlm.nih.gov/sra), aligned to the reference genome for SNP detection (GATK pipeline): vcf file LiuGenotypesSNPs2allelesForCoSearchDuplOK.vcf.gz

The script find_crossing_overs.py was then used to detect crossing overs:

- -l Colony*.list : list of samples in the Colony
- -q SRR* : the colony's queen
- -c 6 : number of crossing-overs allowed simultaneously over all individuals

```
sbatch --wrap="./find_crossing_overs.py \
                -i combinedVcf/LiuGenotypesSNPs2allelesForCoSearchDuplOK.vcf.gz \
                -l Colony1.list \
                -q SRR1424586 \
                -c 6 \
                -e HAv3_1_Chromosomes.list \
                -o _colo1"

sbatch --wrap="./find_crossing_overs.py \
                -i combinedVcf/LiuGenotypesSNPs2allelesForCoSearchDuplOK.vcf.gz \
                -l Colony2.list \
                -q SRR1425460 \
                -c 6 \
                -e HAv3_1_Chromosomes.list \
                -o _colo2"

sbatch --wrap="./find_crossing_overs.py \
                -i combinedVcf/LiuGenotypesSNPs2allelesForCoSearchDuplOK.vcf.gz \
                -l Colony3.list \
                -q SRR1425476 \
                -c 6 \
```

```
                  -e HAv3_1_Chromosomes.list \
                  -o _colo3"
```

Then a genetic map was constructed from the data:

After importing the 3 files Colony3.list, make table with one line when multiple COs at one position and add column indicating the lines that are involved in double COs.

- A double CO is due either to:
  - A non-crossing-over event marked by a single SNP
  - non-crossing-over events marked by more than one SNP will be detected by the distance to the previous and/or the next CO in the same individual. The distance is set here at 10 kb, as suggested by Liu, et al., 2015.
  - A genotyping error (especially in the case of nb_COs = 1 or 2)
  - nb_COs = number of ofspring from a colony, with COs at the same position
  - Problems in the assembly, such as small inversions, especially in the case of nb_COs > 2)
- Detection of double crossing-overs: two lines must be removed
  - Line for the recombinant before the SNP : the previous and the next vectors are identical
  - Line for the recombinant after the SNP : the current examined vector and the vectors two steps back are identical

```
#!/usr/bin/env python3

import pandas as pd
import numpy as np
import re
import csv

#Import data
colo1 = pd.read_csv('~/MappingLiu/testCOs_intervals_v_colo1', sep = "\t")
colo1['colony'] = 'Colony1'
colo2 = pd.read_csv('~/MappingLiu/testCOs_intervals_v_colo2', sep = "\t")
colo2['colony'] = 'Colony2'
colo3 = pd.read_csv('~/MappingLiu/testCOs_intervals_v_colo3', sep = "\t")
colo3['colony'] = 'Colony3'
colos_all = pd.concat([colo1,colo2,colo3], axis=0)
colos_all_indexed = colos_all.set_index(['chrom','pos_end_CO'])

# Create a table for the detection of double COs.
vectors_table = colos_all[['colony','chrom','event','pos_end_CO','nb_COs','vector']]
vectors_table = vectors_table.astype({'vector': 'str'})
vectors_table = vectors_table.sort_values(by=['colony','chrom','pos_end_CO'], ascending=[True,True,True])

# There is one line per individual when multiple individual have a CO in the same interval. Keep only one.
vectors_table = vectors_table.drop_duplicates()
# Add columns with previous and next vectors for comparison
vectors_table['prev_vector2'] =  vectors_table['vector'].shift(2, fill_value=0)
vectors_table['prev_vector1'] =  vectors_table['vector'].shift(1, fill_value=0)
vectors_table['next_vector'] =  vectors_table['vector'].shift(-1, fill_value=0)

# New column to write results
vectors_table['double_CO'] = "no"

# Select lines corresponding to potential crossing-over events
vectors_table_COs = vectors_table[vectors_table.event == 'crossing_over']
vectors_table_COs = vectors_table_COs.reset_index()

# Mark lines detected as being due to double COs
vectors_table_COs.loc[(vectors_table_COs.prev_vector1 == vectors_table_COs.next_vector),'double_CO'] = "yes"
vectors_table_COs.loc[(vectors_table_COs.prev_vector2 == vectors_table_COs.vector),'double_CO'] = "yes"

# Join the table with the marked double_CO with the imported table
vectors_table_COs_indexed = vectors_table_COs.set_index(['chrom','pos_end_CO'])
vectors_table_COs_indexed = vectors_table_COs_indexed.drop(['colony','event','nb_COs', 'vector','index'],
                            axis=1)
complete_table = colos_all_indexed.join(vectors_table_COs_indexed)
complete_table = complete_table.reset_index()
complete_table
```

```
# Select potential CO events and the columns for visualisation
COs_table = complete_table[complete_table.event == 'crossing_over']
COs_table = COs_table[['chrom','name','pos_start_CO','pos_end_CO','interval','nb_COs',
                        'dist_min_prev_CO','dist_min_next_CO',
                        'dist_max_prev_CO','dist_max_next_CO','double_CO']]

#Eliminate double COs detected by comparing the vectors and NCOs by removing COs at a distance < 10 kb
genet_map = COs_table[(COs_table.dist_max_prev_CO > 10000) &
                        (COs_table.dist_max_next_CO > 10000) &
                        (COs_table.double_CO == 'no')]

#Change genbank chromosome identifiers to numbers
genet_map = genet_map[['chrom','pos_start_CO','pos_end_CO']]
genet_map.columns = ['chrom','left','right']
genet_map.loc[genet_map.chrom == 'NC_037638.1',"chrom"] = 1
genet_map.loc[genet_map.chrom == 'NC_037639.1',"chrom"] = 2
genet_map.loc[genet_map.chrom == 'NC_037640.1',"chrom"] = 3
genet_map.loc[genet_map.chrom == 'NC_037641.1',"chrom"] = 4
genet_map.loc[genet_map.chrom == 'NC_037642.1',"chrom"] = 5
genet_map.loc[genet_map.chrom == 'NC_037643.1',"chrom"] = 6
genet_map.loc[genet_map.chrom == 'NC_037644.1',"chrom"] = 7
genet_map.loc[genet_map.chrom == 'NC_037645.1',"chrom"] = 8
genet_map.loc[genet_map.chrom == 'NC_037646.1',"chrom"] = 9
genet_map.loc[genet_map.chrom == 'NC_037647.1',"chrom"] = 10
genet_map.loc[genet_map.chrom == 'NC_037648.1',"chrom"] = 11
genet_map.loc[genet_map.chrom == 'NC_037649.1',"chrom"] = 12
genet_map.loc[genet_map.chrom == 'NC_037650.1',"chrom"] = 13
genet_map.loc[genet_map.chrom == 'NC_037651.1',"chrom"] = 14
genet_map.loc[genet_map.chrom == 'NC_037652.1',"chrom"] = 15
genet_map.loc[genet_map.chrom == 'NC_037653.1',"chrom"] = 16
genet_map['marker'] = 'yes'
genet_map.head()

#add start chromosomes
chromosomes = sorted(genet_map.chrom.unique().tolist())
zeros = pd.DataFrame(chromosomes, columns = ['chrom'])
zeros['left'] = 0
zeros['right'] = 0
zeros['marker'] = 'yes'
genet_map = pd.concat([genet_map,zeros], axis=0)

# add end of chromosomes
lefts = ['27754200','16089512','13619445','13404451',
         '13896941','17789102','14198698','12717210',
         '12354651','12360052','16352600','11514234',
         '11279722','10670842','9534514','7238532']
ends = pd.DataFrame(list(zip(chromosomes,lefts)), columns = ['chrom','left'])
ends['right'] = ends['left']
ends['marker'] = 'no'
ends = ends.astype({"left": int, "right": int})
genet_map = pd.concat([genet_map,ends], axis=0)

#Define marker positions: one before the first CO, one after the last CO and one between each CO
genet_map = genet_map.sort_values(by=['chrom','left'])
genet_map['next'] = genet_map.left.shift(-1)
genet_map['marker_position'] = (genet_map['left'] + genet_map['next']) / 2
#tried using df['left'], but sometimes, markers not in increasing order to to close proximity!

#Define cM values
marker_cM = list()
cM_count = 0
for idx, row in genet_map.iterrows():
    marker_cM.append(cM_count / 43 * 100) # 43 offspring in the 3 colonies dataset
    if row[3] == 'yes':
        cM_count = cM_count + 1
    elif row[3] == 'no':
        cM_count = 0
genet_map['marker_cM']  =   marker_cM

genetic_map = genet_map[genet_map.marker == 'yes']
genetic_map = genetic_map[['chrom','marker_position','marker_cM']]
genetic_map = genetic_map.astype({'marker_position': 'int'})
```

```
genetic_map.to_csv('~/plinkAnalyses/WindowSNPs/RFMix/in/GenetMap_march_2021_AV.txt', \
                        index=False, header=False, sep="\t")
```

#### Run RFMix

Add population colums to IndsReference.list => IndsPopReference.list

```
head IndsPopReference.list
Ab-PacBio       Black
BER10   Yellow
BER11   Yellow
BER12   Yellow
BER13   Yellow
BER14   Yellow
BER15   Yellow
BER16   Yellow
BER18   Yellow
BER19   Yellow
```

```
#!/bin/bash

#LanceRunRFMix.bash

for i in  $(seq 1 16)
do
sbatch --mem=30g ~/plinkAnalyses/WindowSNPs/RFMix/Pure95/scripts/runRFMixWithGenetMap.bash ${i}

done
```

```
#!/bin/bash

#runRFMixWithGenetMap.bash

module load bioinfo/rfmix-9505bfa

rfmix -f ~/plinkAnalyses/WindowSNPs/RFMix/Pure95/in/SeqApiPop_Pure95_MAF001_diAllelic_phased_Query.bcf \
        -r ~/plinkAnalyses/WindowSNPs/RFMix/Pure95/in/SeqApiPop_Pure95_MAF001_diAllelic_phased_Ref.bcf \
        --chromosome=${1} \
        -m ~/plinkAnalyses/WindowSNPs/RFMix/Pure95/in/IndsPopReference.list \
        -g ~/plinkAnalyses/WindowSNPs/RFMix/Pure95/in/GenetMap_march_2021_AV.txt \
        -o ~/plinkAnalyses/WindowSNPs/RFMix/Pure95/out/SeqApiPopRfmixChr${1}
```

##### Couldn't get chromosome 1 to work

The generation of internal simulation samples for estimating the Conditional Random Field Weight went on for ever

For the other chromosomes, the CRF values used by the software after the simulation were as follow:

```
Loading genetic map for chromosome 2 ...  done
Maximum scoring weight is 53 (91.1)
Loading genetic map for chromosome 3 ...  done
Maximum scoring weight is 24 (94.3)
Loading genetic map for chromosome 4 ...  done
Maximum scoring weight is 31 (90.7)
Loading genetic map for chromosome 5 ...  done
Maximum scoring weight is 44 (94.5)
Loading genetic map for chromosome 6 ...  done
Maximum scoring weight is 78 (93.6)
Loading genetic map for chromosome 7 ...  done
```

```
Maximum scoring weight is 57 (92.6)
Loading genetic map for chromosome 8 ...  done
Maximum scoring weight is 33 (88.4)
Loading genetic map for chromosome 9 ...  done
Maximum scoring weight is 51 (89.9)
Loading genetic map for chromosome 10 ...  done
Maximum scoring weight is 26 (91.2)
Loading genetic map for chromosome 11 ...  done
Maximum scoring weight is 23 (88.4)
Loading genetic map for chromosome 12 ...  done
Maximum scoring weight is 29 (93.2)
Loading genetic map for chromosome 13 ...  done
Maximum scoring weight is 77 (93.8)
Loading genetic map for chromosome 14 ...  done
Maximum scoring weight is 94 (91.6)
Loading genetic map for chromosome 15 ...  done
Maximum scoring weight is 53 (93.5)
Loading genetic map for chromosome 16 ...  done
Maximum scoring weight is 49 (93.8)
```

These are not related to the chromosome size.
The mean value is 48, so chromosome 1 was run with this fixed value.

```
#!/bin/bash

#LanceRunRFMix.bash

for i in  $(seq 1 1)
do
sbatch --mem=30g ~/plinkAnalyses/WindowSNPs/RFMix/Pure95/scripts/runRFMixWithGenetMap.bash ${i}
done
```

```
#!/bin/bash

#runRFMixWithGenetMap_Chr1_CRF_48.bash

module load bioinfo/rfmix-9505bfa

rfmix -f ~/plinkAnalyses/WindowSNPs/RFMix/Pure95/in/SeqApiPop_Pure95_MAF001_diAllelic_phased_Query.bcf \
        -r ~/plinkAnalyses/WindowSNPs/RFMix/Pure95/in/SeqApiPop_Pure95_MAF001_diAllelic_phased_Ref.bcf \
        --chromosome=1 \
        -m ~/plinkAnalyses/WindowSNPs/RFMix/Pure95/in/IndsPopReference.list \
        -g ~/plinkAnalyses/WindowSNPs/RFMix/Pure95/in/GenetMap_march_2021_AV.txt \
        -o ~/plinkAnalyses/WindowSNPs/RFMix/Pure95/out/SeqApiPopRfmixChr1_CRF48 \
        --crf-weight=48
```
