## Supplementary material for "Complex population structure and haplotype patterns in Western Europe honey bee from sequencing a large panel of haploid drones": Scripts: bootstrapingAV_2019_Dec.sh

```
#!/bin/bash

# Modify environment
# load all softwares (bwa, samtools, gatk)

module load -f /home/gencel/vignal/save/000_ProgramModules/program_module

# set -e will cause script to terminate on an error, set +e allows it to continue
set -e

RED='\e[1m\e[91m'
GREEN='\e[0m\e[92m'
SKY='\e[0m\e[96m'
BLUE='\e[1m\e[94m'
NC='\e[0m'

# ==============================================================================
# bootstraping.sh
# ==============================================================================

echo -e "${GREEN}===================================================================="
echo -e "\tBootstraping"
echo -e "\tOne sorted/indexed bam -> One bootstrap bam"
echo -e "====================================================================${NC}"


function usage()
{
	echo -e "Usage:"
	echo -e "\t$0 -i <pathToBam> -o <pathForResults> -s <sampleName> -R <refName> -n <populationEffectif> [OPTIONS]"
	echo -e "NECESSARY"
	echo -e "\t-i pathToBam\t\twhere the bam aligned/sorted/indexed file is"
	echo -e "\t-n populationEffectif\tnumber of individuals in the sample"
	echo -e "\t-o pathForResults\twhere you want to put all output files"
	echo -e "\t-R refName\t\treference genome, fasta format.\n\t\t\t\tIndexing files are already do and tidy in the same folder."
	echo -e "\t-s sampleName\t\tname of the sample"
	echo -e "OPTIONS"
	echo -e "\t-b nbBootstrap\t\tloop number of bootstrap to do [default: 2]"
	echo -e "\t-l interval\t\tinterval to study. Use like this"
	echo -e "\t\t\t\t   chrName"
	echo -e "\t\t\t\t   chrName:begin-end"
	echo -e "\t\t\t\t   myFile.list (list of intervals to study)"
	echo -e "\t\t\t\t[default: NULL]"
	echo -e "\t-h\t\t\tprint this help message"
	echo -e "\t-p ploidy\t\tploidy of the species [default: 2]"
}


#************************************************************************#
# 							INITIALISING VARIABLES
#************************************************************************#
BOOT=2
IN=
OUT=
SAMPLE=
N=
PLOIDY=2
REF=
INTERVAL=

while getopts ":b:i:l:n:o:p:R:s:h" opt
do
	case ${opt} in
		b)	BOOT=${OPTARG} ;;
		h) 
			usage
			exit 0
			;;
		i)	IN=${OPTARG} ;;
		l)	INTERVAL=${OPTARG} ;;
		n)	N=${OPTARG} ;;
		o)	OUT=${OPTARG} ;;
		p)	PLOIDY=${OPTARG} ;;
		R)	REF=${OPTARG} ;;
		s)	SAMPLE=${OPTARG} ;;
	esac
done

if [[ -z ${SAMPLE} ]] | [[ -z ${IN} ]] | [[ -z ${OUT} ]] | [[ -z ${REF} ]] | [[ -z ${N} ]]
then
  usage
  exit 1
fi

mkdir -p ${OUT}

#************************************************************************#
# 							BOOTSTRAPING START
#************************************************************************#
#************************************************************************#
# BQSR based on boostrapping HaplotypeCaller and BaseRecalibrator -> GATK
#************************************************************************#
let PLOIDY_N=${N}*${PLOIDY}
echo "number of indivuals * ploidy: ${PLOIDY_N}"

echo -e "${BLUE}"; date
echo -e "${RED}Step 8: Base Quality Score Recalibration (GATK-BQSR) \n\t${GREEN}Number of iterations: ${BOOT} \n\t${SKY}${IN}/${SAMPLE}_sort.bam \n\t${SKY}${OUT}/${SAMPLE}_bootstrap.bam${NC}"

if [[ -z ${INTERVAL} ]] #-z : string is null, that is, has zero length
then
	for (( n = 0; n <= ${BOOT}; n++ ))
	do
		echo -e "${RED}\tProcessing iteration ${n}${NC}"
		echo -e "${BLUE}"; date
		echo -e "${GREEN}Step 8-1: Sort and Index bootstrap bam (SAM)${NC}"
		samtools sort -T ${OUT}/${SAMPLE}_bootstrap -o ${OUT}/${SAMPLE}_bootstrap.bam ${IN}/${SAMPLE}_sort.bam
		samtools index ${OUT}/${SAMPLE}_bootstrap.bam
		
		echo -e "${BLUE}"; date
		echo -e "${GREEN}Step 8-2: HaplotypeCaller on realn BAM file (GATK)${NC}"
		gatk --java-options "-Xmx4g" HaplotypeCaller \
		  -R ${REF} \
		  -I ${OUT}/${SAMPLE}_bootstrap.bam \
		  --genotyping-mode DISCOVERY \
		  -O ${OUT}/${SAMPLE}_GATK_realn_${n}.vcf \
		  -ploidy ${PLOIDY_N}
		
		echo -e "${BLUE}"; date
		echo -e "${GREEN}Step 8-3: Filter SNP for high quality (GATK)${NC}"
		gatk --java-options "-Xmx4g" VariantFiltration \
		  -R ${REF} \
		  -V ${OUT}/${SAMPLE}_GATK_realn_${n}.vcf \
		  --filter-expression "QD < 2.0 || FS > 60.0 || MQ < 40.0" \
		  --filter-name "HQ_fail" \
		  --genotype-filter-expression "GQ < 90.0" \
		  --genotype-filter-name "GQ_fail" \
		  -O ${OUT}/${SAMPLE}_GATK_realn_filtered_${n}.vcf
		  
		echo -e "${BLUE}"; date
		echo -e "${GREEN}Step 8-4: Extract SNPs from VCF that passed the filters (GATK)${NC}"
		gatk --java-options "-Xmx4g" SelectVariants \
		  -R ${REF} \
		  --variant ${OUT}/${SAMPLE}_${ids}_GATK_realn_filtered_${n}.vcf \
		  -O ${OUT}/${SAMPLE}_GATK_realn_pass_${n}.vcf \
		  --exclude-filtered

		echo -e "${BLUE}"; date
		echo -e "${GREEN}Step 8-5: Analyze patterns of covariation in the sequence dataset${NC}"
		gatk --java-options "-Xmx4g" BaseRecalibrator \
		  -R ${REF} \
		  -I ${OUT}/${SAMPLE}_bootstrap.bam \
		  --known-sites ${OUT}/${SAMPLE}_GATK_realn_pass_${n}.vcf \
		  -O ${OUT}/${SAMPLE}_BQSR_${n}.table
				  
		gatk --java-options "-Xmx4g" ApplyBQSR \
		  -R ${REF} \
		  -I ${OUT}/${SAMPLE}_bootstrap.bam \
		  --bqsr-recal-file ${OUT}/${SAMPLE}_BQSR_${n}.table \
		  -O ${OUT}/${SAMPLE}_tmp1.bam

		echo -e "${BLUE}"; date
		echo -e "${GREEN}Step 8-6: Do a second pass to analyze covariation post-recalibration${NC}"
		gatk --java-options "-Xmx4g" BaseRecalibrator \
		  -R ${REF} \
		  -I ${OUT}/${SAMPLE}_bootstrap.bam \
		  --known-sites ${OUT}/${SAMPLE}_GATK_realn_pass_${n}.vcf \
		  -O ${OUT}/${SAMPLE}_post_BQSR_${n}.table
				  
		gatk --java-options "-Xmx4g" ApplyBQSR \
		  -R ${REF} \
		  -I ${OUT}/${SAMPLE}_tmp1.bam \
		  --bqsr-recal-file ${OUT}/${SAMPLE}_post_BQSR_${n}.table \
		  -O ${OUT}/${SAMPLE}_tmp2.bam

		cp ${OUT}/${SAMPLE}_tmp2.bam ${OUT}/${SAMPLE}_bootstrap.bam
	done
	# Re-index
	samtools index ${OUT}/${SAMPLE}_bootstrap.bam

	# Remove surplus bootstrap files and realn.bam
	rm ${OUT}/${SAMPLE}_tmp.ba*
	rm ${OUT}/${SAMPLE}*BQSR*
	rm ${OUT}/${SAMPLE}_GATK_realn*vcf*
       
else
	# 2 cas
	#	si on donne un fichier
	#	si on donne un nom de chromosome
	echo -e "${SKY}\t${INTERVAL}${NC}"
	case ${INTERVAL} in
		*"list")
			tmp=`basename ${INTERVAL}`
			ids=${tmp/.list/}
			
			samtools view -H ${IN}/${SAMPLE}_sort.bam > ${OUT}/${SAMPLE}_${ids}_bootstrap.bam
			
			while read line
			do
				samtools view ${IN}/${SAMPLE}_sort.bam ${line} >> ${OUT}/${SAMPLE}_${ids}_bootstrap.bam
			done < ${INTERVAL}
			;;
		*)
			ids=${INTERVAL}
			samtools view -bh -o ${OUT}/${SAMPLE}_${ids}_bootstrap.bam ${IN}/${SAMPLE}_sort.bam ${INTERVAL}
			;;
	esac
	
	
	# Boostrap a number of times (default 2)
	for (( n = 0; n < ${BOOT}; n++ ))
	do
		echo -e "${RED}\tProcessing iteration ${n}${NC}"
		
		echo -e "${BLUE}"; date
		echo -e "${GREEN}Step 8-1: Sort and Index bootstrap bam${NC}"
		samtools sort -T ${OUT}/${SAMPLE}_${ids}_bootstrap -o ${OUT}/${SAMPLE}_${ids}_bootstrap.bam ${OUT}/${SAMPLE}_${ids}_bootstrap.bam
		samtools index ${OUT}/${SAMPLE}_${ids}_bootstrap.bam
			
		echo -e "${BLUE}"; date
		echo -e "${GREEN}Step 8-2: HaplotypeCaller on realn BAM file${NC}"
	    gatk --java-options "-Xmx4g" HaplotypeCaller \
	      -R ${REF} \
	      -I ${OUT}/${SAMPLE}_${ids}_bootstrap.bam \
	      --genotyping-mode DISCOVERY \
	      -O ${OUT}/${SAMPLE}_${ids}_GATK_realn_${n}.vcf
			
	    echo -e "${BLUE}"; date
		echo -e "${GREEN}Step 8-3: Filter SNPs for high quality${NC}"
	    gatk --java-options "-Xmx4g" VariantFiltration \
	      -R ${REF} \
	      -V ${OUT}/${SAMPLE}_${ids}_GATK_realn_${n}.vcf \
          --filter-expression "QD < 2.0 || FS > 60.0 || MQ < 40.0" \
          --filter-name "HQ_fail" \
          --genotype-filter-expression "GQ < 90.0" \
          --genotype-filter-name "GQ_fail" \
		  -O ${OUT}/${SAMPLE}_${ids}_GATK_realn_filtered_${n}.vcf
				  
		echo -e "${BLUE}"; date
		echo -e "${GREEN}Step 8-4: Extract SNPs from VCF that passed the filters${NC}"
		gatk --java-options "-Xmx4g" SelectVariants \
		  -R ${REF} \
		  --variant ${OUT}/${SAMPLE}_${ids}_GATK_realn_filtered_${n}.vcf \
		  -O ${OUT}/${SAMPLE}_${ids}_GATK_realn_pass_${n}.vcf \
		  --exclude-filtered

		echo -e "${BLUE}"; date
		echo -e "${GREEN}Step 8-5: Analyze patterns of covariation in the sequence dataset${NC}"
		gatk --java-options "-Xmx4g" BaseRecalibrator \
		  -R ${REF} \
		  -I ${OUT}/${SAMPLE}_${ids}_bootstrap.bam \
		  --known-sites ${OUT}/${SAMPLE}_${ids}_GATK_realn_pass_${n}.vcf \
		  -O ${OUT}/${SAMPLE}_${ids}_BQSR_${n}.table
		  
		gatk --java-options "-Xmx4g" ApplyBQSR \
		  -R ${REF} \
		  -I ${OUT}/${SAMPLE}_${ids}_bootstrap.bam \
		  --bqsr-recal-file ${OUT}/${SAMPLE}_${ids}_BQSR_${n}.table \
		  -O ${OUT}/${SAMPLE}_${ids}_tmp1.bam
    
    	echo -e "${BLUE}"; date
		echo -e "${GREEN}Step 8-6: Do a second pass to analyze covariation post-recalibration${NC}"
		gatk --java-options "-Xmx4g" BaseRecalibrator \
		  -R ${REF} \
		  -I ${OUT}/${SAMPLE}_${ids}_bootstrap.bam \
		  --known-sites ${OUT}/${SAMPLE}_${ids}_GATK_realn_pass_${n}.vcf \
		  -O ${OUT}/${SAMPLE}_${ids}_post_BQSR_${n}.table
		  
		gatk --java-options "-Xmx4g" ApplyBQSR \
		  -R ${REF} \
		  -I ${OUT}/${SAMPLE}_${ids}_tmp1.bam \
		  --bqsr-recal-file ${OUT}/${SAMPLE}_${ids}_post_BQSR_${n}.table \
		  -O ${OUT}/${SAMPLE}_${ids}_tmp2.bam

		cp ${OUT}/${SAMPLE}_${ids}_tmp2.bam ${OUT}/${SAMPLE}_${ids}_bootstrap.bam
	done
	
	# Re-index
	samtools index ${OUT}/${SAMPLE}_${ids}_bootstrap.bam

	# Remove surplus bootstrap files
	rm ${OUT}/${SAMPLE}_${ids}_tmp*.ba*
	rm ${OUT}/${SAMPLE}_${ids}*BQSR*
	rm ${OUT}/${SAMPLE}_${ids}_GATK_realn*vcf*
fi


echo "Finnished: ${BLUE}"`date`


#end of file
```
