## Supplementary material for "Complex population structure and haplotype patterns in Western Europe honey bee from sequencing a large panel of haploid drones": Scripts: callingAV_2019_Dec.sh

```
#!/bin/bash

# set -e will cause script to terminate on an error, set +e allows it to continue
set -e

module load -f  /home/gencel/vignal/save/000_ProgramModules/program_module

RED='\e[1m\e[91m'
GREEN='\e[0m\e[92m'
SKY='\e[0m\e[96m'
BLUE='\e[1m\e[94m'
NC='\e[0m'

# ==============================================================================
# BsMaping.bash
# ==============================================================================

echo -e "${GREEN}===================================================================="
echo -e "\tCalling"
echo -e "\t Many bam -> One bam -> One gvcf"
echo -e "====================================================================${NC}"

function usage()
{
	echo -e "Usage:"
	echo -e "\t$0 -i <pathToInputFile> -o <pathForResults> -e <pathToErrorFile> -s <sampleName> -n <individualsNumber> [OPTIONS]"
	echo -e "NECESSARY"
	echo -e "\t-i pathToInputFile\t\tpath of the input file"
	echo -e "\t-n individualsNumber\t\tpass to an integer"
	echo -e "\t-o pathForResults\t\tpath to the output files"
	echo -e "OPTIONS"
	echo -e "\t-c qualityEncoding\t\tpass to a string, default sanger"
	echo -e "\t-e pathToErrorFile\t\tpath of error files"
	echo -e "\t-l identifyListFile\t\tpass to a string"
	echo -e "\t-p numberOfProcess\t\tnumber of cores (qsub -pe), pass to an integer, default 5"
}
function usage()
{
	echo -e "Usage:"
	echo -e "\t$0 -i <pathToBam> -o <pathForResults> -s <sampleName> -R <refName> -n <populationEffectif> [OPTIONS]"
	echo -e "NECESSARY"
	echo -e "\t-i pathToBam\t\twhere the bam aligned/sorted/indexed file is"
	echo -e "\t-n populationEffectif\tnumber of individuals in the sample"
	echo -e "\t-o pathForResults\twhere you want to put all output files"
	echo -e "\t-R refName\t\treference genome, fasta format.\n\t\t\t\tIndexing files are already do and tidy in the same folder."
	echo -e "\t-s sampleName\t\tname of the sample"
	echo -e "OPTIONS"
	echo -e "\t-b nbBootstrap\t\tloop number of bootstrap to do [default: 2]"
	echo -e "\t-l interval\t\tinterval to study. Use like this"
	echo -e "\t\t\t\t   chrName"
	echo -e "\t\t\t\t   chrName:begin-end"
	echo -e "\t\t\t\t   myFile.list (list of intervals to study)"
	echo -e "\t\t\t\t[default: NULL]"
	echo -e "\t-h\t\t\tprint this help message"
	echo -e "\t-p ploidy\t\tploidy of the species [default: 2]"
	echo -e "\t-t threadsNumber\tnumber of cores (qsub -pe, sbatch --cpus-per-task) [default: 1]"
}

#************************************************************************#
#		Initialising variables
#************************************************************************#
N=
SAMPLE=
IN=
OUT=
REF=
PLOIDY=2
THREADS=1

#From options list
while getopts ":i:n:o:p:R:s:t:h" opt
do
	case ${opt} in
		h) 
			usage
			exit 0
			;;
		i)	IN=${OPTARG} ;;
		n)	N=${OPTARG} ;;
		o)	OUT=${OPTARG} ;;
		p)	PLOIDY=${OPTARG} ;;
		R)	REF=${OPTARG} ;;
		s)	SAMPLE=${OPTARG} ;;
		t)	THREADS=${OPTARG} ;;
	esac
done

if [[ -z ${SAMPLE} ]] | [[ -z ${IN} ]] | [[ -z ${OUT} ]] | [[ -z ${N} ]] | [[ -z ${REF} ]]
then
  usage
  exit 1
fi

mkdir -p ${OUT}
cd ${IN}

#************************************************************************#
# Merge bam file
#************************************************************************#
echo -e "${BLUE}"; date
echo -e "${RED}Step 1: Merge chromosome bam files\n\t${SKY}${IN}/${SAMPLE}_*_bootstrap.bam \n\t${OUT}/mapping/${SAMPLE}_bootstrap.bam${NC}"
ls ${IN}/${SAMPLE}_*_bootstrap.bam > ${IN}/${SAMPLE}_bam.list
# les fichiers seront bien un par ligne : /directory/absolu/file.bam
samtools merge -fb ${IN}/${SAMPLE}_bam.list ${IN}/${SAMPLE}_bootstrap.bam

rm ${IN}/${SAMPLE}_*_bootstrap.ba*

# Sort and index BAM file
#-T PREFIX : Write temporary files to PREFIX.nnnn.bam. This option is required. 
echo -e "${BLUE}"; date
echo -e "${RED}Step 2: Sort and index bam file\n\t${SKY}${OUT}/mapping/${SAMPLE}_bootstrap.bam \n\t${OUT}/mapping/${SAMPLE}.bam${NC}"
samtools sort -@ ${THREADS} -m 2G -T ${IN}/${SAMPLE}_bootstrap -o ${IN}/${SAMPLE}.bam ${IN}/${SAMPLE}_bootstrap.bam
samtools index ${IN}/${SAMPLE}.bam

rm ${IN}/${SAMPLE}_bootstrap.bam

# Calling
echo -e "${BLUE}"; date
echo -e "${RED}Step 3: Give the variants (gvcf)\n\t${SKY}${IN}/${SAMPLE}.bam\n\t${SKY}${OUT}/${SAMPLE}.g.vcf.gz${NC}"
let PLOIDY_N=${N}*${PLOIDY}
echo "number of indivuals * ploidy: ${PLOIDY_N}"

gatk --java-options "-Xmx4g" HaplotypeCaller  \
   -R ${REF} \
   -I ${IN}/${SAMPLE}.bam \
   -O ${OUT}/${SAMPLE}.g.vcf.gz \
   -ERC GVCF \
   -ploidy ${PLOIDY_N}

echo "Finnished: "`date`

#end of file
```
