## Supplementary material for "Complex population structure and haplotype patterns in Western Europe honey bee from sequencing a large panel of haploid drones": Scripts: mappingAV_2019_Dec.sh

```
#!/bin/bash

module load -f  /home/gencel/vignal/save/000_ProgramModules/program_module

# set -e will cause script to terminate on an error, set +e allows it to continue
set -e

RED='\e[1m\e[91m'
GREEN='\e[0m\e[92m'
SKY='\e[0m\e[96m'
BLUE='\e[1m\e[94m'
NC='\e[0m'

# ==============================================================================
# mapping.sh
# ==============================================================================

echo -e "${GREEN}===================================================================="
echo -e "\tMapping"
echo -e "\tTwo paired fastq -> One sorted/indexed bam"
echo -e "====================================================================${NC}"

function usage()
{
	echo -e "Usage:"
	echo -e "\t$0 -i <pathToFastqFiles> -o <pathForResults> -s <sampleName> -R <refName> -C <chrList> -U <contigList> -n <populationEffectif> [OPTIONS]"
	echo -e "NECESSARY"
	echo -e "\t-C chrList\t\tfile with chromosome name and size, 1 by line"
	echo -e "\t-i pathToFastqFiles\twhere are the fastq files"
	echo -e "\t-n populationEffectif\tnumber of individuals in the sample"
	echo -e "\t-o pathForResults\twhere you want to put all output files"
	echo -e "\t-R refName\t\treference genome, fasta format.\n\t\t\t\tIndexing files are already do and tidy in the same folder."
	echo -e "\t-s sampleName\t\tname of the sample"
	echo -e "\t-U contigList\t\tfile with just contig name, 1 by line"
	echo -e "OPTIONS"
	echo -e "\t-b nbBootstrap\t\tloop number of bootstrap to do [default: 2]"
	echo -e "\t-c encodingQual\t\tencoding sequencing qualities [default: illumina1.8]"
	echo -e "\t-e pathOutputError\twhere you want put all job informations"
	echo -e "\t-h\t\t\tprint this help message"
	echo -e "\t-p ploidy\t\tploidy of the species [default: 2]"
}


#************************************************************************#
# 							INITIALISING VARIABLES
#************************************************************************#
BOOT=2
IN=
OUT=
SAMPLE=
REF=
CHROMOSOME=
UNKNOWN=
N=
PLOIDY=2
ERROR=
ENCODING="illumina1.8"

while getopts ":b:C:c:e:i:n:o:p:R:s:U:h" opt
do
	case ${opt} in
		b)	BOOT=${OPTARG} ;;
		C)	CHROMOSOME=${OPTARG} ;;
		c)	ENCODING=${OPTARG} ;;
		e)	ERROR=${OPTARG} ;;
		h) 
			usage
			exit 0
			;;
		i)	IN=${OPTARG} ;;
		n)	N=${OPTARG} ;;
		o)	OUT=${OPTARG} ;;
		p)	PLOIDY=${OPTARG} ;;
		R)	REF=${OPTARG} ;;
		s)	SAMPLE=${OPTARG} ;;
		U)	UNKNOWN=${OPTARG} ;;
	esac
done

if [[ -z ${SAMPLE} ]] | [[ -z ${IN} ]] | [[ -z ${OUT} ]] | [[ -z ${REF} ]] | [[ -z ${CHROMOSOME} ]] | [[ -z ${UNKNOWN} ]] | [[ -z ${N} ]] #-z : string is null, that is, has zero length
then
  usage
  exit 1
fi


# Create folders for storing files
mkdir -p ${OUT}/{mapping,metrics}
mkdir -p ${ERROR}/{bootstraping,calling}
cd ${OUT}

#************************************************************************#
# 							MAPPING START
#************************************************************************#
# Change the fatsq encoding qualities if necessary

case ${ENCODING} in
	"illumina1.3")
		mkdir -p ${IN}/real
		mv ${IN}/${SAMPLE}_R*.fastq.gz ${IN}/real/
		seqtk seq -V -Q64 ${IN}/real/${SAMPLE}_R1.fastq.gz > ${IN}/${SAMPLE}_R1.fastq
		seqtk seq -V -Q64 ${IN}/real/${SAMPLE}_R2.fastq.gz > ${IN}/${SAMPLE}_R2.fastq
		gzip ${IN}/${SAMPLE}_R1.fastq
		gzip ${IN}/${SAMPLE}_R2.fastq
		;;
	"illumina1.8")
		:
		;;
	*)
		echo "If it is another encoding than illumina1.3 or illumina1.8, I invite you to add an exception"
		;;
esac

#For BAM header (read in first line of FASTQ)
header=`zcat ${IN}/${SAMPLE}_R1.fastq.gz | head -1 | cut -c 2-`	# enlève le @ des fastq
TECHNO=${header% *}		# récupère tout avant les derniers :	HWI...6

POP=${SAMPLE%%_*}		# récupère tout avant le premier _		BB2
tmp=${SAMPLE#*_}		# recupere tout apres le premier _		CGATGT_L003
LIB=${tmp%%_*}			# récupère tout avant le premier _		CGATGT

READ1=${IN}/${SAMPLE}_R1.fastq.gz
READ2=${IN}/${SAMPLE}_R2.fastq.gz


#************************************************************************#
# Map to the reference -> bwa mem
#************************************************************************#
echo -e "${BLUE}"; date
echo -e "${RED}Step 1: Map to the reference (BWA)\n\t${SKY}${IN}/${SAMPLE}.fastq.gz \n\t${OUT}/mapping/${SAMPLE}_aligned.bam${NC}"
echo -e "\tRead Group: @RG\tID:${SAMPLE}\tSM:${POP}\tPL:ILLUMINA\tLB:${LIB}\tPU:${TECHNO}"

bwa mem -M -R @RG"\t"ID:${SAMPLE}"\t"SM:${POP}"\t"PL:ILLUMINA"\t"LB:${LIB}"\t"PU:${TECHNO} ${REF} ${READ1} ${READ2} | samtools view -bh -o ${OUT}/mapping/${SAMPLE}_aligned.bam -

#************************************************************************#
# Sort, Mark duplicates and Index -> picard.jar
#************************************************************************#

#Sort Sam file
echo -e "${BLUE}"; date
echo -e "${RED}Step 2: Sort sam file [coordinate] (Picard)\n\t${SKY}${OUT}/mapping/${SAMPLE}_aligned.bam \n\t${OUT}/mapping/${SAMPLE}_sorted.bam${NC}"
java -Xmx4g -jar ${PICARD} SortSam \
	INPUT=${OUT}/mapping/${SAMPLE}_aligned.bam \
	OUTPUT=${OUT}/mapping/${SAMPLE}_sorted.bam \
	SORT_ORDER=coordinate

#Mark Duplicates
echo -e "${BLUE}"; date
echo -e "${RED}Step 3: Mark duplicates (Picard)\n\t${SKY}${OUT}/mapping/${SAMPLE}_sorted.bam \n\t${OUT}/mapping/${SAMPLE}_dedup.bam${NC}"
java -Xmx4g -jar ${PICARD} MarkDuplicates \
    INPUT=${OUT}/mapping/${SAMPLE}_sorted.bam \
    OUTPUT=${OUT}/mapping/${SAMPLE}_dedup.bam \
    METRICS_FILE=${OUT}/metrics/${SAMPLE}_dedup.metrics \
    MAX_FILE_HANDLES=2040

rm ${OUT}/mapping/${SAMPLE}_aligned.*
rm ${OUT}/mapping/${SAMPLE}_sorted.bam

#Build Bam Index for the GATK
echo -e "${BLUE}"; date
echo -e "${RED}Step 4: Sort and index\n\t${SKY}${OUT}/mapping/${SAMPLE}_dedup.bam${NC}"
#java -Xmx4g -jar ${PICARD}/picard.jar BuildBamIndex \
#  INPUT=${OUT}/mapping/${SAMPLE}_dedup.bam
samtools sort -T ${OUT}/mapping/${SAMPLE}_dedup -o ${OUT}/mapping/${SAMPLE}_sort.bam ${OUT}/mapping/${SAMPLE}_dedup.bam
samtools index ${OUT}/mapping/${SAMPLE}_sort.bam

rm ${OUT}/mapping/${SAMPLE}_dedup.ba*
#************************************************************************#
# BQSR based on boostrapping HaplotypeCaller and BaseRecalibrator -> GATK
#************************************************************************#
echo -e "${BLUE}"; date
echo -e "${RED}Step 7: Base Quality Score Recalibration (GATK-BQSR) \n\t${GREEN}Number of iterations: ${BOOT} \n\t${SKY}${OUT}/mapping/${SAMPLE}_sort.bam \n\t${SKY}${OUT}/mapping/${SAMPLE}_bootstrap.bam${NC}"

# Call SNPs > BQSR > Call SNPs > BQSR > Call SNPs > BQSR > Call SNPs

#The following were in Noemie's script
#cp ${OUT}/mapping/${SAMPLE}_sort.bam ${OUT}/mapping/${SAMPLE}_bootstrap.bam
#samtools index ${OUT}/mapping/${SAMPLE}_bootstrap.bam

#rm ${OUT}/mapping/${SAMPLE}_sort.bam*

while read line
do
	c=`cut -d ' ' -f1 <<< ${line}`
	let length=`cut -d ' ' -f2 <<< ${line}`
	
	if [ ${length} -gt 50000000 ]			# taille > 50 Mb
	then
		if [ ${length} -gt 100000000 ]		# taille > 100 Mb
		then
			# divise la taille du chromosome en 3 parties
			# calcule les coordonnees des differents morceaux
			let size=${length}/3
			first_part=${c}:1-${size}			# debut du chromosome jusque size
			let ext1=${size}+1
			let ext2=${ext1}+${size}
			second_part=${c}:${ext1}-${ext2}	# milieu du chromosome
			let ext1=${ext2}+1
			third_part=${c}:${ext1}				# fin du chromosome (on ne met pas la coordonnee finale qui correspond implicitement a la taille)
			
			echo ${first_part}
			echo "module load -f  /home/gencel/vignal/save/000_ProgramModules/program_module; bootstrapingAV_2019_Dec.sh -s ${SAMPLE} -i ${OUT}/mapping -o ${OUT}/bootstraping -R ${REF} -n ${N} -p ${PLOIDY} -l ${first_part} -b ${BOOT} > ${ERROR}/bootstraping/${SAMPLE}_${first_part}_bootstraping.o 2> ${ERROR}/bootstraping/${SAMPLE}_${first_part}_bootstraping.e" >> ${OUT}/${SAMPLE}_bootstrap.array
			
			echo ${second_part}
			echo "module load -f  /home/gencel/vignal/save/000_ProgramModules/program_module; bootstrapingAV_2019_Dec.sh -s ${SAMPLE} -i ${OUT}/mapping -o ${OUT}/bootstraping -R ${REF} -n ${N} -p ${PLOIDY} -l ${second_part} -b ${BOOT} > ${ERROR}/bootstraping/${SAMPLE}_${second_part}_bootstraping.o 2> ${ERROR}/bootstraping/${SAMPLE}_${second_part}_bootstraping.e" >> ${OUT}/${SAMPLE}_bootstrap.array
			
			echo ${third_part}
			echo "module load -f  /home/gencel/vignal/save/000_ProgramModules/program_module; bootstrapingAV_2019_Dec.sh -s ${SAMPLE} -i ${OUT}/mapping -o ${OUT}/bootstraping -R ${REF} -n ${N} -p ${PLOIDY} -l ${third_part} -b ${BOOT} > ${ERROR}/bootstraping/${SAMPLE}_${third_part}_bootstraping.o 2> ${ERROR}/bootstraping/${SAMPLE}_${third_part}_bootstraping.e" >> ${OUT}/${SAMPLE}_bootstrap.array
		else
			# diviser la taille du chromosome en 2 parties
			# calcule les coordonnees des differents morceaux
			let size=${length}/2
			first_part=${c}:1-${size}	# debut du chromosome jusque size
			let ext1=${size}+1
			second_part=${c}:${ext1}	# fin du chromosome (on ne met pas la coordonnee finale qui correspond implicitement a la taille)
			
			echo ${first_part}
			echo "module load -f  /home/gencel/vignal/save/000_ProgramModules/program_module; bootstrapingAV_2019_Dec.sh -s ${SAMPLE} -i ${OUT}/mapping -o ${OUT}/bootstraping -R ${REF} -n ${N} -p ${PLOIDY} -l ${first_part} -b ${BOOT} > ${ERROR}/bootstraping/${SAMPLE}_${first_part}_bootstraping.o 2> ${ERROR}/bootstraping/${SAMPLE}_${first_part}_bootstraping.e" >> ${OUT}/${SAMPLE}_bootstrap.array
			
			echo ${second_part}
			echo "module load -f  /home/gencel/vignal/save/000_ProgramModules/program_module; bootstrapingAV_2019_Dec.sh -s ${SAMPLE} -i ${OUT}/mapping -o ${OUT}/bootstraping -R ${REF} -n ${N} -p ${PLOIDY} -l ${second_part} -b ${BOOT} > ${ERROR}/bootstraping/${SAMPLE}_${second_part}_bootstraping.o 2> ${ERROR}/bootstraping/${SAMPLE}_${second_part}_bootstraping.e" >> ${OUT}/${SAMPLE}_bootstrap.array
		fi
		
	else
		echo ${c} 
		echo "module load -f  /home/gencel/vignal/save/000_ProgramModules/program_module; bootstrapingAV_2019_Dec.sh -s ${SAMPLE} -i ${OUT}/mapping -o ${OUT}/bootstraping -R ${REF} -n ${N} -p ${PLOIDY} -l ${c} -b ${BOOT} > ${ERROR}/bootstraping/${SAMPLE}_${c}_bootstraping.o 2> ${ERROR}/bootstraping/${SAMPLE}_${c}_bootstraping.e" >> ${OUT}/${SAMPLE}_bootstrap.array
	fi
	
done < ${CHROMOSOME}


echo "module load -f /home/gencel/vignal/save/000_ProgramModules/program_module; bootstrapingAV_2019_Dec.sh -s ${SAMPLE} -i ${OUT}/mapping -o ${OUT}/bootstraping -R ${REF} -n ${N} -p ${PLOIDY} -l ${UNKNOWN} -b ${BOOT} > ${ERROR}/bootstraping/${SAMPLE}_unknown_bootstraping.o 2> ${ERROR}/bootstraping/${SAMPLE}_unknown_bootstraping.e" >> ${OUT}/${SAMPLE}_bootstrap.array

sarray --cpus-per-task=1 --mem-per-cpu=10G -J ${SAMPLE}_bootstraping ${OUT}/${SAMPLE}_bootstrap.array


job_id_to_hold=`squeue --name=${SAMPLE}_bootstraping | sed 1d | awk '{print $1}'`

sbatch --dependency=afterok:${job_id_to_hold} --cpus-per-task=1 --mem=10G -J ${SAMPLE}_calling \
	-o ${ERROR}/calling/${SAMPLE}_calling.o -e ${ERROR}/calling/${SAMPLE}_calling.e \
	callingAV_2019_Dec.sh -i ${OUT}/bootstraping -o ${OUT}/calling -s ${SAMPLE} -R ${REF} -n ${N} -p ${PLOIDY} -e ${ERROR}

echo "Finnished: "`date`

#end of file
```
