## Supplementary material for "Complex population structure and haplotype patterns in Western Europe honey bee from sequencing a large panel of haploid drones": Scripts: mergingGenologin.sh

```
#!/bin/bash

module load -f  /home/gencel/vignal/save/000_ProgramModules/program_module

# set -e will cause script to terminate on an error, set +e allows it to continue
set -e

RED='\e[1m\e[91m'
GREEN='\e[0m\e[92m'
SKY='\e[0m\e[96m'
BLUE='\e[1m\e[94m'
NC='\e[0m'

function usage()
{
	echo -e "Usage:"
	echo -e "merging"
}

BAMLIST=
OUT=
SAMPLEMERGED=
T=

echo -e "${BLUE}"; date
echo -e "${RED}Step 1: Merging${NC}"

while getopts ":b:o:s:t" opt
do
	case ${opt} in
		b)	BAMLIST=${OPTARG} ;;
		o)	OUT=${OPTARG} ;;
		s)	SAMPLEMERGED=${OPTARG} ;;
		t)	T=${OPTARG} ;;
	esac
done

echo ${BAMLIST}
echo ${SAMPLEMERGED}
echo ${OUT}
echo ${T}

if [[ -z ${BAMLIST} ]] | [[ -z ${SAMPLEMERGED} ]] | [[ -z ${OUT} ]]
then
	usage
	exit 1
fi

samtools merge -f ${OUT}/${SAMPLEMERGED}/mapping/${SAMPLEMERGED}_aligned.bam -b ${BAMLIST}


#************************************************************************#
# Sort, Mark duplicates and Index -> picard.jar
#************************************************************************#

#Sort Sam file
echo -e "${BLUE}"; date
echo -e "${RED}Step 2: Sort sam file [coordinate] (Picard)\n\t${SKY}${OUT}/${SAMPLEMERGED}/mapping/${SAMPLEMERGED}_aligned.bam \n\t${OUT}/${SAMPLEMERGED}/mapping/${SAMPLEMERGED}_sorted.bam${NC}"
java -Xmx4g -jar ${PICARD} SortSam \
	INPUT=${OUT}/${SAMPLEMERGED}/mapping/${SAMPLEMERGED}_aligned.bam \
	OUTPUT=${OUT}/${SAMPLEMERGED}/mapping/${SAMPLEMERGED}_sorted.bam \
	SORT_ORDER=coordinate

#Mark Duplicates
echo -e "${BLUE}"; date
echo -e "${RED}Step 3: Mark duplicates (Picard)\n\t${SKY}${OUT}/${SAMPLEMERGED}/mapping/${SAMPLEMERGED}_sorted.bam \n\t${OUT}/${SAMPLEMERGED}/mapping/${SAMPLEMERGED}_dedup.bam${NC}"
java -Xmx4g -jar ${PICARD} MarkDuplicates \
    INPUT=${OUT}/${SAMPLEMERGED}/mapping/${SAMPLEMERGED}_sorted.bam \
    OUTPUT=${OUT}/${SAMPLEMERGED}/mapping/${SAMPLEMERGED}_dedup.bam \
    METRICS_FILE=${OUT}/${SAMPLEMERGED}/metrics/${SAMPLEMERGED}_dedup.metrics \
    MAX_FILE_HANDLES=2040

rm ${OUT}/${SAMPLEMERGED}/mapping/${SAMPLEMERGED}_aligned.*
rm ${OUT}/${SAMPLEMERGED}/mapping/${SAMPLEMERGED}_sorted.bam

#Build Bam Index for the GATK
echo -e "${BLUE}"; date
echo -e "${RED}Step 4: Sort and index\n\t${SKY}${OUT}/${SAMPLEMERGED}/mapping/${SAMPLEMERGED}_dedup.bam${NC}"
#java -Xmx4g -jar ${PICARD}/picard.jar BuildBamIndex \
#  INPUT=${OUT}/mapping/${SAMPLEMERGED}_dedup.bam
samtools sort -T ${OUT}/${SAMPLEMERGED}/mapping${SAMPLEMERGED}_dedup -o ${OUT}/${SAMPLEMERGED}/mapping/${SAMPLEMERGED}_sort.bam ${OUT}/${SAMPLEMERGED}/mapping/${SAMPLEMERGED}_dedup.bam
samtools index ${OUT}/${SAMPLEMERGED}/mapping/${SAMPLEMERGED}_sort.bam

rm ${OUT}/${SAMPLEMERGED}/mapping/${SAMPLEMERGED}_dedup.ba*
#************************************************************************#
# Now you have to do BQSR based on boostrapping HaplotypeCaller and BaseRecalibrator -> GATK
#************************************************************************#
#echo -e "${BLUE}"; date
#echo -e "${RED}Step 7: Now you have to do Base Quality Score Recalibration (GATK-BQSR) \n\t${GREEN}Number of iterations: ${BOOT} \n\t${SKY}${OUT}/mapping/${SAMPLEMERGED}_sort.bam \n\t${SKY}${OUT}/mapping/${SAMPLEMERGED}_bootstrap.bam${NC}"

# Call SNPs > BQSR > Call SNPs > BQSR > Call SNPs > BQSR > Call SNPs
#cp ${OUT}/${SAMPLEMERGED}/mapping/${SAMPLEMERGED}_sort.bam ${OUT}/${SAMPLEMERGED}/mapping/${SAMPLEMERGED}_bootstrap.bam
#samtools index ${OUT}/${SAMPLEMERGED}/mapping/${SAMPLEMERGED}_bootstrap.bam

#rm ${OUT}/${SAMPLEMERGED}/mapping/${SAMPLEMERGED}_sort.bam*


#end of file
```
