## Supplementary material for "Complex population structure and haplotype patterns in Western Europe honey bee from sequencing a large panel of haploid drones": Scripts: count_phased_geno.py

```
#!/usr/bin/env python3

# Sonia Eynard, March 2020

import sys

directory=sys.argv[1]
file_in=directory+'/'+sys.argv[2]
file_out=directory+'/count_phased_geno.txt'
myfile=open(file_out,'w')
head='CHROM,POS,unphased,phased,missing'
myfile.write(head+'\n')

X=[]
with open(file_in) as f:
	for l in f:
		H=[]
		X=l.strip().split(' ')
		CHROM=X[0]
		POS=X[1]
		X=X[3:len(X)]
		unphased=X.count('0/0')+X.count('1/1')+X.count('2/2')+X.count('3/3')+X.count('0/1')+X.count('0/2')+X.count('0/3')+X.count('1/2')+X.count('1/3')+X.count('2/3')+X.count('1/0')+X.count('2/0')+X.count('3/0')+X.count('2/1')+X.count('3/1')+X.count('3/2')
		phased=X.count('0|0')+X.count('1|1')+X.count('2|2')+X.count('3|3')+X.count('0|1')+X.count('0|2')+X.count('0|3')+X.count('1|2')+X.count('1|3')+X.count('2|3')+X.count('1|0')+X.count('2|0')+X.count('3|0')+X.count('2|1')+X.count('3|1')+X.count('3|2')
		miss=X.count('./.')+X.count('.|.')
		s=str(CHROM)+','+str(POS)+','+str(unphased)+','+str(phased)+','+str(miss)
		myfile.write(str(s)+'\n')
```
