## Supplementary material for "Complex population structure and haplotype patterns in Western Europe honey bee from sequencing a large panel of haploid drones": Scripts: diagnostic.r

```
#module load system/R-3.5.1
#R

# Sonia Eynard, March 2020

#dir='/work/project/dynagen/seynard/NewVcfCleanUp/results'
args<-commandArgs(TRUE)
library(data.table)
library(ggplot2)
library(reshape2)
library(viridis)
ecdf_fun<-function(x,perc) ecdf(x)(perc)
#obligatory arguments
dir<-args[1]
quantile_prob_above_threshold<-as.numeric(args[2])
kept_above_threshold<-args[3]
args_above=args[4]
ABOVE<-data.frame(filt=unlist(strsplit(kept_above_threshold,'_')),Filt=unlist(strsplit(args_above,'_')),info='above')
quantile_prob_below_threshold<-as.numeric(args[5])
kept_below_threshold<-args[6]
args_below=args[7]
BELOW<-data.frame(filt=unlist(strsplit(kept_below_threshold,'_')),Filt=unlist(strsplit(args_below,'_')),info='below')

filter_info<-rbind(ABOVE,BELOW)
filter_info$Filt<-as.numeric(as.character(filter_info$Filt))
print(filter_info)

plot_filter<-function(infile,filter,value_filter){
	hist(infile[,filter][!is.na(infile[,filter])],breaks=200,main=paste0('Histogram of ',filter),xlab=filter,ylab='Frequency of markers',axes=F,freq=T)
	axis(side=2)
	ticks=c(seq(min(infile[,filter][!is.na(infile[,filter])]),max(infile[,filter][!is.na(infile[,filter])]),length.out=10),value_filter)
	axis(side=1,at=round(ticks,2),labels=round(ticks,2),las=2)
	abline(v=value_filter,lty=2,lwd=2,col='blue')
	plot(ecdf(infile[,filter][!is.na(infile[,filter])]),main=paste0('ECDF ',filter),xlab=filt,ylab='Proportion of markers',axes=F)
	abline(h=quantile_prob,lty=2,lwd=2,col='blue')
	abline(v=value_filter,lty=2,lwd=2,col='blue')
	ticks_x=c(seq(min(infile[,filter][!is.na(infile[,filter])]),max(infile[,filter][!is.na(infile[,filter])]),length.out=10),value_filter)
	axis(side=1,at=round(ticks_x,2),labels=round(ticks_x,2),las=2)
	ticks_y=c(0,0.2,0.4,0.6,0.8,1,quantile_prob)
	axis(side=2,at=round(ticks_y,2),labels=round(ticks_y,2),las=2)
}

option=paste0(filter_info$filt,collapse='_')
pdf(paste0(dir,'/plot_decision_',option,'.pdf'),width=10,height=10)

for(x in 1:nrow(filter_info)){
	filt<-filter_info$filt[x]
	Filt<-filter_info$Filt[x]
	if(filter_info$info[x]=='below'){quantile_prob=quantile_prob_below_threshold
	}else if (filter_info$info[x]=='above'){quantile_prob=quantile_prob_above_threshold}
	info<-fread(paste0(dir,'/info.txt'),data.table=F,header=T)

	if(filt %in% colnames(info)){
		for(i in 5:ncol(info)){info[,i][info[,i]=='.']<-NA
		info[,i]<-as.numeric(info[,i])}
		f<-which(colnames(info)==filt)
		for(i in 5:ncol(info)){
			if(max(info[,i][!is.na(info[,i])])-min(info[,i][!is.na(info[,i])]) > 100){info[,i]<-log10(info[,i])
			colnames(info)[i]<-paste0('log_',colnames(info)[i])
			info[,i][info[,i]=='-Inf']<-min(info[,i][info[,i]!='-Inf'])
			info[,i][info[,i]=='Inf']<-max(info[,i][info[,i]!='Inf'])
			}
			}
		filt<-colnames(info)[f]
		if(Filt==-999){Filt<-quantile(info[,filt][!is.na(info[,filt])],probs=quantile_prob)
		}else {quantile_prob<-ecdf_fun(info[,filt][!is.na(info[,filt])],Filt)}
		print(paste0(filt,' threshold for ',quantile_prob,' quantile ',Filt))
		if(grepl('log',filt)==T){
			if(filter_info$Filt[x]!=-999){Filt<-log10(as.numeric(filter_info$Filt[x]))}
			filt2<-gsub('log_','',filt)
			print(paste0(filt2,' threshold for ',quantile_prob,' quantile ',10^Filt))
			write.table(10^Filt,paste0(dir,'/',filt2,'_value.txt'),row.names=F,col.names=F,quote=F)
		}else{write.table(Filt,paste0(dir,'/',filt,'_value.txt'),row.names=F,col.names=F,quote=F)}
		plot_filter(info,filt,Filt)

	}else if (filt=='allele'){
	## allele (keep up to triallelic)
		allele_star<-info[grepl('\\*',info$ALT),c('CHROM','POS')]
		quadri<-subset(info[,c('CHROM','POS')],nchar(info$ALT)>Filt & grepl('\\*',info$ALT)==F)
		tri<-subset(info[,c('CHROM','POS')],nchar(info$ALT)==Filt & grepl('\\*',info$ALT)==F)
		bi<-subset(info[,c('CHROM','POS')],nchar(info$ALT)==1 & grepl('\\*',info$ALT)==F)
		snp_kept<-rbind(bi,tri)
		snp_not_kept<-rbind(allele_star,quadri)
		write.table(snp_kept,sep='\t',paste0(dir,'/snp_kept_nb_allele.txt'),col.names=F,row.names=F,quote=F)
		write.table(snp_not_kept,sep='\t',paste0(dir,'/snp_not_kept_nb_allele.txt'),col.names=F,row.names=F,quote=F)
	}
}

if ('miss' %in% filter_info$filt & 'het' %in% filter_info$filt){
	if(filter_info$info[filter_info$filt=='miss']=='below'){quantile_prob=quantile_prob_below_threshold
	}else if (filter_info$info[filter_info$filt=='miss']=='above'){quantile_prob=quantile_prob_above_threshold}
## genotypes
	gt_list<-list.files(path=dir,pattern='.gt')
#% missing and % heterozygous
	hetero<-c('0/1','0|1','0/2','0|2','0/3','0|3','1/2','1|2','1/3','1|3','2/3','2|3')
	missi<-c('./.','.|.')
	m<-list()
	h<-list()
	for(i in 1:length(gt_list)){
		print(i)
		gt<-fread(paste0(dir,'/',gt_list[i]),data.table=F,header=F)
		colnames(gt)[1]<-'CHROM'
		colnames(gt)[2]<-'POS'
		m[[i]]<-data.frame(gt$CHROM,gt$POS,rowSums(sapply(gt,`%in%`,missi))/(ncol(gt)-2))
		h[[i]]<-data.frame(gt$CHROM,gt$POS,rowSums(sapply(gt,`%in%`,hetero))/(ncol(gt)-2))
		}
	d_miss<-do.call(rbind,m)
	colnames(d_miss)<-c('CHROM','POS','Miss')
	d_het<-do.call(rbind,h)
	colnames(d_het)<-c('CHROM','POS','Het')
#% missing
	if(filter_info$Filt[filter_info$filt=='miss']==-999){Miss<-quantile(d_miss$Miss,probs=quantile_prob)
	write.table(Miss,paste0(dir,'/miss_value.txt'),row.names=F,col.names=F,quote=F)
	}else {Miss<-filter_info$Filt[filter_info$filt=='miss']
	write.table(Miss,paste0(dir,'/miss_value.txt'),row.names=F,col.names=F,quote=F)
	quantile_prob<-ecdf_fun(d_miss$Miss,Miss)}
	print(paste0('% of missing genotype for ',quantile_prob,' quantile ',Miss))
	plot_filter(d_miss,'Miss',Miss)
	write.table(d_miss,paste0(dir,'/missing.txt'),col.names=T,row.names=F,quote=F)
#% heterozygous
	if(filter_info$Filt[filter_info$filt=='het']==-999){Het<-quantile(d_het$Het,probs=quantile_prob)
	write.table(Het,paste0(dir,'/het_value.txt'),row.names=F,col.names=F,quote=F)
	}else {Het<-filter_info$Filt[filter_info$filt=='het']
	write.table(Het,paste0(dir,'/het_value.txt'),row.names=F,col.names=F,quote=F)
	quantile_prob<-ecdf_fun(d_het$Het,Het)}
	print(paste0('% of heterozygous genotype for ',quantile_prob,' quantile ',Het))
	plot_filter(d_het,'Het',Het)
	write.table(d_het,paste0(dir,'/heterozygous.txt'),col.names=T,row.names=F,quote=F)
}

if ('GQ' %in% filter_info$filt & 'GQfiltered' %in% filter_info$filt){
## Genotype quality
	gq_list<-intersect(list.files(path=dir,pattern='.gq'),list.files(path=dir,pattern='^gq_*'))
	g_v<-list()
	g_p<-list()
	for(i in 1:length(gq_list)){
		print(i)
		gq<-fread(paste0(dir,'/',gq_list[i]),data.table=F,header=F)
		colnames(gq)[1]<-'CHROM'
		colnames(gq)[2]<-'POS'
		gq[gq=='.']<-NA
		GQ<-gq[,3:ncol(gq)]
		GQ<-sapply(GQ,as.numeric)
		gv<-c(GQ)
		gv<-gv[!is.na(gv)]
		x<-vector()
		for(y in 1:100){x[y]<-length(gv[gv==y])}
		g_v[[i]]<-x
	}
	GQ<-do.call(cbind,g_v)
	GQ<-cbind(seq(1,100,by=1),rowSums(GQ))
	GQ<-cbind(GQ,cumsum(GQ[,2]))
	colnames(GQ)<-c('value','GQ','GQ_sum')
	GQ<-as.data.frame(GQ)
	if(filter_info$info[filter_info$filt=='GQ']=='below'){quantile_prob=quantile_prob_below_threshold
	}else if (filter_info$info[filter_info$filt=='GQ']=='above'){quantile_prob=quantile_prob_above_threshold}
	if(filter_info$Filt[filter_info$filt=='GQ']==-999){
	Gq<-max(GQ$value[GQ$GQ_sum<quantile_prob*max(GQ$GQ_sum)])
	write.table(Gq,paste0(dir,'/GQ_value.txt'),row.names=F,col.names=F,quote=F)
	}else {Gq<-filter_info$Filt[filter_info$filt=='GQ']
	write.table(Gq,paste0(dir,'/GQ_value.txt'),row.names=F,col.names=F,quote=F)
	quantile_prob<-GQ$GQ_sum[GQ$value==Gq]/max(GQ$GQ_sum)}
	print(paste0('Genotype quality threshold for ',quantile_prob,' quantile ',Gq))
	plot(GQ[,1:2],type='l',main='Histogram of GQ',xlab='GQ',ylab='Frequency of markers',axes=F)
	axis(side=2)
	ticks=c(seq(0,100,by=10),Gq)
	axis(side=1,at=round(ticks,2),labels=round(ticks,2),las=2)
	abline(v=Gq,lty=2,lwd=2,col='blue')
	plot(GQ$GQ_sum,pch=19,main='ECDF GQ',xlab='GQ',ylab='Proportion of markers',axes=F)
	abline(h=quantile_prob*max(GQ$GQ_sum),lty=2,lwd=2,col='blue')
	abline(v=Gq,lty=2,lwd=2,col='blue')
	ticks_x=c(seq(0,100,by=10),Gq)
	axis(side=1,at=round(ticks_x,2),labels=round(ticks_x,2),las=2)
	ticks_y=c(0,0.2*max(GQ$GQ_sum),0.4*max(GQ$GQ_sum),0.6*max(GQ$GQ_sum),0.8*max(GQ$GQ_sum),1*max(GQ$GQ_sum),quantile_prob*max(GQ$GQ_sum))
	axis(side=2,at=round(ticks_y,2),labels=c(0,0.2,0.4,0.6,0.8,1,quantile_prob),las=2)
#% GQ
	g_p<-list()
	for(i in 1:length(gq_list)){
		print(i)
		gq<-fread(paste0(dir,'/',gq_list[i]),data.table=F,header=F)
		colnames(gq)[1]<-'CHROM'
		colnames(gq)[2]<-'POS'
		GQ<-gq[,3:ncol(gq)]
		GQ<-sapply(GQ,as.numeric)
		g_p[[i]]<-data.frame(gq$CHROM,gq$POS,rowSums(GQ<Gq,na.rm=T)/(ncol(gq)-2))
	}
	d_gq<-do.call(rbind,g_p)
	colnames(d_gq)<-c('CHROM','POS','GQfiltered')
	if(filter_info$info[filter_info$filt=='GQfiltered']=='below'){quantile_prob=quantile_prob_below_threshold
	}else if (filter_info$info[filter_info$filt=='GQfiltered']=='above'){quantile_prob=quantile_prob_above_threshold}
	if(filter_info$Filt[filter_info$filt=='GQfiltered']==-999){GQ_p<-quantile(d_gq$GQfiltered,probs=quantile_prob)
	write.table(GQ_p,paste0(dir,'/GQfiltered_value.txt'),row.names=F,col.names=F,quote=F)
	}else{GQ_p<-filter_info$Filt[filter_info$filt=='GQfiltered']
	write.table(GQ_p,paste0(dir,'/GQfiltered_value.txt'),row.names=F,col.names=F,quote=F)
	quantile_prob<-ecdf_fun(d_gq$GQfiltered,GQ_p)}
	print(paste0('% of genotype quality below threshold ',Gq,' for ',quantile_prob,' quantile ', GQ_p))
	plot_filter(d_gq,'GQfiltered',GQ_p)
	colnames(d_gq)[3]<-paste0('GQfiltered',Gq)
	write.table(d_gq,paste0(dir,'/GQfiltered.txt'),col.names=T,row.names=F,quote=F)
}

if('QUAL' %in% filter_info$filt & 'QD' %in% filter_info$filt){
	info<-fread(paste0(dir,'/info.txt'),data.table=F,header=T)
	for(i in 5:ncol(info)){info[,i][info[,i]=='.']<-0
	info[,i]<-as.numeric(info[,i])}
	for(i in 5:ncol(info)){
		if(max(info[,i][!is.na(info[,i])])-min(info[,i][!is.na(info[,i])]) > 100){info[,i]<-log10(info[,i])
		colnames(info)[i]<-paste0('log_',colnames(info)[i])}
		}
	c_qd<-grep('QD',colnames(info))
	cat_qd<-data.frame(start=seq(0,max(info[,c_qd][!is.na(info[,c_qd])]),by=1),stop=seq(1,(max(info[,c_qd][!is.na(info[,c_qd])])+1),by=1))
	cat_qd$cat<-seq(1,nrow(cat_qd),by=1)
	c_qual<-grep('QUAL',colnames(info))
	cat_qual<-data.frame(start=seq(0,max(info[,c_qual][!is.na(info[,c_qual])]),by=0.1),stop=seq(0.1,(max(info[,c_qual][!is.na(info[,c_qual])])+0.1),by=0.1))
	cat_qual$cat<-seq(1,nrow(cat_qual),by=1)
	info$cat_qual<-NA
	info$cat_qd<-NA
	for(i in 1:nrow(cat_qual)){info[info[,c_qual]>=cat_qual$start[i] & info[,c_qual]<cat_qual$stop[i],'cat_qual']<-cat_qual$cat[i]}
	for(i in 1:nrow(cat_qd)){info[info[,c_qd]>=cat_qd$start[i] & info[,c_qd]<cat_qd$stop[i],'cat_qd']<-cat_qd$cat[i]}
	mat<-table(factor(info$cat_qd,levels=seq(1:nrow(cat_qd))),factor(info$cat_qual,levels=seq(1:nrow(cat_qual))))
	rownames(mat)<-cat_qd$start
	colnames(mat)<-cat_qual$start
	melted_mat<-melt(mat)
	ggplot(data=melted_mat,aes(x=Var2,y=Var1,fill=value))+
		geom_tile()+xlab('log10(QUAL)')+ylab('QD')+
		scale_fill_viridis(option='plasma')
	}
dev.off()
```
