## Supplementary material for "Complex population structure and haplotype patterns in Western Europe honey bee from sequencing a large panel of haploid drones": Scripts: filter.r

```
#module load system/R-3.5.1
#R

# Sonia Eynard, March 2020

#dir='/work/project/dynagen/seynard/GWAS/BeeStrongHAV3_1'
library(VennDiagram)
library(data.table)
library(RColorBrewer)
library(reshape2)
library(VennDiagram)
library(grDevices)
args<-commandArgs(TRUE)
#obligatory arguments
dir<-args[1]
#dir<-'/work/project/dynagen/seynard/GWAS/BeeStrongHAV3_1'
kept_above_threshold<-args[2]
#kept_above_threshold<-"MQ_QUAL_QD_GQ_GQ"
args_above=args[3]
#args_above="39_200_20_10_10_"
ABOVE<-data.frame(filt=unlist(strsplit(kept_above_threshold,'_')),Filt=unlist(strsplit(args_above,'_')),info='above')
kept_below_threshold<-args[4]
#kept_below_threshold<-"FS_SOR_allele_miss_het_GQfiltered"
args_below=args[5]
#args_below="61_4_3_0.05_0.01_0.2"
BELOW<-data.frame(filt=unlist(strsplit(kept_below_threshold,'_')),Filt=unlist(strsplit(args_below,'_')),info='below')
filter_info<-rbind(ABOVE,BELOW)
filter_info<-filter_info[!duplicated(filter_info),]
print(filter_info)
filter_info$Filt<-as.numeric(as.character(filter_info$Filt))

info<-fread(paste0(dir,'/info.txt'),header=T,data.table=F)
if('miss'%in%filter_info$filt){miss<-fread(paste0(dir,'/missing.txt'),header=T,data.table=F)
	info$miss<-miss$Miss}
if('het'%in%filter_info$filt){het<-fread(paste0(dir,'/heterozygous.txt'),header=T,data.table=F)
	info$het<-het$Het}
if('GQ'%in%filter_info$filt & 'GQfiltered'%in%filter_info$filt){gq<-fread(paste0(dir,'/GQfiltered.txt'),header=T,data.table=F)
	info$GQfiltered<-gq[,3]}
for(i in 5:ncol(info)){info[,i][info[,i]=='.']<-0; info[,i]<-as.numeric(info[,i])}

run_filter<-function(filter_list,filter_name,filter_value,threshold_direction,infile,previous_markers_name,previous_markers){
	f_list<-list()
	name_filter<-vector()
	for(i in 1:length(filter_list)){
		if(filter_list[i]=='allele'){
			f_list[[i]]<-paste0(infile$CHROM[nchar(infile$ALT)<=filter_value[filter_name==filter_list[i]] & grepl('\\*',infile$ALT)==F],'_',infile$POS[nchar(infile$ALT)<=filter_value[filter_name==filter_list[i]] & grepl('\\*',infile$ALT)==F])
		}else if(filter_list[i]%in%filter_name & filter_list[i]!='allele'){
			if(threshold_direction[filter_name==filter_list[i]]=='below'){
				name_filter[i]<-paste0(filter_name[filter_name==filter_list[i]],'<',filter_value[filter_name==filter_list[i]])
				f_list[[i]]<-paste0(infile$CHROM[infile[,filter_list[i]]<filter_value[filter_name==filter_list[i]]],'_',infile$POS[infile[,filter_list[i]]<filter_value[filter_name==filter_list[i]]])
			}else if (threshold_direction[filter_name==filter_list[i]]=='above'){
				f_list[[i]]<-paste0(infile$CHROM[infile[,filter_list[i]]>filter_value[filter_name==filter_list[i]]],'_',infile$POS[infile[,filter_list[i]]>filter_value[filter_name==filter_list[i]]])
				name_filter[i]<-paste0(filter_name[filter_name==filter_list[i]],'>',filter_value[filter_name==filter_list[i]])
			}
		}
	}
	f_list<-f_list[lapply(f_list,length)>0]
	name_filter<-name_filter[!is.na(name_filter)]
	if(previous_markers_name!='none'){f_list[[length(f_list)+1]]<-(previous_markers)
		name_filter<-c(name_filter,previous_markers_name)}
	if('allele'%in%filter_list){name_filter<-c('allele',name_filter)}
	myCol<-brewer.pal(length(f_list),"Set2")
	plot<-venn.diagram(
		x=f_list,
		category.names=name_filter,
		filename=NULL,
		output=TRUE,
		cex=0.5,
		lwd=2,
		col=myCol,
		fill=myCol,
		alpha=0.3,
		cat.cex=1,
		margin=c(0.3,0.3,0.3,0.3))
	pdf(paste0(dir,'/venn',paste0(name_filter,collapse='_'),'.pdf'),width=10,height=10)
	grid.draw(plot)
	dev.off()
	int<-Reduce(intersect,f_list)
	return(int)
}

#first venn (mapping quality filter)
filter_list<-c('FS','SOR','MQ','MQRankSum','ReadPosRankSum')
int1<-run_filter(filter_list,filter_info$filt,filter_info$Filt,filter_info$info,info,'none',NA)
#second venn quality
filter_list<-c('QUAL','QD')
int2<-run_filter(filter_list,filter_info$filt,filter_info$Filt,filter_info$info,info,'int1',int1)
#third venn allele
filter_list<-c('allele','miss','het','GQfiltered')
int3<-run_filter(filter_list,filter_info$filt,filter_info$Filt,filter_info$info,info,'int2',int2)

list_final<-colsplit(int3,'.1_',c('CHROM','POS'))
list_final$CHROM<-paste0(list_final$CHROM,'.1')
write.table(list_final,paste0(dir,'/list_kept.txt'),col.names=F,row.names=F,quote=F,sep='\t')

print('count initial VCF')
print(data.table(table(info$CHROM)))
print(nrow(info))

list<-colsplit(int1,'_',c('NC','CHROM','POS'))
list$CHROM<-paste0(list$NC,'_',list$CHROM)
list$NC<-NULL
print('count VCF GATK filters')
print(data.table(table(list$CHROM)))
print(length(int1))

list<-colsplit(int2,'_',c('NC','CHROM','POS'))
list$CHROM<-paste0(list$NC,'_',list$CHROM)
list$NC<-NULL
print('count VCF quality filters')
print(data.table(table(list$CHROM)))
print(length(int2))

list<-colsplit(int3,'_',c('NC','CHROM','POS'))
list$CHROM<-paste0(list$NC,'_',list$CHROM)
list$NC<-NULL
print('count VCF allele and biological filters')
print(data.table(table(list$CHROM)))
print(length(int3))
```
