## Supplementary material for "Complex population structure and haplotype patterns in Western Europe honey bee from sequencing a large panel of haploid drones": Scripts: filter_list.r

```
#module load system/R-3.5.1
#R

# Sonia Eynard, March 2020

#dir='/work/project/dynagen/seynard/NewVcfCleanUp/results'
args<-commandArgs(TRUE)
library(data.table)
library(ggplot2)
library(reshape2)
library(viridis)
#obligatory arguments
dir<-args[1]
filt<-args[2]
limits<-args[3]
filter_info<-data.frame(filt=unlist(strsplit(filt,'_')),Filt=unlist(strsplit(limits,'_')))
filter_info$Filt<-as.numeric(as.character(filter_info$Filt))
print(filter_info)
option=paste0(filter_info$filt,collapse='_')

if (filt=='allele'){
		allele_star<-info[grepl('\\*',info$ALT),c('CHROM','POS')]
		quadri<-subset(info[,c('CHROM','POS')],nchar(info$ALT)>Filt & grepl('\\*',info$ALT)==F)
		tri<-subset(info[,c('CHROM','POS')],nchar(info$ALT)==Filt & grepl('\\*',info$ALT)==F)
		bi<-subset(info[,c('CHROM','POS')],nchar(info$ALT)==1 & grepl('\\*',info$ALT)==F)
		snp_kept<-rbind(bi,tri)
		snp_not_kept<-rbind(allele_star,quadri)
		write.table(snp_kept,sep='\t',paste0(dir,'/snp_kept_nb_allele.txt'),col.names=F,row.names=F,quote=F)
		write.table(snp_not_kept,sep='\t',paste0(dir,'/snp_not_kept_nb_allele.txt'),col.names=F,row.names=F,quote=F)
	}

if ('miss' %in% filter_info$filt & 'het' %in% filter_info$filt){
	gt_list<-list.files(path=dir,pattern='.gt')
	het<-c('0/1','0|1','0/2','0|2','0/3','0|3','1/2','1|2','1/3','1|3','2/3','2|3')
	m<-list()
	h<-list()
	for(i in 1:length(gt_list)){
		print(i)
		gt<-fread(paste0(dir,'/',gt_list[i]),data.table=F,header=F)
		colnames(gt)[1]<-'CHROM'
		colnames(gt)[2]<-'POS'
		m[[i]]<-data.frame(gt$CHROM,gt$POS,rowSums(gt%in%c('./.','.|.'))/(ncol(gt)-2))
		h[[i]]<-data.frame(gt$CHROM,gt$POS,rowSums(sapply(gt,`%in%`,het))/(ncol(gt)-2))
		}
	d_miss<-do.call(rbind,m)
	colnames(d_miss)<-c('CHROM','POS','Miss')
	d_het<-do.call(rbind,h)
	colnames(d_het)<-c('CHROM','POS','Het')
	write.table(d_miss,paste0(dir,'/missing.txt'),col.names=T,row.names=F,quote=F)
	write.table(d_het,paste0(dir,'/heterozygous.txt'),col.names=T,row.names=F,quote=F)
}

if ('GQ' %in% filter_info$filt & 'GQfiltered' %in% filter_info$filt){
## Genotype quality
	gq_list<-intersect(list.files(path=dir,pattern='.gq'),list.files(path=dir,pattern='^gq_*'))
	g_v<-list()
	g_p<-list()
	for(i in 1:length(gq_list)){
		print(i)
		gq<-fread(paste0(dir,'/',gq_list[i]),data.table=F,header=F)
		colnames(gq)[1]<-'CHROM'
		colnames(gq)[2]<-'POS'
		gq[gq=='.']<-NA
		GQ<-gq[,3:ncol(gq)]
		GQ<-sapply(GQ,as.numeric)
		gv<-c(GQ)
		gv<-gv[!is.na(gv)]
		x<-vector()
		for(y in 1:100){x[y]<-length(gv[gv==y])}
		g_v[[i]]<-x
	}
	GQ<-do.call(cbind,g_v)
	GQ<-cbind(seq(1,100,by=1),rowSums(GQ))
	GQ<-cbind(GQ,cumsum(GQ[,2]))
	colnames(GQ)<-c('value','GQ','GQ_sum')
	GQ<-as.data.frame(GQ)
	Gq<-filter_info$Filt[filter_info$filt=='GQ']
#% GQ
	g_p<-list()
	for(i in 1:length(gq_list)){
		gq<-fread(paste0(dir,'/',gq_list[i]),data.table=F,header=F)
		colnames(gq)[1]<-'CHROM'
		colnames(gq)[2]<-'POS'
		GQ<-gq[,3:ncol(gq)]
		GQ<-sapply(GQ,as.numeric)
		g_p[[i]]<-data.frame(gq$CHROM,gq$POS,rowSums(GQ<Gq,na.rm=T)/(ncol(gq)-2))
	}
	d_gq<-do.call(rbind,g_p)
	colnames(d_gq)<-c('CHROM','POS','GQfiltered')
	write.table(d_gq,paste0(dir,'/GQfiltered.txt'),col.names=T,row.names=F,quote=F)
}
```
