## Supplementary material for "Complex population structure and haplotype patterns in Western Europe honey bee from sequencing a large panel of haploid drones": Scripts: run_vcf_cleanup.sh

```
#!/bin/bash
### vcf filter (only done once to prepare list SNP positions) ###

# Sonia Eynard, March 2020

############################################ modules #####################################################
module load system/R-3.5.1
module load bioinfo/bcftools-1.6
module load bioinfo/tabix-0.2.5
module load bioinfo/vcftools-0.1.15
module load bioinfo/samtools-1.8
##########################################################################################################

############################################ parameters ##################################################
username=seynard
SCRIPTS='/work/project/dynagen/seynard/scriptGWAS/vcf_cleanup'
DIRIN='/work/project/cytogen/Alain/seqapipopOnHAV3_1/combineGVCFs/The870vcf'
DIROUT='/work/project/dynagen/seynard/GWAS/BeeStrongHAV3_1'
VCFIN='MetaGenotypesCalled870_raw_snps.vcf.gz' #vcf avant filtres
limit_allele=3 #accept up to tri-allelic
#for all filter threshold if x=-999 the value will be assigned eliminate quantile_prob*100% of the data
##########################################################################################################

mkdir -p ${DIROUT}/log

##########################################################################################################
# run diagnostic plots
# adjust values
##########################################################################################################
limit_FS=-999
limit_SOR=-999
limit_MQ=-999
limit_MQRankSum=-999
limit_ReadPosRankSum=-999
limit_QUAL=-999
limit_QD=-999
limit_GQ=-999
limit_miss=-999
limit_het=-999
limit_GQfiltered=-999
quantile_prob_above_threshold=0.1
quantile_prob_below_threshold=0.9
kept_above_threshold="MQ_MQRankSum_ReadPosRankSum_QUAL_QD~GQ~GQ"
kept_below_threshold="FS_SOR_allele~miss_het~GQfiltered"
run='diagnostic'
##########################################################################################################
sbatch -W -J vcf_cleanup -o ${DIROUT}/log/vcf_cleanup.o -e ${DIROUT}/log/vcf_cleanup.e --wrap="${SCRIPTS}/vcf_cleanup.sh ${username} ${SCRIPTS} ${DIRIN} ${DIROUT} ${VCFIN} ${limit_allele} ${limit_FS} ${limit_SOR} ${limit_MQ} ${limit_MQRankSum} ${limit_ReadPosRankSum} ${limit_QUAL} ${limit_QD} ${limit_GQ} ${limit_miss} ${limit_het} ${limit_GQfiltered} ${quantile_prob_above_threshold} ${quantile_prob_below_threshold} ${kept_above_threshold} ${kept_below_threshold} ${run}"

##########################################################################################################
# run diagnostic plots
# fixed values
##########################################################################################################
limit_FS=61
limit_SOR=4
limit_MQ=39
limit_MQRankSum=-12.5
limit_ReadPosRankSum=-8
limit_QUAL=200
limit_QD=20
limit_GQ=10
limit_miss=0.05
limit_het=0.01
limit_GQfiltered=0.2
kept_above_threshold="MQ_QUAL_QD~GQ~GQ"
kept_below_threshold="FS_SOR_allele~miss_het~GQfiltered"
quantile_prob_above_threshold=0.1
quantile_prob_below_threshold=0.9
run='diagnostic'
##########################################################################################################
sbatch -W -J vcf_cleanup -o ${DIROUT}/log/vcf_cleanup.o -e ${DIROUT}/log/vcf_cleanup.e --wrap="${SCRIPTS}/vcf_cleanup.sh ${username} ${SCRIPTS} ${DIRIN} ${DIROUT} ${VCFIN} ${limit_allele} ${limit_FS} ${limit_SOR} ${limit_MQ} ${limit_MQRankSum} ${limit_ReadPosRankSum} ${limit_QUAL} ${limit_QD} ${limit_GQ} ${limit_miss} ${limit_het} ${limit_GQfiltered} ${quantile_prob_above_threshold} ${quantile_prob_below_threshold} ${kept_above_threshold} ${kept_below_threshold} ${run}"

##########################################################################################################
# run filters
# fixed values
##########################################################################################################
limit_FS=61
limit_SOR=4
limit_MQ=39
limit_MQRankSum=-12.5
limit_ReadPosRankSum=-8
limit_QUAL=200
limit_QD=20
limit_GQ=10
limit_miss=0.05
limit_het=0.01
limit_GQfiltered=0.2
kept_above_threshold="MQ_QUAL_QD_GQ"
kept_below_threshold="FS_SOR_allele_miss_het_GQfiltered"
quantile_prob_above_threshold=0.1
quantile_prob_below_threshold=0.9
run='filter_all'
##########################################################################################################
sbatch -W -J vcf_cleanup -o ${DIROUT}/log/vcf_cleanup.o -e ${DIROUT}/log/vcf_cleanup.e --wrap="${SCRIPTS}/vcf_cleanup.sh ${username} ${SCRIPTS} ${DIRIN} ${DIROUT} ${VCFIN} ${limit_allele} ${limit_FS} ${limit_SOR} ${limit_MQ} ${limit_MQRankSum} ${limit_ReadPosRankSum} ${limit_QUAL} ${limit_QD} ${limit_GQ} ${limit_miss} ${limit_het} ${limit_GQfiltered} ${quantile_prob_above_threshold} ${quantile_prob_below_threshold} ${kept_above_threshold} ${kept_below_threshold} ${run}"

rm ${DIROUT}/*.gt ${DIROUT}/*.gq ${DIROUT}/geno*.txt
```
