## Supplementary material for "Complex population structure and haplotype patterns in Western Europe honey bee from sequencing a large panel of haploid drones": Scripts: vcf_cleanup.sh

```
#!/bin/bash
###### Script to clean initial vcf file ######
#Sonia Eynard 2018 - Alain Vignal 2019 - Sonia Eynard 2020

######################
#Variables
######################
username=${1}
SCRIPTS=${2}
DIRIN=${3}
DIROUT=${4}
VCFIN=${5} #vcf before all filters
limit_allele=${6} #accept up to tri-allelic
#for all filter threshold if x=-999 the value will be assigned eliminate quantile_prob*100% of the data 
limit_FS=${7} #Phred-scaled probability that there is strand bias at the site. Often 60
limit_SOR=${8} #Strand Odds Ratio, strand bas. Often 3
limit_MQ=${9} #Root mean square mapping quality over all the reads at the site. Often 40
limit_MQRankSum=${10} #u-based z-approximation from the Rank Sum Test for mapping qualities, compares the mapping qualities of the reads supporting the reference allele and the alternate allele. Often -12.5
limit_ReadPosRankSum=${11} #u-based z-approximation from the Rank Sum Test for site position within reads, compares whether the positions of the reference and alternate alleles are different within the reads. Often -8
limit_QUAL=${12} #Quality of ALT allele. The more samples have the ATL allele, the higher the QUAL score. Often QUAL=200 
limit_QD=${13} #QUAL score normalized by allele depth (AD). Often QD=20
limit_GQ=${14} #Genotype Quality. Often GQ=10
limit_miss=${15} #percent missing data allowed for a SNP. Often Miss=0.05
limit_het=${16} #percent heterozygotes allowed for a SNP. Often Het=0.01
limit_GQfiltered=${17} #limit of missing data due to GQ allowed. Often GQ_p=0.2 (remove markers for which GQ_p% of the values are below limit_GQ)
quantile_prob_above_threshold=${18} #Quantile probability of draw diagnostic plot
quantile_prob_below_threshold=${19} #Quantile probability of draw diagnostic plot
kept_above_threshold="${20}" #list of filters for which we keep markers above threshold
kept_below_threshold="${21}" #list of filters for which we keep markers below threshold
run=${22} #type of run we are doing, can be diagnostic or filter
#define possible filters 'FS','SOR','MQ','MQRankSum','ReadPosRankSum','QUAL','QD','miss','het','GQ','GQfiltered','allele'

vcf_fs=${VCFIN/.vcf.gz/_fs.vcf} #FS
vcf_sor=${VCFIN/.vcf.gz/_sor.vcf} #SOR
vcf_mq=${VCFIN/.vcf.gz/_mq.vcf} #MQ
vcf_qual=${VCFIN/.vcf.gz/_qual.vcf} #quality
vcf_qd=${VCFIN/.vcf.gz/_qd.vcf} #qd
vcf_multi=${VCFIN/.vcf.gz/_multi.vcf} #multiallele
vcf_miss=${VCFIN/.vcf.gz/_miss.vcf} #missing
vcf_het=${VCFIN/.vcf.gz/_het.vcf} #heterozygous
vcf_gq=${VCFIN/.vcf.gz/_gq.vcf} #all filters passed. No limit_GQ set to missing. SNP removed when more than percent_GQfiltered of  GQ < limit_GQ
vcf=${VCFIN/.vcf.gz/_allfilter.vcf} #all filters passed

############################################################################################
#Extract information per markers from vcf to draw diagnostic plots for vcf filtering 
############################################################################################
if [ "${run}" == "diagnostic" ] 
then 
	if [ -f ${DIROUT}/info.txt ]
	then 
		echo 'input files info already exist'
	else 
		bcftools query -f '%CHROM %POS %REF %ALT %FS %SOR %MQ %MQRankSum %ReadPosRankSum %QUAL %QD\n' ${DIRIN}/${VCFIN} > ${DIROUT}/info.txt
		head='CHROM POS REF ALT FS SOR MQ MQRankSum ReadPosRankSum QUAL QD\n'
		sed -i "1s/^/${head}/"  ${DIROUT}/info.txt
	fi
	
	if [ -f ${DIROUT}/geno.txt ]
	then 
		echo 'input files geno already exist'
		if [ -f ${DIROUT}/geno_aa.gt ]
		then 
			echo 'split geno already done'
		else
			split -l 100000 ${DIROUT}/geno.txt ${DIROUT}/geno_
			for file in ${DIROUT}/geno_*; do mv "$file" "${file}.gt"; done
		fi		
	else 
		bcftools query -f '%CHROM %POS [ %GT]\n' ${DIRIN}/${VCFIN} > ${DIROUT}/geno.txt 
		split -l 100000 ${DIROUT}/geno.txt ${DIROUT}/geno_
		for file in ${DIROUT}/geno_*; do mv "$file" "${file}.gt"; done
	fi
	
	if [ -f ${DIROUT}/gq.txt ]
	then 
		echo 'input files GQ already exist'
		if [ -f ${DIROUT}/gq_aa.gq ]
		then 
			echo 'split GQ already done'
		else
			split -l 100000 ${DIROUT}/gq.txt ${DIROUT}/gq_
			for file in ${DIROUT}/gq_*; do mv "$file" "${file}.gq"; done
		fi
		
	else 
		bcftools query -f '%CHROM %POS [ %GQ]\n' ${DIRIN}/${VCFIN}> ${DIROUT}/gq.txt
		split -l 100000 ${DIROUT}/gq.txt ${DIROUT}/gq_
		for file in ${DIROUT}/gq_*; do mv "$file" "${file}.gq"; done
	fi
	wc -l ${DIROUT}/*.txt

	IFS='~'
	read -ra kept_above <<< "${kept_above_threshold}"
	read -ra kept_below <<< "${kept_below_threshold}"
	l_array=$(echo ${#kept_above[@]})
	l_array=$((${l_array}-1))
	for ((i=0; i<=l_array;i++))
	do
		k_above=${kept_above[i]}
		k_below=${kept_below[i]}
		args_above=()
		IFS='_' read -r -a array <<< "${k_above}"
		for j in "${array[@]}"; do v="limit_${j}"; args_above+=${!v}; args_above+='_'; done
		args_below=()
		IFS='_' read -r -a array <<< "${k_below}"
		for j in "${array[@]}"; do v="limit_${j}"; args_below+=${!v}; args_below+='_'; done
		echo ${quantile_prob_above_threshold} 
		echo ${k_above} 
		echo ${args_above}
		echo ${quantile_prob_below_threshold} 
		echo ${k_below} 
		echo ${args_below}
		sbatch --mem=100G -W -J diagnostic_plot_${k_above}_${k_below} -o ${DIROUT}/log/diagnostic_plot_${k_above}_${k_below}.o -e ${DIROUT}/log/diagnostic_plot_${k_above}_${k_below}.e --wrap="Rscript ${SCRIPTS}/diagnostic.r ${DIROUT} ${quantile_prob_above_threshold} ${k_above} ${args_above} ${quantile_prob_below_threshold} ${k_below} ${args_below}"
	done
else
############################################################################################
#Adjust filters 
############################################################################################
	if [ ${limit_FS} = -999 ]; then limit_FS=$(more ${DIROUT}/FS_value.txt); fi
	if [ ${limit_SOR} = -999 ]; then limit_SOR=$(more ${DIROUT}/SOR_value.txt); fi
	if [ ${limit_MQ} = -999 ]; then limit_MQ=$(more ${DIROUT}/MQ_value.txt); fi
	if [ ${limit_MQRankSum} = -999 ]; then limit_MQRankSum=$(more ${DIROUT}/MQRankSum_value.txt); fi
	if [ ${limit_ReadPosRankSum} = -999 ]; then limit_ReadPosRankSum=$(more ${DIROUT}/ReadPosRankSum_value.txt); fi
	if [ ${limit_QUAL} = -999 ]; then limit_QUAL=$(more ${DIROUT}/QUAL_value.txt); fi
	if [ ${limit_QD} = -999 ]; then limit_QD=$(more ${DIROUT}/QD_value.txt); fi
	if [ ${limit_GQ} = -999 ]; then limit_GQ=$(more ${DIROUT}/GQ_value.txt); fi
	if [ ${limit_miss} = -999 ]; then	limit_miss=$(more ${DIROUT}/miss_value.txt); fi
	if [ ${limit_het} = -999 ]; then	limit_het=$(more ${DIROUT}/het_value.txt); fi
	if [ ${limit_GQfiltered} = -999 ]; then limit_GQfiltered=$(more ${DIROUT}/GQfiltered_value.txt); fi
############################################################################################
#Run filters
############################################################################################
	### option 1 group by group
	if [ "${run}" == "filter_sequential" ]
	then
		if [ -f ${DIROUT}/snp_kept_nb_allele.txt ]
		then 
			echo 'snp kept files already exist'
		else 
			sbatch -W --wrap="Rscript ${SCRIPTS}/filter_list.r ${DIROUT} allele ${limit_allele}"
		fi		
		#list markers pass filters
		kept_above_threshold="$(sed s/~/_/g <<<$kept_above_threshold)"
		kept_below_threshold="$(sed s/~/_/g <<<$kept_below_threshold)"
		IFS='_' read -r -a array <<< "${kept_above_threshold}"
		for j in "${array[@]}"; do v="limit_${j}"; args_above+=${!v}; args_above+='_'; done
		args_below=()
		IFS='_' read -r -a array <<< "${kept_below_threshold}"
		for j in "${array[@]}"; do v="limit_${j}"; args_below+=${!v}; args_below+='_'; done
		filters_id=="${kept_above_threshold} ${kept_below_threshold}"
		echo ${filters_id}
		cp ${DIRIN}/${VCFIN} ${DIROUT}/vcf_new.vcf
		
		if [[ ${filters_id}  == *"FS"* ]]
		then
			#filter on FS
			bcftools view -i 'FS<'${limit_FS}'' ${DIROUT}/vcf_new.vcf > ${DIROUT}/tmp.vcf && mv ${DIROUT}/tmp.vcf ${DIROUT}/vcf_new.vcf	
			cp ${DIROUT}/vcf_new.vcf ${DIROUT}/${vcf_fs}
		fi		
		if [[ ${filters_id}  == *"SOR"* ]]
		then
			#filter on SOR
			bcftools view -i 'SOR<'${limit_SOR}'' ${DIROUT}/vcf_new.vcf > ${DIROUT}/tmp.vcf && mv ${DIROUT}/tmp.vcf ${DIROUT}/vcf_new.vcf	
			cp ${DIROUT}/vcf_new.vcf ${DIROUT}/${vcf_sor}
		fi		
		if [[ ${filters_id}  == *"MQ"* ]]
		then
			#filter on MQ
			bcftools view -i 'MQ>'${limit_MQ}'' ${DIROUT}/vcf_new.vcf > ${DIROUT}/tmp.vcf && mv ${DIROUT}/tmp.vcf ${DIROUT}/vcf_new.vcf	
			cp ${DIROUT}/vcf_new.vcf ${DIROUT}/${vcf_mq}
		fi
		if [[ ${filters_id}  == *"QUAL"* ]]
		then
			#filter on QUAL 
			bcftools view -i 'QUAL>'${limit_QUAL}'' ${DIROUT}/vcf_new.vcf > ${DIROUT}/tmp.vcf && mv ${DIROUT}/tmp.vcf ${DIROUT}/vcf_new.vcf	
			cp ${DIROUT}/vcf_new.vcf ${DIROUT}/${vcf_qual}		
		fi
		if [[ ${filters_id}  == *"QD"* ]]
		then
			#filter on QD
			bcftools view -i 'QD>'${limit_QD}'' ${DIROUT}/vcf_new.vcf > ${DIROUT}/tmp.vcf && mv ${DIROUT}/tmp.vcf ${DIROUT}/vcf_new.vcf	
			cp ${DIROUT}/vcf_new.vcf ${DIROUT}/${vcf_qd}		
		fi
		if [[ ${filters_id}  == *"allele"* ]]
		then
			#filter on number of alleles
			awk '!/*/{print $0}' ${DIROUT}/vcf_new.vcf > ${DIROUT}/tmp.vcf && mv ${DIROUT}/tmp.vcf ${DIROUT}/vcf_new.vcf	
			bcftools view --max-alleles ${limit_allele} ${DIROUT}/vcf_new.vcf > ${DIROUT}/tmp.vcf && mv ${DIROUT}/tmp.vcf ${DIROUT}/vcf_new.vcf	
			cp ${DIROUT}/vcf_new.vcf ${DIROUT}/${vcf_multi}		
		fi
		if [[ ${filters_id}  == *"miss"* ]]
		then
			#filter on missing
			bcftools view -i 'F_MISSING<'${limit_miss}'' ${DIROUT}/vcf_new.vcf > ${DIROUT}/tmp.vcf && mv ${DIROUT}/tmp.vcf ${DIROUT}/vcf_new.vcf
			cp ${DIROUT}/vcf_new.vcf ${DIROUT}/${vcf_miss}		
		fi
		if [[ ${filters_id}  == *"het"* ]]
		then		
			#filter on heterozygous
			bcftools +missing2ref ${DIROUT}/vcf_new.vcf  -- -m > ${DIROUT}/tmp.vcf
			bcftools +setGT ${DIROUT}/tmp.vcf -- -t q -i 'GT="het"' -n "./." > ${DIROUT}/tmp2.vcf
			bcftools view -i 'F_MISSING<'${limit_het}'' ${DIROUT}/tmp2.vcf | bcftools query -f '%CHROM %POS \n' > ${DIROUT}/heterozygous_keep.txt
			vcftools --vcf ${DIRIN}/vcf_new.vcf  --positions ${DIROUT}/heterozygous_keep.txt --recode --recode-INFO-all --out ${DIROUT}/tmp.vcf && mv ${DIROUT}/tmp.vcf.recode.vcf ${DIROUT}/vcf_new.vcf
			cp ${DIROUT}/vcf_new.vcf ${DIROUT}/${vcf_het}		
		fi
		if [[ ${filters_id}  == *"GQfiltered"* ]]
		then		
			#filter on GQ 
			bcftools query -f '%CHROM %POS [ %GQ]\n' ${DIROUT}/vcf_new.vcf > ${DIROUT}/tmp.txt
			ncol=$(awk '{print NF;exit}' ${DIROUT}/tmp.txt)
			nind=$(($ncol-2))
			threshold=$( bc -l <<<"${limit_GQfiltered}*${nind}" )
			threshold=${threshold%.*}
			nsnp=$(wc -l < ${DIROUT}/tmp.txt)
			echo -n > ${DIROUT}/gq_keep.txt
			for i in $(seq 1 ${nsnp}) 	
			do
				line=($(sed -n "${i}p" ${DIROUT}/tmp.txt))
				line=("${line[@]:2}")
				n_gq_filt=0
				for j in $(seq 0 1 $((${limit_GQ}-1))) 	
				do
					x=$(grep -o -w ${j} <<< "${line[@]}" | wc -l)
					n_gq_filt=$((n_gq_filt+x))
				done
				if [[ ${n_gq_filt} -lt ${threshold} ]]
				then
					sed -n "${i}p" ${DIROUT}/tmp.txt | awk '{print $1" "$2}' >> ${DIROUT}/gq_keep.txt
				fi
			done
			vcftools --vcf ${DIRIN}/vcf_new.vcf --positions ${DIROUT}/gq_keep.txt --recode --recode-INFO-all --out ${DIROUT}/tmp.vcf && mv ${DIROUT}/tmp.vcf.recode.vcf ${DIROUT}/vcf_new.vcf
			cp ${DIROUT}/vcf_new.vcf ${DIROUT}/${vcf_gq}		
		fi
		
		#final
		cp ${DIROUT}/vcf_new.vcf ${DIROUT}/${vcf}
		bcftools +setGT ${DIROUT}/${vcf} -- -t q -i 'GT="het"' -n "./." > ${DIROUT}/tmp.vcf && mv ${DIROUT}/tmp.vcf ${DIROUT}/${vcf}
		rm ${DIROUT}/tmp.vcf.recode.vcf ${DIROUT}/tmp2.vcf.recode.vcf ${DIROUT}/tmp.vcf.log ${DIROUT}/tmp ${DIROUT}/tmp.vcf ${DIROUT}/tmp2.vcf ${DIROUT}/heterozygous_keep.txt ${DIROUT}/gq_keep.txt
		#Counting SNPs on the chromosomes in the various vcf files
		zgrep -v '#' ${DIRIN}/${VCFIN} | cut -f 1 | sort | uniq -c | awk 'BEGIN {OFS="\t";sum=0}{print $2, $1; sum += $1} END {print "Sum", sum}' > ${DIROUT}/countVcfInitial
		grep -v '#' ${DIROUT}/${vcf_fs} | cut -f 1 | sort | uniq -c | awk 'BEGIN {OFS="\t";sum=0}{print $2, $1; sum += $1} END {print "Sum", sum}' > ${DIROUT}/countVcfAllFilteredSNPs_fs
		grep -v '#' ${DIROUT}/${vcf_sor} | cut -f 1 | sort | uniq -c | awk 'BEGIN {OFS="\t";sum=0}{print $2, $1; sum += $1} END {print "Sum", sum}' > ${DIROUT}/countVcfAllFilteredSNPs_sor
		grep -v '#' ${DIROUT}/${vcf_mq} | cut -f 1 | sort | uniq -c | awk 'BEGIN {OFS="\t";sum=0}{print $2, $1; sum += $1} END {print "Sum", sum}' > ${DIROUT}/countVcfAllFilteredSNPs_mq
		grep -v '#' ${DIROUT}/${vcf_qual} | cut -f 1 | sort | uniq -c | awk 'BEGIN {OFS="\t";sum=0}{print $2, $1; sum += $1} END {print "Sum", sum}' > ${DIROUT}/countVcfAllFilteredSNPs_qual
		grep -v '#' ${DIROUT}/${vcf_qd} | cut -f 1 | sort | uniq -c | awk 'BEGIN {OFS="\t";sum=0}{print $2, $1; sum += $1} END {print "Sum", sum}' > ${DIROUT}/countVcfAllFilteredSNPs_qd
		grep -v '#' ${DIROUT}/${vcf_multi} | cut -f 1 | sort | uniq -c | awk 'BEGIN {OFS="\t";sum=0}{print $2, $1; sum += $1} END {print "Sum", sum}' > ${DIROUT}/countVcfAllFilteredSNPs_multi
		grep -v '#' ${DIROUT}/${vcf_miss} | cut -f 1 | sort | uniq -c | awk 'BEGIN {OFS="\t";sum=0}{print $2, $1; sum += $1} END {print "Sum", sum}' > ${DIROUT}/countVcfAllFilteredSNPs_miss
		grep -v '#' ${DIROUT}/${vcf_het} | cut -f 1 | sort | uniq -c | awk 'BEGIN {OFS="\t";sum=0}{print $2, $1; sum += $1} END {print "Sum", sum}' > ${DIROUT}/countVcfAllFilteredSNPs_het
		grep -v '#' ${DIROUT}/${vcf_gq} | cut -f 1 | sort | uniq -c | awk 'BEGIN {OFS="\t";sum=0}{print $2, $1; sum += $1} END {print "Sum", sum}' > ${DIROUT}/countVcfAllFilteredSNPsFinal_GqProportion
		grep -v '#' ${DIROUT}/${vcf} | cut -f 1 | sort | uniq -c | awk 'BEGIN {OFS="\t";sum=0}{print $2, $1; sum += $1} END {print "Sum", sum}' > ${DIROUT}/countVcfAllFilteredSNPsFinal
		sbatch -W -J count_phased_geno -o ${DIROUT}/log/count_phased_geno.o -e ${DIROUT}/log/count_phased_geno.e --wrap="python ${SCRIPTS}/count_phased_geno.py ${DIROUT} geno.txt"
		mv ${DIROUT}/count_phased_geno.txt ${DIROUT}/count_phased_geno_initial.txt
		bcftools query -f '%CHROM %POS [ %GT]\n' ${DIROUT}/${vcf} > ${DIROUT}/geno2.txt
		sbatch -W -J count_phased_geno -o ${DIROUT}/log/count_phased_geno.o -e ${DIROUT}/log/count_phased_geno.e --wrap="python ${SCRIPTS}/count_phased_geno.py ${DIROUT} geno2.txt"
		mv ${DIROUT}/count_phased_geno.txt ${DIROUT}/count_phased_geno_final.txt
		#Compressing vcfs
		bgzip -c ${DIROUT}/${vcf} > ${DIROUT}/${vcf}.gz
		bgzip -c ${DIROUT}/${vcf_fs} > ${DIROUT}/${vcf_fs}.gz
		bgzip -c ${DIROUT}/${vcf_sor} > ${DIROUT}/${vcf_sor}.gz
		bgzip -c ${DIROUT}/${vcf_mq} > ${DIROUT}/${vcf_mq}.gz
		bgzip -c ${DIROUT}/${vcf_qual} > ${DIROUT}/${vcf_qual}.gz
		bgzip -c ${DIROUT}/${vcf_qd} > ${DIROUT}/${vcf_qd}.gz
		bgzip -c ${DIROUT}/${vcf_multi} > ${DIROUT}/${vcf_multi}.gz
		bgzip -c ${DIROUT}/${vcf_miss} > ${DIROUT}/${vcf_miss}.gz
		bgzip -c ${DIROUT}/${vcf_het} > ${DIROUT}/${vcf_het}.gz
		bgzip -c ${DIROUT}/${vcf_gq} > ${DIROUT}/${vcf_gq}.gz

	#### option 2 all filter at once
	elif [ "${run}" == "filter_all" ]
	then
		if [ -f ${DIROUT}/info.txt ]
		then 
			echo 'input files info already exist'
		else 
			bcftools query -f '%CHROM %POS %REF %ALT %FS %SOR %MQ %MQRankSum %ReadPosRankSum %QUAL %QD\n' ${DIRIN}/${VCFIN} > ${DIROUT}/info.txt
			head='CHROM POS REF ALT FS SOR MQ MQRankSum ReadPosRankSum QUAL QD\n'
			sed -i "1s/^/${head}/"  ${DIROUT}/info.txt
		fi
		if [ -f ${DIROUT}/snp_kept_nb_allele.txt ]
		then 
			echo 'snp kept files already exist'
		else 
			sbatch -W --wrap="Rscript ${SCRIPTS}/filter_list.r ${DIROUT} allele ${limit_allele}"
		fi

		if [ -f ${DIROUT}/missing.txt ]
		then 
			echo 'missing % files already exist'
		else 
			if [ -f ${DIROUT}/geno.txt ]
			then 
				echo 'input files geno already exist'
				if [ -f ${DIROUT}/geno_aa.gt ]
				then 
					echo 'split geno already done'
				else
					split -l 100000 ${DIROUT}/geno.txt ${DIROUT}/geno_
					for file in ${DIROUT}/geno_*; do mv "$file" "${file}.gt"; done
				fi		
			else 
				bcftools query -f '%CHROM %POS [ %GT]\n' ${DIRIN}/${VCFIN} > ${DIROUT}/geno.txt 
				split -l 100000 ${DIROUT}/geno.txt ${DIROUT}/geno_
				for file in ${DIROUT}/geno_*; do mv "$file" "${file}.gt"; done
			fi
			limit=${limit_miss}_${limit_het}
			sbatch -W --mem=100G -W --wrap="Rscript ${SCRIPTS}/filter_list.r ${DIROUT} miss_het ${limit}"
		fi									
		if [ -f ${DIROUT}/GQfiltered.txt ]
		then 
			echo 'GQ % files already exist'
		else 
			if [ -f ${DIROUT}/gq.txt ]
			then 
				echo 'input files geno already exist'
				if [ -f ${DIROUT}/gq_aa.gq ]
				then 
					echo 'split GQ already done'
				else
					split -l 100000 ${DIROUT}/gq.txt ${DIROUT}/gq_
					for file in ${DIROUT}/gq_*; do mv "$file" "${file}.gq"; done
				fi		
			else 
				bcftools query -f '%CHROM %POS [ %GQ]\n' ${DIRIN}/${VCFIN} > ${DIROUT}/gq.txt 
				split -l 100000 ${DIROUT}/gq.txt ${DIROUT}/gq_
				for file in ${DIROUT}/gq_*; do mv "$file" "${file}.gq"; done
			fi
			limit=${limit_GQ}_${limit_GQfiltered}
			sbatch -W --mem=100G -W --wrap="Rscript ${SCRIPTS}/filter_list.r ${DIROUT} GQ_GQfiltered ${limit}"
		fi							
			
		#list markers pass filters
		kept_above_threshold="$(sed s/~/_/g <<<$kept_above_threshold)"
		kept_below_threshold="$(sed s/~/_/g <<<$kept_below_threshold)"
		IFS='_' read -r -a array <<< "${kept_above_threshold}"
		for j in "${array[@]}"; do v="limit_${j}"; args_above+=${!v}; args_above+='_'; done
		args_below=()
		IFS='_' read -r -a array <<< "${kept_below_threshold}"
		for j in "${array[@]}"; do v="limit_${j}"; args_below+=${!v}; args_below+='_'; done
		echo ${kept_above_threshold} 
		echo ${args_above}
		echo ${kept_below_threshold} 
		echo ${args_below}
		sbatch --mem=20G -W -J filter_${args_above}_${args_below} -o ${DIROUT}/log/filter_${args_above}_${args_below}.o -e ${DIROUT}/log/filter_${args_above}_${args_below}.e --wrap="Rscript ${SCRIPTS}/filter.r ${DIROUT} ${kept_above_threshold} ${args_above} ${kept_below_threshold} ${args_below}"

		#filter
		vcftools --gzvcf ${DIRIN}/${VCFIN} --positions ${DIROUT}/list_kept.txt --recode --recode-INFO-all --out ${DIROUT}/${vcf}
		mv ${DIROUT}/${vcf}.recode.vcf ${DIROUT}/${vcf}
		bcftools +setGT ${DIROUT}/${vcf} -- -t q -i 'GT="het"' -n "./." > ${DIROUT}/tmp.vcf && mv ${DIROUT}/tmp.vcf ${DIROUT}/${vcf}

		#Counting SNPs on the chromosomes in the various vcf files
		zgrep -v '#' ${DIRIN}/${VCFIN} | cut -f 1 | sort | uniq -c | awk 'BEGIN {OFS="\t";sum=0}{print $2, $1; sum += $1} END {print "Sum", sum}' > ${DIROUT}/countVcfInitial
		grep -v '#' ${DIROUT}/${vcf} | cut -f 1 | sort | uniq -c | awk 'BEGIN {OFS="\t";sum=0}{print $2, $1; sum += $1} END {print "Sum", sum}' > ${DIROUT}/countVcfAllFilteredSNPsFinal
		sbatch -W -J count_phased_geno -o ${DIROUT}/log/count_phased_geno.o -e ${DIROUT}/log/count_phased_geno.e --wrap="python ${SCRIPTS}/count_phased_geno.py ${DIROUT} geno.txt"
		mv ${DIROUT}/count_phased_geno.txt ${DIROUT}/count_phased_geno_initial.txt
		bcftools query -f '%CHROM %POS [ %GT]\n' ${DIROUT}/${vcf} > ${DIROUT}/geno2.txt
		sbatch -W -J count_phased_geno -o ${DIROUT}/log/count_phased_geno.o -e ${DIROUT}/log/count_phased_geno.e --wrap="python ${SCRIPTS}/count_phased_geno.py ${DIROUT} geno2.txt"
		mv ${DIROUT}/count_phased_geno.txt ${DIROUT}/count_phased_geno_final.txt

		#Compressing vcfs
		bgzip -c ${DIROUT}/${vcf} > ${DIROUT}/${vcf}.gz

	fi
fi
```
