## Supplementary material for "Complex population structure and haplotype patterns in Western Europe honey bee from sequencing a large panel of haploid drones": Scripts: find_crossing_overs.py

```
#!/usr/bin/env python3

#import subprocess
from subprocess import Popen, PIPE
import csv
import argparse
import pandas as pd
from collections import defaultdict

#dataDict = {}
#dataDictAgg = defaultdict(list)


def print_header(names, out_file):
    """
    Prints a header with chromosome, position and all drone samples analysed;
    for the output file with all the informative genotype vectors,
    for which the queen is heterozygote.
    """
    names_line = []
    print('chromosome position', end = " ", file = out_file)
    for line in names:
        names_line.append(line[0])
    print(" ".join(names_line), file = out_file)
    return names_line

def make_vector(genotypes):
    """
    Input: a list of genotypes from the vcf file.
    Output: a list of two lists (genotype vectors),
    each inverse of the other, the best of two will be used in the phasing stage.
    """
    genotype_vector0=[]
    genotype_vector1=[]
    for allele in genotypes:
        if allele[0] != allele[2]:
            genotype_vector0.append("2")
            genotype_vector1.append("2")
        elif allele[0] == allele[2]:
            if allele[0] == "0":
                genotype_vector0.append("0")
                genotype_vector1.append("1")
            if allele[0] == "1":
                genotype_vector0.append("1")
                genotype_vector1.append("0")
            if allele[0] == ".":
                genotype_vector0.append("9")
                genotype_vector1.append("9")
    genotype_vector=[genotype_vector0,genotype_vector1]
    return genotype_vector          #Two lists, inverse of each other

def chromosome_lengths(chrom_lengths, samples, chromosomes): #read file in the form chromosome, length, with no header. return dataframe
#    chrom_lengths = pd.read_csv('/Users/avignal/GenotoulBigWork/seqapipopOnHAV3_1/MappingLiu/HAv3_1_Chromosomes.list', sep = "\t", header = None)
    """
    Input:
    """
    data_chrom_lengths = pd.read_csv(chrom_lengths, sep = "\t", header = None)
    data_chrom_lengths.columns=['chromosome','length']
    data_chrom_lengths = data_chrom_lengths.set_index('chromosome')
    #data_chrom_lengths = pd.DataFrame(columns = ['name','chromosome','position','nb_COs'])
    data_chrom_lengths_dict = defaultdict(list)
    for s in samples:
        for c in chromosomes:
            data_chrom_lengths_dict['name'].append(s)
            data_chrom_lengths_dict['chrom'].append(c)
            data_chrom_lengths_dict['pos_start_CO'].append(data_chrom_lengths.at[c,'length'])
            data_chrom_lengths_dict['pos_end_CO'].append(data_chrom_lengths.at[c,'length'])
            data_chrom_lengths_dict['nb_COs'].append(0)
            data_chrom_lengths_dict['event'].append('end_chromosome')
            data_chrom_lengths_dict["nb_vect_prev"].append(0)
            data_chrom_lengths_dict["vector"].append(0)

    data_chrom_lengths = pd.DataFrame.from_dict(data_chrom_lengths_dict)
    return data_chrom_lengths

#def make_sub_table()


def main (input_file, output_files_version, sample_list, queen_name, non_co_size, nb_simultaneous_COs, chrom_lengths):

    out_file = open("all_vectors_v" + output_files_version, "w")
    """
    Writes a file with vectors for all the informative SNPs.
    Name: all_vectors_version
    """

# Sample list provided: print header, create list of samples and genotypes table (bcftools)

    if not sample_list == "no_sample_list":
        select_samples = Popen(['bcftools','view','--samples-file', sample_list, input_file],
                stdout=PIPE, encoding = 'UTF-8')
        genotypes = Popen(['bcftools','query', '-f', '\'%CHROM %POS[ %GT]\n\''],
                stdin=select_samples.stdout, stdout=PIPE, encoding = 'UTF-8')
        with open(sample_list) as csvfile:
            names = csv.reader(csvfile)
            names_line = print_header(names,out_file)

# No sample list provided: print header, create list of samples and genotypes table (bcftools)

    elif sample_list == "no_sample_list":
        genotypes = Popen(['bcftools', 'query', '-f', '\'%CHROM %POS[ %GT]\n\'', input_file],
                stdout=PIPE, encoding = 'UTF-8')#,
        names_list = Popen(['bcftools', 'query', '-l', input_file],
                stdout=PIPE, encoding = 'UTF-8')
        names = csv.reader(names_list.stdout)
        names_line = print_header(names,out_file)

# Read data

    data = csv.reader(genotypes.stdout, delimiter = ' ')        #Read the genotype data file produced by bcftools

# Queen in data ?
    dict_cos = defaultdict(list)                            #Dictionary of crossing-overs
    dict_first_last = defaultdict(list)                     #Dictionary of first and last informative SNPs
    if not queen_name == "no_queen_name":                   #If a genotyped queen is in the dataset
        queen_index = names_line.index(queen_name)
        first_line = "yes"
        for line in data:
            line[0] = line[0].strip("'")                    #For some reason, using bcftools with python, there are some "'" that need removing"'"
            if len(line) > 1:                               #There is an empty trailing line at the end of the data file
                chromosome, position, *genotypes = line
            elif len(line) == 1:                            #If last (empty) line reached, include the last SNP of the last chromosome
                dict_first_last['chrom'].append(chromosome_old)
                dict_first_last['pos_start_CO'].append(int(position_old))
                dict_first_last['pos_end_CO'].append(int(position_old))
                dict_first_last['nb_COs'].append(0)
                dict_first_last['event'].append('last_informative')
                dict_first_last["nb_vect_prev"].append(count_vectors)
                dict_first_last["vector"].append(vector)
            if genotypes[queen_index][0] != genotypes[queen_index][2]:      #If queen has 2 alleles (heterozygote)
                genotype_vector = make_vector(genotypes)
                if not "9" in genotype_vector[0]:                           #Eliminate SNPs with missing data
                    if first_line == "yes":                                 #Analyse first line
                        count_vectors = 1
                        print(chromosome, position, end = " ", file = out_file)
                        print("".join(genotype_vector[0]), file = out_file)
                        dict_first_last['chrom'].append(chromosome)
                        dict_first_last['pos_start_CO'].append(int(position))
                        dict_first_last['pos_end_CO'].append(int(position))
                        dict_first_last['nb_COs'].append(0)
                        dict_first_last['event'].append('first_informative')
                        dict_first_last["nb_vect_prev"].append(0)
                        dict_first_last["vector"].append("".join(genotype_vector[0]))
                        first_line = "no"
                        genotype_vector_old = genotype_vector[0]
                        chromosome_old = chromosome
                        position_old = position
                        start_co_fragment = [0] * len(names_line)
                    elif first_line == "no":                                #All other lines
                        if chromosome == chromosome_old:
                            print(chromosome, position, end = " ", file = out_file)
                            if genotype_vector[0] == genotype_vector_old:               #Which of the two genotype vectors is in phase and has no difference with the previous SNP
                                print("".join(genotype_vector[0]), file = out_file)
                                position_old = position
                                count_vectors = count_vectors + 1
                            elif genotype_vector[1] == genotype_vector_old:             #Which of the two genotype vectors is in phase and has no difference with the previous SNP
                                print("".join(genotype_vector[1]), file = out_file)
                                position_old = position
                                count_vectors = count_vectors + 1
                            else:                                                       #If none of the two vectors is identical to the previous one.

                                count0 = sum(1 for a, b in zip(genotype_vector[0], genotype_vector_old) if a != b)  #Count differences with previous for genotype vector 1
                                count1 = sum(1 for a, b in zip(genotype_vector[1], genotype_vector_old) if a != b)  #Count differences with previous for genotype vector 2
#                                print(count0,count1)
                                if count0 <= nb_simultaneous_COs:
                                    indiv_with_co = [i for i in range(len(genotype_vector[0])) if genotype_vector[0][i] != genotype_vector_old[i]]
                                    fragment_length = str(int(position) - start_co_fragment[indiv_with_co[0]])
                                    vector = "".join(genotype_vector[0])
                                elif count1 <= nb_simultaneous_COs:
                                    indiv_with_co = [i for i in range(len(genotype_vector[1])) if genotype_vector[1][i] != genotype_vector_old[i]]
                                    fragment_length = str(int(position) - start_co_fragment[indiv_with_co[0]])
                                    vector = "".join(genotype_vector[1])
                                if count0 <= nb_simultaneous_COs or count1 <= nb_simultaneous_COs:
                                    # The following will only print information on the first individual, if more than 1 CO in the interval. Just used as indicator.
                                    print(vector, names_line[indiv_with_co[0]], position_old, position, start_co_fragment[indiv_with_co[0]], min(count0,count1), fragment_length, file = out_file)
                                    start_co_fragment[indiv_with_co[0]] = int(position_old)

#                                    print(indiv_with_co)

#                                dict_cos[names_line[indiv_with_co[0]]].append(chromosome)
                                    for i in indiv_with_co:
                                        dict_cos["name"].append(names_line[i])
                                        dict_cos["chrom"].append(chromosome)
                                        dict_cos["pos_start_CO"].append(int(position_old))
                                        dict_cos["pos_end_CO"].append(int(position))
#                                        dict_cos["fragment_length"].append(fragment_length)
                                        dict_cos["nb_COs"].append(int(min(count0,count1)))
                                        dict_cos["event"].append("crossing_over")
                                        dict_cos["nb_vect_prev"].append(count_vectors)
                                        dict_cos["vector"].append(vector)
                                    position_old = position
                                    genotype_vector_old = list(vector)
                                    count_vectors = 1

#                                dict_cos[names_line[indiv_with_co[0]]][chromosome]["vector"].append(vector)
#                                dict_cos[names_line[indiv_with_co[0]]][chromosome]["length"].append(fragment_length)

                            chromosome_old = chromosome
                        elif chromosome != chromosome_old: #When start a new chromosome
#                            first_line = "yes"
                            dict_first_last['chrom'].append(chromosome_old)
                            dict_first_last['pos_start_CO'].append(int(position_old))
                            dict_first_last['pos_end_CO'].append(int(position_old))
                            dict_first_last['nb_COs'].append(0)
                            dict_first_last['event'].append('last_informative')
                            dict_first_last["nb_vect_prev"].append(count_vectors)
                            dict_first_last["vector"].append(vector)

                            count_vectors = 1
                            print(chromosome, position, end = " ", file = out_file)
                            print("".join(genotype_vector[0]), file = out_file)
                            dict_first_last['chrom'].append(chromosome)
                            dict_first_last['pos_start_CO'].append(int(position))
                            dict_first_last['pos_end_CO'].append(int(position))
                            dict_first_last['nb_COs'].append(0)
                            dict_first_last['event'].append('first_informative')
                            dict_first_last["nb_vect_prev"].append(0)
                            dict_first_last["vector"].append("".join(genotype_vector[0]))
#                        first_line = "no"
                            genotype_vector_old = genotype_vector[0]
                            chromosome_old = chromosome
                            position_old = position
                            start_co_fragment = [0] * len(names_line)


    data_table = pd.DataFrame.from_dict(dict_cos)

    data_out = pd.DataFrame(columns = ['name','chrom','event','nb_vect_prev','nb_vect_next','pos_start_CO','pos_end_CO','interval','nb_COs','pos_prev_CO','pos_next_CO'])

    samples = list(data_table['name'].unique())
    chromosomes = list(data_table['chrom'].unique())

    dict_zeros = defaultdict(list)
    for s in samples:
        for c in chromosomes:
            dict_zeros['name'].append(s)
            dict_zeros['chrom'].append(c)
            dict_zeros['pos_start_CO'].append(0)
            dict_zeros['pos_end_CO'].append(0)
            dict_zeros['nb_COs'].append(0)
            dict_zeros['event'].append('start_chromosome')
            dict_zeros["nb_vect_prev"].append(0)
            dict_zeros["vector"].append(0)

    temp_table = pd.DataFrame.from_dict(dict_zeros)
    data_table = data_table.append(temp_table)

    first_last_table = pd.DataFrame.from_dict(dict_first_last)
    for s in samples:
        temp_table = first_last_table
        temp_table['name'] = s
        data_table = data_table.append(temp_table)

    if not chrom_lengths == "none":     #If a file with lengths of chromosomes provided, create dataframe of chromosome lengths for each sample.
        data_chrom_lengths = chromosome_lengths(chrom_lengths, samples, chromosomes)
        data_table = data_table.append(data_chrom_lengths)

    data_table = data_table.sort_values(by=['name','chrom','pos_start_CO'], ascending=[True,True,True])

    for s in samples:
        for c in chromosomes:
            data_sample_chromosome = data_table[(data_table.name == s) & (data_table.chrom == c)]
            data_sample_chromosome = data_sample_chromosome.reset_index()
            data_sample_chromosome = data_sample_chromosome.drop(['index'], axis =1)
            data_sample_chromosome['pos_prev_CO'] = data_sample_chromosome['pos_end_CO'].shift(1, fill_value=0)
            data_sample_chromosome['pos_next_CO'] = data_sample_chromosome['pos_start_CO'].shift(-1, fill_value=0)
            data_sample_chromosome['nb_vect_next'] = data_sample_chromosome['nb_vect_prev'].shift(-1, fill_value=0)
            data_out = data_out.append(data_sample_chromosome, sort=True)
    data_out['interval'] = data_out['pos_end_CO'] - data_out['pos_start_CO']
    data_out['dist_min_prev_CO'] = data_out['pos_start_CO'] - data_out['pos_prev_CO']
    data_out['dist_min_next_CO'] = data_out['pos_next_CO'] - data_out['pos_end_CO']
    data_out['dist_max_prev_CO'] = data_out['pos_end_CO'] - data_out['pos_prev_CO']
    data_out['dist_max_next_CO'] = data_out['pos_next_CO'] - data_out['pos_start_CO']

    data_out.to_csv("testCOs_intervals_v" + output_files_version, sep="\t", index=False)

    out_file.close()


def parseArguments():
    '''Parse input arguments.
    IN:		None
    OUT:	parsed arguments
    '''
    parser = argparse.ArgumentParser(description='Find Crossing overs in colonies')

    parser.add_argument('-i', '--input_file', nargs='?', required=True,
                        help='input *.vcf.gz')
    parser.add_argument('-o', '--output_files_version', nargs='?', required=False, default='0', #adds a version number to the output files
                        help='output files version. Default = 0')
    parser.add_argument('-l', '--sample_list', nargs='?', required=False, default="no_sample_list",
                        help='list of samples to analyse')
    parser.add_argument('-q', '--queen_name', nargs='?', required=False, default="no_queen_name",
                        help='name of the queen, if present')
    parser.add_argument('-n', '--non_co_size', nargs='?', required=False, type=int, default=10000,
                        help='length for non-CO tracts. Default = 10000 bp')
    parser.add_argument('-c', '--nb_simultaneous_COs', nargs='?', required=False, type=int, default=1,
                        help='number of simultaneous COs allowed in an interval')
    parser.add_argument('-e', '--chrom_lengths', nargs='?', required=False, default="none",
                        help='file with chromosome sequence lengths. Default = none')

    return parser.parse_args()

if __name__ == '__main__':
    args = parseArguments()
    main(args.input_file, args.output_files_version, args.sample_list, args.queen_name, args.non_co_size, args.nb_simultaneous_COs, args.chrom_lengths)

# end of file
```
